# Dicer-to-Argonaute switch controls biogenesis of oncogenic miRNA

**DOI:** 10.1101/2021.08.30.458145

**Authors:** L. Winchester, L. van Bijsterveldt, A. Dhawan, S. Wigfield, C. Triantafyllidis, S. Haider, A. McIntyre, T.C. Humphrey, A.L. Harris, F.M. Buffa

## Abstract

miRNAs are post-transcriptional regulators of gene expression, controlling biological processes from development to pathogenesis. We asked whether the reshaped functional miRNA landscape in cancers is driven by altered transcription of its precursors, or altered biogenesis and maturation of miRNAs. Integrated analysis of genomic and transcriptomic data in 9,111 samples across 10 cancer types and healthy tissues revealed a recurrent genomic switch from DICER-dependent to non-canonical Argonaute-mediated, DICER-independent, miRNA biogenesis. Experimental validation in AGO2-amplified clinical samples and cancer cell lines confirmed that canonical miRNAs can undergo maturation in a DICER-independent manner, and that elevated Argonaute levels promote selective maturation of the oncogenic miR-106b/25 cluster as shown by the altered ratio of mature miRNA to immature pri-miRNA levels. The preferential maturation of these oncogenic miRNAs, whose processing bypasses DICER1, promotes cancer progression and predicts poor prognosis. This highlights the evolution of non-canonical AGO2-dependent oncomiR processing as a novel driver pathway in cancer.

## Introduction

MicroRNAs (miRNAs) are small noncoding RNA molecules, key to normal development [1] and disease, such as cancer [2–5]. The defining characteristic of miRNAs is their ability to rapidly post-transcriptionally regulate the expression of a large number of target genes.

The production of a functional miRNA involves several steps requiring a number of enzymes, as well as processing, stabilising, and transportation factors [6]. Briefly, primary miRNAs (pri-miRNAs) are transcribed [7] and processed by DROSHA, an RNase III enzyme, to generate a 70-nucleotide strand in a hairpin formation. This precursor miRNA (pre-miRNA) is exported to the cytoplasm by Exportin 5 (XPO5), cleaved by DICER into a 20mer [8] and loaded into the miRNA induced silencing complex (miRISC) [6]. Argonaute 2 (*AGO2*) is the central binding module of miRISC and responsible for recruiting miRNAs, thereby silencing the complementary target mRNAs [9]. Recently, an AGO2-driven, DICER-independent, maturation pathway was identified as alternative route to miRNA biogenesis [10], but its role in human biology and disease is yet unknown.

Altered miRNA levels in cancer have been described [11], and aberrant miRNA expression is frequently associated with oncogenesis and poor prognosis [12–15]. In some cases, increased abundance of specific miRNAs has been demonstrated to arise from increased transcription of the pri-miRNA. An obvious example is that of the oncogene *MYC*, which has been linked with increased transcription of oncomiR-1 miR-17-92 (reviewed in [12]), or hsa-miR-210, which is up-regulated at the transcriptional level by hypoxia and associated with poor prognosis in multiple cancer types [13]. On the other hand, it is likely that epigenetic changes, amplification or deletion of the miRNA sequence, and differential usage of alternative and parallel maturation pathways can all alter miRNA profiles and their functional effects in cancer. For example, we have previously shown that epigenetic down-regulation of *DICER1*, which is central to miRNA biogenesis, in hypoxic tumours promotes a stem cell phenotype linked with poor prognosis [15].

A systematic pan-cancer analysis of the interplay between altered pri-miRNA transcription and changes in miRNA biogenesis pathway usage is lacking, and the relative contributions of each of these steps to global miRNA dysregulation in cancer remains unclear. To address this, we integrated genomic, transcriptomic and clinical data in a comprehensive analysis of 9,111 tumour and normal tissue samples from The Cancer Genome Atlas (TCGA), the International Cancer Genome Consortium [16] (ICGC), and Metabric [17] to reveal a global landscape of dysregulation of miRNA biogenesis in cancer. Our analysis showed that not only disrupted pri-miRNA transcription, as previously thought, but also global dysregulation of miRNA biogenesis pathways by recurrent genomic amplification underlies aberrant miRNA expression in cancer. Importantly, we identified a recurrent genomic switch supporting AGO2-mediated, DICER1 independent, selective biogenesis and maturation of prognostic and oncogenic miRNAs, which we validated using a AGO2-amplified breast cancer cell lines. While this miRNA regulatory pathway has only been recently described [10]; our analysis reveals a key role for AGO2-driven, DICER1-independent miRNA biogenesis in human disease, as amplification and over-expression of genes in the AGO2-dependent pathway frequently co-occur with DICER1 mutations and under-expression, proposing it as an important new cancer driver.

## Results

### miRNA maturation and pri-miRNA transcription are both dysregulated in cancer

Our first question was whether we could detect differences in pri-miRNA and mature miRNA expression patterns in cancer. We interrogated 9,111 tumour and normal samples (Table S1) spanning 10 cancer types, for which transcriptomic (mRNA and miRNA) profiles, somatic mutations and CNA matched data were available [18–24]. We mapped, genome-wide, the mature miRNA to their cognate pri-miRNA (Supplementary File 1) and systematically assessed the association of both pri-miRNA and mature miRNA expression, with previously characterized gene expression signatures of cancer hallmarks [5], using penalised generalised linear regression with cross-validation (Figure 1 and Table S1). For each signature and cancer type we identified mature miRNA and/or pri-miRNA that showed association with the hallmark gene signatures’ expression scores (Figure 1A). We then asked which mature miRNA and/or pri-miRNA showed either positive or negative association across cancer types, more so than would be expected by chance alone (see methods and [5]). This identified mature species recurrently associated with hallmark gene signature across cancer types (‘hallmarks-associated’).

**Figure 1.**
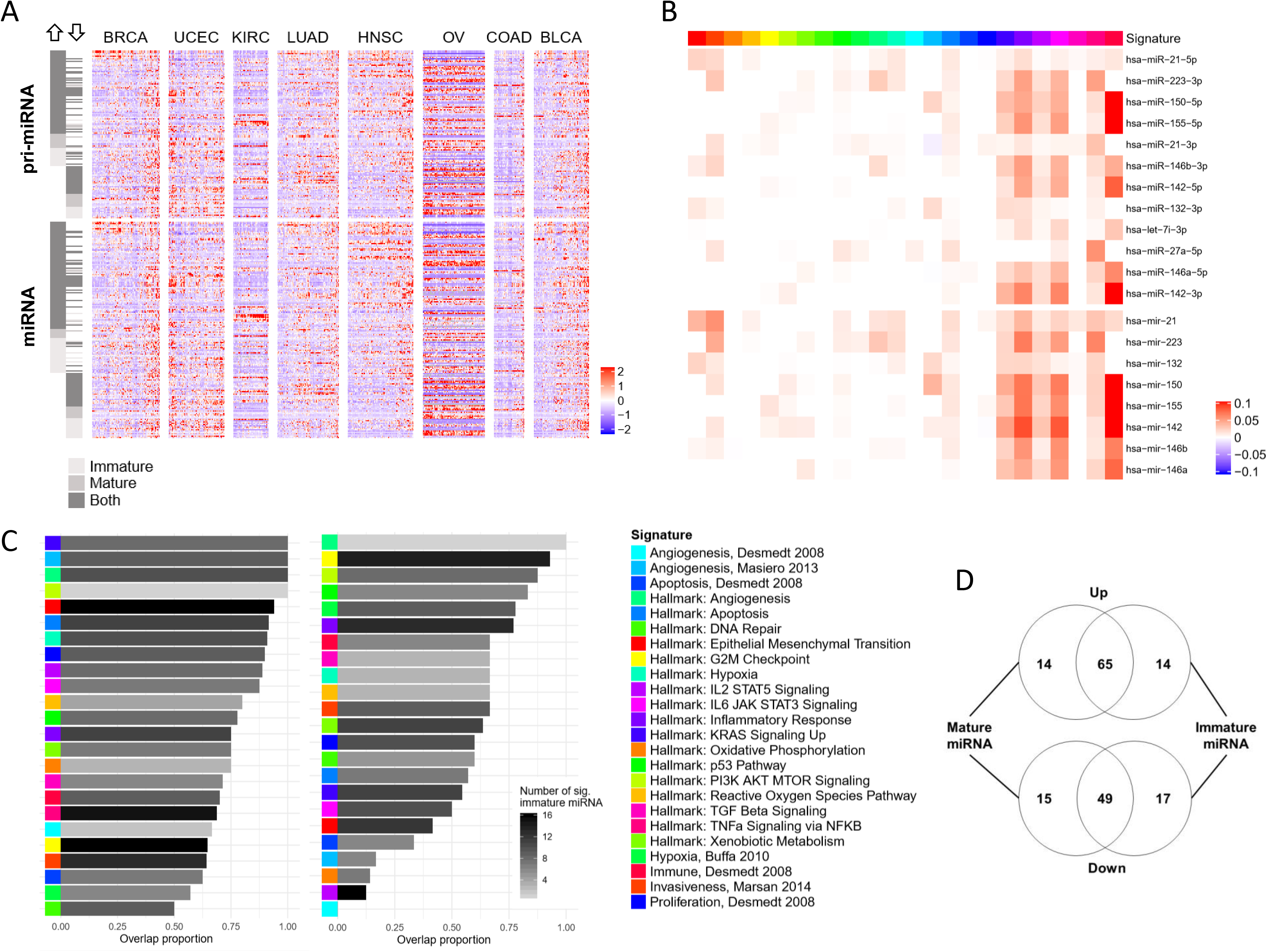
Changes in miRNA precursor transcript and abundance of mature miRNA are both associated with cancer hallmarks. **(A)** A heatmap of expression for miRNA identified as positively and negatively associated with cancer hallmark gene signatures reveals the heterogeneity in expression across mature and immature species. miRNA and pri-miRNA were identified in a pan-cancer penalized GLM regression with cross-validation. Each miRNA identified as associated is indicated by a row of the heatmap, and samples are represented as individual columns of the heatmap. For each miRNA species identified, the leftmost heatmap indicates whether it was positively associated with the hallmarks signatures (’Up’) or negatively associated with the hallmarks signatures (’Down’), and whether it was associated in mature form only, immature form only, or both mature and immature forms. miRNA expression values have been standardised to mean 0 and unit variance for ease of visualisation. **(B)** Association of top concordant miRNAs and pri-miRNAs with different hallmark signatures (Table S2), the coefficient in the GLM optimized model is shown with label. Color-coding of signatures is shown in legend and applies to all Figure 1 panels. **(C)** Concordance of miRNA and pri-miRNA in their association with hallmark signatures for miRNA positively and negatively associated with cancer hallmarks signatures (Table S2). Overlap is between miRNA and pri-miRNA is shown as proportion and the number of pre-miRNA is shown in graded grey. **(D)** Venn diagram showing overlap of miRNA species identified in independent analyses of association to hallmarks gene signatures across cancers. miRNA species positively associated with cancer hallmark gene signature score (denoted ‘Up’) are shown above, and those negatively associated with cancer hallmark gene signature scores (denoted ‘Down’) are shown below.

We observed a variable concordance between mature miRNA and pri-miRNA across hallmark-associated signatures (Figure 1B-C). Overall, our analysis identified 145 pri-miRNAs significantly associated with hallmark gene signatures, of which 79 were positively associated and 66 were negatively associated with multiple hallmarks signatures (Figure 1D, Table S2). Conversely, in the case of mature miRNAs, 79 were positively associated and 64 negatively associated with the hallmark-associated signatures (Figure 1D, Table S2). The expression for these miRNA species is depicted in Figure 1B, where both the heterogeneity of miRNA expression across cancers, and the divergent expression patterns between mature hallmark-associated miRNA and pri-miRNA are shown. In 70% of cases (65/93) where the pri-miRNA was identified as positively associated with hallmark-associated signatures, the cognate mature miRNA was also positively associated with the hallmarks, and likewise in 60% of negatively-associated species (49/81). For these miRNAs, we posit that dysregulated transcription is likely to be the key event, with the change in abundance of the mature miRNA form mostly reflecting changes in pri-miRNA levels. However, a combined change in transcription and maturation cannot be excluded without further evidence. In contrast, we showed differential association between mature and immature forms in a substantial proportion of the species identified (Figure 1D). Namely, 15% (14/93) of the positive signature-miRNA and pri-miRNA associations were non-cognate species (i.e. either mature miRNA or pri-miRNA for a specific miRNA were associated with the hallmarks signatures, but not both); likewise, 19% (15/81) and 21% (17/81) of negatively associated mature miRNA and pri-miRNA respectively were non-cognate species. This points towards an important role for dysregulated, either selectively increased or decreased, miRNA biogenesis and maturation for specific subset of miRNA species.

### Recurrent disruption by amplification of miRNA biogenesis genes

To investigate possible causes driving the observed changes in miRNA biogenesis, we looked at somatic mutation frequencies across all 43 well characterised components of the mammalian canonical miRNA biogenesis pathway [6, 7, 25] (Table S3). This highlighted a mutation level with a mean frequency of 1.05% and standard deviation (st. dev.) of 0.77% (Table S4). Colorectal (COADREAD) and endometrial (UCEC) cancers had higher mutation frequencies of 1.69% (st. dev. 1.27%) and 2.87% (st. dev. 2.32%), respectively. However, a two-sided proportion test revealed that somatic mutation frequency in miRNA biomachinery genes was not significantly different in these cancers from the expected mutation rate (Table S4). Amongst top frequently mutated genes were *DICER1*, a key miRNA processing enzyme responsible for cleavage of the pri-miRNA, and *CNOT1*, a mediator of mRNA degradation by miRNAs, with an overall pan-cancer mutation rate around 3% (2.6% and 3.53% respectively, of which 2% and 3% non-silent mutations). These results confirm our previous findings where mutations in these genes have been identified, with functional consequences on miRNA production and function in cancer [26, 27].

In contrast to somatic mutations, copy-number changes in miRNA biogenesis genes were frequent across most cancer types (Figure 2A, Table S5). Specifically, whilst deletions were rarer (but observed for some genes, for example for DICER1), copy-number amplifications (CNA) at miRNA biogenesis gene locations occurred with a pan-cancer average ranging between 7% (SMAD1) and 52% (EIF2C2), with an overall high rate of co-occurrence (Figure 2A, Table S5).

**Figure 2.**
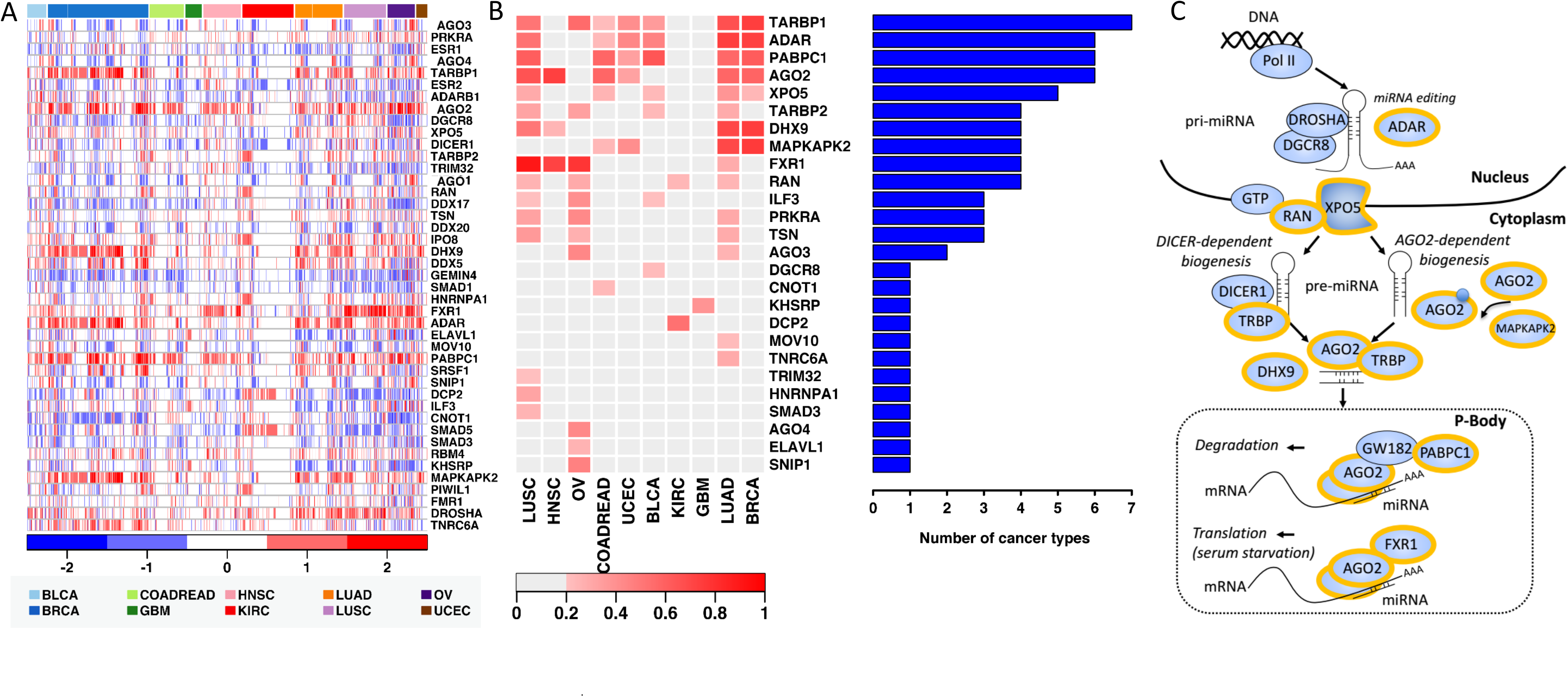
The genomic and transcriptomic landscape of 10 cancer types reveals infrequent mutation but recurring co-amplification and co-expression of genes responsible for miRNA biogenesis. **(A)** CN profiles of miRNA biogenesis genes in 10 cancer types (colour coded, see Methods). Within each cancer type, cases (columns) were visually ordered using hierarchical clustering. Genes included in Table S1; datasets used are as described in Table S2. **(B)** Functional assessment of CNAs. For each gene, the bar chart shows the number of cancer types whereby mRNA and copy-number profiles were significantly correlated (Spearman’s *ρ* > 0.3, *q* < 10^-3^). The heatmap shows the extent (fraction) of genomic aberrations for each gene, in each cancer type. The colour intensity indicates the fraction of cohort in which a given gene had a gain or amplification. Tumour types were ordered using hierarchical clustering. **(C)** A schema of the miRNA processing machinery. Main enzymes and factors are shown (Table S2). Top pan-cancer amplified genes, with correlated expression as described in Figure 2B, are shown highlighted in yellow.

### The AGO2 miRNA biogenesis and maturation pathway is functionally amplified in cancer

We asked whether we could identify gene expression changes associated with the CNAs observed, across cancer types (Figure 2B, S1A). This analysis is based on the accepted assumption that potential driver amplifications, selected for during cancer evolution, will be functional, and thus associated with changed phenotypes, namely increased gene expression. Such events could therefore constitute candidate drivers of miRNA dysregulation in cancer.

We first selected genes that were amplified/gained in at least 20% of samples in each cancer cohort, and over-expressed in tumours with respect to normal tissue (log_2_ fold-change*>* 0.5 and adjusted p<0.05, data in Table S6). Next, we performed a pan-cancer correlation analysis, which identified those CNA events significantly correlated (Spearman’s *ρ*> 0.3, *q*< 10^-3^) with over-expression of genes at the genomic location (Figure 2B, Table S7-8). This revealed that nearly a quarter (10/43) of miRNA biogenesis genes were frequently amplified, with amplification significantly correlated with over-expression in at least 4 out of 10 cancer types (Figure 2B). These genes all have key roles in miRNA biogenesis (Figure 2C). Importantly, four of the genes identified above were found to have a high concordance rate between over-expression and gene amplification in at least 6 out of 10 cancer types (Figure 2B). This included *AGO2,* a key component of miRISC; *PABPC1*, shown to interact with *AGO2* and capable of determining miRNA inhibition efficiency; and *ADAR*, responsible for regulating miRISC loading and miRNA target binding via RNA editing [28, 29]. These results highlight a potential driver role for the AGO2 maturation pathway.

Importantly, this analysis revealed some interesting differences between cancer types, with glioblastoma (GBM) and kidney cancer (KIRC) showing the lowest occurrence of CNA in these genes (Figure 2B), and lung (both LUSC and LUAD) and breast cancer (BRCA) showing the highest occurrence of candidate driver CNA events (Figure 2B, Table S7-8).

### Amplification of AGO2 across cancer cohorts is associated with DICER1 deletion

Breast cancer was the cancer type with most frequent *AGO2* CNA, most consistent co-amplification of miRNA biogenesis genes, with the highest frequency of concomitant over-expression (Figure 2A-B). Thus, we proceeded to investigate the genomic and transcriptomic patterns in breast cancer in greater detail, including the potential clinical consequences. To achieve this, we interrogated three independent well-annotated clinical datasets where matched transcriptomic (coding and non-coding) and genomic CN data were available. These were the TCGA BRCA dataset (Figure 2A), the Metabric dataset (N=1293 complete cases) and our own, smaller (N=180 complete cases), previously published dataset [14, 17, 21].

Firstly, we assessed if the genomic landscape observed in the TCGA BRCA samples (Figure 2) could be cross validated in the Metabric dataset (Figure 3 and S1B). Our analysis confirmed the existence of different gene clusters when considering CNA status in isolation (Figure 3A-B). In both the TCGA and Metabric datasets, this was mostly reflecting common or near-genomic locations. For example, AGO family members 1, 3 and 4, located on chromosome 1, shared a similar CNA correlation profile; whereas *AGO2*, located on chromosome 8, showed a markedly different profile (Figure 3B). Interestingly, *PABPC1* and *AGO2* CNAs were highly correlated despite their genomic locations not being immediately proximal, the first being on 8q22 and the second on 8q24 amplicons. Furthermore, a cluster of patients with no amplification of biogenesis genes (Figure 3A) showed an overall genome-wide lower amplification frequency. Of note, this cluster contains 31.3% of the normal Pam50 subtype.

**Figure 3.**
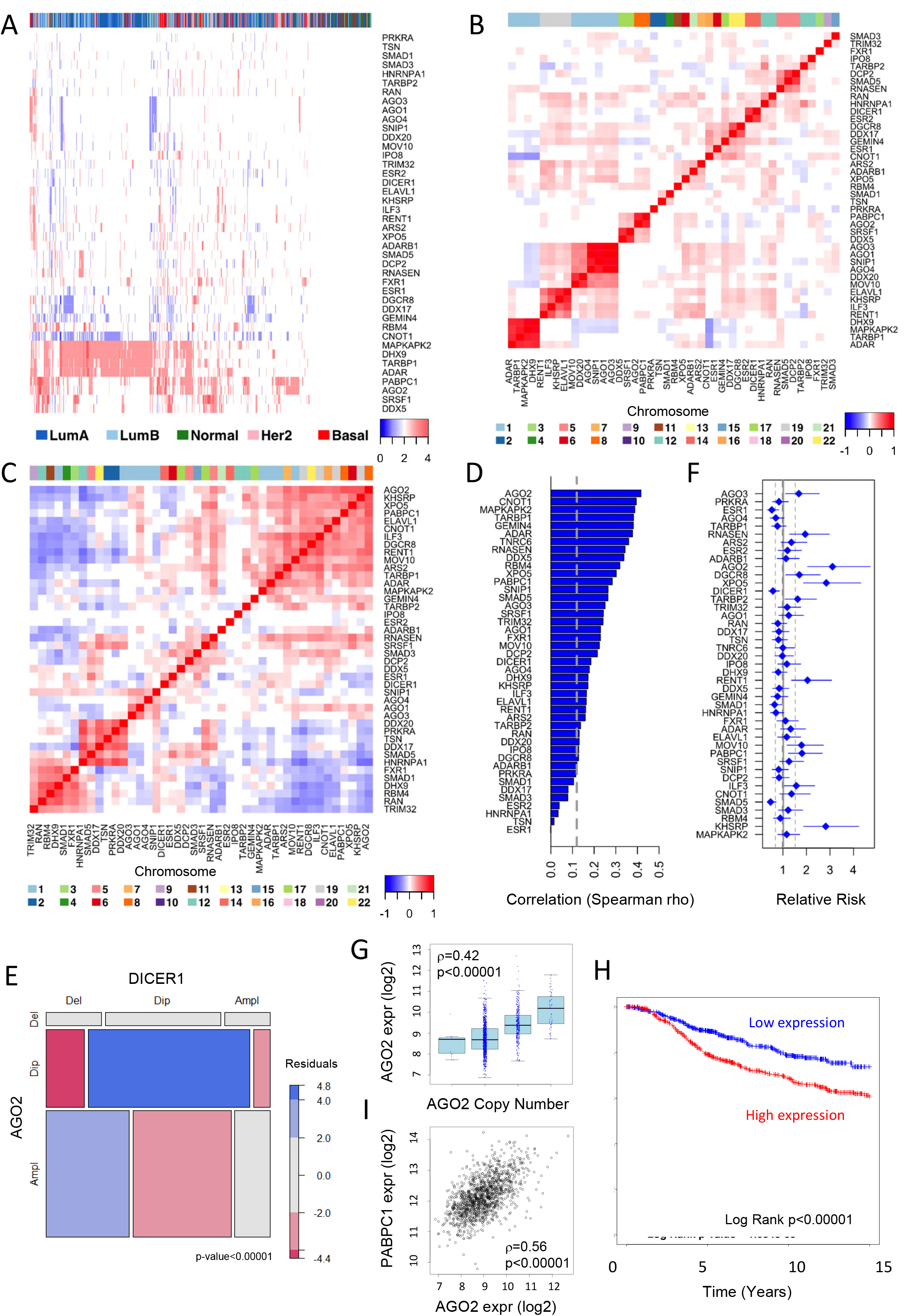
*AGO2* and *PABPC1* frequent amplification and over-expression is associated with poor prognosis in breast cancer. **(A)** CN profiles for 1,991 cases in the Metabric cohort are shown. Cases (columns) and genes (rows) were visually ordered using hierarchical clustering. Genes included are as in Table S2. Mapping to PAM50 classification is show colour coded at the top. **(B)** Auto-correlation of the biogenesis genes CN profiles is shown. CN levels were compared across genes and correlation (Spearman) estimated. Genes are visually clustered using hierarchical clustering. Colour coded chromosomal location is displayed at the top. **(C)** Autocorrelation of biogenesis genes expression profiles is shown. Expression levels were compared across genes and correlation (Spearman) estimated. Genes are visually clustered using hierarchical clustering. Colour coded chromosomal location is displayed at the top. **(D)** Correlation (Spearman rho) between CN and mRNA levels. Dotted line represents the average correlation obtained when resampling (x1000) random sets of 43 genes across the genome. Genes are ranked from highest to lowest correlation to show strongest amplification effect. **(E)** Mosaic plot of copy number aberrations is shown for AGO2 and DICER1. TCGA BRCA data, GISTIC calls (Del=Deep or Shallow Deletions, Dip=Diploids, Ampl=Gain or Amplification) Pearson Residuals are colour coded (negative:below expected, positive: greater than expected. **(F)** Survival Analysis of miRNA biogenesis genes. Cancer-specific overall survival was considered in a Cox survival analysis including the normalize expression of each gene as continuous covariate (see methods). Relative Risk (Hazard Ratio of the univariate Cox model) is shown with 95% confidence intervals. Thresholds plotted in grey show our control; namely, the mean and 95% confidence intervals of randomly genome-wide sampled genes (x1000). **(G)** Box plot of *AGO2* mRNA expression distribution at each CN value. Spearman correlation between expression and CN levels is shown. **(H)** Survival analysis of *AGO2* mRNA expression levels. Gene expression was separated by median into low and high equally sized groups. Cox analysis results are shown (HR = Hazard Ratio). Number of samples shown in associated Figure S1. **(I)** *AGO2* and *PABPC1* mRNA expression levels. Spearman correlation is shown.

On the other hand, the mRNA level-correlation analysis showed similarities in expression patterns across cancers within genes of the same family, such as members of the SMAD family; and interestingly a highly correlated cluster including several members of the *AGO2*-mediated biogenesis pathway as well as components of miRISC, including *AGO2*, *PABPC1*, and *MAPKAPK2* (Figure 3C).

When looking at CNA in combination with gene expression, only one cluster showed both correlated expression (Figure 3C) and CNA-expression correlation (Figure 3D), confirming the subset of frequently amplified and co-expressed genes identified in the analysis of TCGA BRCA samples. Namely, *TARBP1*, *ADAR, PABPC1, AGO2* and *MAPKAPK2* were co-expressed (Figure 3C), and amplification of their genomic location was strongly associated with an increase in their expression (Figure 3D, Table S9). This analysis also identified *DGCR8*, *CNOT1* and *GEMIN4* as deleted, with significant correlation between copy number (CN) and reduced expression levels (Table S10).

As a strong correlation between DNA and mRNA levels is widely recognised as an indicator of driver CNA, the next stage of the analysis focused on miRNA biogenesis genes where this was detected (Figure 3D). Importantly, a resampling analysis considering randomly selected (x1000 times) sets of 43 genes across the genome (same number as our gene set), showed that for the majority of the miRNA biogenesis genes, the correlation between amplification and expression levels was significantly greater than expected by chance alone (mean Spearman rho = 0.12 across these genes, p=1.5e-05).

Interestingly, we could observe a clear inverse correlation between AGO2 and DICER1 CNA (Figure 3B), and mRNA expression levels (Figure 3C). When we further evaluated the relationship between these two genes, we could observe that more AGO2-amplified cases than expected by chance also harboured a DICER1 deletion, while the opposite was not true, namely there was not an enrichment in DICER1 amplifications in AGO2 deleted cases (Figure 3E). Of note, DICER1 expression levels are correlated with its CNA, showing decreased expression levels in samples that harboured shallow and deep deletions in the DICER1 locus (Figure 3D, and S1C). This confirms the in these samples the DICER1 pathway is compromised. Furthermore, the expression of AGO2 was increased in cases with DICER1 shallow and deep deletion, when compared to cases with no deletions (Figure S1D). These results, taken together, suggest a switch from a DICER1 dependent to AGO2-dependent, DICER1 independent, miRNA processing occurs in tumours.

### Amplification of the AGO2 biogenesis pathway is associated with poor prognosis

Next, we asked if CNA and over-expression of frequently amplified miRNA biogenesis genes was significantly associated with clinical outcome. We used the Metabric dataset, as complete annotated clinical data and long-term follow-up is available for 1293 cases. Over-expression of 11 out 43 of the biogenesis genes was significantly associated with decreased survival (Figure 3E); whilst for smaller proportion of these genes, 4/43, over-expression was associated with increased survival (Figure 3E). Importantly, amplification events in the *ADARB1*, *DDX5*, *PABPC1* and *AGO2* cluster were concordantly associated with poor outcome (Table S9). Conversely, genes associated with good prognosis included *DICER1*, supporting our previous findings [15]. Of note, significance was maintained after resampling on the whole genome (as described in previous paragraph), which is a more conservative approach to assess significance (Figure 3E).

Next, using all three independent breast cancer cohorts, we selected highly amplified and concordantly over-expressed genes, asking if both CNA and over-expression significantly associated with prognosis across all the three cohorts. This revealed *AGO2* and *PABPC1* as top candidate drivers (Figure 3, Table S9), with both AGO2 amplification and over-expression associated with low overall survival across all cohorts (Figure 3F-H, Table S9). Of note, there was a significant increase in *AGO2* mRNA expression in high Estrogen Receptor (ER) status breast cancers when compared to low ER-status breast cancers (p-value < 2.2×10^-16^).

*PABPC1* is located at chromosomal position 8q22, which is near but not proximal to *AGO2* location, 8q24. The strong correlation between this gene pair’s CN profiles (rho = 0.90, p-value < 2.2×10^-16^) (Figure 2B) however suggests that CNA of these two genes is co-occurring or co-selected for. Importantly, in our TCGA pan-cancer analysis, these two genes were found to be co-amplified in 8/10 cancer types (Figure 2A), with Fisher exact test P<0.01 in all cancer types but LUSC and GBM. Additionally, we observed a strong correlation between the mRNA levels of these genes irrespective of amplification (rho = 0.57, p-value < 2.2×10^-16^, Figure 3I). Accordingly, increase in either *PABPC1* CNA or expression was also associated with poor prognosis (Figure S2).

These results confirm a key role and suggest a synergetic action between *AGO2*, central component of the miRISC [25] and responsible for miRNA loading, and *PABPC1*, polyA binding protein regulating the extent of inhibition that the miRISC achieves on target genes [30, 31].

### AGO2 amplification is associated with over-expression of oncogenic miRNAs

Given *AGO2*’s role in miRNA maturation, it is expected that changes in its expression would affect the abundance of several miRNAs. Interestingly, a prognostic miRNA expression profile previously identified by our group and validated by independent studies [14], showed a striking association with *AGO2* expression. Specifically, all miRNAs associated with poor prognosis, but one, had significantly increased expression in *AGO2* amplified samples. In contrast, the expression levels of all miRNAs associated with good prognosis were decreased in these samples (Table S11).

To investigate this further, we compared expression of *AGO2* in clinical samples with genome-wide miRNA sequencing data and asked whether any association of AGO2 with miRNA levels could be detected robustly across independent breast cancer datasets. Of note, miRNA expression in these datasets has been measured using different technologies (RNAseq in TCGA, and two different array platforms; see Methods for further details), which enabled us to identify robust associations. Expression of several miRNAs correlated with *AGO2* expression across datasets (Figure 4A, Table S12), including multiple members of the oncomiR-1 family. This family is composed of three paralogue clusters: miR-17/92, miR-106a/363 and miR-106b/25. In our analysis miR-18a, 19b, and 20a, members of the miR-17/92 cluster, and miR-93 and miR-106b, members of the miR-106b/25 cluster, were strongly correlated with AGO2 amplification and AGO2 transcript levels.

**Figure 4.**
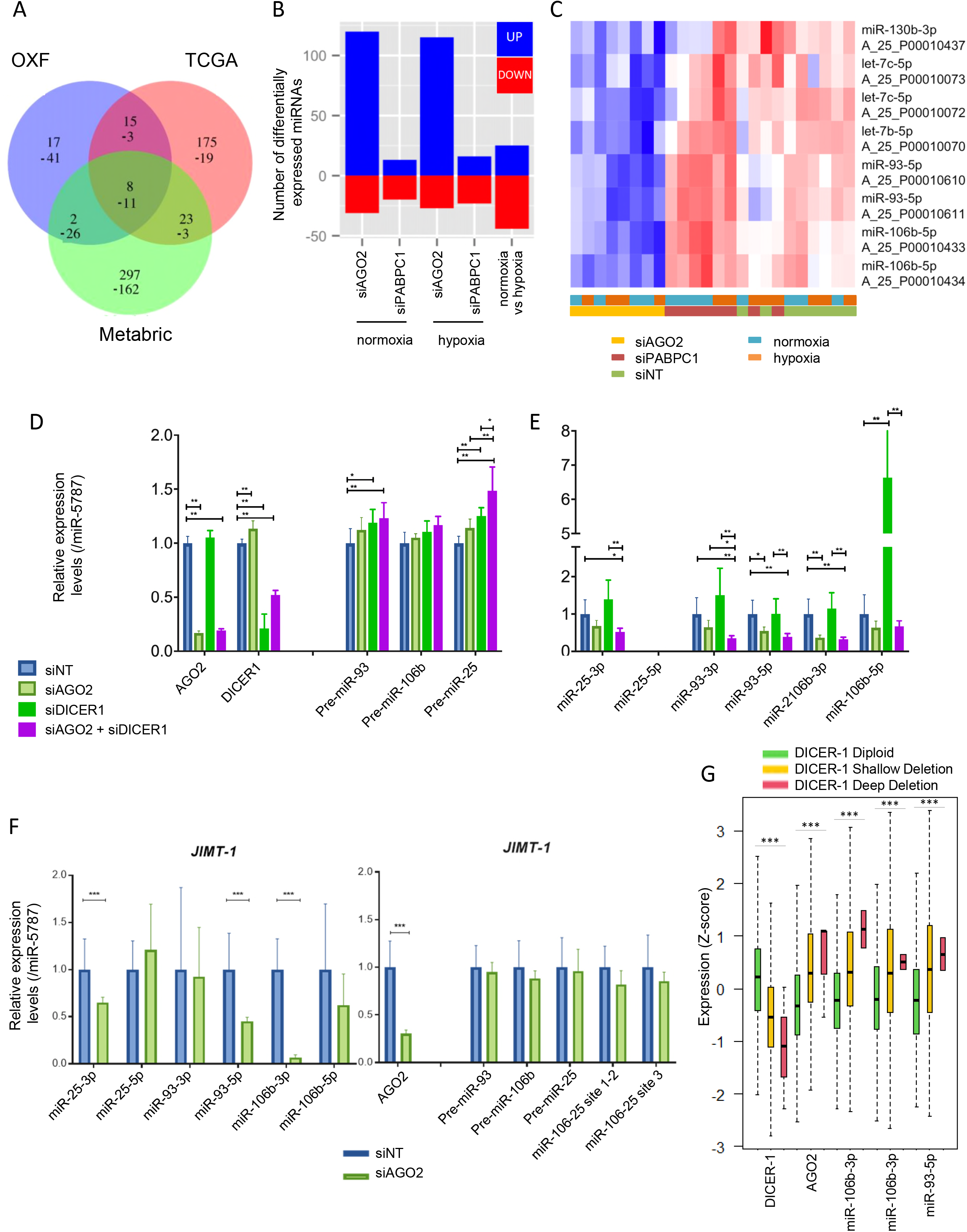
AGO2-dependent, DICER1 independent, maturation in amplified AGO2 human samples and HCC1806 cell lines. **(A)** Overlap between the *AGO2* miRNA profiles derived in the three cohorts using Spearman correlation and Benjamini & Hochberg correction (FDR<0.5). The number of common miRNAs between the three analyses is shown, minus sign “-” indicates inverse correlation with *AGO2* expression (Tables S13 and S14 for details and list of miRNAs). **(B)** *AGO2* or *PABPC1* knockdown in HCC1806. Differentially expressed miRNAs with respect to control are shown. Number of differentially regulated miRNAs are on the vertical axis. Number of miRNAs regulated under hypoxia (1% O2, see Methods) with respect to normoxia are shown. “Up” (in blue) are miRNAs with higher expression levels in the control conditions (i.e. candidate regulated by *AGO2*, PABC1 or hypoxia), “Down” (red) are miRNAs with lower expression in the control condition (i.e. candidate repressed by *AGO2*, PABC1 or hypoxia). Complete analysis in Tables S15-S17. **(C)** Expression of miRNAs correlated with *AGO2* expression in the cancer samples and confirmed regulated in cell lines after *AGO2* knockdown. Rows represent miRNAs (IDs shown) and columns represent triplicate repeats in each experimental condition, namely *AGO2* and *PABPC1* knockdown, and hypoxia 1%O2 (see Methods). When multiple probesets resulted significant, expression for both probesets is shown. miRNA expression profiles were logged base 2 and *z*-transformed, range shown from -3 log-fold changes (dark blue) to 3 log-fold changes (bright red). Hierarchical clustering is used to visually order the samples. **(D)** Real Time PCR quantification of miR-106b/25 cluster precursors normalised to Actin B in HCC1806 transfected with siDICER1 and/or siAGO2. **(E)** TaqMan based quantification of mature miRNAs miR-25-3p, miR-25-5p, miR-93-3p, miR-93-5p, miR-106b-3p and miR-106b-5p normalised to mitochondrial miR-5787 in HCC1806 transfected with siDICER1 and/or siAGO2. **(F)** TaqMan based quantification of mature miRNAs miR-25-3p, miR-25-5p, miR-93-3p, miR-93-5p, miR-106b-3p and miR-106b-5p normalised to mitochondrial miR-5787 and real-time PCR quantification of miR-106b/25 cluster precursors (normalised to Actin B) in JIMT-1 cells (harbouring a DICER1 deep deletion) transfected with siNT or siAGO2. (E) Expression of DICER1, AGO2, miR-106-3p and -5p, and miR-93-5p is shown in DICER1 diploids, shallow deleted and deep deleted cases from the TCGA BRCA dataset (z-score standardised). ***= p<0.00000001, ANOVA linear model test.

Interestingly, similar analysis considering *PABPC1* expression or amplification did not reveal consistently associated miRNA levels in any of the three cohorts (Figure S3A). This supports the known role of *PABPC1*, which is not involved in the production of mature miRNAs, but in the regulation of the extent of target inhibition by the miRNA [30, 31]. Therefore, this serves as an important control as we would not expect *PABPC1* amplification or over-expression to have a global effect in miRNA abundance.

Of the oncomiR-1 paralogue clusters associated with AGO2 amplification, the miR-17/92 cluster has been shown to be regulated directly by *MYC* [11]. Given *MYC* and *AGO2* are on nearby locations on amplicon 8q24, we asked whether *MYC* could be a confounder. Analysis including only *MYC* neutral cases, showed that the abundance of miRNAs of the miR-17/92 oncomiR-1 cluster was not significantly correlated with *AGO2* CNA or expression. Although this could be due to a smaller cohort size, it agrees with the evidence that *MYC* amplification dominates regulation of these specific miRNA.

On the other hand, we identified other miRNA, including oncomiR-1 members, strongly correlated with *AGO2* expression independently of *MYC* amplification across all datasets, notwithstanding the smaller cohort size (Figure S3B, Table S13). Specifically, in *MYC* copy neutral cases AGO2 expression was positively correlated with miR-93 expression, member of the miR-106b/25 oncomiR-1 cluster, and regulator of important cancer suppressor genes [5, 32]; positively correlated with miR-150 [33], linked with breast cancer prognosis in previous studies [14]; and negatively correlated with miR-10a, whose expression has been linked with reduced proliferation [34]. Importantly, miR-93 and -10a were also predictive of prognosis. Specifically, miR-10a abundance was associated with good prognosis (Hazard Ratio = 0.57, p=0.0067); and miR-93 abundance was associated with poor prognosis (Hazard Ratio = 2.35, p<0.0001) (Figure S3C-D), as expected from the patterns of association with AGO2 expression.

*AGO2*-dependent DICER1-independent maturation selects for the oncogenic miR-106b/25 cluster in both cancer samples and cell lines We asked to what extent *AGO2* could influence overall miRNA abundance, and abundance of specific miRNA species. To investigate this *in vitro*, we chose the HCC1806 breast cancer cell line as a model system, as it harbours amplification in AGO2 and PABPC1 and displays associated over-expression at both mRNA and protein levels (Figure S4A-C). As such, we confirmed that knockdown of AGO2 did not affect the survival of HCC1806 cells as measured by colony formation (Figure S4D-E). Comparison of miRNA expression between the *AGO2* siRNA treated and control HCC1806 cells (methods, Suppl. File 4), confirmed that there was a striking effect on miRNA abundance, with a large number of miRNAs (170 probesets, targeting 114 miRNAs) differentially expressed between *AGO2* knock down and non-targeting control treated HCC1806 cells (Figure 4B, Table S14). Of these differentially expressed miRNAs, 131 probesets (targeting 80 miRNAs) were downregulated in the *AGO2* knock down cells (with respect to the control), and 39 probesets (targeting 34 miRNAs) were upregulated in the *AGO2* knock down cells.

In agreement with our analysis of clinical cohorts (Figures 4A, S3A), *AGO2* knock down in HCC1806 revealed a significantly greater effect on expression of mature miRNAs than *PABPC1* knock down in the same cells under the same conditions (Figure 4B, Table S15), where only 22 probesets, targeting 21 miRNAs, were significantly up- or downregulated. Of these, 7 probesets, targeting 6 miRNAs, were upregulated in *PABPC1* silenced cells and 15 probesets/miRNAs, were downregulated. However, only 17 miRNAs were specific to the *PABPC1* silencing, as the others (miR-7-1, miR-4520, miR-320c and e) were downregulated following either *AGO2* or *PABPC1* silencing (Figure S5). Furthermore, amongst miRNAs identified in the *AGO2* knockdown, we were able to validate several of those identified in our analyses of the clinical samples, including both miR-93 and miR-106, members of oncomiR-1 cluster miR-106b/25 (Figure 4C; Tables S14-15).

To understand if the increased abundance in these miRNAs in AGO2 amplified samples was likely caused by changed transcription or processing, we considered changes in miRNA mature levels with respect to changes in the level of miRNA precursor transcript. This showed that the miRNA/precursor ratios for members of the miR-106b/25 cluster were significantly positively correlated (p < 10^-6^) with AGO2 amplification (Table S16), supporting the hypothesis of increased maturation for these miRNAs in samples harbouring AGO2 amplification, rather than increased transcription.

To further validate these results *in vitro*, we quantified the levels of miR-25, 93 and 106b precursors, and their cognate mature miRNAs in siAGO2-treated HCC1806 cells using qRT-PCR quantification. The forward and reverse primers were designed to bind within the hairpin sequence of the precursor miRNA (methods, Suppl. File 4). Knockdown of AGO2 caused a significant downregulation of mature miRNAs miR-93-5p (p=0.0282) and miR-106b-3p (p=<0.0001) (Figure 4E), even though levels of the miR-93/106b precursor transcripts remained unchanged in AGO2-depleted cells (Figure 4D). Importantly, expression of the host gene MCM7, with the TSS 7867 bp upstream of miR-106b, and its transcription factor E2F1 also remained unchanged (Figure S6A).

Knockdown of DICER1, in contrast, resulted in increased levels of miR-93/106b precursor transcripts and their corresponding mature miRNAs, in particular miR-106b-5p (Figure 4E), suggesting that DICER1 is dispensable for miRNA maturation in HCC1806 cells. Interestingly, E2F1, which was previously shown to be a target gene of miR-106b-5p [35], was also significantly downregulated in DICER1-knockdown HCC1806 cells (p=<0.0001, Figure S5C). Decreased levels of mature miRNAs miR-25-3p, miR-93-3-, and miR-106b-3p following AGO2 depletion were confirmed in the epithelial breast cancer cell line, JIMT-1, which harbours a DICER1 deep deletion (Figure 4F), further showing that maturation of these miRNAs is DICER1-independent. These results agree with the genomic and transcriptomic switch observed in the clinical samples between AGO2 and DICER1 (Figure 3E, and SupplS1C-E). Additionally, we could verify that in the tumours harbouring DICER1 shallow and also deep deletions, the expression of oncomiR-1 cluster members miR-93-5p, miR-106b-3p and -5p is significantly increased, with concomitant AGO2 increase, with respect to the DICER1 diploids cases (Figure 4G).

Taken together these results show that *AGO2* has a large global impact on miRNA abundance levels, including the oncomiR-1 cluster, and this is independent from DICER1. Importantly, miRNA regulation has been often reported to differ between cell lines and clinical samples [2], thus the agreement observed here between cancer cell lines and clinical samples provides solid evidence that canonical miRNAs can undergo biogenesis in a DICER-independent manner and that elevated Argonaute levels promote selective maturation of the oncogenic miR-106b/25 cluster as shown by the altered ratio of mature miRNA to immature pri-miRNA levels.

### AGO2 amplification is associated with TP53 mutation in breast cancer

Given the frequency of *AGO2 *CNA, we sought to identify novel methods to target the selective AGO2-dependent maturation of oncogenic miRNAs for therapeutics purposes. As there are currently no FDA-approved Argonaute-2 inhibitors, indirect methods of targeting AGO2 CN amplification, such as through synthetic lethality (SL) approaches, might be better suited[36].

First, we exploited two comprehensive databases where SL pairs from a multitude of studies have been submitted, the SynLethDB database [37] and the Synthetic Lethality Knowledge Graph [38]. These databases revealed two potential AGO2 SL interaction partners: *TP53* and *BRCA1*. To confirm SL interactions with *AGO2* CNA in cancer types where this event is frequent, we interrogated both the TCGA BRCA cancer and the Metabric databases to infer SL interactions based on the frequency at which genes were altered (either through somatic mutation or predicted deleterious mutation) in a (significantly) mutually exclusive manner to *AGO2* CNA [39]. In both clinical datasets, we found that TP53 mutation was more likely to co-occur with AGO2 amplification than non-amplification (Table S17). Specifically, we detected a significantly higher number of samples harbouring both *AGO2* CNA and *TP53* mutation than expected by random distribution (Table S17). In the TCGA breast cancer cohort, the frequency of *TP53* mutation was 45% in cases harbouring AGO2 CNA, but 14% in cases without AGO2 CNA. If only *TP53* deleterious mutations were considered, likely resulting in p53 loss-of-function, the frequency was 29% in cases harbouring AGO2 CNA, but only 8% in cases without AGO2 CNA. Similarly, in the Metabric cohort, where different assays are used to measure CN changes and mutations, the frequency of *TP53* mutation was 47% in cases harbouring AGO2 CNA, to 25% in cases without AGO2 CNA.

In addition to this, we identified PIK3CA as a potential AGO2 dosage SL partner, as the frequency of *PIK3CA* mutation was lower in AGO2 CNA cases than in non-amplified cases (Table S17). In the TCGA breast cancer cohort, the frequency of *PIK3CA* mutation was 28% in cases harbouring AGO2 CNA, but 48% in non-amplified cases. Similarly, in the Metabric cohort, the frequency of *PIK3CA* mutation was 37% in cases harbouring AGO2 CNA, but 47% in non-amplified cases. As the majority of missense mutations in *PIK3CA* are predicted to increase its kinase activity, these associations were not found when only *PIK3CA* deleterious mutations were considered.

Hence, we predict that simultaneous activation of the PIK3CA pathway together with AGO2 CNA in breast cancer could constitute a lethal combination, as these events co-occur less frequently.

As AGO2 is often co-amplified with cMYC, we repeated the analysis in cMYC neutral cases and obtained very similar results. Specifically, in the cMYC neutral TCGA breast cancer cohort, the frequency of *TP53* mutation was 44% in cases harbouring AGO2 CNA and 13% in cases without AGO2 CNA. If only *TP53* deleterious mutations were considered, the frequency was found to be 33% in cases harbouring AGO2 CNA, but only 7% in cases with no AGO2 CNA. In the cMYC neutral Metabric cohort, the frequency of *TP53* mutation decreased from 45% in cases harbouring AGO2 CNA, to 21% in cases with no AGO2 CNA.

### PABPC1, co-amplified with AGO2 and regulator of miRNA target inhibition, is associated with a pro-survival glycolytic shift

Neither analysis of clinical samples nor experiments in HCC1806 cell lines found *PABPC1* associated with miRNA maturation. This is in line with the role of *PABPC1* as regulator of miRNA target inhibition rather than miRNA maturation [30, 31]. If this hypothesis is correct, we would still expect to see differential regulation of miRNA targets genes in samples with amplified or over-expressed *PABPC1*.

Analysis in clinical samples identified a specific gene expression profile strongly associated with PABPC1 expression (Table S18), which included other genes of the Poly(A) Binding Protein family, and genes involved in a broad range of pathway ranging from RNA transport, glycolysis, Wnt signalling, p53, tight junction, and immune related pathways (Table S19). We thus asked if we could observe significant downregulation of some of these pathways when *PABPC1* was silenced in HCC1806 cells. This confirmed mRNA surveillance, tight junction, and glycolysis amongst the top enriched pathways in both cell lines and clinical samples expressing high PABPC1 levels (Figure 5A-B, Tables S20-21).

**Figure 5.**
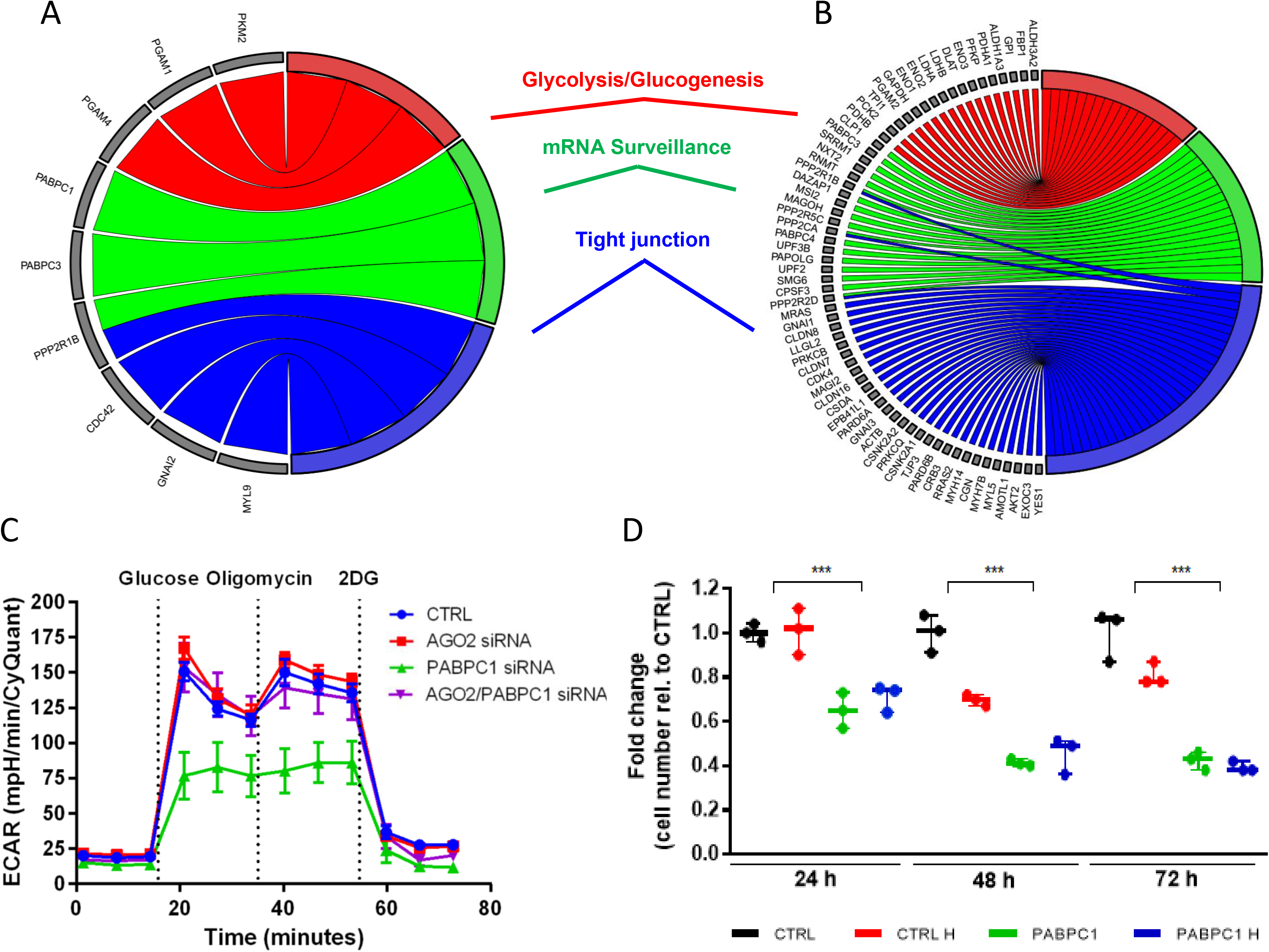
AGO2 co-amplified *PABPC1* associated with pro-survival glycolytic shift, in cancer cells and human samples. **(A)** KEGG pathway analysis for the mRNA *PABPC1* expression profile in cancer samples. Genes for which expression levels significantly correlated (Spearman, Bonferroni adjusted p<0.05, Table S19) with expression levels of *PABPC1* in the Metabric cohort were considered in a Genecodis3 KEGG pathway analysis (see Methods). Enriched pathway (FDR<0.05) which could be validated in the *PABPC1* knockdown experiments are shown. Complete list in Table S19. **(B)** KEGG pathway analysis for the mRNA *PABPC1* expression profile in HCC1806 cell lines. Genes differentially expressed in *PABPC1* knockdown with respect to control cells (Table S20) were considered in a Genecodis3 KEGG pathway analysis (see Methods). Enriched pathway (FDR<0.05) which were in common with the clinical data analysis are shown. List of all significant pathways is shown in Table S23. **(C)** Effect of *PABPC1* and *AGO2* knockdown on the rate of extracellular acidification (ECAR). ECAR (11mM Glucose, 2mM Glutamine) was measured using the Seahorse XF96e Bio-analyser in HCC1806 cells (see Methods and Figure S6). **(D)** Effect of *PABPC1* knockdown on HCC1806 cell lines’ viability at 24, 48 and 72 hours.

To further investigate a possible role of *PABPC1* in regulating glycolysis, we looked more directly at perturbations in glycolysis and oxidative phosphorylation following *PABPC1* knockdown by measuring the rate of extracellular acidification (ECAR) and oxygen consumption rate (OCR) in HCC1806 cells. This highlighted a significant reduction in extracellular acidification rate, as well as oxygen consumption and ATP production, following *PABPC1* knockdown (Figures 5C, S7). Furthermore, PABPC1 silencing caused a significant reduction in cell growth, both in normoxia and hypoxia (Figure 5D).

### AGO2 and PABPC1 are over-expressed by amplification in hypoxic tumours

Hypoxia, low levels or lack of oxygen, is one of the most important microenvironmental differences between cancer and normal tissue, and is associated with progression and prognosis of many solid tumours [40]. We have previously shown that the miRNA biogenesis gene *DICER1* is reduced in hypoxic cells and tumours, and that this promotes a stem cell phenotype [15], and we have also shown alteration in the levels of several miRNAs under hypoxia, both in cell lines and cancer samples [13]. Thus, we asked whether amplification and/or expression of *AGO2* and *PABPC1* were associated with hypoxia.

Using a gene expression signature previously validated in breast cancer as hypoxia biomarker [41] we found that over-expression of *AGO2* was significantly and strongly correlated with this signature in both our Oxford dataset (rho = 0.42, p = 1.3×10^-11^) and the TCGA BRCA dataset (rho = 0.54, p > 2.2×10^-16^), and significantly - albeit less strongly - correlated with hypoxia in the Metabric dataset (rho = 0.1, p = 0.035). Furthermore, *PABPC1* correlation with the hypoxia signature was strong in all three datasets (our dataset: rho = 0.35, p = 1.0×10^-07^; Metabric dataset: rho = 0.22, p = 2.9×10^-16^; TCGA BRCA: rho = 0.50, p < 2.2×10^-16^).

To investigate a possible direct regulation of AGO2 and PABPC1 by hypoxia, we measured protein levels in normoxia and hypoxia (24h, 1% O_2_) in a panel of 7 cell lines, with and without 8q amplification (Figure S3C). Cells exposed to hypoxia did not have altered *AGO2* and *PABPC1* abundance. We then asked if we could observe a differential regulation of miRNA levels when silencing *AGO2* or *PABPC1* in cell lines exposed to hypoxia, with respect to normoxia (Figure 4B). The overall number of miRNAs differentially expressed between the knock down and control siRNA treated cells was low for *PABPC1* both when cells where cultured under hypoxia or normoxia. Conversely for *AGO2* the number of miRNAs differentially expressed between the knock down and control siRNA treated cells was high in either hypoxia or normoxia. Whilst we could identify miRNAs regulated differently when *AGO2* was silenced under hypoxia (Table S22) with respect to normoxia (Table S15), a large proportion (77/114) of these miRNAs was similarly and significantly affected in both hypoxia and normoxia, including members of the oncomiR-1 cluster (Figures 4, S5A).

These results taken together suggest that the association observed between the expression of these two genes and hypoxia could be the result of selection of cells harbouring amplification, rather than direct regulation of these genes by exposure to hypoxia.

### AGO2 defines a role for the 8q chromosome in DICER-independent biogenesis of oncogenic miRNA

AGO2 and PABPC1 genomic location, chromosome 8q, hosts a considerable number of candidate driver genes [42], the most notable examples being *MYC* [43] and *RECQL4* [44]. Such enrichment in cancer drivers could confound analyses of genes located at this region. Thus, we re-analysed the expression data to perform a detailed analysis of this region, including all genes and drivers as those defined in the COSMIC database (Table S23 for a full list). This revealed that a large proportion of genes in this region were either not expressed, or their expression was not correlated with CNA (Figure 6). This emphasizes that focusing on amplification alone might not be a sufficiently powerful approach, and that over-expression is a functional phenotype which needs to be considered. Furthermore, when we investigated how many of the candidate driver genes in this region showed significant association with prognosis, we obtained that in most cases gene expression did not predict prognosis (Figure 6).

**Figure 6.**
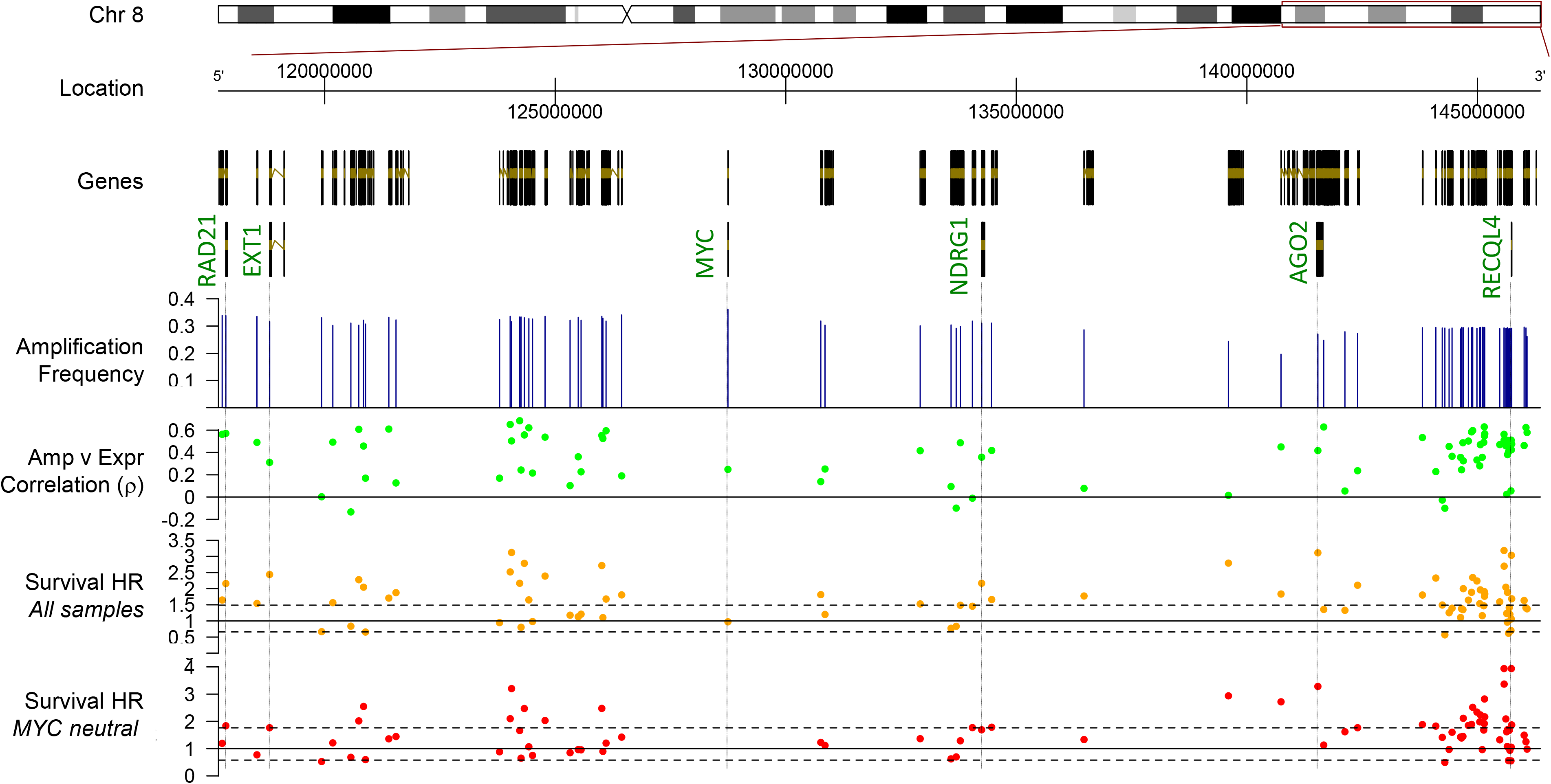
A new role of the 8q24 amplicon in miRNA biogenesis and cancer. The 8q24 amplicon chromosomal location is shown. Gene and driver genes locations are plotted alongside tracks showing values for each gene. Plots beneath genes (one point at each separate location) show amplification frequency (normalised to whole cohort), correlation between amplification and expression (Spearman rho, shown in green), and Hazard Ratio of survival analysis including all samples (in orange), and considering only *MYC* neutral cases (in red). Table S23 provides list of cancer drivers in this region and Table S24 provides full results for these analyses.

On the other hand, *AGO2* expression ranked as one of the top prognostic factors in this region, above known driver genes (Table S24). Given the well-known role of cancer driver *MYC* amplification, we repeated the above analysis in *MYC* not-amplified (neutral) cases. As expected from the role of *MYC* and from the smaller cohort, and thus decreased power, this reduced significantly the overall number of prognostic genes in the amplicon (Figure 6). However, over-expression of both *AGO2* (Figures 6, S8A) and *PABPC1* (Figures 6, S8B) were still significantly associated with poor prognosis in *MYC* neutral cases.

## Discussion

Notwithstanding the recognised role of miRNAs in health and disease, little is known about the role of miRNA biogenesis pathways in oncogenic miRNA production. Namely, the abundance of a functional miRNA can be regulated both at the transcriptional level, by increased or reduced transcription of precursor miRNAs, and by modified miRNA processing, altering the rate or efficiency at which mature miRNAs are produced and bind to target transcripts. To understand the interplay between these two mechanisms of miRNA regulation is crucial, both in terms of health and disease biology, but also in terms of therapeutic approaches in diseases like cancer, where miRNA play a crucial role.

In a pan-cancer analysis considering 9,111 tumour and normal tissue samples from 10 major cancer types, we revealed an overall global dysregulation of miRNA, both via transcription and biogenesis, across multiple cancer types. Specifically, key elements of miRISC (*TARBP1*, *AGO2* and *PABPC1*), and miRNA editing factors (*ADAR*), were frequently amplified and concordantly over-expressed across cancer types. Interestingly, in many cancers these aberrations were co-occurring, suggesting a global reprogramming of miRNA biogenesis. We also observed significant heterogeneity between cancer types, with some cancers, such as breast and lung, being characterised by the highest dysregulation of miRNA biogenesis. Importantly, we discovered *AGO2* and *PABPC1* as the most consistently over-expressed by amplification, both in human samples from multiple clinical cohorts, and *in vitro*. Importantly, we revealed that *AGO2* and *PABPC1* are associated with poor prognosis independently from, and more strongly than, known cancer drivers at nearby genomic locations, suggesting a distinct role for these genes in cancer progression. As an increasing number of small-molecule inhibitors are under development, some of them for example competing with Ago2-miRNA binding [45], targeting this miRNA regulatory gene could provide a promising novel therapeutic approach to cancer. Furthermore, the frequent co-occurrence of *TP53* mutation and AGO2 CNA we could detect across breast cancer cohorts is interesting, especially as p53, one of the most frequently mutated tumour suppressor, has been shown to interact with AGO2 to modulate AGO2’s association of a subset of miRNAs in response to DNA damage [46]. AGO2 amplification could therefore confer advantages for tumour progression in the presence of inactivating *TP53* mutations through impairing the miRNA-mediated DNA damage response. This could lead to alternative treatment for tumours harbouring this frequent mutation.

The involvement of AGO2 in the regulation of miRNA function is well-known, but recent findings have discovered the existence of an additional role. Specifically, a AGO2-mediated miRNA maturation pathway, independent of DICER1, was identified [10]. The role of this biogenesis pathway in human health and disease is still unknown. Here we have revealed recurrent amplification and over-expression of genes in this maturation pathway, including AGO2 and PABPC1, in cancer. Importantly, we observed co-occurrence of DICER1 deletions and under-expression, with amplification and over-expression of genes in the AGO2-dependent pathway, providing evidence for a switch between a DICER1-dependent and -independent miRNA processing.

As AGO2 is the key miRNA guide protein in the miRISC [25], a rise in *AGO2* expression levels would be expected to increase the production of mature miRNAs. Indeed, upon silencing of AGO2 in breast cancer cell lines harbouring amplification or high AGO2 mRNA expression levels, we observed both a significant global dysregulation of miRNA levels and also preferential regulation of specific miRNAs, including the miR-106b/25 cluster of the oncomiR-1 family, in agreement with analysis of the clinical samples. We were also able to confirm selective maturation of this oncomiR-1 cluster by analysing the ratio of mature miRNA to immature pri-miRNA levels in clinical samples, which showed increased levels of mature miRNA with respect to precursor miR-106b/25 miRNA in AGO2 amplified and over-expressing samples. We additionally demonstrated AGO2-dependent, DICER1-independent, maturation, but not transcription, of the miR-106b/25 cluster of the oncomiR-1.

Another paralogue member of the oncomiR-1 family, the miR-17-92 cluster, is regulated by proto-oncogene *MYC*. Accordingly, when we considered only *MYC* neutral cases, abundance of miRNA belonging to this cluster was not prognostic. However, the oncomiR-1 paralogue cluster miR-106b/25, and in particular miR-93, remained a significant prognostic factor irrespective of *MYC* amplification, confirming the hypothesis of a complementary and independent role of these two clusters. Importantly, miR-93 targets include genes such as CDKN1A and E2F1 which are known tumour suppressors [32].

*AGO2* and *PABPC1* polyA binding protein share near, although not proximal, locations on chromosome 8q, 8q24 and 8q22 respectively. We produced strong evidence that these genes are co-amplified and co-expressed. PABPC1 interacts with miRISC to regulate the extent of target inhibition achieved by the miRNA, rather than regulating miRNA maturation per se [30, 31]. More specifically, the *TNRC6A* (*GW182*) protein interacts with *AGO2* protein through an amino-terminal domain containing multiple Gly-Trp (GW) repeats, compete with eukaryotic initiation factor 4G (eIF4G) for binding to *PABPC1* and promote translational repression and mRNA degradation through the interaction of a bipartite silencing domain with *PABPC1* (Figure 1C) [30]. In agreement with this understanding, both our analysis of clinical and *in-vitro* data found that *PABPC1* is less associated with changes in miRNA abundance than AGO2, but is associated with important changes in gene expression and support a pro-survival shift in glycolysis and oxidative phosphorylation. This is key pathway in cancer progression [47, 48].

Previous studies by us and others have shown that hypoxia affects expression of genes in the biogenesis machinery. Examples are EGFR modulation of microRNA maturation in response to hypoxia through phosphorylation of AGO2 [49], and DICER1 suppression by hypoxia, promoting stemness [15]. Here, we find that in clinical samples, hypoxia is strongly associated with the AGO2 cluster amplification and over-expression, but we could not detect direct regulation by hypoxia in cell lines. We have previously observed a similar phenomenon for other cancer drivers [50], whereby over-expression is likely to be caused by selection, rather than by direct regulation. 21

In conclusion, in a comprehensive pan-cancer analysis of somatic mutations, genomic amplifications, and expression of coding and non-coding RNAs, we revealed for the first time a global landscape of dysregulation of miRNA biogenesis in cancer. This demonstrated recurrent genomic alterations of enzymes and factors responsible for the production of functional miRNA, and revealed a switch to AGO2-mediated, DICER-independent, miRNA biogenesis, leading to selective maturation of oncogenic miRNA.

Our findings have important implications for cancer prognosis, as circulating miRNAs are promising non-invasive biomarkers for cancer diagnosis and can provide useful insights with regards to therapy response, as many miRNAs regulate important tumour suppressor genes [5], which can impact on cancer evolution and cancer drug resistance. Finally, as an increasing number of miRNA-based therapeutics, including anti-microRNA constructs, and small-molecule RISC- and Ago2-inhibitors are entering pre-clinical and clinical trials, our findings open up a promising new avenue for cancer treatment.

## Methods

### Clinical Datasets

Cancer and normal tissue samples’ genomic and/or transcriptomic data (Table S1) were downloaded from TCGA (https://cghub.ucsc.edu/) for 9,111 samples, covering 10 major cancer types. Specifically, this dataset consists of 3,990 cancer samples with complete genomic (SNP6) and transcriptomic (mRNA and miRNA-seq) data; 3,990 matched normal tissue samples with genomic (SNP6) data, of which 395 also had transcriptomic (RNA-seq) data; and 1,131 additional cancer/normal tissue samples with gene expression and miRNA microarray data. Additionally, for 3,310 cancer samples we used the previously published TCGA mutation calls [18–24]. An additional independent validation clinical cohort (1,991 cancer and 144 normal tissue samples), the Metabric breast cancer cohort [2, 17], was used by kind permission. Raw Metabric files were downloaded from European genome-phenome archive (EGA) (Study ID: EGAS00000000083). We also considered an additional validation cohort of 198 early breast cancer samples from patients treated in Oxford, with full clinical annotation, complete 10-years follow-up, and global microRNA, mRNA expression and CNA profiles. Clinical data, and generation of mRNA and miRNA expression profiles for this cohort have been described previously [51]; Illumina Cytosnp-12 Infinium arrays were used to estimate CNA profiles using OncoSNP v2.19 [52].

### Pre-processing and filtering

mRNA abundance, DNA copy-number and somatic mutation profiles for the TCGA datasets were downloaded from TCGA DCC (gdac), release 2014-01-15. Amplification was reported both using GISTIC [53] calls and logged base2 intensity. For mRNA abundance, Illumina HiSeq rnaseqv2 level 3 RSEM normalised profiles were used. Pre-processing and filtering of the TCGA data have been previously described [50]. The Metabric breast cancer dataset was pre-processed, summarised and quantile-normalised from the raw expression files generated by Illumina BeadStudio (R packages: be*ADAR*ray v2.4.2 and illuminaHuman v3.db_1.12.2). Probe to gene-level mapping was performed by keeping the most variable (standard deviation) probe. Log_2_ scaled data was used for differential expression analysis. Metabric copy-number segmented genomic regions data, as published in the original study [17] (Human genome assembly: Hg18), was processed to create gene by patient log_2_ ratio and calls profiles.

### Statistical and Bioinformatics Analyses

Penalized Generalized Linear Model (GLM) Regression was used with cross-validation to determine optimal L1 and L2 penalization parameters (Suppl File 4 for details). Pan-cancer mRNA and CN analyses were performed using our previously developed R package iDOS [50]. All analyses were performed using the R statistical environment (https://cran.r-project.org/). Unless otherwise stated we have used ANOVA to establish difference between experimental conditions/treatments. Effects of expression on overall outcome and patient relapse were assessed by Cox proportional hazard models (*survival* R package). Differential gene expression analysis of miRNA was done using *Limma* R package (Suppl. File 4). Analyses described in this manuscript were performed either with R 3.4 or Prism 8.4.2.

### Cell line experiments

HCC1806 cells were obtained from LGC (ATCC) and grown in RPMI media, supplemented with 10% FCS. For hypoxic exposure, cells were grown at 1 % O_2_, 5 % CO_2_ and 37 °C in an INVIVO2 400 (Baker Ruskinn, USA). Transfections of 100 nM siRNA duplexes targeting AGO1, 2, 3 and 4 and *PABPC1* (ON-TARGETplus SMARTpool) or a scramble control (ON-TARGETplus SMARTpool) were performed in Optimem (Life Technologies), using Oligofectamine (Life Technologies). Methods for microarray, Western blots and qPCR are described in Supplementary File 4.

### Metabolic analysis

The rate of extracellular acidification (ECAR) and the oxygen consumption rate (OCR) were measured using the Seahorse XF96e Bio-analyzer (Seahorse Bioscience, North Billerica, MA). Briefly, HCC1806 cells were plated into XF96e cell culture plates (Seahorse Bioscience, North Billerica) with a density of 40,000 cells/well. After overnight incubation, the culture medium was changed to XF base medium (without glucose). After measurement of basal ECAR levels, 11 mM glucose was injected to determine the glycolytic rate, followed by injection of 1 uM oligomycin and finally injection of 50 mM 2-DG. Similarly, after measurement of basal OCR levels, 5 mM glucose was injected, followed by injection of 2 uM oligomy*c*in and finally injection of 50 mM 2-DG. Data was normalised using Cyquant (Life Technologies).

### Availability of data and materials

The code used for penalized GLM used to identify miRNA and pri-miRNA associated with Cancer Hallmarks Signatures can be found in the Buffa Lab GitHub repository. Our open source R package (iDOS) implementing the methodology described here to identify candidate driver genes [50] is available through CRAN (http://cran.r-project.org), and example scripts on how to use the package are also available to through Zenodo with DOI: 10.5281/zenodo.51787 (https://zenodo.org/record/51787).

## Abbreviations

CN: Copy Number; CNA: Copy Number Amplification; miRNA: microRNA; miRNA induced silencing complex (miRISC); The Cancer Genome Atlas (TCGA);

## Authors’ Contributions

FMB conceived the idea and designed the study. LW, AD, SH, CH and FMB performed bioinformatics and computational analyses. LvB, SW and AM performed experimental work with input from ALH, FMB and TH. LW, LvB and FMB wrote the manuscript with contribution from all authors.

## Acknowledgements

We would like to thank Anna Git and Heidi Dvinge for informative discussions on normalization of miRNA array data. This study makes use of data generated by the Molecular Taxonomy of Breast Cancer International Consortium, which was funded by Cancer Research UK and the British Columbia Cancer Agency Branch. The results published here are in part based upon data generated by The Cancer Genome Atlas pilot project established by the NCI and NHGRI. Information about TCGA and the investigators and institutions who constitute the TCGA research network can be found at http://cancergenome.nih.gov/.

