## Supplementary Figures for "Dicer-to-Argonaute switch controls biogenesis of oncogenic miRNA"

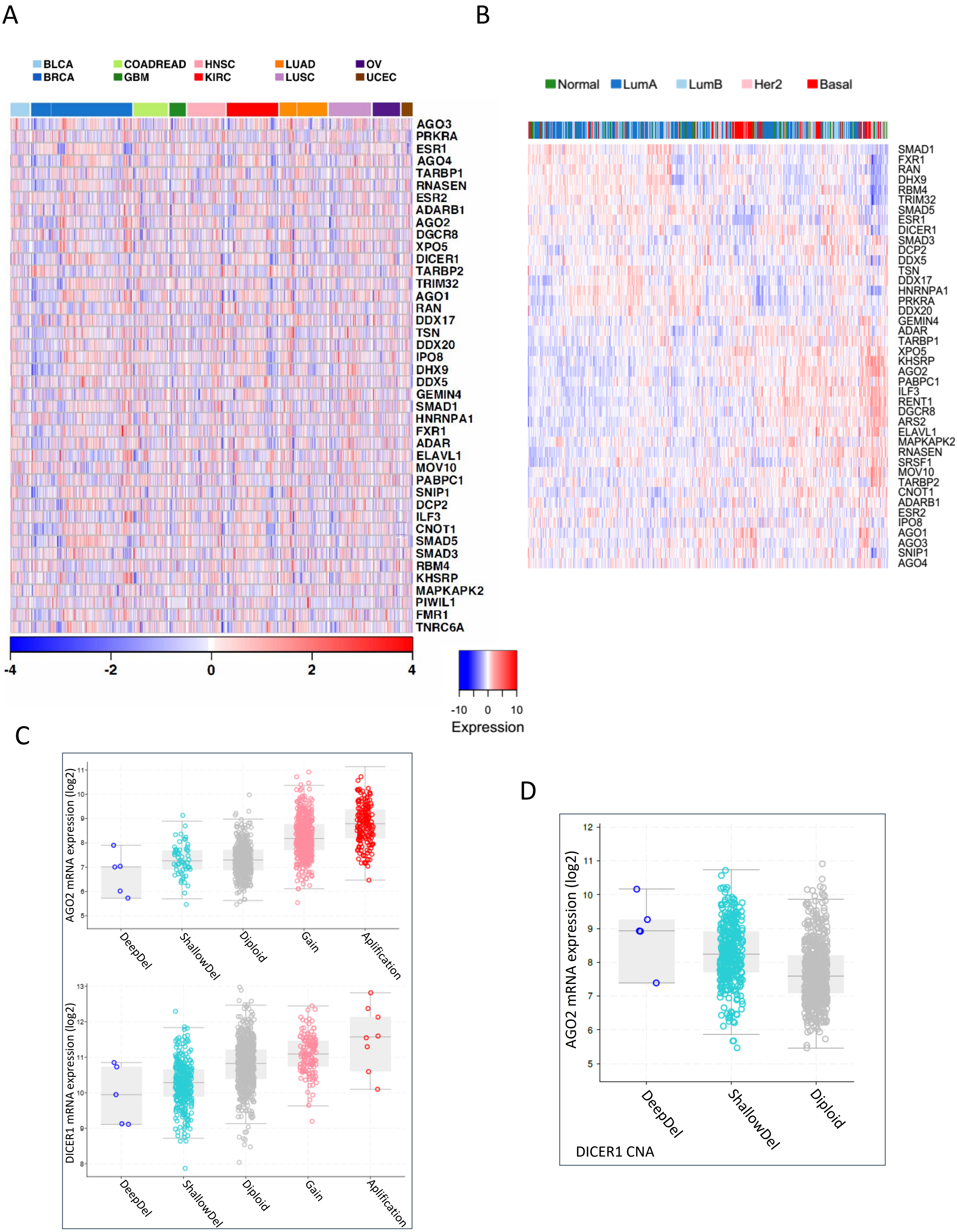

A
B
BLCA
BRCA
COADREAD
GBM
HNSC
KIRC
LUAD
LUSC
OV
UCEC
AGO3
PRKRA
ESR1
AGO4
TARBP1
RNASEN
ESR2
ADARB1
AGO2
DGCR8
XPO5
DICER1
TARBP2
TRIM32
AGO1
RAN
DDX17
TSN
DDX20
IPO8
DHX9
DDX5
GEMIN4
SMAD1
HNRNPA1
FXR1
ADAR
ELAVL1
MOV10
PABPC1
SNIP1
DCP2
ILF3
CNOT1
SMAD5
SMAD3
RBM4
KHSRP
MAPKAPK2
PIWIL1
FMR1
TNRC6A
-4 -2 0 2 4
Normal
LumA
LumB
Her2
Basal
SMAD1
FXR1
RAN
DHX9
RBM4
TRIM32
SMAD5
ESR1
DICER1
SMAD3
DCP2
DDX5
TSN
DDX17
HNRNPA1
PRKRA
DDX20
GEMIN4
ADAR
TARBP1
XPO5
KHSRP
AGO2
PABPC1
ILF3
RENT1
DGCR8
ARS2
ELAVL1
MAPKAPK2
RNASEN
SRSF1
MOV10
TARBP2
CNOT1
ADARB1
ESR2
IPO8
AGO1
AGO3
SNIP1
AGO4
-10 0 10
Expression
C
AGO2 mRNA expression (log2)
DeepDel
ShallowDel
Diploid
Gain
Aplification
DICER1 mRNA expression (log2)
DeepDel
ShallowDel
Diploid
Gain
Aplification
D
AGO2 mRNA expression (log2)
DeepDel
ShallowDel
Diploid
DICER1 CNA

**Figure S1. Pan-cancer expression and copy number aberrations (CNA) for the miRNA biogenesis pathway.** A) Pan-cancer mRNA abundance profiles. Datasets are as described in Table S2. Patients were ordered using hierarchical clustering within each cancer type separately. B) Breast cancer mRNA abundance profiles for the Metabric dataset (Material and Methods). Rows represent genes and columns represent patients. Genes in Table S1 were considered. mRNA profiles for each tumour type was *z*-transformed. C) AGO2 CNA versus gene expression (top) and DICER-1 CNA versus gene expression (bottom) in TGCA BRCA data; D) AGO2 expression in DICER1 deleted and diploid cases.

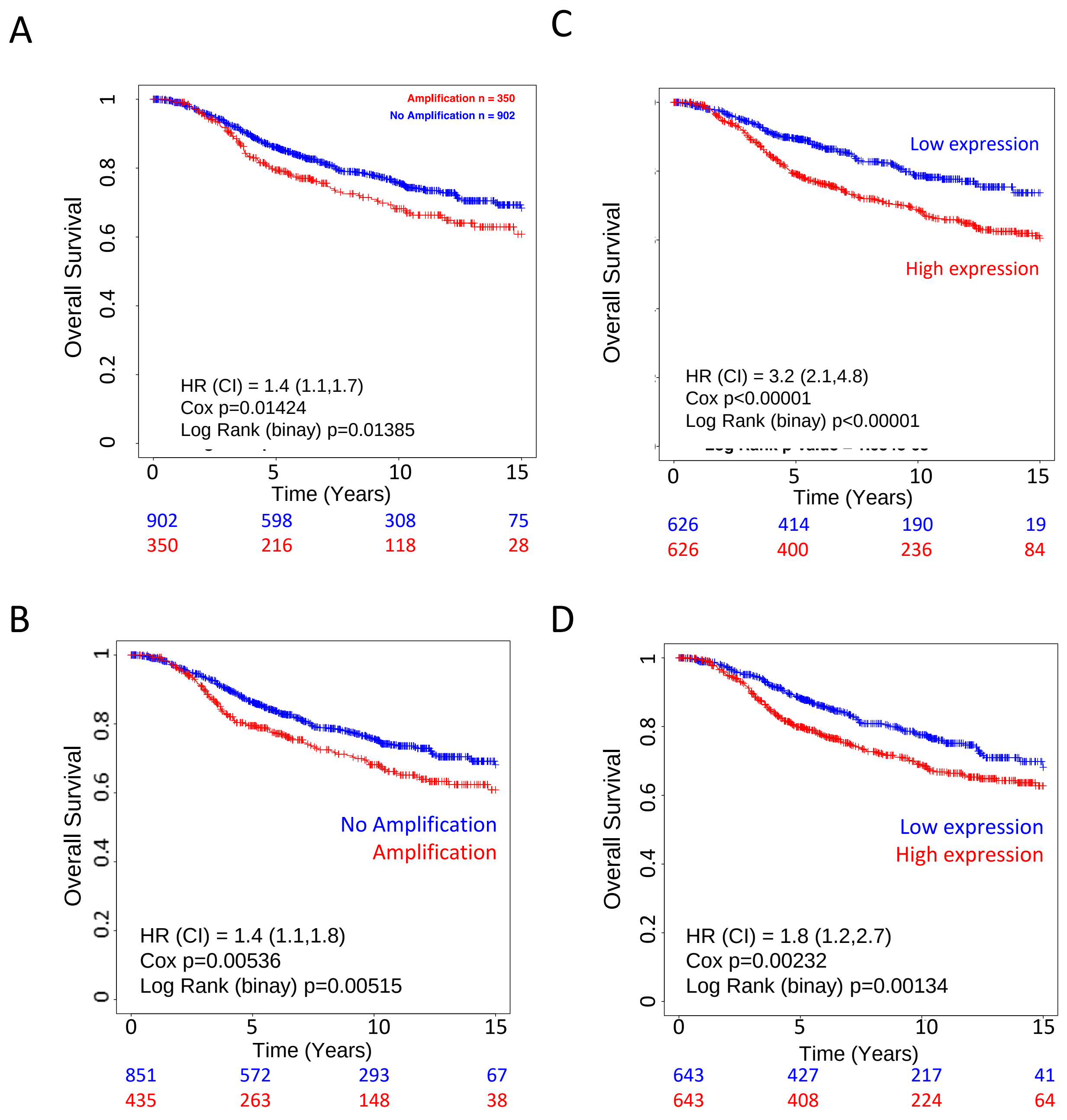


**Figure 2 AGO2 and PABPC1 expression associated with poorer survival in the Metabric cohort.** (A) Survival analysis of AGO2 and (B) *PABPC1* CNA status defined as amplification versus no amplification (neutral and loss). Cox analysis results are shown (HR = Hazard Ratio). (C) Survival analysis of AGO2 and (D) *PABPC1* mRNA expression levels. Gene expression was separated by median into low and high equally sized groups.

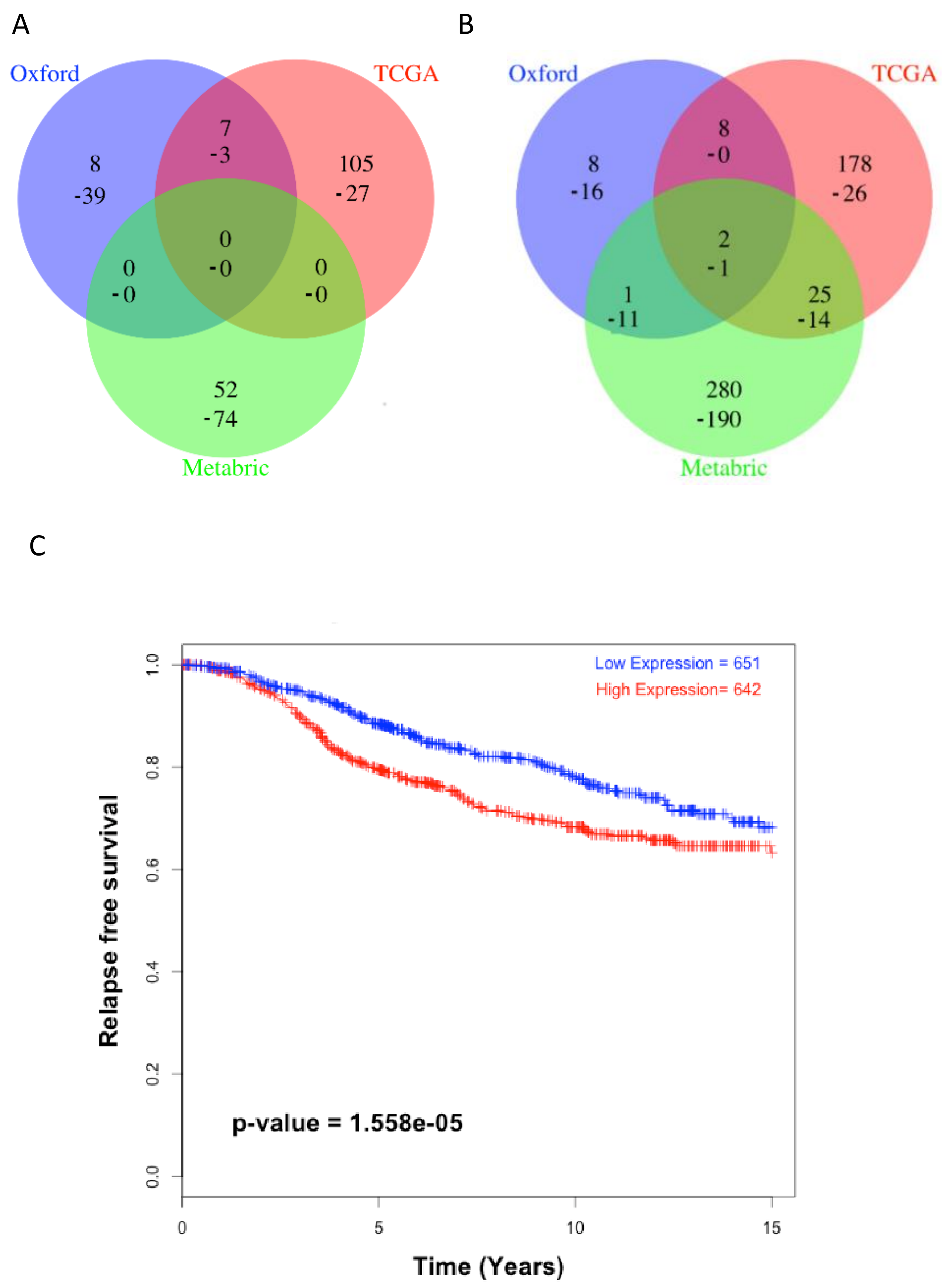

A
Oxford
TCGA
Metabric
8
-39
7
-3
105
-27
0
-0
0
-0
0
-0
52
-74
B
Oxford
TCGA
Metabric
8
-16
8
-0
178
-26
2
-1
1
-11
25
-14
280
-190
C
Low Expression = 651
High Expression= 642
p-value = 1.558e-05
Relapse free survival
Time (Years)
0.0
0.2
0.4
0.6
0.8
1.0
0
5
10
15

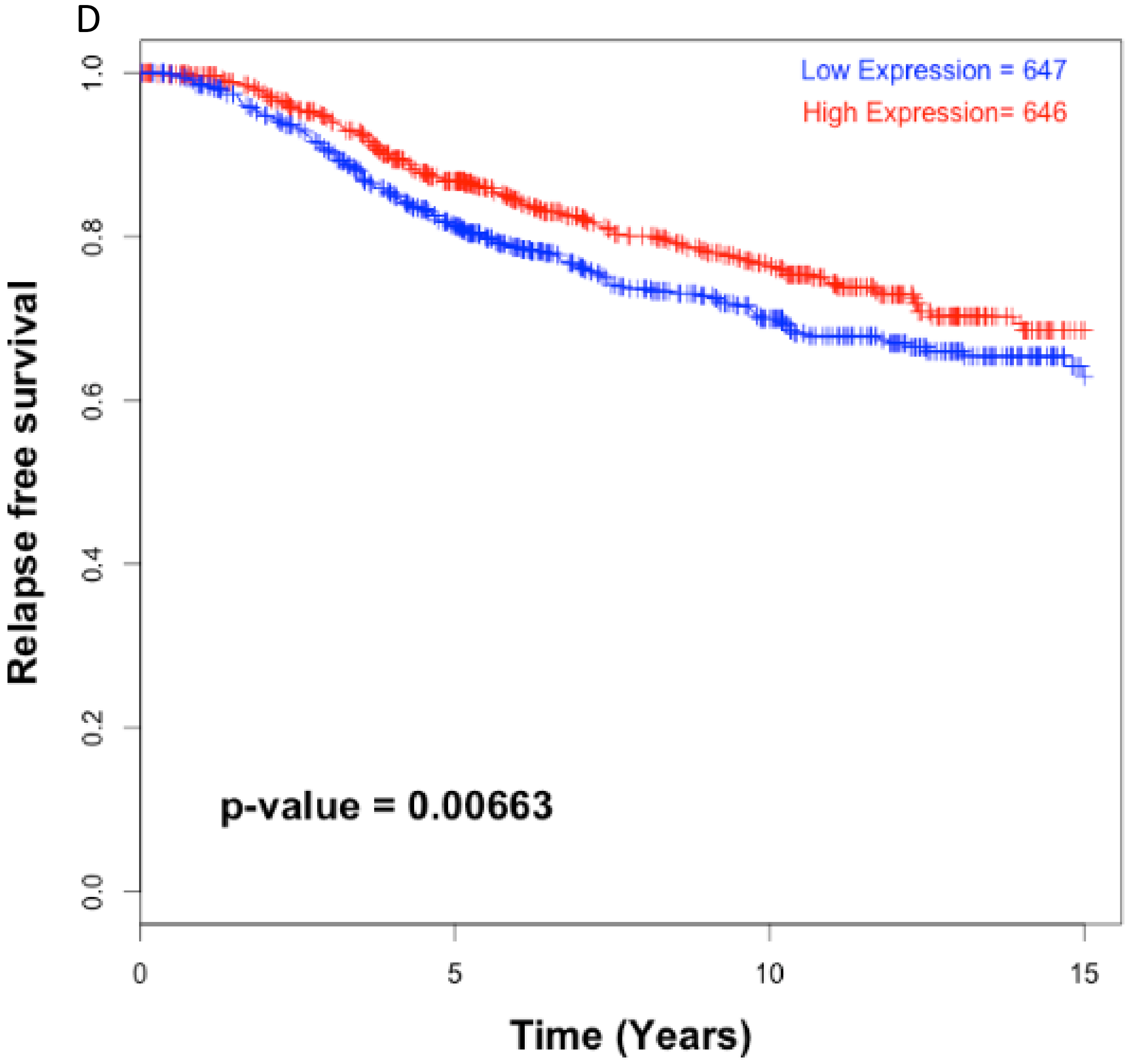


**Figure S3. AGO2 and PABPC1 associated miRNA expression profiles, and their prognostic significance, in breast cancer cohorts.** miRNAs with expression significantly correlated (Spearman) with AGO2 or PABPC1 expression were selected (FDR<0.05) in the three independent breast cancer cohorts considered in this study. **(A)** Overlap between the PABPC1 miRNA profiles. The number of common positively and negatively (indicated with minus sign "-") correlated miRNAs is shown. **(B)** Overlap between the AGO2 miRNA profiles derived in MYC neutral cases in the three cohorts (see also Table S13). **(C)** Survival analysis (relapse free survival, Metabric cohort, N=1,293) for miR-93 (Hazard Ratio = 2.35) and **(D)** miR-10a (Hazard Ratio = 0.57), co- and inversely correlated respectively with AGO2 expression in all three cohorts. Samples were split by median expression into two, "low" and "high", equal groups for the plots, but considered as continuous variables in the Cox analysis.

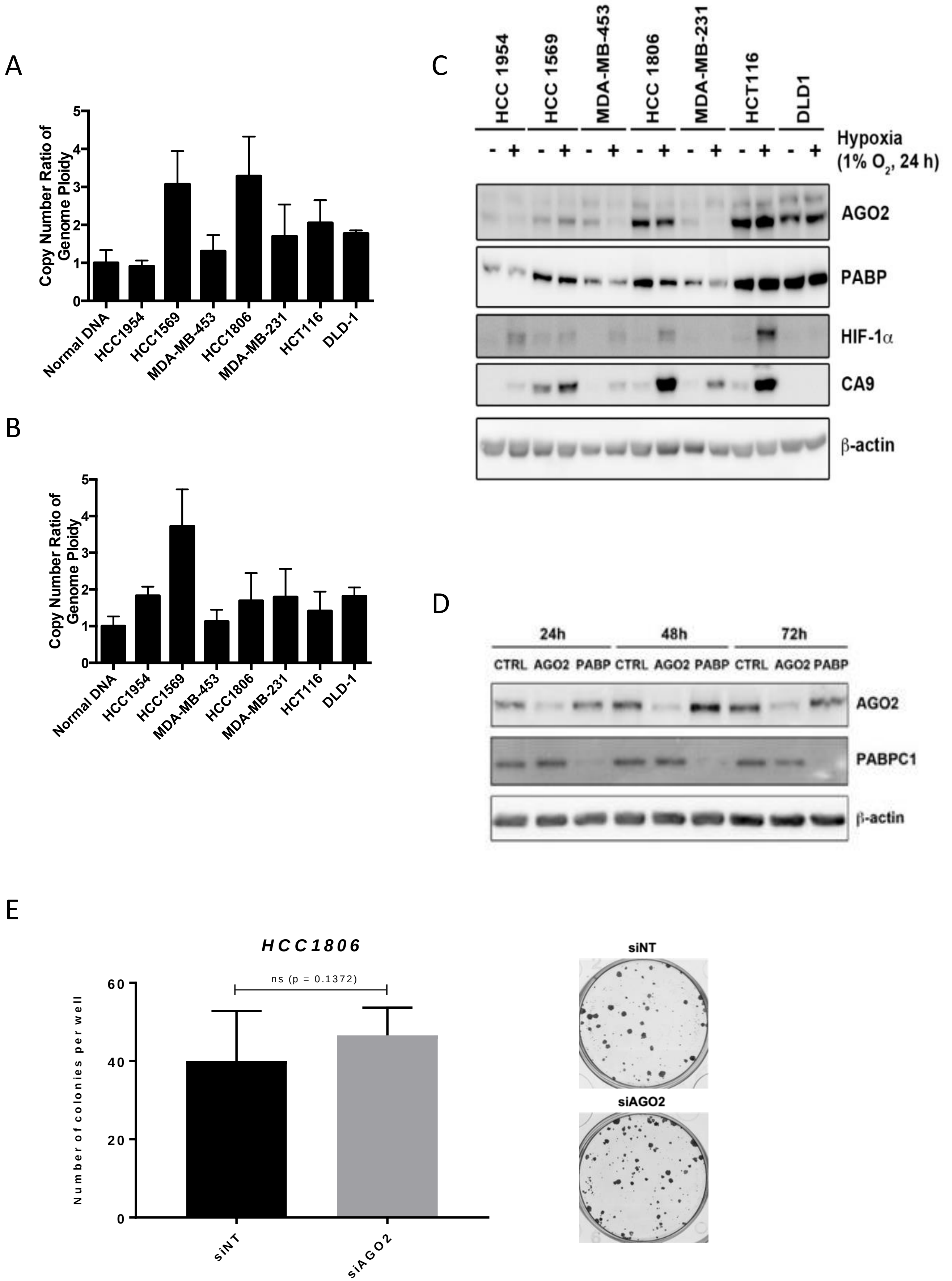

A
Copy Number Ratio of Genome Ploidy
0
1
2
3
4
5
Normal DNA
HCC1954
HCC1569
MDA-MB-453
HCC1806
MDA-MB-231
HCT116
DLD-1
B
Copy Number Ratio of Genome Ploidy
0
1
2
3
4
5
Normal DNA
HCC1954
HCC1569
MDA-MB-453
HCC1806
MDA-MB-231
HCT116
DLD-1
C
HCC 1954
HCC 1569
MDA-MB-453
HCC 1806
MDA-MB-231
HCT116
DLD1
Hypoxia (1% $O_2$, 24 h)
- + - + - + - + - + - + - +
AGO2
PABP
HIF-1α
CA9
β-actin
D
24h
48h
72h
CTRL AGO2 PABP CTRL AGO2 PABP CTRL AGO2 PABP
AGO2
PABPC1
β-actin
E
HCC1806
Number of colonies per well
0
20
40
60
ns (p = 0.1372)
siNT
siAGO2
siNT
siAGO2

**Figure S4. Expression and amplification status of AGO2 and PABPC1 in a panel of cell lines.** (A) The relative genomic copy-number of *AGO2* and (B) *PABPC* (normalized to ploidy). (C) Expression of AGO2 and PABPC1 (PABP) in a panel of cell lines by Western blot analysis. Expression in normoxia and 24 hours hypoxia is shown. Expression of other genes shown as control. "-" denotes normoxia, "+" denotes hypoxia (1% Oxygen, 24 hours). Other genes in the miRNA biogenesis (Drosha and Dicer), and hypoxia regulated genes are shown as positive controls. (D) Validation of *AGO2* and *PABPC1* knockdown by siRNA in HCC1806. (E) Representative colony formation assay 10 days after siNT or siAGO2 treatment (HCC1806 cells). Data points and bars represent the mean and SEM of ≥ three independent experiments.

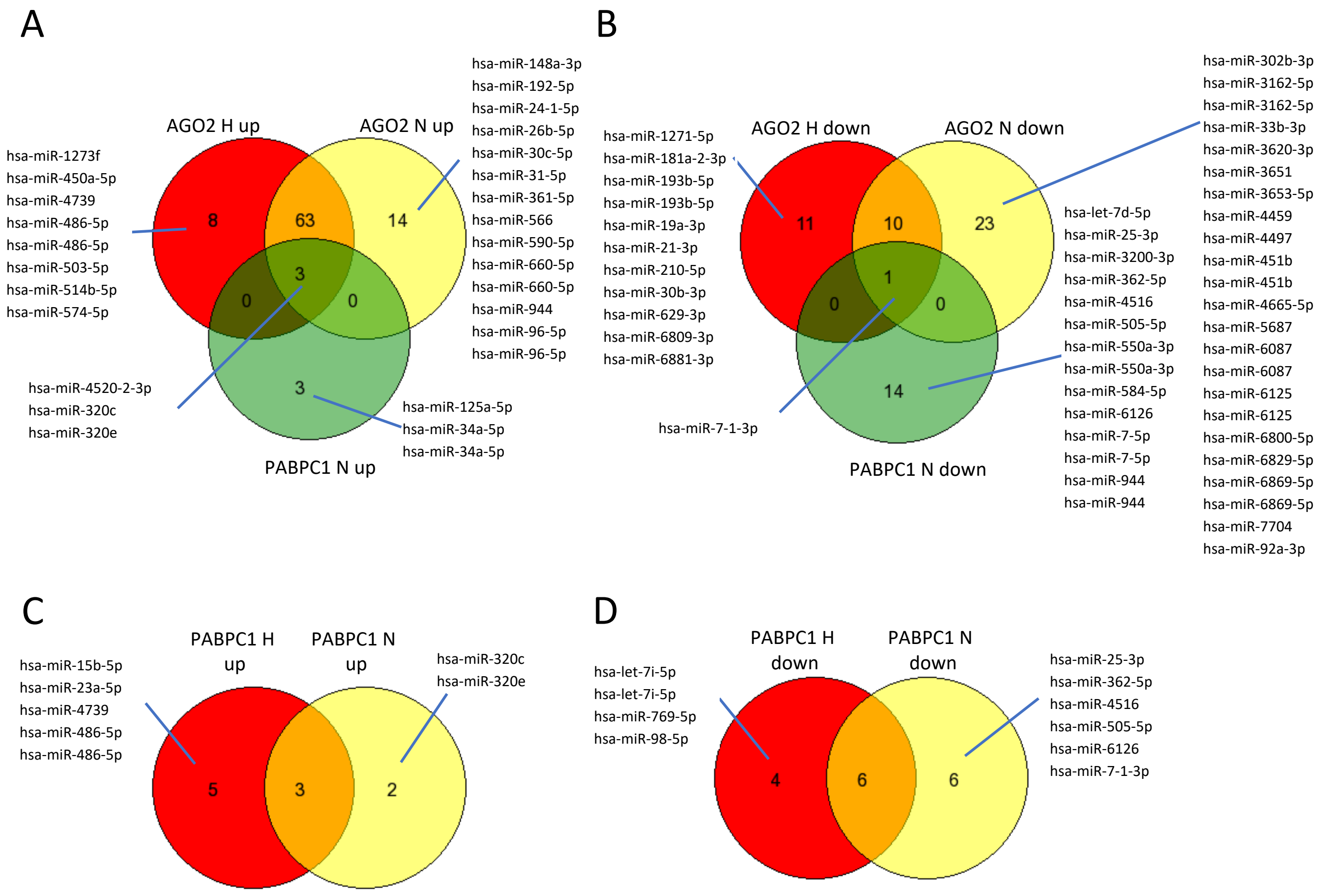


**Figure S5. Differentially expressed miRNAs in AGO2 and PABPC1 knockdown in HCC1806.** Overlap between miRNA profiles obtained by AGO2 or PABPC1 knockdown experiments in hypoxia (H) and normoxia (N). Common and uniquely differentially expressed miRNAs are shown. (A) and (C) upregulated profiles: miRNAs with higher expression in the experimental condition (AGO2, PABPC1 or H). (B) and (D) downregulated profiles: miRNAs with higher expression in the control conditions (Scramble and N). Analysis results in Tables S15-17.

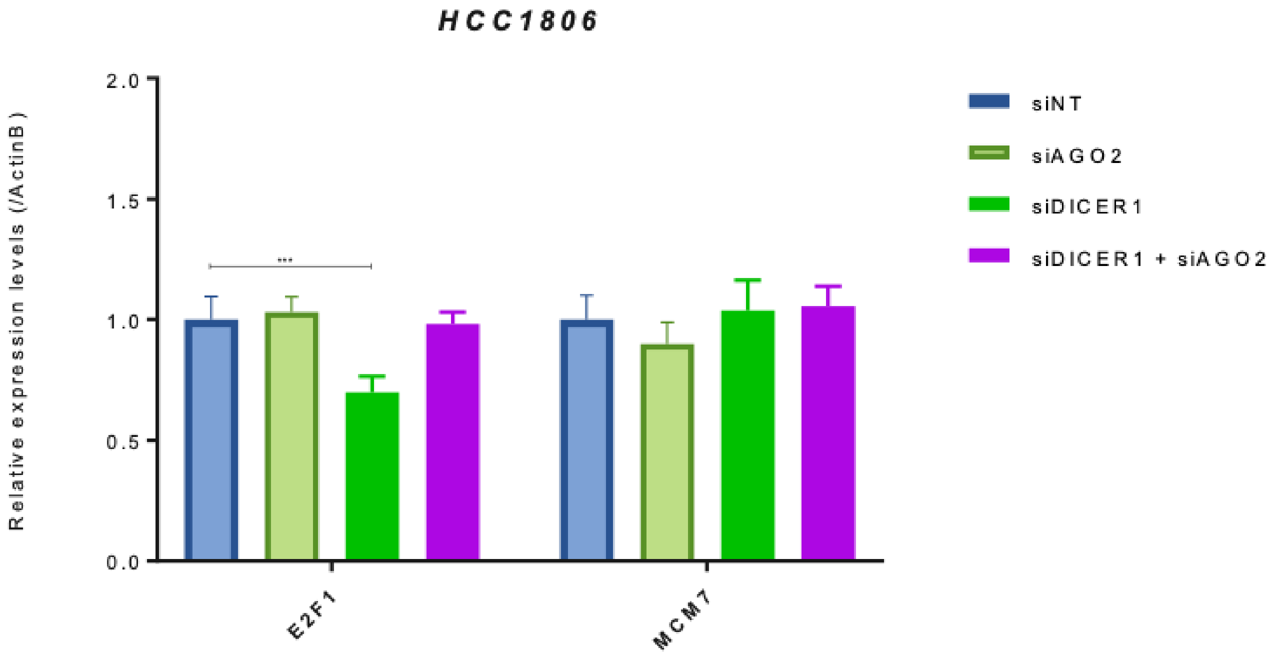


**Figure S6. AGO2 regulates miRNA maturation in HCC1806 and JIMT-1 cells.** HCC1806 cells were transfected with a siRNA against AGO2 and/or DICER, and expression of E2F1 and host gene MCM7 were determined by qRT-PCR. Error bars represent 95% confidence intervals for the differential expression measurements.

A

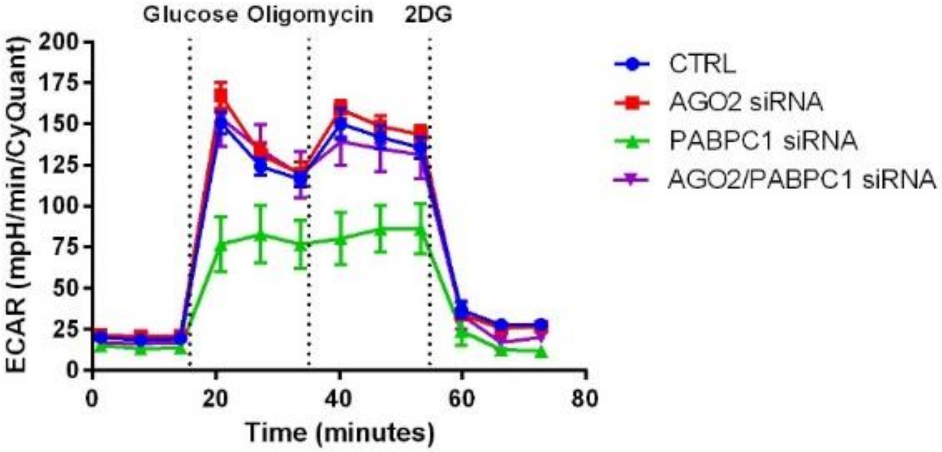
Glucose Oligomycin 2DG
ECAR (mpH/min/CyQuant)
Time (minutes)
CTRL
AGO2 siRNA
PABPC1 siRNA
AGO2/PABPC1 siRNA

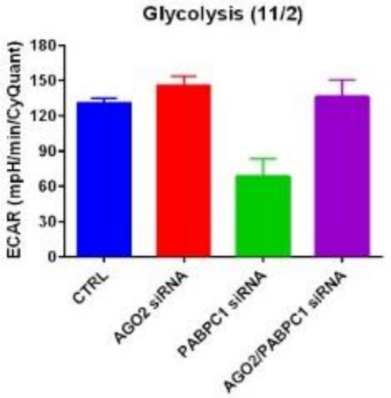
Glycolysis (11/2)
ECAR (mpH/min/CyQuant)
CTRL
AGO2 siRNA
PABPC1 siRNA
AGO2/PABPC1 siRNA

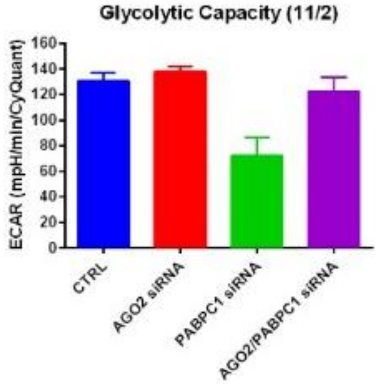
Glycolytic Capacity (11/2)
ECAR (mpH/min/CyQuant)
CTRL
AGO2 siRNA
PABPC1 siRNA
AGO2/PABPC1 siRNA

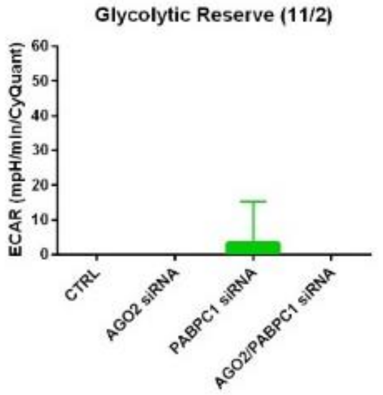
Glycolytic Reserve (11/2)
ECAR (mpH/min/CyQuant)
CTRL
AGO2 siRNA
PABPC1 siRNA
AGO2/PABPC1 siRNA

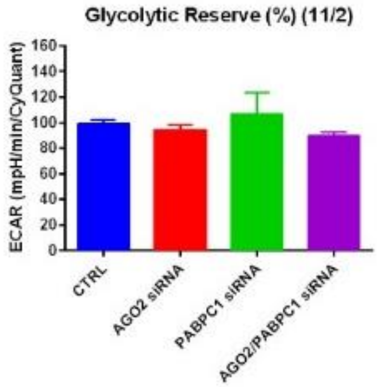
Glycolytic Reserve (%) (11/2)
ECAR (mpH/min/CyQuant)
CTRL
AGO2 siRNA
PABPC1 siRNA
AGO2/PABPC1 siRNA

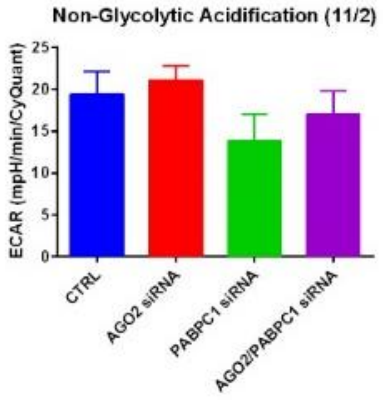
Non-Glycolytic Acidification (11/2)
ECAR (mpH/min/CyQuant)
CTRL
AGO2 siRNA
PABPC1 siRNA
AGO2/PABPC1 siRNA

B

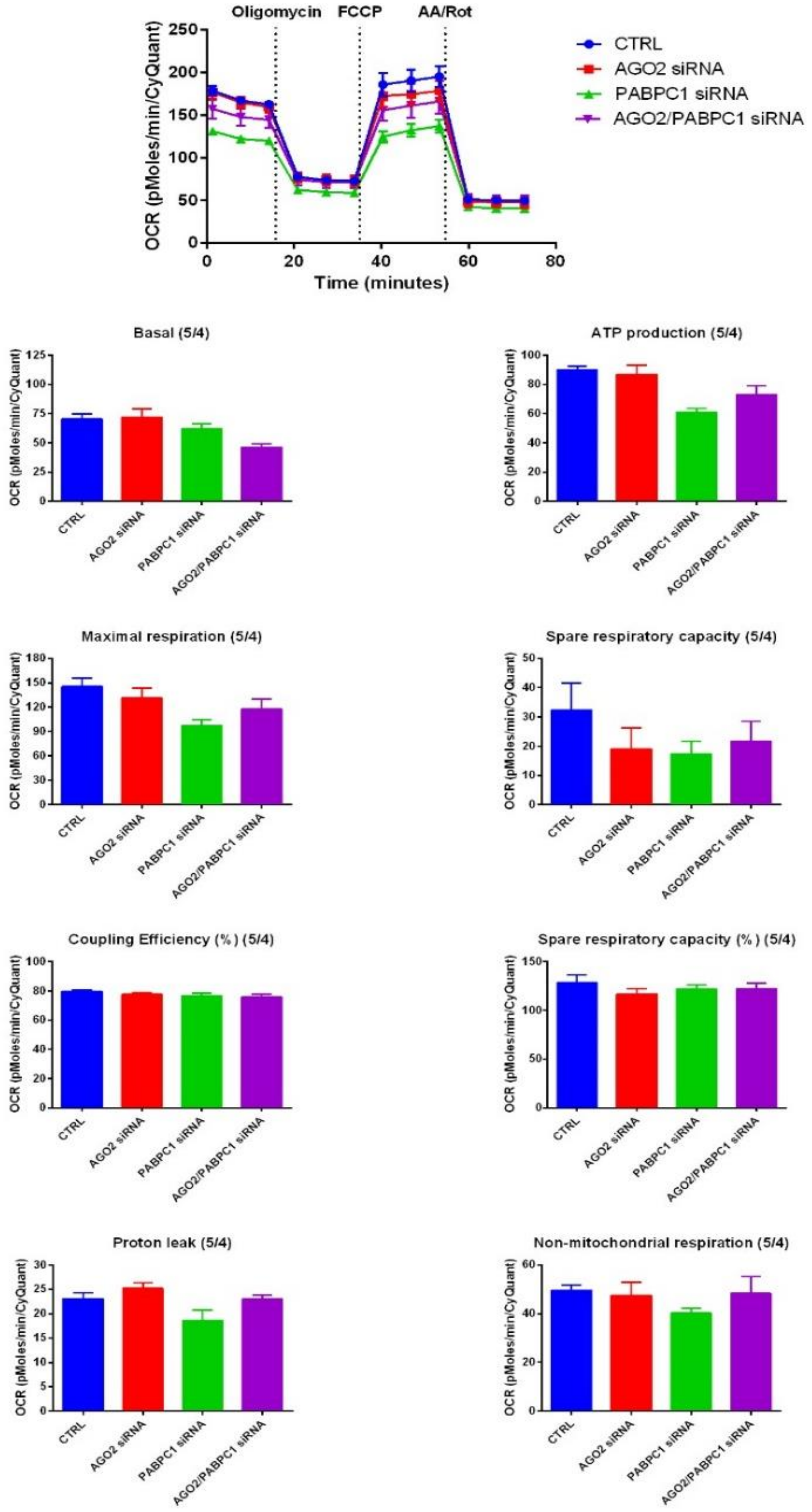

Oligomycin
FCCP
AA/Rot
OCR (pMoles/min/CyQuant)
Time (minutes)
CTRL
AGO2 siRNA
PABPC1 siRNA
AGO2/PABPC1 siRNA
Basal (5/4)
ATP production (5/4)
Maximal respiration (5/4)
Spare respiratory capacity (5/4)
Coupling Efficiency (%) (5/4)
Spare respiratory capacity (%) (5/4)
Proton leak (5/4)
Non-mitochondrial respiration (5/4)

**Figure S7. Effect of *PABPC1* and *AGO2* knockdown on the rate of extracellular acidification (ECAR) and the oxygen consumption rate (OCR).** ECAR (A) and OCR (B) were measured in HCC1806 cells (see Methods). In each condition, average Cyquant normalized data are shown (x3 replicates). High glucose, low glutamine was used in (A) to provide maximum input to measure lactate production; low glucose, high glutamine was used in (B) to maintain the Krebb cycle.

A

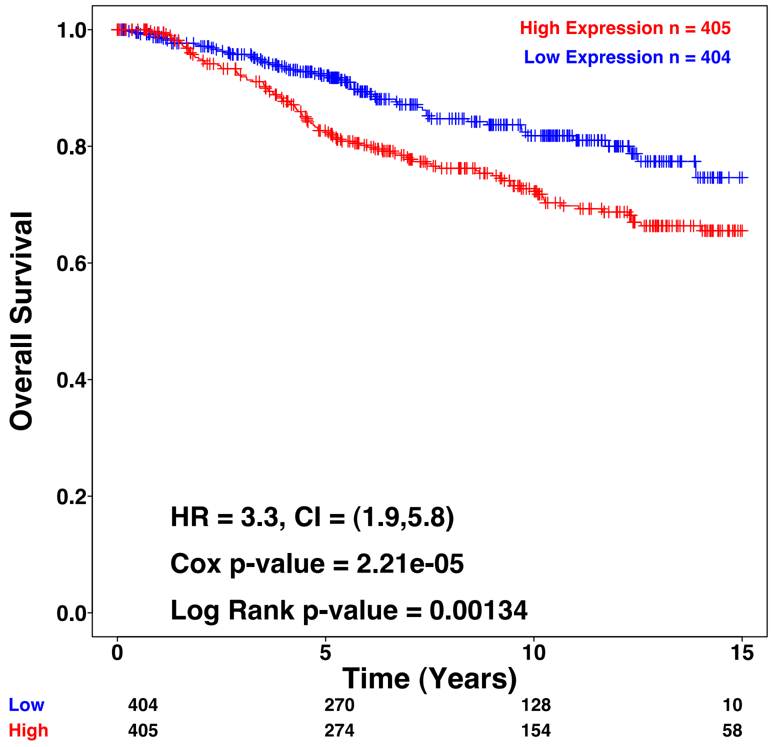


B

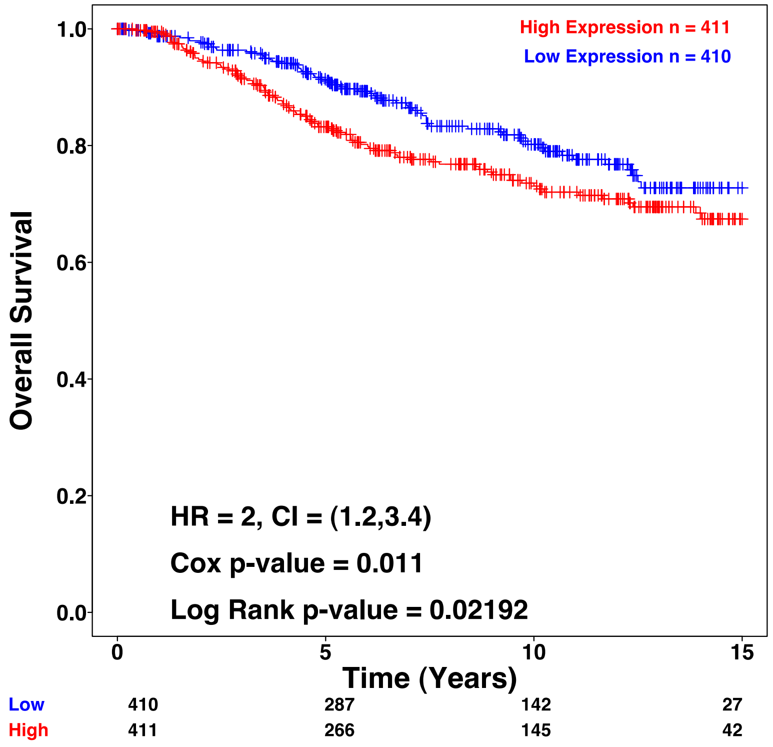


**Figure S8. AGO2 and PABPC1 are prognostic in MYC neutral cases.** Survival analysis of AGO2 (A) and PABPC1 (B) mRNA expression levels using Metabric dataset. Gene expression was separated by median into low and high to provide balanced groups. Only MYC neutral (no amplified) cases were considered.
