## Supplementary Methods for "Dicer-to-Argonaute switch controls biogenesis of oncogenic miRNA"

### A Dicer-to-Argonaute genomic switch regulates miRNA biogenesis in cancer

#### *Copy Number Data Pre-processing in clininca datasets: Oxford Dataset*

Extracted DNA was amplified and hybridised to Illumina Cyto-12 SNP arrays. For each SNP, the Log R Ratio (LRR) was calculated from the expected normalised intensity compared to observed normalised intensity and B Allele Frequency (BAF) was derived from the difference between the expected position of the cluster group and the actual value. Values were used by OncoSNP to predict CN state at each locus. Samples with high level of noise in the raw data were flagged by OncoSNP. Where correction was not possible using alternative parameters these samples were removed from analysis due to the high number of potential false positive calls present. Predicted calls were then smoothed to remove anomalous events from the data.

Each processed sample CN profile comprised of a list of predicted segments with a CN status between 0 – 6 (normal CN = 2). Where required for frequency counts, CN values above 2 were considered amplified and greater than 3 were highly amplified. Genes were considered amplified when the full region of the gene was located in the amplicon segment.

#### *Genomic Data Pre-processing and quality control in clinical datasets*

METABRIC data was downloaded in a pre-processed format from European Genome-Phenome Archive (EMBL-EBI). For each TCGA data type (CN, mRNA and miRNA expression) individual patient files containing RSEM generated read counts were downloaded and combined into a single matrix. The data was then $\log_2$ transformed and filtered to remove transcripts were > 0.75 of patients showed expression. CN prediction data was generated using GISTIC therefore samples were considered amplified only if CN status was 3 or above; continuous log2 values were also considered. For gene expression microarray data, relevant probes were selected for each biogenesis gene. In some cases a probe was not available or was eliminated based on expression range > 2. For Metabric data, a pre-processed matrix was used (methods described by Dvinge et al). Alternative probes were selected from the EGAD00010000210 dataset where genes were underperforming.

To give a parameter for patient amplification frequency an assessment of randomly selected single genes amplification was performed. 50 random genes were selected for each permutation (n = 1000) and the median value was taken. Therefore for the discovery set the frequency of amplification cut-off was 23 (frequency = 0.128), in the Metabric replication set greater than 40 (frequency = 0.031) patients carrying the amplicon were considered highly amplified.

##### *Genomic and transcriptomic analyses in clinical datasets*

Full analysis was implemented in the R environment (3.1.0) using the survival package (2.37-7) in addition to the basic libraries. Plotting libraries used include GenomeGraphs (1.24.0) and gplots (2.16.0).

After selection of the biogenesis machinery genes selection of candidates using three filtering stages in all three datasets. Firstly genes that were highly amplified based on background amplification levels in the studied patient set were selected. Then correlation analysis (Spearman's coefficient) between CN levels and mRNA expression was performed to remove genes not affected by amplification changes. Finally Cox survival tests were performed on amplification status (after removal of the deleted patients).

To isolate candidate genes in the *MYC* region the METABRIC dataset was used and a similar stage analysis was performed assessing frequency of amplification and correlation between mRNA and CN. Cox survival testing was then performed for each mRNA expression of gene in the region.

Assessment of *MYC* independence was performed after removal of *MYC* amplified patients. Patient CN was used to eliminate these samples and then analysis repeated. Given the location of *AGO2* and *MYC* few *AGO2* amplified samples remained after the filter (> 10 for all sample sets). However, the mRNA expression range and variability of *AGO2* was still large enough for an accurate analysis.

Correlation analysis of *AGO2* mRNA expression and genome-wide miRNA and mRNA expression (Spearman's coefficient test) was performed on three independent datasets to reduce the possibility of false positive results. In addition p-values were corrected for multiple testing using the FDR method. Normalized mRNA expression data as well as log2 transformed copy number data were used, for breast cancer (BRCA) samples. Samples considered were only those common between copy number variation data and mRNA expression data (N=1001). Genes expressed in fewer than 25% of clinical samples were removed from further analysis. Spearman correlation was considered for the remaining genes, against *PABPC1* expression, partial to *MYC* copy number, across all samples, with 100x bootstrap resampling used to generate confidence intervals for the correlation coefficient. Genes with a Spearman correlation coefficient-associated p-value < 0.05 (after Bonferroni correction), were considered for further pathway analysis.

##### *mRNA array for in-vitro experiments: experimental protocol and analysis*

HCC1806 cells were transfected with siRNA to *AGO2*, *PABPC1* or control. After incubation for 24 h in normoxia or hypoxia, mRNA extractions was carried out using the mirVANA extraction kit (Life Technologies), according to manufacturers instructions. Each experiment was performed on 4 separate occasions. Gene expression was considered for the HCC1806 cell line under control conditions of normoxia, and cell line with *PABPC1* knockdown by siRNA. Microarray analysis with Illumina HumanHT-12_V4 chip was performed, and gene expression values were obtained. Expression values were quantile normalized between samples, log2 transformed, and using the SamR R package (version 2.00) in R version 3.3.0, a list of differentially expressed genes was obtained between the control cell lines and those with *PABPC1* knockdown in normoxia via a two-sample paired test. Differentially expressed downregulated genes with FDR < 0.05 were considered for further pathway analysis.

##### *miRNA array for in-vitro experiments: experimental protocol and analysis*

HCC1806 cells were transfected with siRNA to *AGO2*, *PABPC1* or control. After incubation for 24 h in normoxia or hypoxia, miRNA was extracted using the Qiagen miRNA extraction kit, according to manufacturers instructions. Each experiment was performed on 4 separate occasions. .  Agilent 8x60K arrays were used (Version 21 with spike in probes).  Probe level signal data was summarised using median IOR [2].  Quantile normalisation was performed before proceeding with fold change analysis (LIMMA).

##### *Pathway analysis*

Pathway analysis was considered using the hypergeometric test for enrichment of genes identified as strongly correlated with *PABPC1* expression in clinical samples, as well as those differentially expressed in cell line data, with the genes of the (Kyoto Encyclopedia of Genes and Genomes) KEGG pathways. Singular enrichment analysis was carried out using Genecodis version 3.0 via the web interface with default settings. Subsequently, statistically significant ($p < 0.05$) overrepresented KEGG pathways in common for both the clinical samples and the cell line data were represented as chord diagrams, with the respective overlapping genes shown in each. Gene Set Enrichment Analysis (GSEA) was also performed using KEGG pathways (software.broadinstitute.org/gsea/index.jsp). When this was done not selection was applied to the gene prior entering the GSEA analysis to avoid bias.

##### *Western Blots*

Cell lysates were separated on 8, 10 or 12 % SDS-PAGE and transferred to a PVDF membrane. Primary antibodies against *CA9* (gift from J. Pastorek, Institute of Virology, Slovak Republic), *HIF1* (BD Biosciences, USA), *β-Actin* (A3854, Sigma, UK), *MYC* (9402, Cell Signalling, USA), Tubulin (Sigma, UK), *AGO1-4, PABPC1, Dicer and DROSHA* were assayed at 1:1000 unless otherwise stated.  Secondary horseradish peroxidase-linked antibodies were then utilised (Dako, UK). Chemiluminescence (Amersham, UK) was applied to detect immunoreactivity and then finally visualised using Image Quant LS4000 mini (GE Healthcare, UK).

##### *Quantitative PCR*

mRNA extractions were performed with Trizol (Invitrogen, CA, USA) as per  manufacturer's instructions. DNA was extracted using DNAzol (Invitrogen, CA, USA) according to the manufacturer's instruction. cDNA was synthesised using cDNA synthesis kit (Life

Technologies) according to manufacturers instructions. Applied Biosystems 7900HT Real-Time PCR System was used for real time analysis as described previously [50]. Primers for precursor miRNAs as previously described (Lo Sardo et al., 2017). Other primers available on request. Gene expression was normalised against the control genes RPL11 and β-Actin.

##### *miRNA TaqMan assay*

Small RNAs were isolated using the mirVana miRNA Isolation kit (Thermo Scientific). MicroRNA expression levels were determined using TaqMan Advanced miRNA Assays (Thermo Scientific) according to the manufacturer's instructions. The following miRNA assays were used: has-miR-25-3p (assay ID: 477994) , has-miR-25-5p (assay ID: 478786), miR-93-3p (assay ID: 002139), miR-93-5p (assay ID: 001090), has-miR-106b-3p (assay ID: 002380), hsa-miR-106b-5p (assay ID: 478412), and miR-5787 (assay ID: 480167), which was identified to be the least variable housekeeping miRNA in the microarray (HCC1806 cells), see above.
