## Supplementary Tables for "Dicer-to-Argonaute switch controls biogenesis of oncogenic miRNA"

**Table S1: Number of samples, genomic/transcriptomic assays and gene coverage for the clinical datasets used in the pan-cancer analysis.** All cancer samples had tumour and normal tissue matching genomic (DNA, CN) data (columns 3-4). These cancer samples had also transcriptomic (coding mRNA and miRNA) data (column 3), whilst the number of normal tissue samples with transcriptomic data is provided in column 5. Numbers in parenthesis correspond to additional samples where gene expression microarrays were available, used for differential expression analysis in GBM and OV. Number of detected genes is shown in column 6. See methods for details on data processing.

| **Tumor Acronym** | **Cancer Type** | **Number of cancer samples with matched genomic and transcriptomic data** | **Number of normal samples with genomic data** | **Number of normal samples with transcriptomic data** | **Number of Genes** |
|---|---|---|---|---|---|
| BLCA | Bladder Urothelial Carcinoma | 220 | 220 | 19 | 18078 |
| BRCA | Breast Invasive Carcinoma | 1001 | 1001 | 108 | 18253 |
| COADREAD | Colorectal Adenocarcinoma | 328 | 328 | 28 | 17922 |
| GBM | Glioblastoma Multiforme | 154 (528) | 154 | 0 (10) | 18364 (12042) |
| HNSC | Head and Neck Squamous Cell Carcinoma | 420 | 420 | 42 | 18213 |
| KIRC | Kidney Renal Clear Cell Carcinoma | 498 | 498 | 72 | 18154 |
| LUAD | Lung Adenocarcinoma | 486 | 486 | 58 | 18061 |

| | | | | | |
|---|---|---|---|---|---|
| LUSC | Lung Squamous Cell Carcinoma | 479 | 479 | 50 | 18279 |
| OV | Ovarian Serous Cystadenocarcinoma | 259 (585) | 259 | 0 (8) | 18730 (12042) |
| UCEC | Uterine Corpus Endometrial Carcinoma | 145 | 145 | 18 | 18075 |

**Table S2: Mature miRNA and immature pre-miRNA consistently associated with Cancer Hallmarks Gene Signatures across cancer types in a cross-validated GLM penalized regression (see Methods)**

| Hallmark | Fraction of immature miRNA also significant in mature miRNA analysis | Fraction of mature miRNA also significant in immature miRNA analysis | Immature miRNA associated with Hallmark Signature | Mature miRNA associated with Hallmark Signature |
|---|---|---|---|---|
| ***Positive Association between miRNA and Hallmarks Signatures*** | | | | |
| Hallmark: Epithelial Mesenchymal Transition | 0.76 | 0.75 | hsa-mir-21; hsa-mir-199a-1; hsa-mir-199a-2; hsa-mir-214; hsa-let-7i; hsa-mir-199b; hsa-mir-145; hsa-mir-379; hsa-mir-22; hsa-let-7e; hsa-mir-24-2; hsa-mir-708; hsa-mir-132; hsa-mir-152; hsa-mir-455; hsa-mir-409; hsa-mir-574 | hsa-mir-199a-5p; hsa-mir-21-5p; hsa-mir-379-5p; hsa-mir-199a-3p; hsa-mir-199b-3p; hsa-mir-214-5p; hsa-mir-22-3p; hsa-let-7i-3p; hsa-let-7e-5p; hsa-let-7i-5p; hsa-mir-145-5p; hsa-mir-708-5p; hsa-mir-152-3p; hsa-mir-132-3p; hsa-mir-574-3p; hsa-mir-181a-3p; hsa-mir-455-5p; hsa-mir-21-3p; hsa-mir-24-3p; hsa-let-7b-5p |

| | | | | |
|---|---|---|---|---|
| Invasiveness, Marsan 2014 | 0.50 | 0.75 | hsa-mir-21; hsa-mir-22; hsa-mir-223; hsa-mir-199a-1; hsa-mir-144; hsa-mir-145; hsa-let-7b; hsa-mir-199a-2; hsa-mir-24-2; hsa-let-7e; hsa-mir-146b; hsa-mir-125b-1; hsa-mir-150; hsa-mir-199b | hsa-mir-146b-3p; hsa-mir-21-5p; hsa-mir-27a-5p; hsa-mir-21-3p; hsa-mir-199a-5p; hsa-mir-223-3p; hsa-mir-22-3p; hsa-let-7b-5p; hsa-mir-199a-3p; hsa-mir-199b-3p; hsa-let-7b-3p; hsa-let-7e-5p |
| Hallmark: Oxidative Phosphorylation | 0.75 | 0.33 | hsa-mir-362; hsa-mir-1307; hsa-mir-34a; hsa-mir-18a | hsa-mir-362-5p; hsa-mir-1307-3p; hsa-mir-22-5p; hsa-let-7d-5p; hsa-mir-15b-3p; hsa-mir-34a-5p; hsa-mir-98-5p; hsa-mir-3607-3p; hsa-mir-192-5p |
| Hallmark: Reactive Oxygen Species Pathway | 0.60 | 0.56 | hsa-mir-193b; hsa-mir-22; hsa-mir-28; hsa-mir-24-2; hsa-mir-21 | hsa-mir-193b-3p; hsa-mir-22-3p; hsa-mir-21-3p; hsa-mir-22-5p; hsa-mir-21-5p; hsa-mir-29b-3p; hsa-mir-24-3p; hsa-mir-148a-5p; hsa-let-7a-3p |
| Hallmark: G2M Checkpoint | 0.65 | 0.93 | hsa-mir-15b; hsa-mir-130b; hsa-mir-7-1; hsa-mir-106b; hsa-mir-16-2; hsa-mir-18a; hsa-mir-155; hsa-mir-942; hsa-mir-20a; hsa-mir-93; hsa-mir-17; hsa-mir-92b; hsa-mir-769; hsa-mir-19b-1; hsa-mir-25; hsa-mir-629; hsa-let-7d | hsa-mir-15b-3p; hsa-mir-15b-5p; hsa-mir-106b-3p; hsa-mir-130b-5p; hsa-mir-130b-3p; hsa-mir-93-5p; hsa-mir-20a-5p; hsa-mir-17-5p; hsa-mir-92b-3p; hsa-mir-9-5p; hsa-mir-155-5p; hsa-mir-25-3p; hsa-mir-18a-5p; hsa-mir-19b-1-5p |
| Hallmark: PI3K AKT MTOR Signaling | 1.00 | 0.67 | hsa-mir-150; hsa-mir-142 | hsa-mir-142-3p; hsa-mir-27a-5p; hsa-mir-150-5p |
| Hallmark: Xenobiotic Metabolism | 0.75 | 0.75 | hsa-mir-22; hsa-mir-34a; hsa-mir-146a; hsa-mir-148a; hsa-mir-223; hsa-mir-142; hsa-mir-192; hsa-mir-29c | hsa-mir-22-3p; hsa-mir-34a-5p; hsa-mir-192-5p; hsa-mir-146a-5p; hsa-mir-223-3p; hsa-mir-29b-2-5p; hsa-mir-148a-3p; hsa-let-7c-5p |
| Hallmark: DNA Repair | 0.50 | 0.56 | hsa-mir-130b; hsa-mir-18a; hsa-mir-19b-1; hsa-mir-339; hsa-mir-362; hsa-mir-625; hsa-mir-130a; hsa-mir-93; hsa-mir-20a; hsa-mir-16-1 | hsa-mir-339-5p; hsa-mir-106b-5p; hsa-mir-17-3p; hsa-let-7d-5p; hsa-mir-19b-1-5p; hsa-mir-18a-5p; hsa-mir-15b-3p; hsa-mir-362-5p; hsa-mir-625-3p |

| | | | | |
|---|---|---|---|---|
| Hallmark: p53 Pathway | 0.78 | 0.82 | hsa-mir-34a; hsa-mir-21; hsa-mir-223; hsa-mir-27a; hsa-mir-22; hsa-mir-150; hsa-mir-205; hsa-mir-584; hsa-let-7b | hsa-mir-34a-5p; hsa-mir-27a-5p; hsa-mir-223-3p; hsa-mir-21-3p; hsa-mir-22-3p; hsa-mir-205-5p; hsa-mir-21-5p; hsa-mir-1976; hsa-mir-221-3p; hsa-mir-584-5p; hsa-mir-27a-3p |
| Hypoxia, Buffa 2010 | 0.43 | 0.50 | hsa-mir-210; hsa-mir-24-2; hsa-mir-98; hsa-mir-223; hsa-mir-27a; hsa-mir-193b; hsa-mir-425 | hsa-mir-210-3p; hsa-mir-24-3p; hsa-mir-21-3p; hsa-mir-193b-3p; hsa-mir-27a-5p; hsa-mir-584-5p |
| Hallmark: Angiogenesis | 0.73 | 0.64 | hsa-mir-22; hsa-let-7e; hsa-mir-21; hsa-mir-223; hsa-mir-455; hsa-mir-708; hsa-mir-145; hsa-mir-199a-1; hsa-mir-218-2; hsa-mir-24-2; hsa-mir-143 | hsa-let-7e-5p; hsa-mir-22-3p; hsa-mir-146b-3p; hsa-mir-223-3p; hsa-mir-199a-5p; hsa-mir-21-5p; hsa-mir-455-5p; hsa-mir-143-3p; hsa-mir-24-3p; hsa-mir-708-3p; hsa-mir-145-5p; hsa-mir-379-5p; hsa-mir-22-5p; hsa-mir-218-5p |
| Hallmark: Hypoxia | 0.82 | 0.71 | hsa-mir-210; hsa-mir-21; hsa-mir-223; hsa-let-7b; hsa-mir-132; hsa-mir-22; hsa-mir-24-2; hsa-mir-574; hsa-mir-144; hsa-mir-193a; hsa-let-7e | hsa-mir-210-3p; hsa-mir-21-5p; hsa-mir-223-3p; hsa-let-7b-5p; hsa-mir-22-3p; hsa-mir-21-3p; hsa-mir-27a-5p; hsa-mir-24-3p; hsa-mir-28-5p; hsa-mir-132-3p; hsa-mir-193a-5p; hsa-let-7e-5p; hsa-mir-574-3p; hsa-mir-28-3p |
| Angiogenesis, Desmedt 2008 | 0.67 | 0.50 | hsa-mir-210; hsa-let-7b; hsa-mir-148a | hsa-let-7b-3p; hsa-let-7b-5p; hsa-mir-27a-5p; hsa-let-7d-3p; hsa-let-7e-3p; hsa-mir-210-3p |
| Angiogenesis, Masiero 2013 | 0.80 | 0.69 | hsa-mir-126; hsa-mir-139; hsa-mir-140; hsa-mir-145; hsa-mir-150; hsa-mir-143; hsa-mir-218-2; hsa-mir-181a-1; hsa-mir-132; hsa-mir-144 | hsa-mir-126-3p; hsa-mir-126-5p; hsa-mir-139-5p; hsa-mir-140-3p; hsa-mir-150-5p; hsa-mir-218-5p; hsa-mir-132-3p; hsa-mir-139-3p; hsa-mir-143-3p; hsa-mir-145-5p; hsa-let-7c-5p; hsa-mir-145-3p; hsa-mir-144-5p; hsa-let-7b-5p; hsa-mir-22-3p; hsa-mir-181a-3p |

| | | | | |
|---|---|---|---|---|
| Hallmark: Apoptosis | 0.75 | 0.71 | hsa-mir-21; hsa-mir-223; hsa-mir-150; hsa-mir-155; hsa-mir-142; hsa-mir-22; hsa-mir-146b; hsa-mir-199a-1; hsa-mir-144; hsa-mir-132; hsa-let-7i; hsa-mir-199a-2 | hsa-mir-27a-5p; hsa-mir-21-5p; hsa-mir-150-5p; hsa-mir-155-5p; hsa-mir-223-3p; hsa-mir-21-3p; hsa-mir-144-5p; hsa-mir-22-3p; hsa-mir-146b-3p; hsa-mir-142-3p; hsa-let-7b-5p; hsa-let-7i-3p; hsa-mir-199a-5p; hsa-mir-199a-3p |
| Apoptosis, Desmedt 2008 | 0.63 | 0.86 | hsa-mir-181c; hsa-let-7c; hsa-let-7b; hsa-mir-181a-2; hsa-let-7a-2; hsa-let-7a-3; hsa-mir-25; hsa-mir-148a | hsa-mir-181c-3p; hsa-let-7b-5p; hsa-mir-25-3p; hsa-let-7a-2-3p; hsa-let-7c-5p; hsa-let-7b-3p; hsa-let-7e-3p |
| Proliferation, Desmedt 2008 | 0.90 | 0.67 | hsa-mir-130b; hsa-mir-18a; hsa-mir-15b; hsa-mir-93; hsa-mir-106b; hsa-mir-942; hsa-mir-16-2; hsa-mir-17; hsa-mir-301a; hsa-mir-25 | hsa-mir-15b-3p; hsa-mir-130b-5p; hsa-mir-93-5p; hsa-mir-18a-5p; hsa-mir-130b-3p; hsa-mir-106b-5p; hsa-mir-9-5p; hsa-mir-128-3p; hsa-mir-15b-5p; hsa-mir-17-5p; hsa-mir-362-5p; hsa-mir-942-5p; hsa-mir-17-3p; hsa-mir-25-3p; hsa-mir-301a-3p; hsa-mir-92b-3p; hsa-mir-98-5p; hsa-mir-19b-1-5p |
| Hallmark: KRAS Signaling Up | 1.00 | 0.72 | hsa-mir-150; hsa-mir-142; hsa-mir-155; hsa-mir-21; hsa-mir-146a; hsa-mir-223; hsa-mir-146b; hsa-mir-132; hsa-let-7i | hsa-mir-150-5p; hsa-mir-142-3p; hsa-mir-155-5p; hsa-mir-146b-3p; hsa-mir-223-3p; hsa-mir-146a-5p; hsa-mir-142-5p; hsa-mir-21-5p; hsa-let-7i-3p; hsa-mir-146b-5p; hsa-mir-132-3p; hsa-mir-199a-5p; hsa-mir-22-3p; hsa-mir-21-3p; hsa-let-7i-5p; hsa-mir-181a-3p; hsa-let-7c-5p; hsa-let-7e-5p |
| Hallmark: Inflammatory Response | 0.75 | 0.92 | hsa-mir-155; hsa-mir-150; hsa-mir-223; hsa-mir-142; hsa-mir-146b; hsa-mir-21; hsa-mir-146a; hsa-mir-22; hsa-mir-132; hsa-let-7i; hsa-mir-342; hsa-mir-144 | hsa-mir-155-5p; hsa-mir-223-3p; hsa-mir-150-5p; hsa-mir-142-3p; hsa-mir-142-5p; hsa-mir-146b-3p; hsa-mir-146a-5p; hsa-mir-146b-5p; hsa-mir-21-5p; hsa-mir-21-3p; hsa-let-7i-3p; hsa-mir-22-3p; hsa-mir-27a-5p |

| | | | | |
|---|---|---|---|---|
| Hallmark: IL2 STAT5 Signaling | 0.89 | 0.83 | hsa-mir-150; hsa-mir-155; hsa-mir-142; hsa-mir-223; hsa-mir-21; hsa-mir-146a; hsa-mir-132; hsa-mir-342; hsa-mir-146b | hsa-mir-150-5p; hsa-mir-155-5p; hsa-mir-223-3p; hsa-mir-146a-5p; hsa-mir-142-3p; hsa-mir-142-5p; hsa-mir-146b-3p; hsa-mir-21-5p; hsa-mir-21-3p; hsa-let-7i-3p; hsa-mir-132-3p; hsa-mir-30a-5p |
| Hallmark: IL6 JAK STAT3 Signaling | 0.88 | 0.71 | hsa-mir-142; hsa-mir-155; hsa-mir-150; hsa-mir-223; hsa-mir-146a; hsa-mir-146b; hsa-mir-21; hsa-mir-132 | hsa-mir-142-3p; hsa-mir-155-5p; hsa-mir-223-3p; hsa-mir-150-5p; hsa-mir-146b-3p; hsa-mir-142-5p; hsa-mir-146a-5p; hsa-mir-146b-5p; hsa-let-7i-3p; hsa-mir-21-3p; hsa-mir-21-5p; hsa-mir-27a-5p; hsa-let-7b-5p; hsa-mir-22-3p |
| Hallmark: TGF Beta Signaling | 0.57 | 0.64 | hsa-mir-21; hsa-let-7e; hsa-mir-23a; hsa-mir-132; hsa-mir-24-2; hsa-mir-144; hsa-let-7b | hsa-mir-199a-5p; hsa-let-7b-5p; hsa-mir-21-3p; hsa-let-7e-5p; hsa-mir-27a-5p; hsa-mir-21-5p; hsa-mir-2355-5p; hsa-let-7b-3p; hsa-mir-24-3p; hsa-mir-23a-3p; hsa-let-7e-3p |
| Hallmark: TNFa Signaling via NFKB | 0.69 | 1.00 | hsa-mir-223; hsa-mir-21; hsa-mir-150; hsa-mir-155; hsa-mir-146a; hsa-mir-132; hsa-mir-144; hsa-mir-142; hsa-mir-222; hsa-mir-22; hsa-mir-24-2; hsa-let-7b; hsa-mir-27a; hsa-mir-146b; hsa-mir-221; hsa-let-7f-1 | hsa-mir-223-3p; hsa-mir-21-3p; hsa-mir-27a-5p; hsa-mir-146a-5p; hsa-mir-150-5p; hsa-mir-21-5p; hsa-mir-146b-3p; hsa-mir-155-5p; hsa-mir-142-3p; hsa-mir-144-5p; hsa-mir-142-5p; hsa-let-7b-5p; hsa-mir-22-3p |
| Immune, Desmedt 2008 | 0.70 | 0.82 | hsa-mir-155; hsa-mir-150; hsa-mir-142; hsa-mir-146a; hsa-mir-146b; hsa-mir-342; hsa-let-7f-1; hsa-mir-21; hsa-mir-330; hsa-mir-221 | hsa-mir-155-5p; hsa-mir-150-5p; hsa-mir-142-3p; hsa-mir-146a-5p; hsa-mir-142-5p; hsa-mir-361-3p; hsa-mir-146b-5p; hsa-mir-146b-3p; hsa-let-7i-3p; hsa-mir-342-3p; hsa-mir-21-5p |
| ***Negative Association between miRNA and Hallmarks Signatures*** | | | | |

| | | | | |
|---|---|---|---|---|
| Hallmark: Epithelial Mesenchymal Transition | 0.33 | 0.67 | hsa-mir-362; hsa-mir-30b; hsa-mir-141; hsa-mir-339; hsa-mir-33a; hsa-mir-92a-2; hsa-mir-660; hsa-mir-423; hsa-mir-769; hsa-mir-942; hsa-mir-148b; hsa-mir-532 | hsa-mir-29c-5p; hsa-mir-362-5p; hsa-mir-30b-5p; hsa-mir-92a-3p; hsa-mir-942-5p; hsa-mir-423-3p |
| Invasiveness, Marsan 2014 | 0.44 | 0.40 | hsa-mir-362; hsa-mir-130b; hsa-let-7f-1; hsa-mir-16-1; hsa-mir-17; hsa-mir-16-2; hsa-mir-33a; hsa-mir-30b; hsa-mir-19b-1 | hsa-mir-362-5p; hsa-mir-423-3p; hsa-mir-17-3p; hsa-mir-16-5p; hsa-mir-192-5p; hsa-let-7a-3p; hsa-mir-130b-3p; hsa-mir-107; hsa-mir-15a-5p; hsa-mir-30b-5p |
| Hallmark: Oxidative Phosphorylation | 0.14 | 0.33 | hsa-mir-574; hsa-let-7e; hsa-mir-26a-2; hsa-mir-101-1; hsa-mir-23a; hsa-mir-193a; hsa-mir-331 | hsa-let-7e-5p; hsa-let-7b-5p; hsa-let-7b-3p |
| Hallmark: Reactive Oxygen Species Pathway | 0.33 | 0.50 | hsa-mir-181b-1; hsa-mir-92a-1; hsa-mir-30d | hsa-mir-181b-5p; hsa-mir-30d-5p |
| Hallmark: G2M Checkpoint | 0.64 | 0.59 | hsa-mir-29a; hsa-mir-22; hsa-mir-26b; hsa-mir-125a; hsa-mir-30d; hsa-mir-30b; hsa-mir-26a-2; hsa-mir-328; hsa-mir-29c; hsa-mir-34a; hsa-mir-101-1; hsa-mir-574; hsa-mir-30c-2; hsa-mir-101-2 | hsa-mir-125a-5p; hsa-mir-29c-5p; hsa-mir-29b-2-5p; hsa-mir-29a-3p; hsa-mir-26b-5p; hsa-mir-22-5p; hsa-mir-22-3p; hsa-mir-30b-5p; hsa-mir-30d-5p; hsa-mir-30c-5p; hsa-mir-26a-5p; hsa-mir-182-5p; hsa-mir-34a-5p; hsa-mir-181c-5p; hsa-mir-101-3p; hsa-mir-328-3p; hsa-mir-335-5p |
| Hallmark: PI3K AKT MTOR Signaling | 0.88 | 0.53 | hsa-mir-125a; hsa-let-7b; hsa-let-7e; hsa-mir-30b; hsa-mir-19b-2; hsa-let-7d; hsa-mir-29a; hsa-mir-15a | hsa-let-7b-5p; hsa-let-7a-3p; hsa-let-7e-3p; hsa-let-7b-3p; hsa-let-7d-5p; hsa-mir-30b-5p; hsa-let-7e-5p; hsa-mir-17-5p; hsa-let-7f-1-3p; hsa-mir-15a-5p; hsa-mir-24-2-5p; hsa-mir-29a-3p; hsa-mir-26b-5p; hsa-mir-20a-5p; hsa-mir- |

| | | | | |
|---|---|---|---|---|
| | | | | 101-3p; hsa-mir-125a-5p; hsa-mir-20a-3p |
| Hallmark: Xenobiotic Metabolism | 0.45 | 0.58 | hsa-mir-106b; hsa-mir-17; hsa-mir-20a; hsa-mir-7-1; hsa-mir-92a-2; hsa-mir-93; hsa-mir-19b-1; hsa-mir-769; hsa-mir-16-2; hsa-mir-24-2; hsa-mir-19b-2 | hsa-mir-17-5p; hsa-mir-92a-3p; hsa-mir-20a-5p; hsa-mir-19b-1-5p; hsa-mir-106b-3p; hsa-let-7a-3p; hsa-mir-106b-5p; hsa-mir-24-2-5p; hsa-let-7f-1-3p; hsa-mir-20a-3p; hsa-let-7d-3p; hsa-mir-16-5p |
| Hallmark: DNA Repair | 0.40 | 0.18 | hsa-mir-574; hsa-let-7e; hsa-mir-26a-2; hsa-mir-197; hsa-mir-22 | hsa-let-7b-3p; hsa-mir-29b-2-5p; hsa-let-7e-5p; hsa-mir-326; hsa-mir-26a-5p; hsa-mir-22-3p; hsa-let-7d-3p; hsa-let-7b-5p; hsa-mir-144-5p; hsa-mir-23a-5p; hsa-mir-23b-3p |
| Hallmark: p53 Pathway | 0.83 | 0.75 | hsa-mir-130b; hsa-mir-25; hsa-mir-17; hsa-mir-106b; hsa-mir-93; hsa-mir-769 | hsa-mir-25-3p; hsa-mir-769-5p; hsa-mir-130b-3p; hsa-mir-106a-5p; hsa-mir-17-5p; hsa-mir-335-3p; hsa-mir-93-5p; hsa-mir-130b-5p |
| Hypoxia, Buffa 2010 | 0.44 | 0.42 | hsa-mir-101-1; hsa-mir-30e; hsa-mir-328; hsa-mir-30c-2; hsa-let-7g; hsa-mir-26a-2; hsa-mir-125a; hsa-mir-30a; hsa-mir-23b | hsa-mir-30e-5p; hsa-mir-181c-5p; hsa-mir-101-3p; hsa-mir-125a-5p; hsa-mir-26a-5p; hsa-mir-29c-5p; hsa-mir-30e-3p; hsa-mir-30a-3p; hsa-let-7b-3p; hsa-mir-29b-2-5p; hsa-mir-30c-5p; hsa-mir-328-3p |
| Hallmark: Angiogenesis | 1.00 | 0.25 | hsa-mir-362 | hsa-mir-362-5p; hsa-mir-361-5p; hsa-mir-29c-5p; hsa-mir-16-5p |
| Hallmark: Hypoxia | 0.67 | 0.25 | hsa-mir-362; hsa-mir-19b-1; hsa-mir-335 | hsa-mir-362-5p; hsa-mir-19b-1-5p; hsa-mir-106b-5p; hsa-mir-17-5p; hsa-mir-16-5p; hsa-let-7a-3p; hsa-mir-25-3p; hsa-mir-29c-5p |

| | | | | |
|---|---|---|---|---|
| Angiogenesis, Desmedt 2008 | 0.00 | 0.00 | hsa-let-7f-1; hsa-mir-98; hsa-let-7a-2 | hsa-let-7d-5p |
| Angiogenesis, Masiero 2013 | 0.17 | 1.00 | hsa-mir-141; hsa-mir-210; hsa-mir-33a; hsa-mir-339; hsa-mir-7-1; hsa-mir-16-2 | hsa-mir-210-3p |
| Hallmark: Apoptosis | 0.57 | 0.80 | hsa-mir-30b; hsa-mir-20a; hsa-mir-362; hsa-mir-769; hsa-mir-19b-1; hsa-mir-339; hsa-mir-19b-2 | hsa-mir-30b-5p; hsa-mir-362-5p; hsa-mir-19b-1-5p; hsa-mir-769-5p; hsa-mir-92a-3p |
| Apoptosis, Desmedt 2008 | 0.33 | 0.67 | hsa-mir-30b; hsa-mir-29a; hsa-mir-193b; hsa-let-7f-2; hsa-let-7d; hsa-mir-30c-2 | hsa-mir-30b-5p; hsa-mir-29a-3p; hsa-let-7a-3p |
| Proliferation, Desmedt 2008 | 0.40 | 0.57 | hsa-mir-26a-2; hsa-mir-101-1; hsa-mir-574; hsa-mir-30e; hsa-mir-125a; hsa-mir-22; hsa-mir-30d; hsa-mir-331; hsa-mir-26b; hsa-mir-140 | hsa-mir-29b-2-5p; hsa-mir-574-3p; hsa-mir-26a-5p; hsa-mir-101-3p; hsa-mir-30d-5p; hsa-mir-125a-5p; hsa-mir-22-3p |
| Hallmark: KRAS Signaling Up | 0.45 | 0.45 | hsa-mir-362; hsa-mir-16-2; hsa-mir-30b; hsa-mir-339; hsa-mir-769; hsa-mir-33a; hsa-mir-93; hsa-mir-92a-2; hsa-mir-942; hsa-mir-17; hsa-mir-629 | hsa-mir-362-5p; hsa-mir-92a-3p; hsa-let-7a-3p; hsa-mir-30b-5p; hsa-mir-93-5p; hsa-mir-942-5p; hsa-mir-769-5p; hsa-mir-106b-5p; hsa-let-7d-5p; hsa-mir-423-3p; hsa-mir-16-1-3p |
| Hallmark: Inflammatory Response | 0.62 | 0.50 | hsa-mir-30b; hsa-mir-769; hsa-mir-33a; hsa-mir-20a; hsa-mir-324; hsa-mir-92a-2; hsa-mir-19b-2; hsa-mir-362; hsa-mir-423; hsa-mir-93; hsa-mir-96; hsa-mir-130b; hsa-mir-30c-2 | hsa-mir-30b-5p; hsa-mir-92a-3p; hsa-let-7a-3p; hsa-mir-345-5p; hsa-mir-106b-5p; hsa-mir-423-3p; hsa-mir-769-5p; hsa-mir-96-5p; hsa-mir-361-5p; hsa-mir-362-5p; hsa-mir-20a-5p; hsa-mir-19b-3p; hsa-mir-1180-3p; hsa-mir-93-5p; hsa-mir-1296-5p; hsa-mir-33a-5p |

| | | | | |
|---|---|---|---|---|
| Hallmark: IL2 STAT5 Signaling | 0.06 | 0.50 | hsa-mir-1307; hsa-mir-33a; hsa-mir-17; hsa-mir-96; hsa-mir-339; hsa-mir-141; hsa-mir-19b-2; hsa-let-7e; hsa-mir-16-2; hsa-mir-19b-1; hsa-mir-362; hsa-mir-92a-2; hsa-mir-30b; hsa-mir-15a; hsa-let-7a-3; hsa-let-7a-2 | hsa-let-7a-3p; hsa-mir-15a-5p |
| Hallmark: IL6 JAK STAT3 Signaling | 0.40 | 0.50 | hsa-mir-30b; hsa-mir-362; hsa-mir-324; hsa-mir-769; hsa-mir-92a-2; hsa-mir-33a; hsa-mir-423; hsa-mir-93; hsa-mir-335; hsa-mir-96 | hsa-mir-30b-5p; hsa-mir-769-5p; hsa-mir-92a-3p; hsa-mir-106b-5p; hsa-let-7a-3p; hsa-mir-423-3p; hsa-mir-19b-3p; hsa-mir-324-3p |
| Hallmark: TGF Beta Signaling | 0.67 | 0.25 | hsa-mir-362; hsa-mir-1307; hsa-mir-148a | hsa-mir-362-5p; hsa-mir-339-5p; hsa-mir-17-3p; hsa-mir-148a-5p; hsa-mir-15a-5p; hsa-mir-16-5p; hsa-mir-19b-1-5p; hsa-let-7d-5p |
| Hallmark: TNFa Signaling via NFKB | 0.00 | 1.00 | | hsa-mir-362-5p |
| Immune, Desmedt 2008 | 0.50 | 0.60 | hsa-mir-130b; hsa-mir-92a-2; hsa-mir-20a; hsa-mir-337; hsa-mir-30b; hsa-mir-93 | hsa-mir-92a-3p; hsa-mir-30b-5p; hsa-mir-1296-5p; hsa-mir-130b-3p; hsa-mir-20a-5p |

**Table S3: miRNA biogenesis genes.**

Full list of the 43 miRNA Processing Machinery Gene considered in this study with symbols, gene names, IDs and literature source.

| **Gene symbol** | **Full Gene Name** | **Ensembl ID** | **Comments** | **Reference (Author, PMID, Citation)** |
|---|---|---|---|---|

| | | | | |
|---|---|---|---|---|
| ADAR | adenosine deaminase, RNA specific | ENSG00000160710 | Blocks dicer/drosha processing by changing RNA sequence | Krol et al, 20661255, [1] |
| ADARB1 | adenosine deaminase, RNA specific B1 | ENSG00000197381 | Editing of specific microRNAs | Buffa et al, 21737487,[2] |
| AGO1 | argonaute 1, miRISC catalytic component | ENSG00000092847 | RISC | Meister G, 23732335,[3] |
| AGO2 | argonaute 2, miRISC catalytic component | ENSG00000123908 | RISC | Meister G, 23732335,[3] |
| AGO3 | argonaute 3, miRISC catalytic component | ENSG00000126070 | RISC | Meister G, 23732335,[3] |
| AGO4 | argonaute 4, miRISC catalytic component | ENSG00000134698 | RISC | Meister G, 23732335,[3] |
| CNOT1 | CCR4-NOT transcription complex subunit 1 | ENSG00000125107 | Interact with AGO1/2 | Krol et al, 20661255, [1] |
| DCP2 | decapping mRNA 2 | ENSG00000172795 | Interact with AGO2 | Krol et al, 20661255, [1] |
| DDX17 | DEAD-box helicase 17 | ENSG00000100201 | ATPase activated by several RNA species | Krol et al, 20661255, [1] |
| DDX20 | DEAD-box helicase 20 | ENSG00000064703 | putative RNA helicase | Krol et al, 20661255, [1] |
| DDX5 | DEAD-box helicase 5 | ENSG00000108654 | putative RNA helicases | Krol et al, 20661255, [1] |
| DDX6 | DEAD-box helicase 6 | ENSG00000110367 | Interacts with AGO1 and AGO2; depletion relieves miRNA mediated repression | Krol et al, 20661255, [1] |
| DGCR8 | DGCR8, microprocessor complex subunit | ENSG00000128191 | Microprocessor | Buffa et al, 21737487,[2] |
| DHX9 | DExH-box helicase 9 | ENSG00000135829 | helicase enzyme involed in unwinding of double-stranded RNA and DNA-RNA complexes. Contributes to the formation of miRISC. | Lee et al, 27034008, [4] |
| DICER1 | dicer 1, ribonuclease III | ENSG00000100697 | DICER | Krol et al, 20661255 , [1] |
| DND1 | DND microRNA-mediated repression inhibitor 1 | ENSG00000256453 | Blocks miRNA/mRNA interaction to protects mRNAs from miRNA repression | Krol et al, 20661255 , [1] |
| DROSHA | drosha ribonuclease III | ENSG00000113360 | Microprocessor | Krol et al, 20661255 , [1] |
| ELAVL1 | ELAV like RNA binding protein 1 | ENSG00000066044 | Relieves miRNA mediated repression in response to stress | Lu et al, 25533351, [5] |

| | | | | |
|---|---|---|---|---|
| ESR1 | estrogen receptor 1 | ENSG00000091831 | p68 and p72 interaction | Buffa et al, 21737487,[2] |
| ESR2 | estrogen receptor 2 | ENSG00000140009 | p68 and p72 interaction | Buffa et al, 21737487,[2] |
| FXR1 | FMR1 autosomal homolog 1 | ENSG00000114416 | Component of RISC complex, activating transcription. Required for AGO2 to activate translation of Tnf! mRNA in growth arrested cells | Gessert et al, 20197067, [6]<br>Anderson et al, 20029446, [7]<br>Shobha et al, 17382880, [8] |
| GEMIN4 | gem nuclear organelle associated protein 4 | ENSG00000179409 | Part of Gemini bodies | Nelson et al, 14970384, [9] |
| HNRNPA1 | heterogeneous nuclear ribonucleoprotein A1 | ENSG00000135486 | DROSHA component | Krol et al, 20661255 , [1] |
| ILF3 | interleukin enhancer binding factor 3 | ENSG00000129351 | Reduces DGCR8 accessibility (TF) | Krol et al, 20661255 , [1] |
| IPO8 | importin 8 | ENSG00000133704 | importin-alpha/beta complex and the GTPase Ran mediate nuclear import of proteins | Hock et al, 17932509 [10] |
| KHSRP | KH-type splicing regulatory protein | ENSG00000088247 | Component of Drosha and Dicer complexes | Krol et al, 20661255, [1] |
| MAPKAPK2 | mitogen-activated protein kinase-activated protein kinase 2 | ENSG00000162889 | Catalyses Ser387 phosphorylation of AGO2 | Krol et al, 20661255, [1]<br>Li Z et al, 25011539, [11] |
| MOV10 | Mov10 RISC complex RNA helicase | ENSG00000155363 | Interacts with AGO1 and AGO2 | Krol et al, 20661255, [1] |
| PABPC1 | poly(A) binding protein cytoplasmic 1 | ENSG00000070756 | Interacts with GW182 (AGO2) | Krol et al, 20661255, [1] |
| PIWIL1 | piwi like RNA-mediated gene silencing 1 | ENSG00000275051 | PIWI subfamily of Argonaute proteins | Meister G, 23732335,[3] |
| PRKRA | protein activator of interferon induced protein kinase EIF2AK2 | ENSG00000180228 | DICER | Buffa et al, 21737487,[2] |
| RAN | RAN, member RAS oncogene family | ENSG00000132341 | Transport | Buffa et al, 21737487, [2] |
| RBM4 | RNA binding motif protein 4 | ENSG00000173933 | Interacts with AGO1 and AGO2; depletion reduces miRNA mediated repression | Krol et al, 20661255 , [1] |
| SMAD1 | SMAD family member 1 | ENSG00000170365 | Interacting with p68 co-activators of Drosha | Krol et al, 20661255, [1] |

| | | | | |
|---|---|---|---|---|
| SMAD3 | SMAD family member 3 | ENSG00000166949 | Interacting with p68 co-activators of Drosha | Krol et al, 20661255, [1] |
| SMAD5 | SMAD family member 5 | ENSG00000113658 | Interacting with p68 co-activators of Drosha | Krol et al, 20661255, [1] |
| SNIP1 | Smad nuclear interacting protein 1 | ENSG00000163877 | Regulates maturation of miR subset when complexed with DROSHA | Krol et al, 20661255, [1] |
| SRSF1 | serine and arginine rich splicing factor 1 | ENSG00000136450 | Splicing factor for Drosha-mediated cleavage | Krol et al, 20661255, [1] |
| TARBP1 | TAR (HIV-1) RNA binding protein 1 | ENSG00000059588 | DICER | Buffa et al, 21737487,[2] |
| TARBP2 | TARBP2, RISC loading complex RNA binding subunit | ENSG00000139546 | DICER | Buffa et al, 21737487,[2] |
| TNRC6A | trinucleotide repeat containing 6A | ENSG00000090905 | Part of AGO2 complex | Nishi et al, 23150874 [12] |
| TP53 | tumor protein p53 | ENSG00000141510 | Interacting with p68 facilitates Drosha complex assembly | Krol et al, 20661255 , [1] |
| TRIM32 | tripartite motif containing 32 | ENSG00000119401 | Binding to miRISC, enhancing microRNA activity | Buffa et al, 21737487,[2] |
| TSN | translin | ENSG00000211460 | RISC loading function | Meister G, 23732335,[3] |
| UPF1 | UPF1, RNA helicase and ATPase | ENSG00000005007 | Role in mRNA surveillance and nuclear export | Meister G, 23732335,[3] |
| XPO5 | exportin 5 | ENSG00000124571 | Transport | Krol et al, 20661255 , [1] |

**Table S4: Pan-cancer somatic mutation frequency of miRNA biogenesis genes.**

Data are expressed as percentages (0-100%) of respective cohorts.

| | BLCA | BRCA | COADREAD | GBM | HNSC | KIRC | LUAD | LUSC | OV | UCEC | Pan-cancer Mean % |
|---|---|---|---|---|---|---|---|---|---|---|---|
| **n** | 28 | 986 | 227 | 283 | 306 | 491 | 248 | 177 | 316 | 248 | |
| **Two-sided Proportion Test P-value** | 0.3203 | 1.0000 | 0.4078 | 1.0000 | 0.8268 | 1.0000 | 0.2553 | 0.5159 | 1.0000 | 0.0775 | |
| EIF2C3 | 0.00 | 0.41 | 2.64 | 0.35 | 2.29 | 0.61 | 1.21 | 1.69 | 0.00 | 4.44 | 1.52 |
| PRKRA | 0.00 | 0.20 | 1.76 | 0.35 | 0.00 | 0.20 | 0.40 | 0.56 | 0.00 | 1.21 | 0.52 |
| ESR1 | 0.00 | 0.71 | 4.85 | 0.35 | 0.98 | 0.00 | 0.40 | 3.39 | 0.00 | 4.44 | 1.68 |
| EIF2C4 | 0.00 | 0.41 | 2.64 | 0.00 | 1.31 | 0.41 | 0.40 | 0.56 | 0.32 | 2.02 | 0.90 |
| TARBP1 | 0.00 | 1.22 | 2.20 | 0.35 | 1.63 | 1.22 | 3.23 | 3.95 | 0.63 | 4.84 | 2.14 |
| RNASEN | 0.00 | 0.00 | 0.00 | 0.71 | 0.65 | 0.20 | 3.23 | 7.34 | 0.63 | 0.00 | 1.42 |
| ARS2 | 0.00 | 0.00 | 0.00 | 0.00 | 0.00 | 0.00 | 0.00 | 0.00 | 0.00 | 0.00 | 0.00 |
| ESR2 | 0.00 | 0.41 | 0.88 | 1.06 | 0.33 | 0.20 | 1.21 | 1.13 | 0.00 | 1.61 | 0.76 |
| ADARB1 | 0.00 | 0.20 | 0.44 | 0.00 | 0.33 | 0.00 | 2.42 | 0.00 | 0.00 | 1.61 | 0.56 |
| EIF2C2 | 0.00 | 0.91 | 1.32 | 0.00 | 1.31 | 0.61 | 2.42 | 0.56 | 0.32 | 3.63 | 1.23 |
| DGCR8 | 0.00 | 0.41 | 1.32 | 0.35 | 0.65 | 0.20 | 0.40 | 1.69 | 0.00 | 3.63 | 0.96 |
| XPO5 | 3.57 | 0.71 | 0.44 | 0.00 | 0.33 | 0.41 | 1.21 | 1.13 | 0.00 | 2.82 | 0.78 |
| DICER1 | 0.00 | 0.81 | 3.08 | 1.41 | 2.29 | 0.41 | 3.23 | 6.21 | 0.32 | 9.27 | 3.00 |
| TARBP2 | 0.00 | 0.61 | 1.32 | 0.35 | 0.33 | 0.00 | 0.40 | 1.13 | 0.00 | 2.02 | 0.68 |
| TRIM32 | 3.57 | 0.30 | 2.20 | 0.00 | 0.65 | 0.00 | 0.40 | 2.26 | 0.00 | 5.24 | 1.23 |
| EIF2C1 | 3.57 | 0.71 | 3.96 | 0.00 | 1.96 | 0.20 | 2.02 | 1.13 | 0.63 | 1.61 | 1.36 |
| RAN | 0.00 | 0.10 | 0.88 | 0.00 | 0.33 | 0.20 | 0.00 | 0.00 | 0.00 | 0.40 | 0.21 |
| DDX17 | 0.00 | 0.30 | 2.64 | 0.71 | 0.00 | 0.20 | 1.21 | 0.56 | 0.00 | 2.02 | 0.85 |
| TSN | 0.00 | 0.20 | 0.00 | 0.00 | 0.00 | 0.00 | 0.40 | 0.56 | 0.00 | 1.61 | 0.31 |
| TNRC6 | 0.00 | 0.00 | 0.00 | 0.00 | 0.00 | 0.00 | 0.00 | 0.00 | 0.00 | 0.00 | 0.00 |

| | | | | | | | | | | | |
|---|---|---|---|---|---|---|---|---|---|---|---|
| DDX20 | 0.00 | 0.41 | 1.76 | 0.35 | 0.98 | 0.81 | 0.40 | 0.00 | 0.00 | 1.61 | 0.70 |
| IPO8 | 7.14 | 0.51 | 4.41 | 0.35 | 0.65 | 0.20 | 4.44 | 1.69 | 0.00 | 6.05 | 2.03 |
| DHX9 | 7.14 | 0.81 | 1.76 | 0.00 | 1.63 | 0.61 | 1.21 | 1.13 | 0.63 | 4.03 | 1.31 |
| RENT1 | 0.00 | 0.00 | 0.00 | 0.00 | 0.00 | 0.00 | 0.00 | 0.00 | 0.00 | 0.00 | 0.00 |
| DDX5 | 0.00 | 0.51 | 0.88 | 1.41 | 0.00 | 0.41 | 1.61 | 0.56 | 0.63 | 2.82 | 0.98 |
| GEMIN4 | 0.00 | 0.41 | 1.32 | 0.35 | 1.31 | 0.41 | 0.40 | 0.56 | 0.63 | 2.02 | 0.82 |
| SMAD1 | 0.00 | 0.10 | 0.88 | 0.00 | 0.65 | 0.00 | 1.21 | 1.69 | 0.32 | 0.81 | 0.63 |
| HNRNPA1 | 0.00 | 0.20 | 1.32 | 0.00 | 0.33 | 0.00 | 0.81 | 1.69 | 0.32 | 1.21 | 0.65 |
| FXR1 | 0.00 | 0.91 | 2.20 | 0.35 | 1.96 | 0.00 | 0.81 | 2.26 | 0.00 | 4.84 | 1.48 |
| ADAR | 0.00 | 0.41 | 3.08 | 0.00 | 0.65 | 0.81 | 3.23 | 1.69 | 1.27 | 2.82 | 1.55 |
| ELAVL1 | 0.00 | 0.41 | 1.32 | 0.35 | 0.33 | 0.00 | 1.21 | 0.56 | 0.00 | 3.23 | 0.82 |
| MOV10 | 3.57 | 0.61 | 1.76 | 0.71 | 0.65 | 0.20 | 2.02 | 1.13 | 0.95 | 2.02 | 1.12 |
| PABPC1 | 0.00 | 0.20 | 1.32 | 0.00 | 0.00 | 1.63 | 1.21 | 2.26 | 0.63 | 3.23 | 1.16 |
| SRSF1 | 0.00 | 0.00 | 0.00 | 0.00 | 0.00 | 0.00 | 0.00 | 0.00 | 0.00 | 0.00 | 0.00 |
| SNIP1 | 0.00 | 0.00 | 1.32 | 0.00 | 0.00 | 0.20 | 1.61 | 2.82 | 0.00 | 1.21 | 0.80 |
| DCP2 | 0.00 | 0.20 | 2.20 | 0.00 | 0.65 | 0.00 | 0.00 | 0.00 | 0.00 | 3.63 | 0.74 |
| ILF3 | 0.00 | 0.61 | 1.76 | 0.71 | 0.00 | 0.20 | 1.61 | 0.56 | 0.63 | 1.21 | 0.81 |
| CNOT1 | 0.00 | 1.52 | 4.85 | 0.35 | 3.59 | 1.83 | 3.23 | 3.95 | 1.27 | 8.87 | 3.27 |
| SMAD5 | 0.00 | 0.20 | 0.44 | 0.00 | 0.65 | 0.00 | 0.81 | 0.00 | 0.00 | 4.03 | 0.68 |
| SMAD3 | 0.00 | 0.41 | 3.52 | 0.00 | 0.65 | 0.20 | 1.21 | 0.56 | 0.00 | 2.82 | 1.04 |
| RBM4 | 0.00 | 0.10 | 1.32 | 0.35 | 0.98 | 0.00 | 0.00 | 0.56 | 0.32 | 1.21 | 0.54 |
| KHSRP | 0.00 | 0.41 | 0.88 | 0.71 | 0.65 | 0.00 | 0.00 | 0.56 | 0.00 | 2.02 | 0.58 |
| MAPKAPK2 | 0.00 | 0.10 | 1.32 | 0.35 | 0.00 | 0.00 | 0.81 | 0.56 | 0.00 | 2.42 | 0.62 |
| PIWIL1 | 0.00 | 1.72 | 2.64 | 0.71 | 2.61 | 0.41 | 4.44 | 4.52 | 0.32 | 5.65 | 2.56 |
| FMR1 | 0.00 | 0.71 | 1.76 | 0.00 | 0.65 | 0.61 | 2.42 | 2.26 | 0.63 | 6.45 | 1.72 |
| DROSHA | 0.00 | 0.00 | 0.00 | 0.00 | 0.00 | 0.00 | 0.00 | 0.00 | 0.00 | 0.00 | 0.00 |
| TNRC6A | 3.57 | 1.12 | 2.64 | 0.00 | 3.27 | 0.61 | 3.63 | 3.39 | 1.27 | 8.47 | 2.71 |
| **Average** | 0.68 | 0.45 | 1.69 | 1.12 | 0.81 | 0.30 | 1.33 | 1.46 | 0.27 | 2.87 | 1.05 |
| **Standard Deviation** | 1.77 | 0.40 | 1.27 | 5.82 | 0.89 | 0.41 | 1.25 | 1.64 | 0.38 | 2.32 | 0.77 |

| Maximum | 7.14 | 1.72 | 4.85 | 40.08 | 3.59 | 1.83 | 4.44 | 7.34 | 1.27 | 9.27 | 3.27 |
|---|---|---|---|---|---|---|---|---|---|---|---|

**Table S5. Pan-cancer copy number gain/amplification frequency at miRNA biogenesis genes' locations**

Data are expressed as fraction (0-1) of respective cohorts (Table S2). White-red color scale from lowest to highest frequency.

| Gene | BLCA | BRCA | COADREAD | GBM | HNSC | KIRC | LUAD | LUSC | OV | UCEC |
|---|---|---|---|---|---|---|---|---|---|---|
| EIF2C3 | 0.234973 | 0.115546 | 0.049231 | 0.189542 | 0.148876 | 0.022495 | 0.267261 | 0.131313 | 0.416988 | 0.144231 |
| PRKRA | 0.131148 | 0.095588 | 0.233846 | 0.065359 | 0.205056 | 0.155419 | 0.305122 | 0.323232 | 0.409266 | 0.259615 |
| ESR1 | 0.081967 | 0.155462 | 0.175385 | 0.03268 | 0.098315 | 0.01636 | 0.089087 | 0.189394 | 0.162162 | 0.230769 |
| EIF2C4 | 0.234973 | 0.112395 | 0.049231 | 0.183007 | 0.143258 | 0.022495 | 0.267261 | 0.123737 | 0.393822 | 0.144231 |
| TARBP1 | 0.344262 | 0.732143 | 0.212308 | 0.150327 | 0.266854 | 0.108384 | 0.657016 | 0.525253 | 0.579151 | 0.480769 |
| ESR2 | 0.224044 | 0.132353 | 0.08 | 0.045752 | 0.317416 | 0.03272 | 0.258352 | 0.25 | 0.119691 | 0.163462 |
| ADARB1 | 0.360656 | 0.22584 | 0.058462 | 0.156863 | 0.078652 | 0.102249 | 0.198218 | 0.116162 | 0.247104 | 0.192308 |
| EIF2C2 | 0.568306 | 0.582983 | 0.593846 | 0.137255 | 0.724719 | 0.151329 | 0.579065 | 0.669192 | 0.760618 | 0.471154 |
| DGCR8 | 0.251366 | 0.12605 | 0.055385 | 0.084967 | 0.264045 | 0.0818 | 0.175947 | 0.507576 | 0.146718 | 0.134615 |
| XPO5 | 0.213115 | 0.234244 | 0.258462 | 0.058824 | 0.168539 | 0.026585 | 0.376392 | 0.290404 | 0.401544 | 0.288462 |
| DICER1 | 0.213115 | 0.135504 | 0.067692 | 0.052288 | 0.323034 | 0.030675 | 0.233853 | 0.290404 | 0.216216 | 0.221154 |
| TARBP2 | 0.240437 | 0.194328 | 0.187692 | 0.098039 | 0.160112 | 0.239264 | 0.289532 | 0.313131 | 0.332046 | 0.201923 |
| TRIM32 | 0.142077 | 0.141807 | 0.16 | 0.156863 | 0.33427 | 0.026585 | 0.093541 | 0.207071 | 0.150579 | 0.028846 |
| EIF2C1 | 0.234973 | 0.113445 | 0.049231 | 0.183007 | 0.143258 | 0.022495 | 0.267261 | 0.126263 | 0.413127 | 0.144231 |
| RAN | 0.251366 | 0.180672 | 0.169231 | 0.091503 | 0.13764 | 0.237219 | 0.249443 | 0.255051 | 0.285714 | 0.125 |
| DDX17 | 0.163934 | 0.109244 | 0.036923 | 0.098039 | 0.227528 | 0.08589 | 0.144766 | 0.482323 | 0.081081 | 0.153846 |

| | | | | | | | | | | |
|---|---|---|---|---|---|---|---|---|---|---|
| TSN | 0.169399 | 0.085084 | 0.215385 | 0.071895 | 0.176966 | 0.145194 | 0.302895 | 0.356061 | 0.250965 | 0.25 |
| DDX20 | 0.218579 | 0.146008 | 0.046154 | 0.163399 | 0.073034 | 0.03681 | 0.229399 | 0.118687 | 0.30888 | 0.173077 |
| IPO8 | 0.333333 | 0.255252 | 0.2 | 0.111111 | 0.300562 | 0.237219 | 0.327394 | 0.477273 | 0.525097 | 0.192308 |
| DHX9 | 0.382514 | 0.732143 | 0.236923 | 0.183007 | 0.255618 | 0.114519 | 0.694878 | 0.520202 | 0.57529 | 0.557692 |
| DDX5 | 0.497268 | 0.407563 | 0.221538 | 0.117647 | 0.207865 | 0.07771 | 0.494432 | 0.449495 | 0.277992 | 0.192308 |
| GEMIN4 | 0.131148 | 0.068277 | 0.049231 | 0.071895 | 0.129213 | 0.04908 | 0.080178 | 0.111111 | 0.169884 | 0.096154 |
| SMAD1 | 0.098361 | 0.106092 | 0.027692 | 0.045752 | 0.08427 | 0.02863 | 0.109131 | 0.050505 | 0.111969 | 0.067308 |
| HNRNPA1 | 0.251366 | 0.20063 | 0.184615 | 0.098039 | 0.16573 | 0.237219 | 0.289532 | 0.308081 | 0.359073 | 0.201923 |
| FXR1 | 0.52459 | 0.301471 | 0.2 | 0.215686 | 0.744382 | 0.167689 | 0.280624 | 0.90404 | 0.803089 | 0.394231 |
| ADAR | 0.47541 | 0.72479 | 0.236923 | 0.176471 | 0.266854 | 0.120654 | 0.732739 | 0.532828 | 0.629344 | 0.605769 |
| ELAVL1 | 0.15847 | 0.162815 | 0.166154 | 0.346405 | 0.098315 | 0.096115 | 0.03118 | 0.184343 | 0.262548 | 0.144231 |
| MOV10 | 0.224044 | 0.155462 | 0.043077 | 0.156863 | 0.073034 | 0.03681 | 0.231626 | 0.118687 | 0.312741 | 0.192308 |
| PABPC1 | 0.63388 | 0.612395 | 0.590769 | 0.124183 | 0.688202 | 0.132924 | 0.565702 | 0.621212 | 0.65251 | 0.461538 |
| SRSF1 | 0.42623 | 0.357143 | 0.212308 | 0.137255 | 0.19382 | 0.06953 | 0.465479 | 0.414141 | 0.23166 | 0.182692 |
| SNIP1 | 0.245902 | 0.125 | 0.046154 | 0.176471 | 0.151685 | 0.026585 | 0.267261 | 0.14899 | 0.444015 | 0.163462 |
| DCP2 | 0.081967 | 0.184874 | 0.08 | 0.071895 | 0.067416 | 0.529652 | 0.162584 | 0.022727 | 0.096525 | 0.019231 |
| ILF3 | 0.185792 | 0.186975 | 0.172308 | 0.359477 | 0.123596 | 0.096115 | 0.037862 | 0.19697 | 0.42471 | 0.25 |
| CNOT1 | 0.218579 | 0.096639 | 0.215385 | 0.039216 | 0.241573 | 0.192229 | 0.198218 | 0.181818 | 0.065637 | 0.048077 |
| SMAD5 | 0.071038 | 0.196429 | 0.083077 | 0.071895 | 0.070225 | 0.605317 | 0.14922 | 0.037879 | 0.146718 | 0.009615 |
| SMAD3 | 0.114754 | 0.108193 | 0.033846 | 0.058824 | 0.171348 | 0.05317 | 0.089087 | 0.247475 | 0.146718 | 0.019231 |
| RBM4 | 0.300546 | 0.256303 | 0.123077 | 0.03268 | 0.348315 | 0.067485 | 0.327394 | 0.287879 | 0.343629 | 0.182692 |
| KHSRP | 0.136612 | 0.152311 | 0.175385 | 0.326797 | 0.081461 | 0.09407 | 0.03118 | 0.176768 | 0.146718 | 0.076923 |
| MAPKAPK2 | 0.31694 | 0.747899 | 0.24 | 0.169935 | 0.266854 | 0.112474 | 0.697105 | 0.517677 | 0.486486 | 0.480769 |
| PIWIL1 | 0.251366 | 0.176471 | 0.169231 | 0.091503 | 0.13764 | 0.237219 | 0.25167 | 0.257576 | 0.293436 | 0.125 |
| FMR1 | 0.136612 | 0.165966 | 0.2 | 0.084967 | 0.264045 | 0.05726 | 0.278396 | 0.217172 | 0.351351 | 0.192308 |
| DROSHA | 0.480874 | 0.328782 | 0.190769 | 0.065359 | 0.44382 | 0.327198 | 0.616927 | 0.770202 | 0.498069 | 0.192308 |
| TNRC6A | 0.251366 | 0.523109 | 0.243077 | 0.058824 | 0.210674 | 0.214724 | 0.291759 | 0.209596 | 0.166023 | 0.125 |

**Table S6. Differentially expressed miRNA biogenesis genes.**

Cross-cancer differential gene expression analysis with $log_2$ fold change and adjusted P-values (see Methods). NA entries correspond to the genes which were either not present on the platform (microarray) or dropped due to lack of abundance in RNA-seq data.

| | BLCA.FC | BLCA.Padj | BRCA.FC | BRCA.Padj | COADREAD.FC | COADREAD.Padj | GBM.FC | GBM.Padj | HNSC.FC | HNSC.Padj | KIRC.FC | KIRC.Padj | LUAD.FC | LUAD.Padj | LUSC.FC | LUSC.Padj | OV.FC | OV.Padj | UCEC.FC | UCEC.Padj |
|---|---|---|---|---|---|---|---|---|---|---|---|---|---|---|---|---|---|---|---|---|
| EIF2C3 | -0.05 | 0.8385 | -0.20 | 0.0005 | 0.15 | 0.0511 | 0.72 | 0.0002 | 0.52 | 0.0000 | -0.15 | 0.0007 | 0.37 | 0.0000 | 0.48 | 0.0000 | 1.03 | 0.0015 | 0.72 | 0.0000 |
| PRKRA | -0.27 | 0.0204 | -0.02 | 0.4982 | 0.14 | 0.0637 | 0.30 | 0.0464 | -0.18 | 0.0066 | -0.17 | 0.0000 | 0.21 | 0.0000 | 0.38 | 0.0000 | 0.40 | 0.0457 | 0.08 | 0.2906 |
| ESR1 | -2.80 | 0.0000 | 0.40 | 0.0098 | -1.91 | 0.0000 | -0.25 | 0.0003 | -0.36 | 0.1190 | -0.74 | 0.0000 | -0.09 | 0.4588 | -1.57 | 0.0000 | -0.33 | 0.0393 | -1.68 | 0.0000 |
| EIF2C4 | -0.57 | 0.0101 | -0.84 | 0.0000 | -0.36 | 0.0031 | 0.13 | 0.0018 | -0.60 | 0.0000 | 0.09 | 0.0678 | -0.56 | 0.0000 | -0.77 | 0.0000 | 0.18 | 0.0418 | -0.84 | 0.0000 |
| TARBP1 | 0.97 | 0.0000 | 0.57 | 0.0000 | 1.11 | 0.0000 | -0.26 | 0.1557 | 0.11 | 0.1775 | 0.12 | 0.0095 | 0.66 | 0.0000 | 1.25 | 0.0000 | 1.11 | 0.0013 | 0.97 | 0.0026 |
| RNASEN | 0.45 | 0.0001 | 0.11 | 0.0000 | 0.44 | 0.0000 | 0.07 | 0.5693 | 0.37 | 0.0000 | -0.08 | 0.0208 | 0.78 | 0.0000 | 1.06 | 0.0000 | 0.10 | 0.6080 | 0.22 | 0.0029 |
| ARS2 | NA | NA | NA | NA | NA | NA | 0.45 | 0.0000 | NA | NA | NA | NA | NA | NA | NA | NA | 0.28 | 0.0008 | NA | NA |
| ESR2 | -0.20 | 0.6761 | -1.87 | 0.0000 | -0.83 | 0.0000 | 0.09 | 0.0037 | 0.58 | 0.0001 | -0.84 | 0.0000 | 0.18 | 0.1145 | 0.37 | 0.0110 | 0.05 | 0.2306 | -0.08 | 0.8287 |
| ADARB1 | -1.89 | 0.0020 | -0.68 | 0.0000 | -0.69 | 0.0025 | -0.77 | 0.0000 | 0.45 | 0.0054 | 0.40 | 0.0000 | -1.87 | 0.0000 | -2.46 | 0.0000 | -0.18 | 0.0945 | -1.90 | 0.0000 |
| EIF2C2 | 0.38 | 0.1015 | 0.38 | 0.0000 | 1.07 | 0.0000 | NA | NA | 0.83 | 0.0000 | 0.12 | 0.0672 | 0.63 | 0.0000 | 1.17 | 0.0000 | NA | NA | 1.15 | 0.0000 |
| DGCR8 | 0.25 | 0.0011 | 0.02 | 0.6818 | 0.10 | 0.1354 | 0.25 | 0.0214 | 0.08 | 0.1486 | 0.01 | 0.8181 | 0.24 | 0.0000 | 0.46 | 0.0000 | 0.12 | 0.3751 | -0.01 | 0.9107 |
| XPO5 | 0.79 | 0.0000 | 0.63 | 0.0000 | 1.31 | 0.0000 | NA | NA | 0.60 | 0.0000 | -0.14 | 0.0003 | 1.02 | 0.0000 | 1.40 | 0.0000 | NA | NA | 1.12 | 0.0000 |

| | | | | | | | | | | | | | | | | | | | | |
|---|---|---|---|---|---|---|---|---|---|---|---|---|---|---|---|---|---|---|---|---|
| DICER1 | -0.19 | 0.1820 | -0.42 | 0.0000 | -0.29 | 0.0018 | 0.22 | 0.0104 | -0.13 | 0.2056 | -0.66 | 0.0000 | -0.33 | 0.0000 | 0.00 | 0.9319 | -0.01 | 0.9413 | -0.34 | 0.0035 |
| TARBP2 | 1.14 | 0.0000 | 0.62 | 0.0000 | 0.55 | 0.0000 | 1.21 | 0.0000 | 0.33 | 0.0000 | -0.02 | 0.7472 | 0.66 | 0.0000 | 0.88 | 0.0000 | 1.02 | 0.0064 | 1.18 | 0.0000 |
| TRIM32 | 0.06 | 0.7602 | -0.08 | 0.0028 | 0.21 | 0.0031 | -0.06 | 0.6708 | 0.18 | 0.0537 | -0.11 | 0.0089 | 0.20 | 0.0001 | 0.44 | 0.0000 | -0.47 | 0.0455 | -0.78 | 0.0000 |
| EIF2C1 | -0.05 | 0.7882 | -0.28 | 0.0000 | -0.19 | 0.0331 | 0.78 | 0.0001 | 0.20 | 0.0054 | 0.18 | 0.0001 | 0.02 | 0.6917 | 0.10 | 0.1161 | 0.05 | 0.7916 | -0.12 | 0.1288 |
| RAN | 0.62 | 0.0001 | 0.72 | 0.0000 | 0.83 | 0.0000 | 0.29 | 0.0060 | 0.51 | 0.0000 | 0.22 | 0.0000 | 0.75 | 0.0000 | 1.23 | 0.0000 | 0.33 | 0.0064 | 1.00 | 0.0000 |
| DDX17 | -0.42 | 0.0015 | -0.50 | 0.0000 | -0.03 | 0.5947 | -0.08 | 0.6720 | -0.06 | 0.3387 | 0.28 | 0.0000 | -0.23 | 0.0006 | -0.20 | 0.0050 | -0.22 | 0.4706 | -0.46 | 0.0008 |
| TSN | 0.13 | 0.1231 | 0.11 | 0.0000 | 0.41 | 0.0000 | 0.60 | 0.0004 | 0.30 | 0.0000 | -0.27 | 0.0000 | 0.37 | 0.0000 | 0.65 | 0.0000 | 0.44 | 0.0064 | 0.33 | 0.0004 |
| DDX20 | 0.15 | 0.0592 | -0.02 | 0.5897 | 0.33 | 0.0000 | NA | NA | 0.29 | 0.0000 | 0.09 | 0.0128 | -0.11 | 0.0007 | 0.04 | 0.3309 | NA | NA | -0.12 | 0.0919 |
| IPO8 | -0.41 | 0.0027 | -0.27 | 0.0000 | -0.26 | 0.0000 | 0.21 | 0.0104 | -0.05 | 0.3129 | -0.46 | 0.0000 | -0.18 | 0.0000 | -0.02 | 0.5927 | 0.17 | 0.1578 | -0.28 | 0.0318 |
| DHX9 | 0.07 | 0.3168 | 0.27 | 0.0000 | 0.48 | 0.0000 | 0.25 | 0.0013 | 0.24 | 0.0004 | -0.49 | 0.0000 | 0.31 | 0.0000 | 0.56 | 0.0000 | 0.25 | 0.2694 | 0.10 | 0.2906 |
| DDX5 | -0.49 | 0.0061 | -0.23 | 0.0000 | 0.12 | 0.1148 | 0.96 | 0.0000 | -0.16 | 0.0537 | 0.12 | 0.0052 | -0.32 | 0.0000 | -0.49 | 0.0000 | 0.28 | 0.0745 | -0.36 | 0.0035 |
| GEMIN4 | 0.06 | 0.6063 | -0.39 | 0.0000 | 0.51 | 0.0000 | 0.62 | 0.0000 | 0.06 | 0.4090 | 0.06 | 0.1379 | 0.01 | 0.7659 | 0.42 | 0.0000 | 0.27 | 0.0457 | 0.24 | 0.0244 |
| SMAD1 | -0.10 | 0.4144 | -0.14 | 0.0003 | -0.50 | 0.0000 | 1.85 | 0.0000 | -0.02 | 0.7413 | -0.39 | 0.0000 | 0.43 | 0.0000 | 0.65 | 0.0000 | -0.18 | 0.4383 | -0.90 | 0.0001 |
| HNRNPA1 | 0.01 | 0.9466 | 0.03 | 0.5594 | 0.66 | 0.0000 | 0.38 | 0.0000 | -0.10 | 0.0961 | 0.44 | 0.0000 | 0.23 | 0.0000 | 0.33 | 0.0000 | -0.23 | 0.0008 | -0.26 | 0.0367 |
| FXR1 | 0.10 | 0.2753 | -0.01 | 0.8535 | 0.34 | 0.0000 | 0.62 | 0.0001 | 0.34 | 0.0122 | 0.18 | 0.0000 | 0.29 | 0.0000 | 1.44 | 0.0000 | 0.48 | 0.0079 | 0.02 | 0.8359 |
| ADAR | 0.49 | 0.0000 | 0.77 | 0.0000 | 0.22 | 0.0026 | -0.49 | 0.0003 | 0.78 | 0.0000 | -0.17 | 0.0000 | 0.52 | 0.0000 | 0.17 | 0.0041 | 0.00 | 0.9796 | 0.53 | 0.0000 |
| ELAVL1 | 0.32 | 0.0001 | 0.55 | 0.0000 | 0.51 | 0.0000 | 2.09 | 0.0000 | 0.25 | 0.0000 | 0.04 | 0.0667 | 0.25 | 0.0000 | 0.52 | 0.0000 | 0.26 | 0.0457 | 0.32 | 0.0000 |
| MOV10 | 0.93 | 0.0001 | 0.62 | 0.0000 | 0.16 | 0.0885 | NA | NA | 0.42 | 0.0001 | 0.21 | 0.0000 | 0.64 | 0.0000 | 0.43 | 0.0000 | NA | NA | 0.95 | 0.0000 |
| PABPC1 | 0.62 | 0.0011 | 0.32 | 0.0000 | 1.01 | 0.0000 | 0.60 | 0.0000 | -0.15 | 0.1944 | 0.71 | 0.0000 | 0.96 | 0.0000 | 1.10 | 0.0000 | -0.07 | 0.3751 | 0.29 | 0.0042 |

| | | | | | | | | | | | | | | | | | | | | |
|---|---|---|---|---|---|---|---|---|---|---|---|---|---|---|---|---|---|---|---|---|
| SNIP1 | -0.23 | 0.0323 | -0.21 | 0.0000 | -0.28 | 0.0000 | 0.45 | 0.0001 | 0.13 | 0.0335 | -0.01 | 0.7590 | -0.10 | 0.0119 | -0.07 | 0.1272 | 0.48 | 0.0008 | -0.32 | 0.0068 |
| DCP2 | 0.50 | 0.0000 | -0.03 | 0.5263 | 0.07 | 0.2590 | 0.41 | 0.0018 | 0.27 | 0.0003 | 0.16 | 0.0002 | -0.18 | 0.0010 | -0.18 | 0.0027 | 0.13 | 0.5836 | 0.11 | 0.1331 |
| ILF3 | 0.53 | 0.0000 | 0.36 | 0.0000 | 0.71 | 0.0000 | 0.71 | 0.0000 | 0.43 | 0.0000 | 0.18 | 0.0000 | 0.38 | 0.0000 | 0.74 | 0.0000 | 0.63 | 0.0078 | 0.48 | 0.0001 |
| CNOT1 | 0.01 | 0.9466 | -0.27 | 0.0000 | 0.21 | 0.0104 | 0.42 | 0.0002 | -0.22 | 0.0382 | -0.24 | 0.0000 | 0.16 | 0.0014 | 0.29 | 0.0000 | -0.27 | 0.0688 | 0.14 | 0.1331 |
| SMAD5 | -0.40 | 0.0323 | -0.47 | 0.0000 | 0.21 | 0.0118 | 1.16 | 0.0000 | -0.13 | 0.1052 | -0.08 | 0.0378 | 0.08 | 0.0524 | 0.08 | 0.1053 | 0.42 | 0.0737 | -1.26 | 0.0000 |
| SMAD3 | -0.17 | 0.4022 | -0.64 | 0.0000 | -0.20 | 0.0492 | -0.24 | 0.0049 | -0.14 | 0.2776 | -0.21 | 0.0000 | -0.05 | 0.4681 | 0.53 | 0.0000 | -0.31 | 0.0629 | -1.47 | 0.0000 |
| RBM4 | 0.21 | 0.0607 | 0.48 | 0.0000 | 0.09 | 0.0382 | 0.60 | 0.0000 | 0.19 | 0.0005 | -0.23 | 0.0000 | 0.37 | 0.0000 | 0.53 | 0.0000 | 0.48 | 0.0008 | 0.63 | 0.0000 |
| KHSRP | 0.41 | 0.0000 | 0.30 | 0.0000 | 0.53 | 0.0000 | 0.36 | 0.0004 | 0.28 | 0.0001 | 0.23 | 0.0000 | 0.25 | 0.0000 | 0.84 | 0.0000 | 0.11 | 0.4195 | 0.05 | 0.3880 |
| MAPKAPK2 | -0.20 | 0.2225 | 0.94 | 0.0000 | 0.22 | 0.0001 | 0.65 | 0.0000 | 0.12 | 0.1100 | 0.14 | 0.0000 | 0.11 | 0.0285 | 0.03 | 0.6934 | 0.14 | 0.0737 | 0.27 | 0.0222 |
| PIWIL1 | 0.50 | 0.0161 | 0.29 | 0.0000 | 3.18 | 0.0000 | -0.14 | 0.0036 | 0.51 | 0.0005 | -0.90 | 0.0000 | NA | NA | NA | NA | 0.06 | 0.2267 | 0.37 | 0.3880 |
| FMR1 | 0.17 | 0.1360 | 0.13 | 0.0010 | 0.40 | 0.0000 | 0.16 | 0.1347 | 0.31 | 0.0000 | -0.34 | 0.0000 | 0.01 | 0.8118 | 0.10 | 0.0231 | 1.02 | 0.0013 | -0.11 | 0.1993 |
| TNRC6A | -0.02 | 0.8671 | -0.35 | 0.0000 | -0.06 | 0.5947 | NA | NA | 0.23 | 0.0004 | 0.35 | 0.0000 | 0.20 | 0.0105 | 0.07 | 0.3097 | NA | NA | -0.85 | 0.0000 |

**Table S7. Spearman's correlation statistics of candidate metabolic genes.**

Cross-cancer Spearman's correlation statistics of genes (Table S1) that passed the pre-selection criteria of over-expression (Table S4) and gained fraction (Table S5).

| | rho | P | Q |
|---|---|---|---|
| **BLCA** | | | |
| PABPC1 | 0.611 | 7.29E-24 | 7.29E-23 |
| XPO5 | 0.54 | 4.42E-18 | 2.21E-17 |
| TARBP1 | 0.467 | 2.42E-13 | 8.07E-13 |
| ILF3 | 0.43 | 2.69E-11 | 6.72E-11 |
| ADAR | 0.423 | 5.94E-11 | 1.19E-10 |
| DGCR8 | 0.354 | 7.01E-08 | 1.17E-07 |
| TARBP2 | 0.332 | 4.68E-07 | 6.69E-07 |
| MOV10 | 0.297 | 7.63E-06 | 9.53E-06 |
| RAN | 0.271 | 4.77E-05 | 5.30E-05 |
| PIWIL1 | 0.003 | 0.962563 | 0.962563 |
| **BRCA** | | | |
| PABPC1 | 0.647 | 1.15E-119 | 8.02E-119 |
| EIF2C2 | 0.644 | 2.32E-118 | 8.13E-118 |
| XPO5 | 0.592 | 7.30E-96 | 1.70E-95 |
| TARBP1 | 0.511 | 1.09E-67 | 1.91E-67 |
| ADAR | 0.487 | 1.21E-60 | 1.69E-60 |

| | | | |
|---|---|---|---|
| MAPKAPK2 | 0.477 | 6.23E-58 | 7.27E-58 |
| DHX9 | 0.321 | 1.84E-25 | 1.84E-25 |
| **COADREAD** | | | |
| PABPC1 | 0.728 | 2.12E-55 | 1.70E-54 |
| XPO5 | 0.517 | 8.16E-24 | 3.26E-23 |
| MAPKAPK2 | 0.479 | 3.32E-20 | 8.86E-20 |
| EIF2C2 | 0.451 | 7.58E-18 | 1.52E-17 |
| ADAR | 0.364 | 1.00E-11 | 1.60E-11 |
| TARBP1 | 0.336 | 4.00E-10 | 5.33E-10 |
| CNOT1 | 0.334 | 5.17E-10 | 5.91E-10 |
| DHX9 | 0.293 | 6.46E-08 | 6.46E-08 |
| **GBM** | | | |
| KHSRP | 0.375 | 1.65E-06 | 4.96E-06 |
| ILF3 | 0.293 | 0.000229 | 0.000343 |
| ELAVL1 | 0.281 | 0.000423 | 0.000423 |
| **HNSC** | | | |
| FXR1 | 0.749 | 9.83E-77 | 6.88E-76 |
| EIF2C2 | 0.379 | 8.40E-16 | 2.94E-15 |
| DHX9 | 0.332 | 2.87E-12 | 6.70E-12 |
| TNRC6A | 0.298 | 4.68E-10 | 8.19E-10 |
| ADAR | 0.239 | 7.13E-07 | 9.99E-07 |
| FMR1 | 0.187 | 0.000112 | 0.000129 |
| ESR2 | 0.186 | 0.000129 | 0.000129 |

| | | | |
|---|---|---|---|
| **KIRC** | | | |
| RAN | 0.408 | 1.89E-21 | 7.56E-21 |
| DCP2 | 0.373 | 7.73E-18 | 1.55E-17 |
| TNRC6A | 0.12 | 0.007193 | 0.009591 |
| HNRNPA1 | 0.038 | 0.396906 | 0.396906 |
| **LUAD** | | | |
| PABPC1 | 0.704 | 8.25E-74 | 1.32E-72 |
| FXR1 | 0.674 | 1.03E-65 | 8.27E-65 |
| RAN | 0.56 | 1.56E-41 | 8.30E-41 |
| XPO5 | 0.529 | 2.21E-36 | 8.84E-36 |
| EIF2C2 | 0.482 | 1.39E-29 | 4.45E-29 |
| TARBP2 | 0.477 | 4.99E-29 | 1.33E-28 |
| TARBP1 | 0.469 | 6.90E-28 | 1.58E-27 |
| ADAR | 0.463 | 3.91E-27 | 7.82E-27 |
| MAPKAPK2 | 0.458 | 1.48E-26 | 2.63E-26 |
| MOV10 | 0.442 | 1.33E-24 | 2.12E-24 |
| PRKRA | 0.422 | 2.17E-22 | 3.15E-22 |
| TSN | 0.404 | 1.65E-20 | 2.20E-20 |
| DHX9 | 0.391 | 3.14E-19 | 3.87E-19 |
| TNRC6A | 0.331 | 7.05E-14 | 8.06E-14 |
| EIF2C3 | 0.325 | 1.87E-13 | 1.99E-13 |
| HNRNPA1 | 0.295 | 3.27E-11 | 3.27E-11 |
| **LUSC** | | | |
| FXR1 | 0.833 | 7.70E-125 | 1.39E-123 |

| | | | |
|---|---|---|---|
| TRIM32 | 0.625 | 2.58E-53 | 2.32E-52 |
| PRKRA | 0.571 | 9.07E-43 | 5.44E-42 |
| XPO5 | 0.55 | 3.67E-39 | 1.65E-38 |
| TARBP1 | 0.528 | 8.81E-36 | 3.17E-35 |
| PABPC1 | 0.517 | 3.83E-34 | 1.15E-33 |
| EIF2C2 | 0.51 | 4.14E-33 | 1.07E-32 |
| TARBP2 | 0.502 | 6.45E-32 | 1.45E-31 |
| ADAR | 0.466 | 3.35E-27 | 6.70E-27 |
| TSN | 0.462 | 9.82E-27 | 1.77E-26 |
| RAN | 0.45 | 2.93E-25 | 4.80E-25 |
| DHX9 | 0.429 | 7.98E-23 | 1.20E-22 |
| HNRNPA1 | 0.364 | 1.76E-16 | 2.43E-16 |
| SMAD3 | 0.363 | 2.24E-16 | 2.88E-16 |
| ILF3 | 0.36 | 4.33E-16 | 5.20E-16 |
| ESR2 | 0.239 | 1.15E-07 | 1.29E-07 |
| DGCR8 | 0.237 | 1.47E-07 | 1.56E-07 |
| FMR1 | 0.236 | 1.82E-07 | 1.82E-07 |
| **OV** | | | |
| ELAVL1 | 0.762 | 2.48E-50 | 2.97E-49 |
| SNIP1 | 0.727 | 7.21E-44 | 4.33E-43 |
| EIF2C3 | 0.643 | 1.32E-31 | 3.95E-31 |
| ILF3 | 0.643 | 1.28E-31 | 3.95E-31 |
| FXR1 | 0.623 | 3.24E-29 | 7.78E-29 |
| PRKRA | 0.619 | 8.15E-29 | 1.63E-28 |
| TSN | 0.612 | 5.15E-28 | 8.83E-28 |
| TARBP2 | 0.553 | 4.17E-22 | 6.26E-22 |
| TARBP1 | 0.548 | 1.05E-21 | 1.40E-21 |

| | | | |
|---|---|---|---|
| RAN | 0.544 | 2.34E-21 | 2.81E-21 |
| EIF2C4 | 0.413 | 4.57E-12 | 4.99E-12 |
| FMR1 | 0.229 | 0.000198 | 0.000198 |
| **UCEC** | | | |
| MAPKAPK2 | 0.629 | 2.47E-17 | 1.23E-16 |
| ADAR | 0.563 | 1.75E-13 | 4.37E-13 |
| PABPC1 | 0.511 | 5.32E-11 | 8.87E-11 |
| EIF2C2 | 0.464 | 4.33E-09 | 5.41E-09 |
| TARBP1 | 0.426 | 9.36E-08 | 9.36E-08 |

**Table S8. Pan-cancer inclusion frequency of candidate miRNA biogenesis genes.**

Cross-cancer inclusion frequency of genes whereby mRNA and copy-number profiles are significantly correlated ($\rho > 0.3$, $q < 10^{-3}$) (as depicted in Table S6, and described in Methods). '1' represents presence of correlated genes, while '0' represents genes failing the above-mentioned criteria. Genes not included in any of the cancer types are not listed.

| | BLCA.All | BRCA.All | COADREAD.All | GBM.All | HNSC.All | KIRC.All | LUAD.All | LUSC.All | OV.All | UCEC.All | summary |
|---|---|---|---|---|---|---|---|---|---|---|---|
| TARBP1 | 1 | 1 | 1 | 0 | 0 | 0 | 1 | 1 | 1 | 1 | 7 |
| ADAR | 1 | 1 | 1 | 0 | 0 | 0 | 1 | 1 | 0 | 1 | 6 |
| PABPC1 | 1 | 1 | 1 | 0 | 0 | 0 | 1 | 1 | 0 | 1 | 6 |
| EIF2C2 | 0 | 1 | 1 | 0 | 1 | 0 | 1 | 1 | 0 | 1 | 6 |
| XPO5 | 1 | 1 | 1 | 0 | 0 | 0 | 1 | 1 | 0 | 0 | 5 |
| TARBP2 | 1 | 0 | 0 | 0 | 0 | 0 | 1 | 1 | 1 | 0 | 4 |
| DHX9 | 0 | 1 | 0 | 0 | 1 | 0 | 1 | 1 | 0 | 0 | 4 |
| MAPKAPK2 | 0 | 1 | 1 | 0 | 0 | 0 | 1 | 0 | 0 | 1 | 4 |
| FXR1 | 0 | 0 | 0 | 0 | 1 | 0 | 1 | 1 | 1 | 0 | 4 |
| RAN | 0 | 0 | 0 | 0 | 0 | 1 | 1 | 1 | 1 | 0 | 4 |
| ILF3 | 1 | 0 | 0 | 0 | 0 | 0 | 0 | 1 | 1 | 0 | 3 |
| PRKRA | 0 | 0 | 0 | 0 | 0 | 0 | 1 | 1 | 1 | 0 | 3 |
| TSN | 0 | 0 | 0 | 0 | 0 | 0 | 1 | 1 | 1 | 0 | 3 |
| EIF2C3 | 0 | 0 | 0 | 0 | 0 | 0 | 1 | 0 | 1 | 0 | 2 |
| DGCR8 | 1 | 0 | 0 | 0 | 0 | 0 | 0 | 0 | 0 | 0 | 1 |
| CNOT1 | 0 | 0 | 1 | 0 | 0 | 0 | 0 | 0 | 0 | 0 | 1 |
| KHSRP | 0 | 0 | 0 | 1 | 0 | 0 | 0 | 0 | 0 | 0 | 1 |
| DCP2 | 0 | 0 | 0 | 0 | 0 | 1 | 0 | 0 | 0 | 0 | 1 |
| MOV10 | 0 | 0 | 0 | 0 | 0 | 0 | 1 | 0 | 0 | 0 | 1 |
| TNRC6A | 0 | 0 | 0 | 0 | 0 | 0 | 1 | 0 | 0 | 0 | 1 |

| | | | | | | | | | | | |
|---|---|---|---|---|---|---|---|---|---|---|---|
| TRIM32 | 0 | 0 | 0 | 0 | 0 | 0 | 0 | 1 | 0 | 0 | 1 |
| HNRNPA1 | 0 | 0 | 0 | 0 | 0 | 0 | 0 | 1 | 0 | 0 | 1 |
| SMAD3 | 0 | 0 | 0 | 0 | 0 | 0 | 0 | 1 | 0 | 0 | 1 |
| EIF2C4 | 0 | 0 | 0 | 0 | 0 | 0 | 0 | 0 | 1 | 0 | 1 |
| ELAVL1 | 0 | 0 | 0 | 0 | 0 | 0 | 0 | 0 | 1 | 0 | 1 |
| SNIP1 | 0 | 0 | 0 | 0 | 0 | 0 | 0 | 0 | 1 | 0 | 1 |

**Table S9. microRNA biogenesis genes in independent breast cancer cohorts: amplification frequency, and association with expression and survival.** Amplification (amp) events, test for correlation between copy number (CN) status and expression (Exp), and ability to predict prognosis (HR=Hazard Ratio in Cox survival analysis) are shown for miRNA biogenesis genes (Table S1). Only patients with matched genomic (CN), transcriptomic (both coding and non-coding mRNA) and clinical follow-up data were considered for each dataset.

| Gene Identifiers | | Oxford Samples (N=180) | | | | | | Metabric Samples (N=1293) | | | | | |
|---|---|---|---|---|---|---|---|---|---|---|---|---|---|
| Symbol | Ensembl | Total Patients with amp | Total Patients with high amp | Correlation CN - Exp p | Correlation CN - Exp rho | Cox survival Amp p-value | Cox survival Amp HR | Total Patients with amp | Total Patients with high amp | Correlation CN - Exp p | Correlation CN - Exp rho | Cox survival Amp p-value | Cox survival Amp HR |

| | | | | | | | | | | | | | |
|---|---|---|---|---|---|---|---|---|---|---|---|---|---|
| ADAR | ENSG00000160710 | 60 | 4 | 1.49E-08 | 0.41 | 0.9732 | 1.0 | 498 | 18 | 1.32E-44 | 0.38 | 0.4343 | 0.9 |
| ARS2 | ENSG00000087087 | 11 | 1 | 0.0242 | 0.17 | 0.1255 | 1.9 | 68 | 1 | 1.07E-08 | 0.16 | 0.5934 | 0.9 |
| CNOT1 | ENSG00000125107 | 11 | 0 | 0.0093 | 0.19 | 0.1298 | 1.9 | 12 | 0 | 5.89E-49 | 0.39 | 0.2395 | 0.3 |
| DDX5 | ENSG00000108654 | 32 | 5 | 0.0086 | 0.20 | 0.2695 | 1.4 | 176 | 22 | 6.72E-35 | 0.34 | 0.0012 | 1.6 |
| DGCR8 | ENSG00000128191 | 2 | 1 | 0.0029 | 0.24 | 0.5577 | 1.8 | 70 | 5 | 4.24E-06 | 0.13 | 0.9201 | 1.0 |
| DHX9 | ENSG00000135829 | 71 | 8 | 1.81E-05 | 0.31 | 0.1669 | 1.4 | 516 | 23 | 6.20E-10 | 0.17 | 0.1901 | 0.9 |
| EIF2C1 (AGO1) | ENSG00000092847 | 3 | 0 | 0.0044 | 0.21 | 0.9951 | 0.0 | 28 | 1 | 1.21E-16 | 0.23 | 0.1719 | 1.6 |
| EIF2C2 (AGO2) | ENSG00000123908 | 47 | 13 | 9.99E-06 | 0.33 | 0.0008 | 2.4 | 350 | 42 | 5.65E-54 | 0.42 | 0.0140 | 1.4 |
| ELAVL1 | ENSG00000066044 | 5 | 2 | 0.0004 | 0.26 | 0.0048 | 4.3 | 35 | 0 | 3.56E-09 | 0.16 | 0.1097 | 1.6 |
| GEMIN4 | ENSG00000179409 | 11 | 0 | 0.0204 | 0.17 | 0.0196 | 2.6 | 16 | 0 | 1.11E-41 | 0.38 | 0.7566 | 1.2 |
| KHSRP | ENSG00000088247 | 6 | 2 | 0.0018 | 0.23 | 0.0009 | 4.7 | 26 | 0 | 6.39E-10 | 0.17 | 0.4622 | 1.3 |
| MAPKAPK2 | ENSG00000162889 | 74 | 13 | 1.74E-05 | 0.32 | 0.3736 | 1.3 | 536 | 20 | 3.64E-47 | 0.39 | 0.8220 | 1.0 |
| PABPC1 | ENSG00000070756 | 52 | 18 | 1.65E-06 | 0.35 | 0.0048 | 2.0 | 435 | 53 | 3.34E-25 | 0.28 | 0.0058 | 1.4 |
| PRKRA | ENSG00000180228 | 8 | 1 | 0.0010 | 0.24 | 0.8507 | 0.9 | 14 | 0 | 3.48E-05 | 0.12 | 0.1743 | 1.8 |
| RAN | ENSG00000132341 | 7 | 0 | 0.0005 | 0.26 | 0.7792 | 0.8 | 48 | 1 | 2.14E-06 | 0.13 | 0.0754 | 1.6 |
| RBM4 | ENSG00000173933 | 21 | 6 | 1.70E-05 | 0.32 | 0.9049 | 1.0 | 138 | 23 | 1.75E-31 | 0.32 | 0.2429 | 1.2 |
| RENT1 | ENSG00000005007 | 8 | 2 | 8.29E-05 | 0.29 | 0.2848 | 1.7 | 72 | 3 | 7.09E-09 | 0.16 | 0.0620 | 1.5 |

| | | | | | | | | | | | | | |
|---|---|---|---|---|---|---|---|---|---|---|---|---|---|
| RNASEN (DROSHA) | ENSG00000113360 | 13 | 2 | 0.0025 | 0.22 | 0.0076 | 2.6 | 99 | 3 | 1.66E-34 | 0.34 | 0.2128 | 1.3 |
| SRSF1 | ENSG00000136450 | 26 | 5 | 0.0104 | 0.19 | 0.2076 | 1.5 | 206 | 47 | 1.02E-18 | 0.24 | 0.0404 | 1.3 |
| TARBP1 | ENSG00000059588 | 76 | 9 | 4.25E-05 | 0.30 | 0.3276 | 1.3 | 493 | 12 | 1.23E-45 | 0.38 | 0.9462 | 1.0 |

**Table S10. miRNA Processing Machinery Genes disrupted by Deletion in validation cohorts.**

Deletion Frequency of miRNA Biogenesis genes in Breast Cancer Patients, and their ability to predict prognosis (HR = Hazard Ratio in Cox survival analysis). Only patients with matched genomic (CN), transcriptomic (both coding and non-coding mRNA) and clinical follow-up data were considered for each dataset.

| | **Oxford (N=180)** | | | **Metabric (N=1293)** | | |
|---|---|---|---|---|---|---|
| | Frequency | Cox p-value | Cox HR | Frequency | Cox p-value | Cox HR |
| ADAR | 0.000 | NA | NA | 0.002 | NA | NA |
| ADARB1 | 0.006 | NA | NA | 0.010 | 0.2055 | 0.3 |
| ARS2 | 0.028 | 0.2401 | 2.0 | 0.026 | 0.7467 | 0.9 |
| CNOT1 | 0.067 | 0.7462 | 0.8 | 0.246 | 0.1700 | 0.8 |
| DCP2 | 0.033 | 0.5884 | 1.4 | 0.034 | 0.1301 | 1.5 |
| DDX17 | 0.094 | 0.7771 | 0.9 | 0.114 | 0.5566 | 1.1 |
| DDX20 | 0.039 | 0.6733 | 0.7 | 0.030 | 9.72E-05 | 2.5 |
| DDX5 | 0.006 | NA | NA | 0.010 | 0.4265 | 0.5 |
| DGCR8 | 0.072 | 0.2672 | 0.5 | 0.085 | 0.4847 | 1.1 |
| DHX9 | 0.000 | NA | NA | 0.002 | NA | NA |
| DICER1 | 0.022 | NA | NA | 0.038 | 0.6687 | 0.9 |
| DND1 | 0.022 | NA | NA | NA | NA | NA |
| EIF2C1 | 0.022 | NA | NA | 0.033 | 0.0336 | 1.7 |
| EIF2C2 | 0.000 | NA | NA | 0.005 | 0.9908 | 0.0 |

| | | | | | | |
|---|---|---|---|---|---|---|
| EIF2C3 | 0.022 | NA | NA | 0.033 | 0.0347 | 1.7 |
| EIF2C4 | 0.022 | NA | NA | 0.032 | 0.2163 | 1.4 |
| ELAVL1 | 0.039 | 0.5624 | 1.4 | 0.025 | 0.2004 | 1.5 |
| ESR1 | 0.006 | NA | NA | 0.054 | 0.9101 | 1.0 |
| ESR2 | 0.022 | NA | NA | 0.039 | 0.5264 | 0.8 |
| FMR1 | 0.028 | 0.6020 | 1.4 | NA | NA | NA |
| FXR1 | 0.006 | NA | NA | 0.003 | NA | NA |
| GEMIN4 | 0.078 | 0.5021 | 1.3 | 0.131 | 0.6946 | 1.1 |
| HNRNPA1 | 0.022 | NA | NA | 0.006 | 0.1339 | 2.1 |
| ILF3 | 0.033 | 0.9347 | 0.9 | 0.017 | 0.8437 | 1.1 |
| IPO8 | 0.022 | NA | NA | 0.012 | 0.2066 | 1.8 |
| KHSRP | 0.050 | 0.4606 | 1.5 | 0.024 | 0.3893 | 1.3 |
| LIN28A | 0.067 | 0.1324 | 1.8 | NA | NA | NA |
| MAPKAPK2 | 0.006 | NA | NA | 0.001 | NA | NA |
| MOV10 | 0.039 | 0.6733 | 0.7 | 0.033 | 0.0012 | 2.2 |
| PABPC1 | 0.000 | NA | NA | 0.001 | NA | NA |
| PIWIL1 | 0.022 | NA | NA | NA | NA | NA |
| PRKRA | 0.011 | NA | NA | 0.005 | 0.5864 | 0.6 |
| RAN | 0.022 | NA | NA | 0.020 | 0.3682 | 1.4 |
| RBM4 | 0.000 | NA | NA | 0.016 | 0.7828 | 1.1 |
| RENT1 | 0.006 | NA | NA | 0.010 | 0.7251 | 1.2 |
| RNASEN | 0.017 | NA | NA | 0.005 | 0.4139 | 0.4 |

| | | | | | | |
|---|---|---|---|---|---|---|
| SMAD1 | 0.000 | NA | NA | 0.014 | 0.7532 | 0.8 |
| SMAD3 | 0.033 | 0.0301 | 2.8 | 0.011 | 0.9632 | 1.0 |
| SMAD5 | 0.011 | NA | NA | 0.016 | 0.9715 | 1.0 |
| SNIP1 | 0.022 | NA | NA | 0.034 | 0.0028 | 2.0 |
| SRSF1 | 0.006 | NA | NA | 0.015 | 0.7435 | 1.2 |
| TARBP1 | 0.006 | NA | NA | 0.002 | NA | NA |
| TARBP2 | 0.017 | NA | NA | 0.010 | 0.0004 | 4.3 |
| TNRC6 | 0.006 | NA | NA | 0.008 | 0.7213 | 1.2 |
| TP53 | 0.089 | 0.1555 | 1.7 | NA | NA | NA |
| TRIM32 | 0.044 | 0.9922 | 1.0 | 0.029 | 0.7068 | 0.9 |
| TSN | 0.006 | NA | NA | 0.008 | 0.7061 | 1.2 |
| XPO5 | 0.006 | NA | NA | 0.020 | 0.8942 | 1.1 |

**Table S11. Correlation of AGO2 expression with the abundance of miRNA included in a previously identified prognostic profile [2]**

| | | | Correlation (Spearman, miRNA /mRNA expression) | | | Cox Survival Analysis | | | |
|---|---|---|---|---|---|---|---|---|---|
| miRNA name | Prognosis group for cases with high miRNA expression | Tumour ER status | p-value | Rho | miRNA expression range | AGO2 CN p-value | AGO2 CN Hazard Ratio | miRNA expression p-value | miRNA expression Hazard Ratio |
| hsa-miR-30c | Good | ER - | $5.10x10^{-05}$ | -0.30 | 0.81 | 0.00175291 | 2.33 | 0.00223087 | 0.23 |
| hsa-miR-342 | Good | ER - | 0.002075407 | -0.23 | 2.03 | 0.003809174 | 2.19 | 0.00825625 | 0.29 |
| hsa-miR-135a | Good | ER + | 0.002254837 | -0.23 | 6.48 | 0.002624873 | 2.25 | 0.00027126 | 0.17 |
| hsa-miR-29c | Good | | 0.002435096 | -0.23 | 1.35 | 0.004943668 | 2.14 | 0.00201997 | 0.23 |
| hsa-miR-30e-3p | Good | ER + | 0.004893281 | -0.21 | 1.39 | 0.006173892 | 2.10 | 0.01518604 | 0.32 |
| hsa-miR-642 | Good | | 0.012892392 | -0.19 | 4.23 | 0.010207844 | 2.01 | 0.00298975 | 0.24 |
| hsa-miR-486 | Good | ER + | 0.019855065 | -0.18 | 5.24 | 0.006566612 | 2.11 | 0.10313816 | 0.47 |
| hsa-miR-27b | Poor | ER - | 0.194377158 | -0.10 | 1.04 | 0.000588683 | 2.56 | 0.01544890 | 3.21 |
| hsa-miR-128a | Poor | ER + | 0.424772339 | -0.06 | 5.61 | 0.006412847 | 2.12 | 0.19773071 | 1.85 |
| hsa-miR-768-3p | Good | ER - | 0.747827168 | -0.02 | 1.60 | 0.00711838 | 2.10 | 0.13646394 | 0.50 |
| hsa-miR-144 | Poor | ER - | 0.334332519 | 0.07 | 3.17 | 0.001608099 | 2.34 | 0.13032243 | 2.04 |

| | | | | | | | | | |
|---|---|---|---|---|---|---|---|---|---|
| hsa-miR-429 | Poor | | 0.222476553 | 0.09 | 0.65 | 0.002109675 | 2.29 | 0.03252385 | 2.69 |
| hsa-miR-92 | Good | ER - | 0.193784512 | 0.10 | 2.08 | 0.010128255 | 2.01 | 0.00305176 | 0.23 |
| hsa-miR-767-3p | Poor | ER + | 0.143213348 | 0.11 | 1.03 | 0.005182663 | 2.14 | 0.09151217 | 2.22 |
| hsa-miR-769-3p | Poor | ER + | 0.037323489 | 0.16 | 2.68 | 0.01794491 | 1.93 | 0.00892986 | 3.58 |
| hsa-miR-150 | Good | ER - | 0.008251167 | 0.20 | 2.25 | 0.006806755 | 2.10 | 0.07988931 | 0.44 |
| hsa-miR-210 | Poor | ER - | 0.001050079 | 0.25 | 5.57 | 0.023194558 | 1.86 | 0.00000671 | 11.21 |
| hsa-miR-548d | Poor | ER + | 0.000556342 | 0.26 | 2.32 | 0.032023028 | 1.82 | 0.00413195 | 4.30 |
| hsa-miR-130b | Poor | ER + | 0.000167489 | 0.28 | 3.55 | 0.008658258 | 2.04 | 0.00119179 | 5.03 |

**Table S12. miRNA profile associated with AGO2 expression across independent invasive breast cancer cohorts.**

miRNAs with abundance significantly correlated with mRNA levels of AGO2, in all three breast cancer datasets considered in this study (see methods), are shown. Correlations calculated between mRNA and miRNA expression and adjusted p-value quoted (Benjamini & Hochberg). Only patients with matched genomic (CN), transcriptomic (both coding and non-coding mRNA) and clinical follow-up data were considered.

| | Oxford (N=180) | | Metabric (N=1293) | | TCGA (N=1029) | |
|---|---|---|---|---|---|---|
| | Correlation | p-value (adjusted BH) | Correlation | p-value (adjusted BH) | Correlation | p-value (adjusted BH) |
| hsa-miR-18a | 0.32 | 0.0048 | 0.22 | 5.77E-15 | 0.48 | 8.32E-39 |
| hsa-miR-20a | 0.29 | 0.0098 | 0.10 | 0.0003 | 0.25 | 1.10E-10 |
| hsa-miR-93 | 0.26 | 0.0243 | 0.27 | 8.74E-22 | 0.31 | 8.85E-16 |
| hsa-miR-130b | 0.23 | 0.0392 | 0.11 | 8.54E-05 | 0.48 | 0 |
| hsa-miR-19a | 0.21 | 0.0591 | 0.24 | 1.74E-18 | 0.28 | 2.82E-13 |
| hsa-miR-224 | 0.20 | 0.0600 | 0.12 | 1.35E-05 | 0.32 | 2.97E-17 |
| hsa-miR-25 | 0.18 | 0.0969 | 0.17 | 1.66E-09 | 0.28 | 5.49E-13 |
| hsa-miR-106b | 0.14 | 0.1926 | 0.24 | 3.94E-18 | 0.49 | 0 |
| hsa-miR-204 | -0.14 | 0.1980 | -0.12 | 5.54E-05 | -0.13 | 0.0012 |
| hsa-miR-432 | -0.17 | 0.1098 | -0.27 | 1.18E-21 | -0.13 | 0.001212216 |
| hsa-miR-375 | -0.19 | 0.0703 | -0.12 | 1.78E-05 | -0.27 | 2.19E-12 |
| hsa-let-7b | -0.20 | 0.0652 | -0.35 | 3.00E-38 | -0.22 | 1.30E-08 |

| | | | | | | |
|---|---|---|---|---|---|---|
| hsa-miR-10b | -0.20 | 0.0636 | -0.26 | 4.49E-20 | -0.23 | 2.86E-09 |
| hsa-let-7c | -0.21 | 0.0575 | -0.29 | 2.96E-26 | -0.12 | 0.001842784 |
| hsa-miR-29c | -0.21 | 0.0542 | -0.26 | 3.88E-20 | -0.32 | 5.18E-17 |
| hsa-miR-195 | -0.21 | 0.0542 | -0.24 | 2.59E-18 | -0.21 | 1.26E-07 |
| hsa-miR-379 | -0.23 | 0.0392 | -0.35 | 5.77E-38 | -0.10 | 0.010781664 |
| hsa-miR-410 | -0.24 | 0.0312 | -0.29 | 4.66E-25 | -0.12 | 0.001884708 |
| hsa-miR-10a | -0.36 | 0.0010 | -0.30 | 3.59E-27 | -0.25 | 5.76E-11 |

**Table S13. miRNA profile associated with AGO2 expression in MYC neutral cases across independent breast cancer cohorts**

Only patients with MYC CN=2 (no deletion nor gain/amplification), and with matched genomic (CN), transcriptomic (both coding and non-coding mRNA) and clinical follow-up data were considered for each dataset.

| | **Oxford (N=111)** | | **Metabric (N=826)** | | **TCGA (N=806)** | |
|---|---|---|---|---|---|---|
| | Correlation | p-value (adjusted BH) | Correlation | p-value (adjusted BH) | Correlation | p-value (adjusted BH) |
| hsa-miR-150 | 0.28 | 0.0126 | 0.13 | 3.17E-04 | 0.18 | 5.66E-05 |
| hsa-miR-93 | 0.26 | 0.0243 | 0.13 | 1.91E-04 | 0.24 | 7.03E-08 |

| | | | | | | |
|---|---|---|---|---|---|---|
| hsa-miR-10a | -0.36 | 0.0010 | -0.27 | 8.26E-15 | -0.20 | 9.14E-06 |

**Table S14. miRNA expression in control HCC1806 cells compared with AGO2 silenced HCC1806 cells.**

Results of Limma analysis are shown, positive t values correspond to lower expression in AGO2 silenced cells (hence downregulated by loss of AGO2 expression), negative t values correspond to higher expression in AGO2 silenced (hence upregulated by loss of AGO2 expression). See Methods for further experimental and analysis details.

| GeneName | ProbeName | AveExpr | t | P.Value | adj.P.Val | B |
|---|---|---|---|---|---|---|
| hsa-let-7i-5p | A_25_P00012145 | 316.11 | 18.01 | 1.14E-13 | 6.72E-10 | 16.52 |
| hsa-miR-4291 | A_25_P00015568 | 226.12 | 17.19 | 2.70E-13 | 7.95E-10 | 16.08 |
| hsa-miR-324-5p | A_25_P00010154 | 271.75 | 16.47 | 5.94E-13 | 1.17E-09 | 15.65 |
| hsa-miR-183-5p | A_25_P00012098 | 227.92 | 14.89 | 3.73E-12 | 5.49E-09 | 14.61 |
| hsa-miR-324-5p | A_25_P00010153 | 384.54 | 14.53 | 5.84E-12 | 6.75E-09 | 14.34 |
| hsa-let-7i-5p | A_25_P00012146 | 460.68 | 14.40 | 6.88E-12 | 6.75E-09 | 14.24 |
| hsa-miR-24-3p | A_25_P00010677 | 4480.34 | 14.05 | 1.07E-11 | 8.99E-09 | 13.97 |
| hsa-miR-4291 | A_25_P00015567 | 162.93 | 13.62 | 1.86E-11 | 1.37E-08 | 13.63 |
| hsa-miR-148b-3p | A_25_P00010133 | 159.54 | 12.40 | 9.72E-11 | 6.36E-08 | 12.55 |
| hsa-miR-24-3p | A_25_P00010676 | 5186.72 | 12.27 | 1.17E-10 | 6.90E-08 | 12.42 |
| hsa-let-7d-5p | A_25_P00011980 | 258.87 | 12.09 | 1.52E-10 | 8.17E-08 | 12.24 |
| hsa-miR-183-5p | A_25_P00012099 | 231.41 | 11.93 | 1.92E-10 | 9.45E-08 | 12.08 |
| hsa-miR-429 | A_25_P00010273 | 526.06 | 11.75 | 2.51E-10 | 1.09E-07 | 11.90 |
| hsa-miR-4520-2-3p | A_25_P00016397 | 40.68 | 11.72 | 2.60E-10 | 1.09E-07 | 11.88 |

| | | | | | | |
|---|---|---|---|---|---|---|
| hsa-miR-429 | A_25_P00014860 | 433.16 | 11.41 | 4.11E-10 | 1.61E-07 | 11.55 |
| hsa-miR-107 | A_25_P00011069 | 1001.22 | 10.78 | 1.10E-09 | 3.80E-07 | 10.85 |
| hsa-miR-221-3p | A_25_P00010689 | 243.73 | 10.35 | 2.14E-09 | 6.79E-07 | 10.36 |
| hsa-miR-107 | A_25_P00011068 | 407.83 | 10.34 | 2.19E-09 | 6.79E-07 | 10.34 |
| hsa-miR-183-5p | A_25_P00012097 | 106.71 | 9.98 | 3.93E-09 | 1.16E-06 | 9.90 |
| hsa-let-7g-5p | A_25_P00012142 | 563.62 | 9.90 | 4.49E-09 | 1.26E-06 | 9.80 |
| hsa-miR-99b-5p | A_25_P00014849 | 191.60 | 9.86 | 4.85E-09 | 1.30E-06 | 9.74 |
| hsa-miR-30d-5p | A_25_P00010682 | 158.87 | 9.72 | 6.10E-09 | 1.56E-06 | 9.57 |
| hsa-miR-1307-3p | A_25_P00015256 | 55.21 | 9.70 | 6.36E-09 | 1.56E-06 | 9.54 |
| hsa-miR-301a-3p | A_25_P00010839 | 258.07 | 9.56 | 7.97E-09 | 1.88E-06 | 9.36 |
| hsa-miR-320c | A_25_P00015037 | 360.35 | 9.25 | 1.37E-08 | 3.11E-06 | 8.94 |
| hsa-miR-505-3p | A_25_P00012654 | 62.74 | 9.04 | 1.97E-08 | 4.31E-06 | 8.66 |
| hsa-let-7d-5p | A_25_P00011981 | 620.69 | 8.92 | 2.45E-08 | 4.98E-06 | 8.49 |
| hsa-miR-342-3p | A_25_P00012358 | 174.11 | 8.76 | 3.28E-08 | 6.45E-06 | 8.26 |
| hsa-miR-4520-2-3p | A_25_P00016398 | 41.13 | 8.55 | 4.79E-08 | 9.11E-06 | 7.96 |
| hsa-let-7g-5p | A_25_P00012141 | 248.02 | 8.50 | 5.24E-08 | 9.65E-06 | 7.88 |
| hsa-miR-138-5p | A_25_P00012170 | 108.65 | 8.47 | 5.58E-08 | 9.97E-06 | 7.83 |
| hsa-miR-3200-3p | A_25_P00015619 | 42.28 | 8.45 | 5.78E-08 | 1.00E-05 | 7.80 |
| hsa-miR-3200-3p | A_25_P00015620 | 48.50 | 8.12 | 1.06E-07 | 1.78E-05 | 7.31 |
| hsa-let-7e-5p | A_25_P00011984 | 499.08 | 7.88 | 1.67E-07 | 2.73E-05 | 6.94 |
| hsa-miR-99b-5p | A_25_P00010597 | 109.28 | 7.83 | 1.86E-07 | 2.96E-05 | 6.85 |
| hsa-miR-30d-5p | A_25_P00010683 | 129.74 | 7.79 | 2.01E-07 | 3.12E-05 | 6.79 |
| hsa-miR-320c | A_25_P00015036 | 65.68 | 7.68 | 2.49E-07 | 3.77E-05 | 6.61 |
| hsa-miR-505-5p | A_25_P00013608 | 51.38 | 7.59 | 2.94E-07 | 4.26E-05 | 6.47 |
| hsa-miR-183-3p | A_25_P00013323 | 69.85 | 7.59 | 2.96E-07 | 4.26E-05 | 6.47 |
| hsa-miR-106b-5p | A_25_P00010434 | 438.24 | 7.44 | 3.98E-07 | 5.59E-05 | 6.22 |

| | | | | | | |
|---|---|---|---|---|---|---|
| hsa-miR-98-5p | A_25_P00010047 | 127.77 | 7.24 | 5.86E-07 | 8.03E-05 | 5.90 |
| hsa-miR-222-3p | A_25_P00012126 | 246.51 | 7.18 | 6.59E-07 | 8.82E-05 | 5.80 |
| hsa-miR-23a-3p | A_25_P00010843 | 1294.36 | 7.14 | 7.25E-07 | 9.50E-05 | 5.72 |
| hsa-miR-221-3p | A_25_P00010690 | 859.49 | 7.06 | 8.45E-07 | 0.00010823 | 5.59 |
| hsa-miR-425-5p | A_25_P00014045 | 61.19 | 6.96 | 1.05E-06 | 0.00013142 | 5.41 |
| hsa-miR-103a-3p | A_25_P00011005 | 876.76 | 6.84 | 1.34E-06 | 0.00016466 | 5.20 |
| hsa-miR-138-5p | A_25_P00012169 | 68.32 | 6.81 | 1.42E-06 | 0.00017061 | 5.15 |
| hsa-miR-330-3p | A_25_P00011046 | 50.93 | 6.75 | 1.59E-06 | 0.00018481 | 5.05 |
| hsa-miR-17-5p | A_25_P00011991 | 173.72 | 6.75 | 1.60E-06 | 0.00018481 | 5.05 |
| hsa-miR-320d | A_25_P00015270 | 120.82 | 6.65 | 1.97E-06 | 0.00022336 | 4.87 |
| hsa-miR-660-5p | A_25_P00010460 | 44.53 | 6.59 | 2.23E-06 | 0.00024781 | 4.76 |
| hsa-miR-331-3p | A_25_P00014026 | 229.98 | 6.55 | 2.42E-06 | 0.00026388 | 4.69 |
| hsa-miR-362-5p | A_25_P00013984 | 57.84 | 6.50 | 2.71E-06 | 0.00028898 | 4.59 |
| hsa-miR-15b-5p | A_25_P00011102 | 1711.71 | 6.49 | 2.75E-06 | 0.00028898 | 4.58 |
| hsa-miR-320e | A_25_P00015664 | 360.91 | 6.44 | 3.03E-06 | 0.00031326 | 4.50 |
| hsa-miR-362-5p | A_25_P00013983 | 42.79 | 6.43 | 3.10E-06 | 0.00031485 | 4.48 |
| hsa-miR-301b-3p | A_25_P00013027 | 112.87 | 6.39 | 3.42E-06 | 0.00034164 | 4.39 |
| hsa-miR-103a-3p | A_25_P00011004 | 2138.82 | 6.33 | 3.87E-06 | 0.00037373 | 4.29 |
| hsa-miR-301a-3p | A_25_P00013973 | 91.66 | 6.10 | 6.38E-06 | 0.00059702 | 3.85 |
| hsa-miR-23b-3p | A_25_P00010882 | 121.66 | 6.05 | 7.02E-06 | 0.00064597 | 3.77 |
| hsa-miR-331-3p | A_25_P00014027 | 423.51 | 6.04 | 7.22E-06 | 0.00065502 | 3.75 |
| hsa-miR-342-3p | A_25_P00012357 | 89.58 | 5.88 | 1.02E-05 | 0.00091172 | 3.44 |
| hsa-miR-23b-3p | A_25_P00010881 | 344.01 | 5.84 | 1.10E-05 | 0.00093608 | 3.38 |
| hsa-miR-744-5p | A_25_P00012986 | 72.71 | 5.84 | 1.10E-05 | 0.00093608 | 3.38 |
| hsa-miR-182-5p | A_25_P00012095 | 87.58 | 5.84 | 1.11E-05 | 0.00093608 | 3.37 |
| hsa-miR-320d | A_25_P00015271 | 481.06 | 5.83 | 1.13E-05 | 0.00094104 | 3.35 |

| | | | | | | |
|---|---|---|---|---|---|---|
| hsa-miR-128-3p | A_25_P00012162 | 127.92 | 5.82 | 1.16E-05 | 0.00094618 | 3.34 |
| hsa-miR-301b-3p | A_25_P00013026 | 58.35 | 5.76 | 1.34E-05 | 0.00106347 | 3.21 |
| hsa-miR-106b-5p | A_25_P00010433 | 1084.13 | 5.74 | 1.39E-05 | 0.00107824 | 3.18 |
| hsa-miR-877-5p | A_25_P00012999 | 47.50 | 5.74 | 1.39E-05 | 0.00107824 | 3.17 |
| hsa-miR-505-5p | A_25_P00013607 | 46.73 | 5.72 | 1.44E-05 | 0.00110406 | 3.14 |
| hsa-miR-744-5p | A_25_P00012987 | 88.86 | 5.65 | 1.69E-05 | 0.00126632 | 3.00 |
| hsa-miR-652-3p | A_25_P00012834 | 65.00 | 5.64 | 1.73E-05 | 0.0012777 | 2.98 |
| hsa-miR-532-5p | A_25_P00014178 | 45.95 | 5.62 | 1.82E-05 | 0.00132152 | 2.94 |
| hsa-miR-148b-3p | A_25_P00010134 | 49.01 | 5.61 | 1.86E-05 | 0.00133515 | 2.92 |
| hsa-miR-105-5p | A_25_P00012043 | 67.27 | 5.55 | 2.13E-05 | 0.0014929 | 2.80 |
| hsa-miR-181a-5p | A_25_P00014832 | 158.37 | 5.50 | 2.33E-05 | 0.00161498 | 2.72 |
| hsa-miR-98-5p | A_25_P00010048 | 47.49 | 5.42 | 2.79E-05 | 0.00190865 | 2.56 |
| hsa-miR-182-5p | A_25_P00012094 | 52.39 | 5.41 | 2.88E-05 | 0.00195183 | 2.53 |
| hsa-miR-17-5p | A_25_P00013841 | 582.84 | 5.40 | 2.97E-05 | 0.00198774 | 2.51 |
| hsa-miR-128-3p | A_25_P00012161 | 64.53 | 5.39 | 3.04E-05 | 0.00201221 | 2.49 |
| hsa-miR-182-5p | A_25_P00012093 | 53.14 | 5.27 | 3.91E-05 | 0.00256154 | 2.26 |
| hsa-let-7c-5p | A_25_P00010073 | 120.83 | 5.24 | 4.23E-05 | 0.00273918 | 2.19 |
| hsa-miR-330-3p | A_25_P00014009 | 40.19 | 5.23 | 4.29E-05 | 0.00274967 | 2.18 |
| hsa-let-7f-5p | A_25_P00010089 | 519.69 | 5.14 | 5.24E-05 | 0.00331848 | 2.00 |
| hsa-miR-200a-3p | A_25_P00010208 | 153.96 | 5.12 | 5.47E-05 | 0.00342722 | 1.97 |
| hsa-miR-17-5p | A_25_P00014819 | 1723.12 | 5.08 | 5.98E-05 | 0.00367254 | 1.89 |
| hsa-miR-301b-3p | A_25_P00013025 | 43.92 | 5.07 | 6.20E-05 | 0.00376861 | 1.85 |
| hsa-miR-152-3p | A_25_P00012196 | 92.79 | 5.06 | 6.35E-05 | 0.00382139 | 1.83 |
| hsa-miR-320e | A_25_P00015663 | 83.00 | 4.89 | 9.26E-05 | 0.00524654 | 1.50 |
| hsa-miR-660-5p | A_25_P00010459 | 69.55 | 4.73 | 0.00013489 | 0.00727927 | 1.16 |
| hsa-miR-203a-3p | A_25_P00010628 | 141.79 | 4.73 | 0.00013588 | 0.00727927 | 1.15 |

| | | | | | | |
|---|---|---|---|---|---|---|
| hsa-miR-222-3p | A_25_P00012125 | 57.85 | 4.67 | 0.00015489 | 0.00814953 | 1.04 |
| hsa-let-7b-5p | A_25_P00010070 | 2052.89 | 4.62 | 0.00017268 | 0.00900538 | 0.94 |
| hsa-miR-769-5p | A_25_P00011955 | 50.88 | 4.58 | 0.00018946 | 0.00970857 | 0.86 |
| hsa-miR-135b-5p | A_25_P00012382 | 78.77 | 4.55 | 0.00020212 | 0.01026819 | 0.80 |
| hsa-miR-130a-3p | A_25_P00010440 | 1859.09 | 4.54 | 0.00020869 | 0.01051108 | 0.77 |
| hsa-miR-361-5p | A_25_P00010894 | 158.47 | 4.49 | 0.00023463 | 0.01161898 | 0.67 |
| hsa-miR-454-3p | A_25_P00012871 | 51.77 | 4.48 | 0.00024096 | 0.01180415 | 0.64 |
| hsa-miR-425-5p | A_25_P00010977 | 105.73 | 4.47 | 0.00024237 | 0.01180415 | 0.64 |
| hsa-miR-200a-3p | A_25_P00010209 | 505.37 | 4.45 | 0.00025821 | 0.01222481 | 0.58 |
| hsa-miR-532-5p | A_25_P00014179 | 60.27 | 4.44 | 0.00025931 | 0.01222481 | 0.58 |
| hsa-miR-31-5p | A_25_P00012019 | 186.62 | 4.39 | 0.00029634 | 0.01385965 | 0.46 |
| hsa-miR-93-5p | A_25_P00010610 | 1183.24 | 4.35 | 0.00032203 | 0.01482573 | 0.38 |
| hsa-miR-137 | A_25_P00010617 | 88.78 | 4.33 | 0.00033551 | 0.01532688 | 0.35 |
| hsa-miR-93-5p | A_25_P00010611 | 245.15 | 4.30 | 0.00036554 | 0.01644352 | 0.27 |
| hsa-miR-4690-5p | A_25_P00017268 | 49.53 | 4.29 | 0.00037271 | 0.01663927 | 0.25 |
| hsa-miR-769-5p | A_25_P00011954 | 66.14 | 4.23 | 0.00042155 | 0.01853879 | 0.14 |
| hsa-miR-30c-5p | A_25_P00010815 | 66.81 | 4.22 | 0.00043414 | 0.0189509 | 0.12 |
| hsa-let-7e-5p | A_25_P00011985 | 826.98 | 4.21 | 0.00045114 | 0.0195485 | 0.08 |
| hsa-miR-137 | A_25_P00010616 | 101.76 | 4.15 | 0.00051018 | 0.02162937 | -0.03 |
| hsa-miR-23a-3p | A_25_P00014820 | 4287.60 | 4.14 | 0.00053057 | 0.02233309 | -0.06 |
| hsa-miR-566 | A_25_P00010946 | 36.00 | 4.11 | 0.00055802 | 0.0231579 | -0.11 |
| hsa-let-7c-5p | A_25_P00010072 | 655.84 | 4.11 | 0.00056256 | 0.02318299 | -0.12 |
| hsa-miR-148a-3p | A_25_P00010131 | 185.95 | 4.05 | 0.00064323 | 0.02632345 | -0.24 |
| hsa-miR-96-5p | A_25_P00012034 | 326.40 | 4.02 | 0.00068952 | 0.02780877 | -0.30 |
| hsa-miR-135b-5p | A_25_P00012383 | 101.90 | 4.02 | 0.0006925 | 0.02780877 | -0.30 |
| hsa-miR-205-5p | A_25_P00010504 | 19172.69 | 4.02 | 0.00069369 | 0.02780877 | -0.31 |

| | | | | | | |
|---|---|---|---|---|---|---|
| hsa-miR-590-5p | A_25_P00014257 | 148.33 | 3.95 | 0.00082364 | 0.03257526 | -0.46 |
| hsa-miR-203a-3p | A_25_P00010629 | 288.86 | 3.92 | 0.00086791 | 0.03388442 | -0.51 |
| hsa-miR-155-5p | A_25_P00012271 | 229.26 | 3.89 | 0.00094433 | 0.03661153 | -0.58 |
| hsa-miR-455-5p | A_25_P00014182 | 60.99 | 3.88 | 0.00095745 | 0.03687757 | -0.59 |
| hsa-miR-194-5p | A_25_P00011007 | 45.97 | 3.87 | 0.00098241 | 0.03759315 | -0.62 |
| hsa-miR-192-5p | A_25_P00010868 | 52.02 | 3.86 | 0.00100303 | 0.03806087 | -0.64 |
| hsa-miR-944 | A_25_P00013107 | 52.85 | 3.86 | 0.00100755 | 0.03806087 | -0.64 |
| hsa-miR-130b-3p | A_25_P00010437 | 497.29 | 3.82 | 0.00110352 | 0.04049837 | -0.72 |
| hsa-miR-24-1-5p | A_25_P00014494 | 43.21 | 3.81 | 0.00113963 | 0.04145587 | -0.75 |
| hsa-miR-96-5p | A_25_P00012035 | 468.42 | 3.80 | 0.00116006 | 0.04194004 | -0.77 |
| hsa-miR-186-5p | A_25_P00012243 | 47.71 | 3.79 | 0.00116882 | 0.04199899 | -0.77 |
| hsa-miR-877-5p | A_25_P00012998 | 43.87 | 3.78 | 0.001209 | 0.04317963 | -0.80 |
| hsa-miR-26b-5p | A_25_P00012001 | 186.61 | 3.78 | 0.00122052 | 0.04332838 | -0.81 |
| hsa-miR-7-1-3p | A_25_P00013295 | 61.99 | -3.73 | 0.00134254 | 0.04653858 | -0.90 |
| hsa-miR-6087 | A_25_P00017893 | 716.98 | -3.74 | 0.00133619 | 0.04653858 | -0.89 |
| hsa-miR-6800-5p | A_25_P00018316 | 180.37 | -3.74 | 0.00133284 | 0.04653858 | -0.89 |
| hsa-miR-5687 | A_25_P00017467 | 35.32 | -3.75 | 0.00129575 | 0.04572371 | -0.87 |
| hsa-miR-302b-3p | A_25_P00010620 | 37.52 | -3.82 | 0.00110644 | 0.04049837 | -0.72 |
| hsa-miR-3651 | A_25_P00016223 | 512.49 | -3.82 | 0.00109725 | 0.04049837 | -0.72 |
| hsa-miR-33b-3p | A_25_P00013651 | 47.22 | -3.83 | 0.00108881 | 0.04049837 | -0.71 |
| hsa-miR-3620-3p | A_25_P00016149 | 42.11 | -3.83 | 0.00106469 | 0.0399631 | -0.69 |
| hsa-miR-30e-3p | A_25_P00014610 | 48.44 | -3.92 | 0.00086824 | 0.03388442 | -0.51 |
| hsa-miR-6829-5p | A_25_P00019059 | 85.59 | -3.97 | 0.00078304 | 0.03117863 | -0.41 |
| hsa-miR-3653-5p | A_25_P00019153 | 37.08 | -4.13 | 0.00053518 | 0.02236731 | -0.07 |
| hsa-miR-1260a | A_25_P00015173 | 3216.88 | -4.16 | 0.00050381 | 0.02151426 | -0.02 |
| hsa-miR-1202 | A_25_P00015075 | 210.43 | -4.18 | 0.00047359 | 0.02037112 | 0.04 |

| | | | | | | |
|---|---|---|---|---|---|---|
| hsa-miR-6125 | A_25_P00017944 | 348.58 | -4.26 | 0.0003975 | 0.01761269 | 0.19 |
| hsa-miR-7704 | A_25_P00018224 | 345.68 | -4.33 | 0.00033949 | 0.01538948 | 0.34 |
| hsa-miR-205-3p | A_25_P00015381 | 127.02 | -4.36 | 0.00031687 | 0.01470311 | 0.40 |
| hsa-miR-451b | A_25_P00016355 | 38.97 | -4.46 | 0.00025223 | 0.01208445 | 0.60 |
| hsa-miR-1273g-3p | A_25_P00017530 | 2245.75 | -4.46 | 0.00025033 | 0.01208445 | 0.61 |
| hsa-miR-6869-5p | A_25_P00018777 | 380.42 | -4.49 | 0.0002328 | 0.01161898 | 0.67 |
| hsa-miR-451b | A_25_P00016354 | 38.78 | -4.61 | 0.00017522 | 0.00905743 | 0.93 |
| hsa-miR-4497 | A_25_P00016920 | 201.94 | -4.71 | 0.00014174 | 0.00752496 | 1.12 |
| hsa-miR-486-3p | A_25_P00012468 | 44.33 | -4.74 | 0.00013052 | 0.00712179 | 1.19 |
| hsa-miR-3162-5p | A_25_P00015692 | 481.50 | -4.82 | 0.00010851 | 0.0059763 | 1.36 |
| hsa-miR-4665-5p | A_25_P00016953 | 36.31 | -4.83 | 0.0001074 | 0.0059709 | 1.36 |
| hsa-miR-4459 | A_25_P00016804 | 1038.81 | -4.89 | 9.39E-05 | 0.00527174 | 1.48 |
| hsa-miR-6125 | A_25_P00017945 | 418.53 | -4.92 | 8.69E-05 | 0.00497223 | 1.55 |
| hsa-miR-3162-5p | A_25_P00015693 | 481.96 | -4.93 | 8.46E-05 | 0.00488501 | 1.58 |
| hsa-miR-6869-5p | A_25_P00018778 | 411.46 | -4.94 | 8.34E-05 | 0.00486376 | 1.59 |
| hsa-miR-4286 | A_25_P00015774 | 3845.49 | -4.98 | 7.54E-05 | 0.00444071 | 1.68 |
| hsa-miR-141-5p | A_25_P00013415 | 56.19 | -5.03 | 6.82E-05 | 0.0040623 | 1.77 |
| hsa-miR-92a-3p | A_25_P00012031 | 333.79 | -5.11 | 5.68E-05 | 0.00352524 | 1.93 |
| hsa-miR-6087 | A_25_P00017892 | 388.78 | -5.59 | 1.95E-05 | 0.00138126 | 2.88 |
| hsa-miR-22-5p | A_25_P00013178 | 46.43 | -5.65 | 1.70E-05 | 0.00126632 | 3.00 |
| hsa-miR-30e-3p | A_25_P00014611 | 78.78 | -5.82 | 1.17E-05 | 0.00094729 | 3.32 |
| hsa-miR-4286 | A_25_P00015773 | 310.55 | -5.86 | 1.06E-05 | 0.00092896 | 3.42 |
| hsa-miR-1202 | A_25_P00015076 | 217.28 | -6.14 | 5.79E-06 | 0.0005507 | 3.94 |
| hsa-miR-205-3p | A_25_P00015382 | 226.49 | -6.34 | 3.75E-06 | 0.0003681 | 4.32 |
| hsa-miR-19b-1-5p | A_25_P00013163 | 55.52 | -9.00 | 2.13E-08 | 4.48E-06 | 8.60 |
| hsa-miR-486-3p | A_25_P00012469 | 54.71 | -11.34 | 4.61E-10 | 1.70E-07 | 11.47 |

**Table S15: miRNA expression in control HCC1806 cells compared with PABPC1 silenced HCC1806 cells.**

Results of Limma analysis are shown, positive t values correspond to lower expression in PABPC1 silenced cells (hence downregulated by loss of PABPC1 expression), negative t values correspond to higher expression in PABPC1 silenced (hence upregulated by loss of PABPC1 expression). See Methods for further experimental and analysis details.

| GeneName | ProbeName | AveExpr | t | P.Value | adj.P.Val | B |
|---|---|---|---|---|---|---|
| hsa-miR-4520-2-3p | A_25_P00016397 | 40.68 | 11.60 | 3.13E-10 | 1.85E-06 | 11.63 |
| hsa-miR-4520-2-3p | A_25_P00016398 | 41.13 | 8.71 | 3.60E-08 | 7.07E-05 | 8.12 |
| hsa-miR-34a-5p | A_25_P00012086 | 299.81 | 6.50 | 2.69E-06 | 0.00224137 | 4.57 |
| hsa-miR-125a-5p | A_25_P00012209 | 401.88 | 5.48 | 2.49E-05 | 0.014648269 | 2.65 |
| hsa-miR-320c | A_25_P00015037 | 360.35 | 5.16 | 5.03E-05 | 0.02201374 | 2.03 |
| hsa-miR-34a-5p | A_25_P00012085 | 143.97 | 4.71 | 0.000140019 | 0.039292122 | 1.12 |
| hsa-miR-320e | A_25_P00015664 | 360.91 | 4.66 | 0.000159147 | 0.042629814 | 1.01 |
| hsa-miR-505-5p | A_25_P00013608 | 51.38 | -4.92 | 8.76E-05 | 0.025824456 | 1.54 |
| hsa-miR-6126 | A_25_P00017954 | 54.70 | -4.94 | 8.23E-05 | 0.025520742 | 1.59 |
| hsa-miR-944 | A_25_P00013107 | 52.85 | -4.98 | 7.56E-05 | 0.024735636 | 1.67 |
| hsa-miR-944 | A_25_P00013106 | 47.50 | -5.03 | 6.82E-05 | 0.023642512 | 1.76 |
| hsa-miR-4516 | A_25_P00016923 | 528.70 | -5.07 | 6.13E-05 | 0.022585923 | 1.85 |
| hsa-miR-25-3p | A_25_P00010990 | 190.76 | -5.11 | 5.60E-05 | 0.02201374 | 1.93 |
| hsa-miR-550a-3p | A_25_P00014661 | 47.74 | -5.12 | 5.59E-05 | 0.02201374 | 1.93 |
| hsa-miR-362-5p | A_25_P00013983 | 42.79 | -5.18 | 4.80E-05 | 0.02201374 | 2.07 |
| hsa-miR-7-1-3p | A_25_P00013294 | 45.98 | -5.33 | 3.45E-05 | 0.018507112 | 2.36 |

| | | | | | | |
|---|---|---|---|---|---|---|
| hsa-miR-584-5p | A_25_P00010634 | 63.68 | -6.39 | 3.42E-06 | 0.00224137 | 4.36 |
| hsa-miR-550a-3p | A_25_P00014662 | 49.36 | -6.39 | 3.39E-06 | 0.00224137 | 4.37 |
| hsa-miR-3200-3p | A_25_P00015620 | 48.50 | -6.62 | 2.11E-06 | 0.00206945 | 4.78 |
| hsa-miR-7-5p | A_25_P00012077 | 88.82 | -6.67 | 1.88E-06 | 0.00206945 | 4.87 |
| hsa-let-7d-5p | A_25_P00011980 | 258.87 | -7.89 | 1.65E-07 | 0.000242735 | 6.90 |
| hsa-miR-7-5p | A_25_P00012078 | 318.30 | -11.05 | 7.15E-10 | 2.11E-06 | 11.05 |

**Table S16: miRNA significantly positively associated with AGO2 expression in the TCGA breast cancer cohort, when comparing mature:precursor ratio (corrected for multiple testing by Bonferroni correction).**

| miRNA name | miRNA mature:immature expression ratio vs AGO2 gene expression<br><br>Adjusted P-value for positive correlation coefficient |
|---|---|
| hsa-miR-191-3p | $< 10^{-16}$ |
| hsa-miR-15b-3p | 6.55E-15 |
| hsa-miR-335-3p | 1.53E-11 |
| hsa-miR-3065-5p | 3.51E-11 |
| hsa-miR-744-3p | 5.74E-11 |
| hsa-miR-199a-3p | 1.91E-10 |
| hsa-miR-30c-5p | 1.39E-09 |

| | |
|---|---|
| hsa-miR-199b-3p | 1.60E-09 |
| hsa-miR-589-3p | 1.80E-09 |
| hsa-miR-185-3p | 8.28E-09 |
| hsa-miR-29b-3p | 1.19E-08 |
| hsa-miR-21-3p | 5.39E-08 |
| hsa-miR-224-5p | 8.83E-08 |
| hsa-miR-7-5p | 8.83E-08 |
| hsa-let-7a-3p | 1.58E-07 |
| hsa-miR-200c-5p | 1.68E-07 |
| hsa-miR-29a-5p | 1.73E-07 |
| hsa-miR-491-3p | 2.14E-07 |
| hsa-miR-125b-1-3p | 2.97E-07 |
| hsa-let-7g-3p | 4.73E-07 |
| hsa-miR-30e-5p | 4.86E-07 |
| hsa-miR-423-3p | 6.56E-07 |
| hsa-miR-29b-1-5p | 6.93E-07 |
| hsa-miR-431-5p | 1.21E-06 |

| | |
|---|---|
| hsa-let-7a-2-3p | 1.30E-06 |
| hsa-miR-330-5p | 1.73E-06 |
| hsa-miR-148b-5p | 7.99E-06 |
| hsa-miR-99b-3p | 8.34E-06 |
| hsa-let-7f-1-3p | 9.15E-06 |
| hsa-miR-125a-3p | 9.84E-06 |
| hsa-miR-340-3p | 1.18E-05 |
| hsa-miR-106b-3p | 1.24E-05 |
| hsa-miR-17-5p | 2.48E-05 |
| hsa-miR-24-3p | 3.45E-05 |
| hsa-miR-452-3p | 3.75E-05 |
| hsa-miR-377-3p | 3.79E-05 |
| hsa-let-7i-3p | 4.14E-05 |
| hsa-miR-34c-3p | 5.69E-05 |
| hsa-miR-455-3p | 5.93E-05 |
| hsa-miR-887-3p | 7.66E-05 |
| hsa-miR-497-3p | 8.09E-05 |

**Table S17. Synthetic dosage lethality with AGO2 amplification. Analysis of samples in the TCGA BRCA and Metabric cancer cohorts.**

| *TCGA BRCA cohort. AGO2 amplification/non-amplification versus mutation/WT of genes with mutation frequency >20%* | | | | |
|---|---|---|---|---|
| Gene (HUGO symb) | Contingency table frequencies (Gene MT in AGO2 CNA/GAIN, Gene MT in AGO2 WT/DEL, Gene WT in AGO2 CNA/GAIN, Gene WT in AGO2 WT/DEL) | Mutation (MT) Freq | pvalues | adjusted_pvalue |
| TP53 | (309,58,382,343) | 370/1106 (0.33) | 6.07E-26 | 4.25E-25 |
| PIK3CA | (195,211,500,225) | 408/1145 (0.36) | 7.34E-12 | 2.57E-11 |
| CDH1 | (55,82,620,321) | 140/1093 (0.13) | 1.47E-08 | 3.42E-08 |
| GATA3 | (69,66,604,340) | 138/1093 (0.13) | 0.004384 | 0.007673 |
| TTN | (201,153,538,329) | 364/1240 (0.29) | 0.093518 | 0.130925 |
| MAP3K1 | (89,48,614,367) | 137/1132 (0.12) | 0.63738 | 0.74361 |
| MUC16 | (100,58,603,357) | 158/1132 (0.14) | 0.92944 | 0.92944 |
| *TCGA BRCA cohort. AGO2 amplification/non-amplification versus deleterious mutation/WT of genes with mutation frequency >20%* | | | | |
| Gene (HUGO symb) | Contingency table frequencies (Gene MT in AGO2 CNA/GAIN, Gene MT in AGO2 WT/DEL, Gene WT in AGO2 CNA/GAIN, Gene WT in AGO2 WT/DEL) | Mutation (MT) Freq | pvalues | adjusted_pvalue |
| TP53 | (156,29,382,343) | 370/1106 (0.33) | 4.36E-16 | 3.05E-15 |
| PIK3CA | (82,101,500,225) | 408/1145 (0.36) | 2.98E-09 | 1.04E-08 |

| | | | | |
|---|---|---|---|---|
| GATA3 | (4,9,604,340) | 138/1093 (0.13) | 0.018868 | 0.044025 |
| MAP3K1 | (16,8,614,367) | 137/1132 (0.12) | 0.831693 | 1 |
| CDH1 | (8,4,620,321) | 140/1093 (0.13) | 1 | 1 |
| MUC16 | (NA,NA,603,357) | 158/1132 (0.14) | 1 | 1 |
| TTN | (NA,NA,538,329) | 364/1240 (0.29) | 1 | 1 |
| ***TCGA BRCA cohort. cMyc WT Neutral Cases. AGO2 amplification/non-amplification versus mutation/WT of genes with mutation frequency >20%*** | | | | |
| Gene (HUGO symb) | Contingency table frequencies (Gene MT in AGO2 CNA/GAIN, Gene MT in AGO2 WT/DEL, Gene WT in AGO2 CNA/GAIN, Gene WT in AGO2 WT/DEL) | Mutation (MT) Freq | pvalues | adjusted_pvalue |
| TP53 | (11,46,14,314) | 370/1104 (0.34) | 0.000254 | 0.001775 |
| TTN | (16,131,14,297) | 364/1238 (0.29) | 0.014316 | 0.050105 |
| MAP3K1 | (NA,48,25,326) | 137/1130 (0.12) | 0.057074 | 0.133173 |
| CDH1 | (2,76,23,286) | 140/1091 (0.13) | 0.193419 | 0.338483 |
| PIK3CA | (10,188,17,205) | 408/1143 (0.36) | 0.322303 | 0.451224 |
| MUC16 | (2,54,23,320) | 158/1130 (0.14) | 0.553914 | 0.646233 |
| GATA3 | (3,59,22,306) | 138/1092 (0.13) | 0.779803 | 0.779803 |
| ***TCGA BRCA cohort. cMyc WT Neutral Cases. AGO2 amplification/non-amplification versus deleterious mutation/WT of genes with mutation frequency >20%*** | | | | |
| Gene (HUGO symb) | Contingency table frequencies (Gene MT in AGO2 CNA/GAIN, Gene MT in AGO2 WT/DEL, Gene WT in AGO2 CNA/GAIN, Gene WT in AGO2 WT/DEL) | Mutation (MT) Freq | pvalues | adjusted_pvalue |
| TP53 | (7,22,14,314) | 370/1104 (0.34) | 0.00059 | 0.004129 |
| PIK3CA | (4,93,17,205) | 408/1143 (0.36) | 0.328408 | 1 |
| CDH1 | (NA,3,23,286) | 140/1091 (0.13) | 1 | 1 |

| | | | | |
|---|---|---|---|---|
| GATA3 | (NA,8,22,306) | 138/1092 (0.13) | 1 | 1 |
| MAP3K1 | (NA,8,25,326) | 137/1130 (0.12) | 1 | 1 |
| MUC16 | (NA,NA,23,320) | 158/1130 (0.14) | 1 | 1 |
| TTN | (NA,NA,14,297) | 364/1238 (0.29) | 1 | 1 |
| ***Metabric cohort. AGO2 amplification/non-amplification versus mutation/WT of genes with mutation frequency >20%*** | | | | |
| Gene (HUGO symb) | Contingency table frequencies (Gene MT in AGO2 CNA/GAIN, Gene MT in AGO2 WT/DEL, Gene WT in AGO2 CNA/GAIN, Gene WT in AGO2 WT/DEL) | Mutation (MT) Freq | pvalues | adjusted_pvalue |
| TP53 | (450,315,506,926) | 888/2320 (0.38) | 3.80E-25 | 5.43E-26 |
| PIK3CA | (363,620,621,697) | 1112/2430 (0.46) | 4.11E-06 | 1.17E-06 |
| CDH1 | (67,137,878,1097) | 243/2218 (0.11) | 0.003262 | 0.001398 |
| AHNAK | (101,114,850,1132) | 246/2228 (0.11) | 0.463387 | 0.277052 |
| MUC16 | (196,236,777,1042) | 499/2318 (0.22) | 0.463387 | 0.330991 |
| AHNAK2 | (195,273,801,1034) | 537/2372 (0.23) | 0.541822 | 0.464419 |
| MAP3K1 | (127,171,858,1105) | 337/2300 (0.15) | 0.754069 | 0.754069 |
| ***Metabric cohort. cMyc WT Neutral Cases. AGO2 amplification/non-amplification versus mutation/WT of genes with mutation frequency >20%*** | | | | |
| Gene (HUGO symb) | Contingency table frequencies (Gene MT in AGO2 CNA/GAIN, Gene MT in AGO2 WT/DEL, Gene WT in AGO2 CNA/GAIN, Gene WT in AGO2 WT/DEL) | Mutation (MT) Freq | pvalues | adjusted_pvalue |
| TP53 | (20,220,24,851) | 888/2255 (0.39) | 0.000273 | 0.001909 |
| AHNAK2 | (19,236,32,896) | 537/2266 (0.24) | 0.008428 | 0.029496 |
| CDH1 | (1,130,43,940) | 243/2105 (0.12) | 0.05225 | 0.121918 |

| | | | | |
|---|---|---|---|---|
| MAP3K1 | (4,162,41,949) | 337/2185 (0.15) | 0.386294 | 0.676014 |
| PIK3CA | (25,556,22,591) | 1112/2338 (0.48) | 0.55419 | 0.775866 |
| MUC16 | (6,194,38,908) | 499/2211 (0.23) | 0.684778 | 0.798908 |
| AHNAK | (4,107,40,975) | 246/2113 (0.12) | 1 | 1 |

**Table S18. Gene expression profile associated with PABPC1 expression levels.**

Top 500 significant (Bonferroni adjusted p<0.05) positively correlated with PABPC1 in the TCGA BRCA dataset (Table S2) are shown. Top and bottom 95% quantiles, generated by 100x bbootstrap resampling, are shown. See Supplementary methods for more details.

| Gene ID | rho | rho 2.5% | rho 97.5% | P |
|---|---|---|---|---|
| PABPC3 | 0.88474099 | 0.85895417 | 0.90517278 | 0 |
| EIF3E | 0.60814235 | 0.56155423 | 0.65338531 | 3.50E-102 |
| PABPC4 | 0.54917524 | 0.51086936 | 0.58850406 | 7.44E-80 |
| PTDSS1 | 0.53524199 | 0.49286614 | 0.57645517 | 3.32E-75 |
| RPL7 | 0.52591744 | 0.48545024 | 0.57304067 | 3.26E-72 |
| RPL30 | 0.48541152 | 0.42245381 | 0.53784391 | 3.05E-60 |
| DCAF13 | 0.46937311 | 0.42037246 | 0.5398576 | 6.28E-56 |
| RPL5 | 0.45992802 | 0.42232061 | 0.50404334 | 1.71E-53 |
| RPS7 | 0.45783352 | 0.41453301 | 0.49972873 | 5.79E-53 |
| RPS27A | 0.44454682 | 0.38939441 | 0.49575708 | 1.09E-49 |
| TAF1D | 0.44187871 | 0.39111928 | 0.48058805 | 4.77E-49 |
| RPS25 | 0.42965491 | 0.38417799 | 0.47738238 | 3.47E-46 |
| SERBP1 | 0.42749941 | 0.37797263 | 0.46858678 | 1.08E-45 |

| | | | | |
|---|---|---|---|---|
| TATDN1 | 0.42677227 | 0.36831585 | 0.49245117 | 1.58E-45 |
| RPS12 | 0.42497945 | 0.36408128 | 0.47647127 | 4.02E-45 |
| EEF1B2 | 0.42354224 | 0.36756698 | 0.46843908 | 8.48E-45 |
| RPS24 | 0.42221563 | 0.36683503 | 0.47629719 | 1.68E-44 |
| RPS18 | 0.41992852 | 0.36917704 | 0.46748295 | 5.44E-44 |
| PDE7A | 0.41739318 | 0.36145954 | 0.46355073 | 1.98E-43 |
| MKI67IP | 0.41671826 | 0.35627646 | 0.46423979 | 2.78E-43 |
| YWHAZ | 0.41280372 | 0.34072694 | 0.47719163 | 1.99E-42 |
| RPL4 | 0.41080002 | 0.34541339 | 0.45810171 | 5.38E-42 |
| EEF1G | 0.40884088 | 0.35111286 | 0.46583074 | 1.42E-41 |
| ZNF706 | 0.40758937 | 0.33704988 | 0.46252341 | 2.62E-41 |
| NUDCD1 | 0.40517384 | 0.35210214 | 0.46761683 | 8.52E-41 |
| RPLP0 | 0.40504439 | 0.35017851 | 0.46267856 | 9.07E-41 |
| RPS8 | 0.40419698 | 0.36108977 | 0.45458183 | 1.37E-40 |
| YBX1 | 0.39984162 | 0.3551649 | 0.44385192 | 1.11E-39 |
| WDR43 | 0.39819823 | 0.3385972 | 0.45017763 | 2.43E-39 |
| TEX10 | 0.39601968 | 0.33602888 | 0.4430563 | 6.82E-39 |
| RPS20 | 0.39369304 | 0.34435019 | 0.45069293 | 2.03E-38 |
| RPL6 | 0.39309941 | 0.33923065 | 0.44352923 | 2.68E-38 |
| RPL12 | 0.39266268 | 0.32841583 | 0.45083216 | 3.29E-38 |
| TAF4B | 0.3896171 | 0.33118542 | 0.43358116 | 1.35E-37 |
| NAP1L1 | 0.3890623 | 0.32346801 | 0.43672212 | 1.74E-37 |
| PTPN2 | 0.38812721 | 0.33706152 | 0.44308192 | 2.67E-37 |
| RPS10 | 0.38702168 | 0.3353936 | 0.44639522 | 4.43E-37 |
| RPS6 | 0.3839847 | 0.32608332 | 0.43287432 | 1.76E-36 |

| | | | | |
|---|---|---|---|---|
| INTS8 | 0.38306379 | 0.3143376 | 0.42895625 | 2.67E-36 |
| EEF1A1 | 0.38230341 | 0.32054685 | 0.42708619 | 3.76E-36 |
| POLR1E | 0.38032496 | 0.31407582 | 0.43037315 | 9.12E-36 |
| PPA1 | 0.378455 | 0.32274511 | 0.43661897 | 2.10E-35 |
| MPP6 | 0.37811307 | 0.32462464 | 0.43313122 | 2.44E-35 |
| KCMF1 | 0.37585227 | 0.32256667 | 0.43098832 | 6.61E-35 |
| RIPK2 | 0.37462136 | 0.3017932 | 0.42856039 | 1.13E-34 |
| RPS2 | 0.37171832 | 0.3223298 | 0.4198936 | 4.01E-34 |
| RPL17 | 0.37070557 | 0.30563646 | 0.42136935 | 6.22E-34 |
| SLC5A6 | 0.3705287 | 0.31581284 | 0.43970586 | 6.71E-34 |
| SLC25A32 | 0.36908925 | 0.30117447 | 0.41990469 | 1.25E-33 |
| MTERFD1 | 0.36835057 | 0.31392711 | 0.42068379 | 1.71E-33 |
| CCNB1IP1 | 0.36788255 | 0.32668199 | 0.42731799 | 2.09E-33 |
| PUS3 | 0.36700681 | 0.32088779 | 0.42545381 | 3.04E-33 |
| MEMO1 | 0.36699681 | 0.31717851 | 0.40982355 | 3.05E-33 |
| TMEM123 | 0.36528353 | 0.30970469 | 0.41134602 | 6.31E-33 |
| TAF5 | 0.36425877 | 0.31189203 | 0.41947714 | 9.74E-33 |
| KIAA0020 | 0.36378101 | 0.30887989 | 0.41676704 | 1.19E-32 |
| RPL35A | 0.36357124 | 0.29250156 | 0.4144892 | 1.30E-32 |
| UQCRB | 0.36341289 | 0.29550078 | 0.42454664 | 1.39E-32 |
| RPSAP58 | 0.3630639 | 0.30329909 | 0.41585651 | 1.61E-32 |
| C8orf59 | 0.35863992 | 0.29670512 | 0.41480002 | 1.02E-31 |
| RPS29 | 0.35580309 | 0.3067762 | 0.41534939 | 3.27E-31 |
| RPS3A | 0.35513754 | 0.2927149 | 0.42111595 | 4.29E-31 |
| EIF3L | 0.35395858 | 0.29296157 | 0.40559559 | 6.94E-31 |

| | | | | |
|---|---|---|---|---|
| RIOK1 | 0.35339046 | 0.28649477 | 0.4054801 | 8.74E-31 |
| RPL32 | 0.35319579 | 0.30011227 | 0.40734272 | 9.46E-31 |
| SEH1L | 0.35311079 | 0.30045411 | 0.41107683 | 9.79E-31 |
| PDSS1 | 0.35277746 | 0.30640421 | 0.4143527 | 1.12E-30 |
| RPL39 | 0.35168647 | 0.29747623 | 0.41886954 | 1.74E-30 |
| NOP58 | 0.35143093 | 0.29734937 | 0.40684362 | 1.93E-30 |
| EIF2S3 | 0.35086943 | 0.28157097 | 0.41822742 | 2.42E-30 |
| DKC1 | 0.35074795 | 0.3006642 | 0.40206782 | 2.54E-30 |
| LMO4 | 0.35024716 | 0.29322312 | 0.40789823 | 3.11E-30 |
| WDR12 | 0.34968516 | 0.28918647 | 0.4002918 | 3.89E-30 |
| RPL22 | 0.34948148 | 0.30302048 | 0.39118267 | 4.22E-30 |
| RPL18A | 0.34908523 | 0.29775797 | 0.39960876 | 4.95E-30 |
| FAM60A | 0.34881808 | 0.29230107 | 0.40908477 | 5.51E-30 |
| PRKX | 0.3486029 | 0.29071211 | 0.41115313 | 6.00E-30 |
| RPL10A | 0.34763887 | 0.27582974 | 0.40642272 | 8.81E-30 |
| RDH10 | 0.34664906 | 0.29728135 | 0.40272135 | 1.30E-29 |
| SNORA8 | 0.34638755 | 0.29122839 | 0.39135947 | 1.45E-29 |
| RPS4X | 0.34612325 | 0.28238192 | 0.39588937 | 1.61E-29 |
| RPL37 | 0.34239602 | 0.28937071 | 0.4079502 | 6.94E-29 |
| GPR126 | 0.34195262 | 0.28049349 | 0.41033698 | 8.25E-29 |
| SRPK1 | 0.34104374 | 0.29250949 | 0.38672591 | 1.17E-28 |
| NT5C2 | 0.34099634 | 0.28592954 | 0.40153127 | 1.20E-28 |
| RPS17 | 0.34076112 | 0.30567326 | 0.40286831 | 1.31E-28 |
| RPL7A | 0.34039824 | 0.26919578 | 0.39674361 | 1.51E-28 |
| CSDA | 0.34028292 | 0.26943264 | 0.39231613 | 1.58E-28 |

| | | | | |
|---|---|---|---|---|
| RPL31 | 0.33821799 | 0.2722331 | 0.39022432 | 3.49E-28 |
| MSL3 | 0.33762271 | 0.28225279 | 0.39396066 | 4.39E-28 |
| ENO1 | 0.33707471 | 0.28525572 | 0.39386044 | 5.42E-28 |
| GDI2 | 0.33705524 | 0.26825072 | 0.39116509 | 5.46E-28 |
| A2ML1 | 0.33574837 | 0.28203699 | 0.38793408 | 8.98E-28 |
| ADAT2 | 0.33542534 | 0.29028079 | 0.38940705 | 1.02E-27 |
| UCK2 | 0.3351901 | 0.27335852 | 0.39996588 | 1.11E-27 |
| RPSA | 0.33516063 | 0.26683129 | 0.41254539 | 1.12E-27 |
| C10orf2 | 0.33477244 | 0.26644715 | 0.38979656 | 1.30E-27 |
| TBPL1 | 0.33451928 | 0.27837714 | 0.38600177 | 1.43E-27 |
| ESRP1 | 0.33440433 | 0.28427876 | 0.38762589 | 1.50E-27 |
| MINA | 0.33244613 | 0.27203324 | 0.38231241 | 3.13E-27 |
| RPS19 | 0.33199208 | 0.27890138 | 0.39023329 | 3.72E-27 |
| RPS15A | 0.33130274 | 0.29432862 | 0.39079767 | 4.81E-27 |
| ETV6 | 0.33092475 | 0.27962815 | 0.37842641 | 5.54E-27 |
| ZNF22 | 0.32994053 | 0.27857745 | 0.38235079 | 8.00E-27 |
| YES1 | 0.32954763 | 0.2722917 | 0.38544368 | 9.26E-27 |
| NCL | 0.32773429 | 0.28487617 | 0.37010512 | 1.81E-26 |
| ANP32B | 0.3276868 | 0.26533187 | 0.39462925 | 1.85E-26 |
| CDCA7 | 0.32755299 | 0.2703804 | 0.37946962 | 1.94E-26 |
| EIF3H | 0.32646241 | 0.2642104 | 0.3759103 | 2.90E-26 |
| RARRES1 | 0.32493083 | 0.27149636 | 0.38333671 | 5.08E-26 |
| PUS7 | 0.32490674 | 0.26600682 | 0.38393725 | 5.13E-26 |
| AMD1 | 0.32422528 | 0.27163739 | 0.36315901 | 6.58E-26 |
| SUV39H2 | 0.3241232 | 0.25505513 | 0.38559719 | 6.83E-26 |

| | | | | |
|---|---|---|---|---|
| NOLC1 | 0.32351182 | 0.26783364 | 0.3815574 | 8.53E-26 |
| PELI1 | 0.32294333 | 0.25499607 | 0.37849028 | 1.05E-25 |
| USP6NL | 0.32263247 | 0.2658608 | 0.3718084 | 1.17E-25 |
| NRBF2 | 0.32188379 | 0.25445463 | 0.36067146 | 1.54E-25 |
| UTP14A | 0.32174213 | 0.2640777 | 0.37684292 | 1.62E-25 |
| POP1 | 0.32126468 | 0.25362128 | 0.3794902 | 1.92E-25 |
| PADI2 | 0.32098445 | 0.25227094 | 0.38483588 | 2.13E-25 |
| RRP1B | 0.31972511 | 0.2562227 | 0.36259604 | 3.35E-25 |
| NUS1 | 0.3196077 | 0.26560473 | 0.37379967 | 3.49E-25 |
| SRD5A1 | 0.31927724 | 0.27615291 | 0.36502569 | 3.93E-25 |
| MTAP | 0.31894203 | 0.25421689 | 0.36689402 | 4.43E-25 |
| PPRC1 | 0.31871517 | 0.26620006 | 0.36172301 | 4.80E-25 |
| CDK8 | 0.3184919 | 0.27280839 | 0.38844657 | 5.20E-25 |
| RPL10 | 0.31820208 | 0.24720984 | 0.3773846 | 5.77E-25 |
| MPZL1 | 0.31818677 | 0.26224255 | 0.37896691 | 5.80E-25 |
| WDR75 | 0.31789354 | 0.26079766 | 0.3688527 | 6.44E-25 |
| E2F3 | 0.31779556 | 0.27029216 | 0.36051098 | 6.67E-25 |
| C1orf106 | 0.31708713 | 0.24876548 | 0.36734043 | 8.57E-25 |
| RPS16 | 0.3165404 | 0.25717079 | 0.37585269 | 1.04E-24 |
| UBE2J1 | 0.31651252 | 0.25628152 | 0.3605938 | 1.05E-24 |
| MPZL2 | 0.31576543 | 0.25753131 | 0.35950196 | 1.37E-24 |
| FBL | 0.31556248 | 0.25271286 | 0.37767849 | 1.47E-24 |
| RPL27A | 0.31524119 | 0.24270804 | 0.38492666 | 1.65E-24 |
| MID1 | 0.31507569 | 0.26565418 | 0.37804641 | 1.75E-24 |
| ZNF695 | 0.31493167 | 0.26728295 | 0.36306642 | 1.84E-24 |

| | | | | |
|---|---|---|---|---|
| RPF2 | 0.3148518 | 0.25910319 | 0.36659196 | 1.89E-24 |
| SKP2 | 0.31464456 | 0.24914771 | 0.37043922 | 2.03E-24 |
| NFIB | 0.31434805 | 0.26356472 | 0.37218211 | 2.26E-24 |
| ZNF146 | 0.31431108 | 0.25418525 | 0.36894866 | 2.28E-24 |
| RPL11 | 0.31392629 | 0.25322669 | 0.36217846 | 2.62E-24 |
| SNHG6 | 0.31327709 | 0.25620079 | 0.36670901 | 3.28E-24 |
| HNRNPA1L2 | 0.31316602 | 0.25238148 | 0.36362899 | 3.41E-24 |
| RPS3 | 0.31255379 | 0.26313193 | 0.36757785 | 4.23E-24 |
| FRAT2 | 0.31084916 | 0.24975739 | 0.36533045 | 7.65E-24 |
| TIMM8A | 0.31082689 | 0.24629347 | 0.37302236 | 7.70E-24 |
| CSDAP1 | 0.31074019 | 0.25527483 | 0.37052614 | 7.94E-24 |
| PPPDE2 | 0.31063982 | 0.23811889 | 0.36412342 | 8.22E-24 |
| SNX3 | 0.31034434 | 0.26225283 | 0.36590946 | 9.11E-24 |
| OBFC2A | 0.30972625 | 0.24791169 | 0.35971293 | 1.13E-23 |
| CSDE1 | 0.30965076 | 0.26241756 | 0.37266647 | 1.16E-23 |
| NPM1 | 0.30963943 | 0.2605471 | 0.37298875 | 1.16E-23 |
| SNHG1 | 0.30923097 | 0.24206145 | 0.37097879 | 1.34E-23 |
| DDX10 | 0.30919994 | 0.25111952 | 0.37072886 | 1.35E-23 |
| ZNF93 | 0.30917693 | 0.26393136 | 0.36248715 | 1.36E-23 |
| EIF3M | 0.30866149 | 0.23935629 | 0.35892258 | 1.63E-23 |
| BCL11A | 0.30826316 | 0.25489453 | 0.34671791 | 1.86E-23 |
| NHSL1 | 0.30822569 | 0.25375592 | 0.36839497 | 1.89E-23 |
| TGS1 | 0.30810014 | 0.25165978 | 0.37107861 | 1.97E-23 |
| USP39 | 0.30753174 | 0.25715825 | 0.3649877 | 2.40E-23 |
| NOL10 | 0.30680993 | 0.24879612 | 0.36715274 | 3.07E-23 |

| | | | | |
|---|---|---|---|---|
| SET | 0.30652536 | 0.24352925 | 0.36472914 | 3.38E-23 |
| RPL24 | 0.30651881 | 0.24185317 | 0.36432082 | 3.39E-23 |
| EBPL | 0.30643895 | 0.24281855 | 0.35065107 | 3.48E-23 |
| LTV1 | 0.30570129 | 0.24187566 | 0.37155255 | 4.47E-23 |
| RPS13 | 0.30530128 | 0.24680857 | 0.35781862 | 5.12E-23 |
| PROM1 | 0.30503875 | 0.24454908 | 0.36667695 | 5.60E-23 |
| SNHG4 | 0.30473368 | 0.24700383 | 0.36194196 | 6.21E-23 |
| RPL15 | 0.30458035 | 0.25775011 | 0.35264602 | 6.54E-23 |
| FAM136A | 0.30450948 | 0.23858934 | 0.35875035 | 6.70E-23 |
| EML4 | 0.30400659 | 0.25283014 | 0.35432857 | 7.94E-23 |
| SYNCRIP | 0.30349424 | 0.24984765 | 0.34354025 | 9.44E-23 |
| ULBP3 | 0.30338559 | 0.24036497 | 0.36401354 | 9.79E-23 |
| PSAT1 | 0.30330227 | 0.24218067 | 0.36184917 | 1.01E-22 |
| TPT1 | 0.30298136 | 0.25516064 | 0.34909611 | 1.12E-22 |
| MTHFD2 | 0.3028116 | 0.24751575 | 0.36147531 | 1.19E-22 |
| UQCRH | 0.30276754 | 0.24109781 | 0.35170178 | 1.20E-22 |
| MRPL15 | 0.30256254 | 0.23614586 | 0.34972219 | 1.29E-22 |
| ZC3H8 | 0.30229067 | 0.22870722 | 0.36165074 | 1.41E-22 |
| TMEM65 | 0.30202768 | 0.2471423 | 0.36695631 | 1.54E-22 |
| CTPS | 0.30196929 | 0.24789314 | 0.35748855 | 1.57E-22 |
| EI24 | 0.30170734 | 0.24846427 | 0.35769512 | 1.72E-22 |
| RSU1 | 0.30162063 | 0.24617059 | 0.35918561 | 1.77E-22 |
| EIF5B | 0.3013915 | 0.23762948 | 0.3609563 | 1.91E-22 |
| GPR156 | 0.30138573 | 0.2430637 | 0.36573058 | 1.91E-22 |
| SND1 | 0.30057613 | 0.24448314 | 0.3506032 | 2.51E-22 |

| | | | | |
|---|---|---|---|---|
| UQCRHL | 0.30042448 | 0.24691206 | 0.3567182 | 2.64E-22 |
| NFIL3 | 0.30037912 | 0.25475442 | 0.36011721 | 2.68E-22 |
| WDR3 | 0.30031746 | 0.24318463 | 0.35444698 | 2.73E-22 |
| PAK1IP1 | 0.30024391 | 0.25168738 | 0.34758498 | 2.80E-22 |
| PDCD11 | 0.29970279 | 0.24584441 | 0.35704957 | 3.35E-22 |
| OLA1 | 0.29917107 | 0.23609214 | 0.35869973 | 4.00E-22 |
| NHEJ1 | 0.29879346 | 0.24202218 | 0.3571084 | 4.53E-22 |
| TTLL4 | 0.29828055 | 0.24474654 | 0.34648116 | 5.37E-22 |
| NKRF | 0.29807501 | 0.22232534 | 0.35676023 | 5.74E-22 |
| NSMAF | 0.29742331 | 0.23790457 | 0.34881274 | 7.12E-22 |
| UTP11L | 0.2967663 | 0.24492446 | 0.35818496 | 8.83E-22 |
| EIF3D | 0.29659008 | 0.24086428 | 0.3497529 | 9.36E-22 |
| E2F5 | 0.29653533 | 0.24321242 | 0.35188286 | 9.53E-22 |
| MTDH | 0.29641591 | 0.22944482 | 0.35696575 | 9.91E-22 |
| GMPS | 0.2963351 | 0.24444009 | 0.3439311 | 1.02E-21 |
| CEP57 | 0.29632504 | 0.24286816 | 0.35309604 | 1.02E-21 |
| ZFAND1 | 0.29568741 | 0.23635413 | 0.36708249 | 1.26E-21 |
| ZNF124 | 0.29556558 | 0.23425564 | 0.35996392 | 1.31E-21 |
| SDHD | 0.29501278 | 0.24549366 | 0.34927468 | 1.57E-21 |
| PRPF38A | 0.2941239 | 0.24715302 | 0.35928029 | 2.09E-21 |
| TES | 0.29383925 | 0.22720294 | 0.34161825 | 2.30E-21 |
| IPO5 | 0.29364463 | 0.23337037 | 0.35924411 | 2.44E-21 |
| MEX3A | 0.29364042 | 0.23737866 | 0.34332627 | 2.45E-21 |
| SKI | 0.29358754 | 0.23819571 | 0.34528478 | 2.49E-21 |
| DBF4 | 0.29356995 | 0.24428657 | 0.35541066 | 2.50E-21 |

| | | | | |
|---|---|---|---|---|
| ODC1 | 0.29327245 | 0.22433549 | 0.36293375 | 2.76E-21 |
| HSPD1 | 0.29325187 | 0.24025466 | 0.33729826 | 2.78E-21 |
| KIAA1804 | 0.29313296 | 0.2389973 | 0.35082003 | 2.89E-21 |
| RPL3 | 0.29297651 | 0.22936951 | 0.34076228 | 3.03E-21 |
| ZNF202 | 0.29256313 | 0.24436112 | 0.3471258 | 3.47E-21 |
| AHCY | 0.29191773 | 0.22880394 | 0.34426562 | 4.27E-21 |
| GAS5 | 0.29173153 | 0.23961278 | 0.36466828 | 4.53E-21 |
| LAD1 | 0.29148057 | 0.22630532 | 0.35852677 | 4.91E-21 |
| RSL1D1 | 0.29100511 | 0.2269692 | 0.35765428 | 5.72E-21 |
| GATAD2A | 0.29076462 | 0.24037145 | 0.34748503 | 6.18E-21 |
| ABCE1 | 0.29001764 | 0.22265055 | 0.32930338 | 7.85E-21 |
| TIGD2 | 0.29001087 | 0.23168519 | 0.35659604 | 7.87E-21 |
| MRPL37 | 0.2899112 | 0.21248788 | 0.35692015 | 8.12E-21 |
| SUPV3L1 | 0.28986537 | 0.23347992 | 0.34390442 | 8.24E-21 |
| RPS11 | 0.28984657 | 0.22401698 | 0.34741192 | 8.29E-21 |
| MTMR2 | 0.28950265 | 0.23993371 | 0.35112949 | 9.25E-21 |
| C1orf198 | 0.28938427 | 0.22603537 | 0.33858852 | 9.61E-21 |
| MED30 | 0.28838519 | 0.22836867 | 0.34295033 | 1.32E-20 |
| CCT6A | 0.28816136 | 0.23043764 | 0.34077528 | 1.42E-20 |
| IVNS1ABP | 0.28803148 | 0.23933808 | 0.34874291 | 1.48E-20 |
| KATNA1 | 0.28781468 | 0.22492084 | 0.33407142 | 1.58E-20 |
| TRPV6 | 0.2877882 | 0.21982793 | 0.33355406 | 1.60E-20 |
| ZNF280C | 0.28767768 | 0.23454849 | 0.34284428 | 1.65E-20 |
| RPP40 | 0.28762308 | 0.22700122 | 0.34449426 | 1.68E-20 |
| TRIT1 | 0.28751057 | 0.234312 | 0.34854019 | 1.74E-20 |

| | | | | |
|---|---|---|---|---|
| KCNQ4 | 0.28715166 | 0.22608303 | 0.3465248 | 1.95E-20 |
| REXO2 | 0.28714864 | 0.24216679 | 0.33717939 | 1.95E-20 |
| ZNF259 | 0.28712565 | 0.22632049 | 0.33793339 | 1.97E-20 |
| UBE2E3 | 0.28710492 | 0.22777052 | 0.34131982 | 1.98E-20 |
| FAM98A | 0.28700985 | 0.23007573 | 0.3345047 | 2.04E-20 |
| LDHB | 0.28614976 | 0.22175751 | 0.34066959 | 2.68E-20 |
| CCNJ | 0.28576633 | 0.22819442 | 0.34258764 | 3.02E-20 |
| C9orf30 | 0.28572547 | 0.2326882 | 0.33415559 | 3.06E-20 |
| FARSB | 0.2854826 | 0.21663647 | 0.35365067 | 3.30E-20 |
| STARD7 | 0.28541361 | 0.23448447 | 0.34569604 | 3.37E-20 |
| STT3A | 0.28531531 | 0.23018509 | 0.34776562 | 3.48E-20 |
| UBE2D1 | 0.28519652 | 0.2225942 | 0.34644834 | 3.61E-20 |
| RAP2A | 0.28492097 | 0.20705705 | 0.33997783 | 3.93E-20 |
| NACA | 0.28487674 | 0.22998263 | 0.35127091 | 3.99E-20 |
| GGH | 0.28483191 | 0.20565802 | 0.34870882 | 4.04E-20 |
| LAPTM4B | 0.28445229 | 0.22137654 | 0.34143547 | 4.55E-20 |
| AZIN1 | 0.28441229 | 0.2364204 | 0.33477908 | 4.61E-20 |
| PNRC1 | 0.2842191 | 0.21597686 | 0.33395076 | 4.90E-20 |
| POLR3G | 0.28348532 | 0.23847582 | 0.34181191 | 6.16E-20 |
| RPL14 | 0.2832323 | 0.21062152 | 0.34742191 | 6.66E-20 |
| PTCD3 | 0.28319545 | 0.22708266 | 0.35100548 | 6.74E-20 |
| FAM123B | 0.28311863 | 0.21595666 | 0.34930437 | 6.90E-20 |
| IL22RA2 | 0.28305425 | 0.22086169 | 0.34605729 | 7.04E-20 |
| R3HDM1 | 0.28300351 | 0.22606875 | 0.33040526 | 7.15E-20 |
| SRM | 0.28277409 | 0.21987428 | 0.3393595 | 7.68E-20 |

| | | | | |
|---|---|---|---|---|
| EIF4EBP2 | 0.28246565 | 0.23764803 | 0.33313846 | 8.45E-20 |
| C11orf57 | 0.28192506 | 0.2266141 | 0.33015586 | 9.99E-20 |
| ERP44 | 0.28157399 | 0.23198477 | 0.33740493 | 1.11E-19 |
| TRAF3IP2 | 0.28144517 | 0.23219016 | 0.34352319 | 1.16E-19 |
| TMEM74 | 0.28093082 | 0.22716256 | 0.35983858 | 1.36E-19 |
| IL12RB2 | 0.28047876 | 0.22459677 | 0.3315868 | 1.56E-19 |
| CHAC2 | 0.28004611 | 0.21761201 | 0.34610717 | 1.78E-19 |
| SLC35F2 | 0.27970526 | 0.22654972 | 0.34472865 | 1.98E-19 |
| HMGA1 | 0.27956993 | 0.22373024 | 0.32968182 | 2.06E-19 |
| RNF138 | 0.27945022 | 0.22293989 | 0.33728766 | 2.14E-19 |
| CCNY | 0.27938183 | 0.22579279 | 0.32749959 | 2.18E-19 |
| MICAL3 | 0.27924457 | 0.20994813 | 0.33424683 | 2.28E-19 |
| OSBPL3 | 0.2792441 | 0.23344085 | 0.32946327 | 2.28E-19 |
| RPL22L1 | 0.27875433 | 0.21940384 | 0.33923221 | 2.64E-19 |
| JRKL | 0.27863396 | 0.22141344 | 0.33023021 | 2.74E-19 |
| SSRP1 | 0.27836333 | 0.2090467 | 0.33204038 | 2.98E-19 |
| RPL37A | 0.27834595 | 0.22478986 | 0.33312371 | 3.00E-19 |
| ADSL | 0.27818164 | 0.21776783 | 0.32795833 | 3.15E-19 |
| NCAPD2 | 0.27772148 | 0.21951251 | 0.33852062 | 3.62E-19 |
| RPL13A | 0.27744314 | 0.20069972 | 0.35706594 | 3.94E-19 |
| ACTR3 | 0.27740053 | 0.21955801 | 0.32802318 | 3.99E-19 |
| ALDH18A1 | 0.27733188 | 0.21975704 | 0.33219704 | 4.08E-19 |
| HNRNPA1 | 0.27720536 | 0.21874514 | 0.32078016 | 4.24E-19 |
| FETUB | 0.27661431 | 0.207761 | 0.33755368 | 5.07E-19 |
| DDX6 | 0.2764801 | 0.22671342 | 0.33466515 | 5.28E-19 |

| | | | | |
|---|---|---|---|---|
| ZBED4 | 0.2761623 | 0.21353387 | 0.34213897 | 5.81E-19 |
| LBR | 0.27604933 | 0.2212469 | 0.33788475 | 6.01E-19 |
| PITPNB | 0.27558901 | 0.21990887 | 0.333853 | 6.90E-19 |
| B3GNT5 | 0.27556857 | 0.21835636 | 0.34398435 | 6.95E-19 |
| KDM1A | 0.27514982 | 0.21452407 | 0.34191397 | 7.88E-19 |
| HSD17B2 | 0.27496456 | 0.2150539 | 0.33100815 | 8.33E-19 |
| OPN3 | 0.27467926 | 0.22103602 | 0.32329444 | 9.08E-19 |
| NDUFAF4 | 0.27452271 | 0.21582689 | 0.33061687 | 9.51E-19 |
| UBA52 | 0.27449712 | 0.20638555 | 0.32950393 | 9.58E-19 |
| HECA | 0.27426304 | 0.21034447 | 0.3324765 | 1.03E-18 |
| LOC157381 | 0.27417184 | 0.21320261 | 0.32318217 | 1.06E-18 |
| ATP6V1C2 | 0.27376377 | 0.22853732 | 0.33208151 | 1.19E-18 |
| FBXO32 | 0.27348105 | 0.22288508 | 0.32158216 | 1.30E-18 |
| TET3 | 0.27342445 | 0.21970124 | 0.32394074 | 1.32E-18 |
| APLF | 0.27324999 | 0.20423462 | 0.33147037 | 1.39E-18 |
| SOD2 | 0.27313314 | 0.20655165 | 0.32675903 | 1.44E-18 |
| CCT4 | 0.27293022 | 0.21679866 | 0.32635998 | 1.53E-18 |
| TCF7L2 | 0.27266888 | 0.22623135 | 0.31687955 | 1.65E-18 |
| MYC | 0.27260654 | 0.19826319 | 0.3295499 | 1.69E-18 |
| CEP55 | 0.27194524 | 0.21767487 | 0.32414352 | 2.05E-18 |
| PALM2 | 0.27183171 | 0.20761553 | 0.32458225 | 2.12E-18 |
| ACN9 | 0.2717531 | 0.21559878 | 0.32390737 | 2.17E-18 |
| RPIA | 0.27163142 | 0.20730325 | 0.33016639 | 2.25E-18 |
| ZW10 | 0.27116698 | 0.2084013 | 0.32983247 | 2.58E-18 |
| RCC2 | 0.27106214 | 0.2074838 | 0.32531187 | 2.66E-18 |

| | | | | |
|---|---|---|---|---|
| ST8SIA1 | 0.27085822 | 0.21643065 | 0.32334079 | 2.83E-18 |
| NPM3 | 0.27084181 | 0.21299317 | 0.32249519 | 2.84E-18 |
| FUT3 | 0.27055533 | 0.21669874 | 0.31472112 | 3.09E-18 |
| GNB2L1 | 0.27051709 | 0.21042393 | 0.3241742 | 3.13E-18 |
| RNF19A | 0.27050189 | 0.20372725 | 0.33125803 | 3.14E-18 |
| DCPS | 0.27046665 | 0.20979082 | 0.33134732 | 3.18E-18 |
| PPPDE1 | 0.27005142 | 0.21216348 | 0.32180322 | 3.59E-18 |
| RPS23 | 0.26963753 | 0.20566082 | 0.32588012 | 4.06E-18 |
| METTL8 | 0.26928618 | 0.21633192 | 0.32808038 | 4.50E-18 |
| RNF145 | 0.26889937 | 0.20617623 | 0.32774627 | 5.04E-18 |
| NCK2 | 0.26881498 | 0.20036383 | 0.3395607 | 5.16E-18 |
| RPL8 | 0.2686439 | 0.198929 | 0.32384773 | 5.43E-18 |
| HDAC2 | 0.26833897 | 0.22406671 | 0.32879144 | 5.93E-18 |
| UGT8 | 0.26814513 | 0.19559871 | 0.32087776 | 6.28E-18 |
| DNAJC2 | 0.26795243 | 0.19824953 | 0.32083307 | 6.64E-18 |
| TBX19 | 0.26785214 | 0.21154945 | 0.31554653 | 6.84E-18 |
| GAL | 0.26777069 | 0.20810602 | 0.34165734 | 7.00E-18 |
| CHRAC1 | 0.26758303 | 0.21146001 | 0.34673849 | 7.40E-18 |
| PPP3R1 | 0.26747633 | 0.21107068 | 0.33090945 | 7.63E-18 |
| C9orf85 | 0.26731751 | 0.21083325 | 0.32645931 | 7.99E-18 |
| EIF2C2 (AGO2) | 0.26730991 | 0.19388476 | 0.33611442 | 8.01E-18 |
| LRRC8D | 0.2669371 | 0.21802239 | 0.31316088 | 8.93E-18 |
| TRDMT1 | 0.26648046 | 0.19676438 | 0.31088924 | 1.02E-17 |
| RGS10 | 0.26640713 | 0.20404454 | 0.32517682 | 1.04E-17 |
| TTK | 0.2663735 | 0.20486832 | 0.32560409 | 1.05E-17 |

| | | | | |
|---|---|---|---|---|
| MRPL3 | 0.26614968 | 0.2069067 | 0.32777183 | 1.12E-17 |
| IFRD1 | 0.26604236 | 0.21628054 | 0.3235229 | 1.16E-17 |
| ABLIM1 | 0.26597931 | 0.20421414 | 0.32294358 | 1.18E-17 |
| RPL34 | 0.26551597 | 0.20554761 | 0.32311302 | 1.35E-17 |
| CDC123 | 0.26517912 | 0.21294567 | 0.31619562 | 1.48E-17 |
| GLYATL2 | 0.26486448 | 0.20612873 | 0.3056771 | 1.63E-17 |
| MED17 | 0.26474844 | 0.20681611 | 0.34056428 | 1.68E-17 |
| GLRX3 | 0.26474779 | 0.207828 | 0.30716021 | 1.68E-17 |
| AIF1L | 0.26466755 | 0.21258096 | 0.31537143 | 1.72E-17 |
| DPH2 | 0.2646252 | 0.19550556 | 0.31712897 | 1.74E-17 |
| FUT4 | 0.26458611 | 0.193979 | 0.31225383 | 1.76E-17 |
| JOSD1 | 0.26449541 | 0.21231505 | 0.31070046 | 1.81E-17 |
| DUSP7 | 0.26431853 | 0.21784011 | 0.31536547 | 1.90E-17 |
| TKT | 0.2642611 | 0.20250136 | 0.3246674 | 1.93E-17 |
| GTF3C2 | 0.26415748 | 0.20326053 | 0.30980779 | 1.99E-17 |
| THOC5 | 0.2639059 | 0.19812936 | 0.30834631 | 2.14E-17 |
| PRKRIR | 0.26379936 | 0.20066791 | 0.33175044 | 2.21E-17 |
| HPDL | 0.26366826 | 0.20295944 | 0.30704302 | 2.29E-17 |
| SLCO5A1 | 0.26358915 | 0.19774571 | 0.31587175 | 2.34E-17 |
| CMPK1 | 0.26355143 | 0.20649568 | 0.31482934 | 2.37E-17 |
| NCOA7 | 0.2635401 | 0.20479324 | 0.32183078 | 2.38E-17 |
| FMO6P | 0.26293406 | 0.20284752 | 0.31035922 | 2.83E-17 |
| GART | 0.26282104 | 0.20195207 | 0.32226924 | 2.92E-17 |
| LPO | 0.26275273 | 0.21547961 | 0.31263047 | 2.98E-17 |
| RRS1 | 0.2626142 | 0.20206341 | 0.32209493 | 3.10E-17 |

| | | | | |
|---|---|---|---|---|
| ATP5L | 0.26260787 | 0.19507234 | 0.30749338 | 3.10E-17 |
| USP28 | 0.26236868 | 0.20801205 | 0.32749189 | 3.32E-17 |
| FAM171A1 | 0.26235982 | 0.18865289 | 0.33259022 | 3.33E-17 |
| RPL26 | 0.26234493 | 0.19560435 | 0.31536479 | 3.34E-17 |
| PDK1 | 0.2623149 | 0.20947867 | 0.32053836 | 3.37E-17 |
| PCGF6 | 0.26227769 | 0.19934998 | 0.32670918 | 3.41E-17 |
| TOMM5 | 0.26211601 | 0.21235991 | 0.3253474 | 3.57E-17 |
| GSDMC | 0.2619751 | 0.200217 | 0.31314367 | 3.72E-17 |
| ACE2 | 0.26164651 | 0.19374567 | 0.30930067 | 4.08E-17 |
| GTPBP4 | 0.26160136 | 0.20492703 | 0.33052704 | 4.13E-17 |
| RPL36 | 0.26150314 | 0.20415188 | 0.32768752 | 4.25E-17 |
| MTIF2 | 0.26127386 | 0.2083262 | 0.31116211 | 4.53E-17 |
| GPN1 | 0.26115589 | 0.19977906 | 0.3269179 | 4.69E-17 |
| DTD1 | 0.26071403 | 0.20131385 | 0.32729936 | 5.31E-17 |
| CREB3L2 | 0.26054032 | 0.19714398 | 0.30910977 | 5.58E-17 |
| GTPBP8 | 0.26044982 | 0.20295534 | 0.31663703 | 5.73E-17 |
| FMNL2 | 0.26043487 | 0.20304053 | 0.31688672 | 5.75E-17 |
| PSMG2 | 0.26031685 | 0.19625956 | 0.31746989 | 5.95E-17 |
| RPLP2 | 0.26029989 | 0.20174729 | 0.31942834 | 5.97E-17 |
| PHF5A | 0.26025984 | 0.2100591 | 0.32923729 | 6.04E-17 |
| DARS | 0.26015154 | 0.20491664 | 0.31317674 | 6.23E-17 |
| SACS | 0.2599332 | 0.20879971 | 0.3224849 | 6.63E-17 |
| RDX | 0.2598422 | 0.19684699 | 0.32285778 | 6.80E-17 |
| MTPAP | 0.25979926 | 0.19971577 | 0.30811126 | 6.88E-17 |
| ACSL5 | 0.25973402 | 0.19736502 | 0.3213596 | 7.01E-17 |

| | | | | |
|---|---|---|---|---|
| MTFR1 | 0.25966616 | 0.19968585 | 0.33638488 | 7.14E-17 |
| LYAR | 0.25957889 | 0.20463715 | 0.31621082 | 7.32E-17 |
| MPHOSPH10 | 0.25956423 | 0.20242648 | 0.32104273 | 7.35E-17 |
| PDCD2 | 0.25948087 | 0.19259956 | 0.31148605 | 7.53E-17 |
| POLR2D | 0.25885776 | 0.20435344 | 0.30477742 | 8.97E-17 |
| C6orf115 | 0.25885748 | 0.22000749 | 0.31443267 | 8.97E-17 |
| PLCH1 | 0.25808666 | 0.1987616 | 0.31186604 | 1.11E-16 |
| C16orf57 | 0.25800881 | 0.21489224 | 0.32612763 | 1.14E-16 |
| SLC25A37 | 0.25798647 | 0.1915139 | 0.31544775 | 1.14E-16 |
| SEPHS1 | 0.25790631 | 0.19993842 | 0.30079357 | 1.17E-16 |
| MTHFD1L | 0.2578794 | 0.20038868 | 0.30161525 | 1.18E-16 |
| EIF2A | 0.25786699 | 0.20191133 | 0.3009782 | 1.18E-16 |
| SHMT2 | 0.25738496 | 0.19735689 | 0.30831315 | 1.35E-16 |
| STAC | 0.25730471 | 0.20894997 | 0.31471926 | 1.38E-16 |
| SKA3 | 0.25719494 | 0.18567112 | 0.31263819 | 1.43E-16 |
| RASGEF1B | 0.25702139 | 0.19946443 | 0.31300341 | 1.50E-16 |
| PTMA | 0.25682488 | 0.1958133 | 0.31612227 | 1.58E-16 |
| SLC7A1 | 0.25681199 | 0.18782812 | 0.30521909 | 1.59E-16 |
| ENY2 | 0.25680843 | 0.18593759 | 0.31804179 | 1.59E-16 |
| DTNB | 0.25679972 | 0.20010941 | 0.31439101 | 1.59E-16 |
| SEC63 | 0.25661841 | 0.18921558 | 0.30281455 | 1.68E-16 |
| BCL11B | 0.25655004 | 0.19306119 | 0.30032626 | 1.71E-16 |
| EIF3F | 0.25649727 | 0.20309485 | 0.31487528 | 1.73E-16 |
| FAM188B | 0.25647218 | 0.19292085 | 0.30105502 | 1.75E-16 |
| IFNAR2 | 0.25637598 | 0.20834373 | 0.31708059 | 1.79E-16 |

| | | | | |
|---|---|---|---|---|
| ELF5 | 0.25631132 | 0.1838155 | 0.31161797 | 1.82E-16 |
| TEX261 | 0.25623047 | 0.2003807 | 0.30563458 | 1.87E-16 |
| MRRF | 0.25596695 | 0.18573732 | 0.32087511 | 2.01E-16 |
| NOB1 | 0.25596456 | 0.20914517 | 0.30860925 | 2.01E-16 |
| RPL13 | 0.25583173 | 0.19945809 | 0.31148245 | 2.08E-16 |
| MAML2 | 0.25579752 | 0.19657272 | 0.31228434 | 2.10E-16 |
| GTF2F2 | 0.25568412 | 0.20106343 | 0.29711502 | 2.17E-16 |
| C12orf45 | 0.2556642 | 0.20348606 | 0.30449995 | 2.18E-16 |
| HEBP2 | 0.2554398 | 0.2006817 | 0.31183636 | 2.32E-16 |
| ZNF883 | 0.25520888 | 0.1858676 | 0.31547807 | 2.48E-16 |
| POLR1D | 0.255117 | 0.20273926 | 0.3105891 | 2.54E-16 |
| SMNDC1 | 0.25493893 | 0.20274639 | 0.31139787 | 2.67E-16 |
| ESYT3 | 0.25482051 | 0.18808411 | 0.3087857 | 2.76E-16 |
| CCDC82 | 0.25481791 | 0.19768895 | 0.30579821 | 2.76E-16 |
| SSR2 | 0.25456976 | 0.18668457 | 0.30389167 | 2.95E-16 |
| RPUSD4 | 0.25436646 | 0.20874599 | 0.30564231 | 3.12E-16 |
| LOC150622 | 0.25417696 | 0.19031176 | 0.30375081 | 3.29E-16 |
| ZNF462 | 0.25401306 | 0.1929816 | 0.31551249 | 3.44E-16 |
| MCTP2 | 0.25395277 | 0.1902571 | 0.31329853 | 3.50E-16 |
| DDN | 0.25374648 | 0.18936174 | 0.31186971 | 3.70E-16 |
| C3orf17 | 0.25324691 | 0.1924794 | 0.30920101 | 4.25E-16 |
| PIWIL4 | 0.25294128 | 0.19870051 | 0.30611813 | 4.62E-16 |
| C3orf26 | 0.25288834 | 0.18224304 | 0.31141989 | 4.69E-16 |
| VGLL1 | 0.25279017 | 0.19755286 | 0.30753012 | 4.81E-16 |
| CXCL1 | 0.25276221 | 0.19112287 | 0.30834942 | 4.85E-16 |

| | | | | |
|---|---|---|---|---|
| RRAS2 | 0.25272678 | 0.18998547 | 0.29807472 | 4.90E-16 |
| GARS | 0.25268906 | 0.20185247 | 0.31897161 | 4.95E-16 |
| ODF2L | 0.25235994 | 0.20165742 | 0.30928568 | 5.41E-16 |
| DSCC1 | 0.25228666 | 0.18667707 | 0.31439421 | 5.52E-16 |
| MCART6 | 0.25211987 | 0.19824705 | 0.32274016 | 5.78E-16 |
| ASS1 | 0.25197129 | 0.18635287 | 0.30334153 | 6.02E-16 |
| RGMA | 0.25190089 | 0.1945923 | 0.30266693 | 6.14E-16 |
| ACTR3B | 0.25188377 | 0.17757961 | 0.31566067 | 6.16E-16 |
| DPH5 | 0.25137537 | 0.19440243 | 0.30203884 | 7.08E-16 |
| LRPPRC | 0.2510087 | 0.18911472 | 0.31151778 | 7.82E-16 |
| PPP1R14C | 0.2509324 | 0.18793466 | 0.3122417 | 7.98E-16 |
| SH2D2A | 0.25059475 | 0.19708869 | 0.31391671 | 8.75E-16 |
| SNHG3 | 0.25050587 | 0.1899867 | 0.30256773 | 8.96E-16 |
| METAP1 | 0.25048928 | 0.19019063 | 0.31847066 | 9.00E-16 |
| MARS2 | 0.2504221 | 0.18494693 | 0.31984192 | 9.17E-16 |
| C1orf174 | 0.25037963 | 0.20602492 | 0.2956627 | 9.27E-16 |
| WASF1 | 0.25012207 | 0.19103207 | 0.31116272 | 9.94E-16 |
| TNFRSF21 | 0.24999783 | 0.20526875 | 0.30482027 | 1.03E-15 |
| NOC3L | 0.2499571 | 0.19276383 | 0.30596442 | 1.04E-15 |
| PHGDH | 0.24981645 | 0.17634147 | 0.30737889 | 1.08E-15 |
| PREP | 0.24937607 | 0.19733115 | 0.30419434 | 1.22E-15 |
| DSC2 | 0.24917206 | 0.19489963 | 0.31174616 | 1.28E-15 |
| PDCD5 | 0.24913034 | 0.17169483 | 0.31730474 | 1.30E-15 |
| DKK1 | 0.24906114 | 0.18845045 | 0.30794737 | 1.32E-15 |
| CNGA1 | 0.24860404 | 0.19073266 | 0.31284243 | 1.50E-15 |

| | | | | |
|---|---|---|---|---|
| KCNG1 | 0.24859866 | 0.20009271 | 0.29596425 | 1.50E-15 |
| GAPDH | 0.24839612 | 0.19064413 | 0.30218039 | 1.58E-15 |
| LSM2 | 0.24764035 | 0.17683245 | 0.30319159 | 1.94E-15 |
| REPS1 | 0.24762539 | 0.17983536 | 0.30569315 | 1.94E-15 |
| B3GALNT2 | 0.24724513 | 0.18030072 | 0.30377849 | 2.15E-15 |
| LIPG | 0.24716243 | 0.19670699 | 0.29200777 | 2.20E-15 |
| DDX26B | 0.24698586 | 0.18900605 | 0.30633503 | 2.31E-15 |
| RCC1 | 0.24663717 | 0.18582246 | 0.30275959 | 2.53E-15 |
| TSPAN14 | 0.24641023 | 0.19993834 | 0.30616492 | 2.69E-15 |
| CALCB | 0.24625309 | 0.19014935 | 0.30813259 | 2.80E-15 |
| TXNDC5 | 0.24593778 | 0.19912997 | 0.30190941 | 3.05E-15 |
| DNMT3B | 0.24550299 | 0.19365932 | 0.31721434 | 3.42E-15 |
| UCHL3 | 0.24531331 | 0.20139932 | 0.30690308 | 3.59E-15 |
| BID | 0.2453021 | 0.19117985 | 0.30460903 | 3.61E-15 |
| SLC15A2 | 0.24526733 | 0.18335073 | 0.30613218 | 3.64E-15 |
| NDC80 | 0.24524739 | 0.19295224 | 0.31952212 | 3.66E-15 |
| C1orf116 | 0.24519494 | 0.19435424 | 0.30444387 | 3.71E-15 |
| COL4A4 | 0.24502912 | 0.18660495 | 0.29862392 | 3.87E-15 |
| RPL29 | 0.2449805 | 0.1819445 | 0.29782483 | 3.92E-15 |
| CCL20 | 0.24490138 | 0.18261421 | 0.28571083 | 4.01E-15 |
| RBM28 | 0.2448658 | 0.17548523 | 0.2983606 | 4.05E-15 |
| DNTTIP2 | 0.24475947 | 0.18355871 | 0.30704297 | 4.16E-15 |
| PPP1CB | 0.24471534 | 0.19002547 | 0.29004276 | 4.21E-15 |
| CHEK1 | 0.2444695 | 0.17980653 | 0.31703623 | 4.49E-15 |
| MMACHC | 0.24443587 | 0.19202903 | 0.30868612 | 4.53E-15 |

| | | | | |
|---|---|---|---|---|
| RPL23A | 0.2443061 | 0.18308209 | 0.32962177 | 4.69E-15 |
| NFYA | 0.24414731 | 0.18562974 | 0.30119947 | 4.89E-15 |
| POLR1B | 0.24405511 | 0.19207233 | 0.32095184 | 5.01E-15 |
| ACTL8 | 0.24398858 | 0.18565177 | 0.30845652 | 5.10E-15 |
| SF3B14 | 0.24391859 | 0.19891704 | 0.30498383 | 5.19E-15 |
| GLB1L2 | 0.2439102 | 0.18313758 | 0.29896893 | 5.20E-15 |
| PKP1 | 0.24385687 | 0.18121943 | 0.29567979 | 5.28E-15 |
| TMEM71 | 0.24377502 | 0.17908734 | 0.30442747 | 5.39E-15 |
| ZNF485 | 0.24305363 | 0.18738082 | 0.29295729 | 6.51E-15 |
| KPNA3 | 0.24303093 | 0.17420761 | 0.30431361 | 6.55E-15 |
| PABPC1P2 | 0.24299133 | 0.1907631 | 0.30511124 | 6.62E-15 |
| XPO5 | 0.24287835 | 0.17982362 | 0.28528641 | 6.82E-15 |

**Table S19. KEGG pathway analysis of genes co-expressed with PABPC1.**

Results of a KEGG pathway enrichment analysis using Genecois3 (http://www.genecodis.cnb.csic.es) for genes co-expressed with PABPC1 in the clinical samples (top 500 shown in Table S19). Only significantly (adjusted p<0.05) enriched pathway are show.

| Pathway Details | Support | Reference Support | Hypergeometric Test p-value (FDR adjusted) |
|---|---|---|---|
| Neurotrophin signaling pathway | 15 | 124 | 0.0399482 |
| Arachidonic acid metabolism,VEGF signaling pathway | 4 | 16 | 0.0416621 |
| RNA transport | 45 | 145 | 1.36E-16 |
| RNA transport,Ribosome biogenesis in eukaryotes | 6 | 18 | 0.00433798 |
| RNA transport,mRNA surveillance pathway | 7 | 30 | 0.0115692 |
| Glycerophospholipid metabolism,Fc gamma R-mediated phagocytosis,Ether lipid metabolism,Glutamatergic synapse,GnRH signaling pathway | 3 | 5 | 0.0108321 |
| Glycerophospholipid metabolism,Ether lipid metabolism,Glutamatergic synapse,GnRH signaling pathway | 4 | 17 | 0.0468142 |
| Glycolysis / Gluconeogenesis | 10 | 62 | 0.021515 |
| Cysteine and methionine metabolism | 9 | 36 | 0.00288736 |
| Pyruvate metabolism | 8 | 39 | 0.0127847 |
| Pyruvate metabolism,Citrate cycle (TCA cycle) | 4 | 10 | 0.0113403 |
| Insulin signaling pathway,Regulation of actin cytoskeleton,Focal adhesion | 5 | 21 | 0.028981 |
| Cytokine-cytokine receptor interaction | 50 | 259 | 6.71E-10 |
| Cytokine-cytokine receptor interaction,Jak-STAT signaling pathway | 13 | 95 | 0.0262434 |
| Cytokine-cytokine receptor interaction,Jak-STAT signaling pathway,Toxoplasmosis,Measles,Natural killer cell mediated cytotoxicity,Tuberculosis,Leishmaniasis,Chagas disease (American trypanosomiasis),Osteoclast differentiation | 3 | 3 | 0.00219324 |

| | | | |
|---|---|---|---|
| Cytokine-cytokine receptor interaction,Jak-STAT signaling pathway,Toxoplasmosis,Measles,Tuberculosis,African trypanosomiasis,Type I diabetes mellitus,Leishmaniasis,Chagas disease (American trypanosomiasis),Amoebiasis,Allograft rejection | 3 | 3 | 0.00219324 |
| Cytokine-cytokine receptor interaction,Jak-STAT signaling pathway,Toxoplasmosis,Measles,Tuberculosis,Leishmaniasis,Chagas disease (American trypanosomiasis) | 5 | 5 | 4.23E-05 |
| Cytokine-cytokine receptor interaction,Jak-STAT signaling pathway,Measles | 9 | 28 | 0.00080582 |
| Cytokine-cytokine receptor interaction,Jak-STAT signaling pathway,Measles,Natural killer cell mediated cytotoxicity,Osteoclast differentiation | 4 | 6 | 0.00218312 |
| Cytokine-cytokine receptor interaction,Jak-STAT signaling pathway,Measles,Endocytosis | 3 | 3 | 0.00219324 |
| Cytokine-cytokine receptor interaction,NOD-like receptor signaling pathway | 5 | 8 | 0.0009351 |
| Cytokine-cytokine receptor interaction,NOD-like receptor signaling pathway,Chemokine signaling pathway | 4 | 5 | 0.00126446 |
| Cytokine-cytokine receptor interaction,NOD-like receptor signaling pathway,Chemokine signaling pathway,Rheumatoid arthritis,Epithelial cell signaling in Helicobacter pylori infection | 3 | 3 | 0.00219324 |
| Cytokine-cytokine receptor interaction,NOD-like receptor signaling pathway,Rheumatoid arthritis | 4 | 7 | 0.00338038 |
| Cytokine-cytokine receptor interaction,Toxoplasmosis | 6 | 15 | 0.00231127 |
| Cytokine-cytokine receptor interaction,Toxoplasmosis,Malaria,Allograft rejection | 3 | 5 | 0.0108321 |
| Cytokine-cytokine receptor interaction,Toxoplasmosis,Allograft rejection | 4 | 6 | 0.00218312 |
| Cytokine-cytokine receptor interaction,Pathways in cancer,Epithelial cell signaling in Helicobacter pylori infection | 3 | 3 | 0.00219324 |
| Cytokine-cytokine receptor interaction,Chemokine signaling pathway | 19 | 62 | 6.76E-07 |
| Cytokine-cytokine receptor interaction,Chemokine signaling pathway,Toll-like receptor signaling pathway | 4 | 7 | 0.00338038 |
| Cytokine-cytokine receptor interaction,Chemokine signaling pathway,Rheumatoid arthritis | 6 | 10 | 0.00033366 |

| | | | |
|---|---|---|---|
| Cytokine-cytokine receptor interaction,Toll-like receptor signaling pathway | 7 | 28 | 0.00835914 |
| Cytokine-cytokine receptor interaction,Toll-like receptor signaling pathway,RIG-I-like receptor signaling pathway,Chagas disease (American trypanosomiasis),Amoebiasis | 3 | 3 | 0.00219324 |
| Cytokine-cytokine receptor interaction,Toll-like receptor signaling pathway,Chagas disease (American trypanosomiasis) | 4 | 8 | 0.00561468 |
| Cytokine-cytokine receptor interaction,Measles | 10 | 37 | 0.00134043 |
| Cytokine-cytokine receptor interaction,Measles,Natural killer cell mediated cytotoxicity | 5 | 26 | 0.0499864 |
| Cytokine-cytokine receptor interaction,Measles,Natural killer cell mediated cytotoxicity,Chagas disease (American trypanosomiasis) | 4 | 6 | 0.00218312 |
| Cytokine-cytokine receptor interaction,Measles,African trypanosomiasis,Type I diabetes mellitus,Chagas disease (American trypanosomiasis),Allograft rejection | 4 | 5 | 0.00126446 |
| Cytokine-cytokine receptor interaction,Measles,Chagas disease (American trypanosomiasis) | 6 | 11 | 0.00051547 |
| Cytokine-cytokine receptor interaction,Tuberculosis | 6 | 32 | 0.0406529 |
| Cytokine-cytokine receptor interaction,Tuberculosis,African trypanosomiasis | 4 | 7 | 0.00338038 |
| Cytokine-cytokine receptor interaction,Tuberculosis,African trypanosomiasis,Malaria | 3 | 6 | 0.0168365 |
| Cytokine-cytokine receptor interaction,African trypanosomiasis | 5 | 9 | 0.00157116 |
| Cytokine-cytokine receptor interaction,Malaria | 6 | 16 | 0.00262021 |
| Cytokine-cytokine receptor interaction,Malaria,Rheumatoid arthritis | 3 | 9 | 0.0418808 |
| Cytokine-cytokine receptor interaction,Malaria,Chagas disease (American trypanosomiasis),Amoebiasis | 3 | 9 | 0.0418808 |
| Cytokine-cytokine receptor interaction,Endocytosis | 7 | 20 | 0.00198878 |
| Cytokine-cytokine receptor interaction,Rheumatoid arthritis | 9 | 32 | 0.00174548 |
| Cytokine-cytokine receptor interaction,Rheumatoid arthritis,Amoebiasis | 3 | 9 | 0.0418808 |
| Cytokine-cytokine receptor interaction,Epithelial cell signaling in Helicobacter pylori infection | 5 | 7 | 0.00046345 |

| | | | |
|---|---|---|---|
| Cytokine-cytokine receptor interaction,Chagas disease (American trypanosomiasis) | 8 | 23 | 0.00101306 |
| Cytokine-cytokine receptor interaction,Chagas disease (American trypanosomiasis),Amoebiasis | 4 | 10 | 0.0113403 |
| Cytokine-cytokine receptor interaction,Osteoclast differentiation | 5 | 19 | 0.0208959 |
| Cytokine-cytokine receptor interaction,Amoebiasis | 6 | 14 | 0.00177582 |
| Cytokine-cytokine receptor interaction,Allograft rejection | 5 | 11 | 0.0028532 |
| Cytokine-cytokine receptor interaction,Primary immunodeficiency | 4 | 6 | 0.00218312 |
| Jak-STAT signaling pathway | 21 | 153 | 0.00586095 |
| Jak-STAT signaling pathway,Toxoplasmosis,Toll-like receptor signaling pathway,Measles,Chagas disease (American trypanosomiasis),Amoebiasis | 3 | 10 | 0.0491861 |
| Jak-STAT signaling pathway,Toxoplasmosis,Measles,Natural killer cell mediated cytotoxicity,Chagas disease (American trypanosomiasis),Osteoclast differentiation | 4 | 11 | 0.0153891 |
| Jak-STAT signaling pathway,Toxoplasmosis,Measles,Chagas disease (American trypanosomiasis) | 6 | 16 | 0.00262021 |
| Jak-STAT signaling pathway,Toxoplasmosis,Measles,Chagas disease (American trypanosomiasis),Amoebiasis | 4 | 11 | 0.0153891 |
| Jak-STAT signaling pathway,Measles | 11 | 53 | 0.00362581 |
| Jak-STAT signaling pathway,Measles,Natural killer cell mediated cytotoxicity,Osteoclast differentiation | 5 | 14 | 0.00714564 |
| Jak-STAT signaling pathway,Natural killer cell mediated cytotoxicity | 6 | 33 | 0.0426965 |
| Jak-STAT signaling pathway,Endocytosis | 4 | 7 | 0.00338038 |
| Jak-STAT signaling pathway,Primary immunodeficiency | 3 | 3 | 0.00219324 |
| Protein digestion and absorption,Salivary secretion | 3 | 10 | 0.0491861 |
| Purine metabolism | 29 | 158 | 2.14E-05 |
| Purine metabolism,Pyrimidine metabolism | 14 | 68 | 0.00150026 |
| Purine metabolism,Pyrimidine metabolism,Cytosolic DNA-sensing pathway,RNA polymerase | 5 | 17 | 0.0145043 |
| Purine metabolism,Pyrimidine metabolism,RNA polymerase | 8 | 28 | 0.00244242 |

| | | | |
|---|---|---|---|
| Pyrimidine metabolism | 20 | 93 | 5.86E-05 |
| NOD-like receptor signaling pathway | 12 | 57 | 0.0022792 |
| Alzheimer's disease | 19 | 163 | 0.0294693 |
| Alzheimer's disease,Pathways in cancer,p53 signaling pathway,Apoptosis | 3 | 6 | 0.0168365 |
| Alzheimer's disease,Natural killer cell mediated cytotoxicity,Apoptosis | 3 | 10 | 0.0491861 |
| Parkinson's disease,Huntington's disease | 13 | 99 | 0.0330598 |
| Parkinson's disease,Huntington's disease,Calcium signaling pathway | 3 | 8 | 0.0330655 |
| Huntington's disease | 24 | 180 | 0.00431182 |
| Huntington's disease,Calcium signaling pathway | 4 | 17 | 0.0468142 |
| Axon guidance | 20 | 128 | 0.00237918 |
| Axon guidance,Pathogenic Escherichia coli infection,T cell receptor signaling pathway | 3 | 5 | 0.0108321 |
| Fc gamma R-mediated phagocytosis | 20 | 92 | 5.10E-05 |
| Fc gamma R-mediated phagocytosis,Regulation of actin cytoskeleton | 11 | 41 | 0.00080107 |
| Fc gamma R-mediated phagocytosis,Regulation of actin cytoskeleton,Pathogenic Escherichia coli infection,Bacterial invasion of epithelial cells,Shigellosis | 4 | 10 | 0.0113403 |
| Fc gamma R-mediated phagocytosis,Regulation of actin cytoskeleton,Bacterial invasion of epithelial cells | 8 | 23 | 0.00101306 |
| Fc gamma R-mediated phagocytosis,Regulation of actin cytoskeleton,Bacterial invasion of epithelial cells,Shigellosis | 7 | 15 | 0.00046167 |
| Fc gamma R-mediated phagocytosis,MAPK signaling pathway,Vascular smooth muscle contraction,VEGF signaling pathway,Fc epsilon RI signaling pathway,Glutamatergic synapse,Long-term depression,GnRH signaling pathway,Pancreatic secretion | 3 | 5 | 0.0108321 |
| Fc gamma R-mediated phagocytosis,VEGF signaling pathway,Fc epsilon RI signaling pathway | 5 | 24 | 0.0413546 |
| Fc gamma R-mediated phagocytosis,VEGF signaling pathway,Fc epsilon RI signaling pathway,Pathways in cancer,Non-small cell lung cancer,Phosphatidylinositol signaling system,Natural killer cell mediated | 3 | 10 | 0.0491861 |

| | | | |
|---|---|---|---|
| cytotoxicity,B cell receptor signaling pathway,Leukocyte transendothelial migration,ErbB signaling pathway,Glioma | | | |
| Fc gamma R-mediated phagocytosis,Fc epsilon RI signaling pathway | 7 | 34 | 0.0194356 |
| Fc gamma R-mediated phagocytosis,Fc epsilon RI signaling pathway,Long-term depression | 4 | 10 | 0.0113403 |
| Fc gamma R-mediated phagocytosis,Fc epsilon RI signaling pathway,B cell receptor signaling pathway | 5 | 25 | 0.0472131 |
| Fc gamma R-mediated phagocytosis,Glutamatergic synapse,GnRH signaling pathway | 4 | 9 | 0.00841625 |
| Fc gamma R-mediated phagocytosis,Pathways in cancer | 5 | 26 | 0.0499864 |
| Fc gamma R-mediated phagocytosis,Inositol phosphate metabolism,Phosphatidylinositol signaling system | 3 | 9 | 0.0418808 |
| Fc gamma R-mediated phagocytosis,Phosphatidylinositol signaling system | 4 | 17 | 0.0468142 |
| Fc gamma R-mediated phagocytosis,Adherens junction | 3 | 10 | 0.0491861 |
| Regulation of actin cytoskeleton | 30 | 209 | 0.00077715 |
| Regulation of actin cytoskeleton,Pathogenic Escherichia coli infection,Bacterial invasion of epithelial cells,Shigellosis | 5 | 13 | 0.00572625 |
| Regulation of actin cytoskeleton,Adherens junction,Bacterial invasion of epithelial cells,Shigellosis | 3 | 9 | 0.0418808 |
| Regulation of actin cytoskeleton,Bacterial invasion of epithelial cells | 9 | 34 | 0.00219052 |
| Regulation of actin cytoskeleton,Bacterial invasion of epithelial cells,Shigellosis | 8 | 21 | 0.00057957 |
| Regulation of actin cytoskeleton,Shigellosis | 10 | 31 | 0.00043272 |
| Basal transcription factors | 10 | 39 | 0.00174225 |
| MAPK signaling pathway | 33 | 262 | 0.00234899 |
| MAPK signaling pathway,Vascular smooth muscle contraction,VEGF signaling pathway,Fc epsilon RI signaling pathway,Glutamatergic synapse,Long-term depression,GnRH signaling pathway,Pancreatic secretion | 4 | 17 | 0.0468142 |
| MAPK signaling pathway,Vascular smooth muscle contraction,Glutamatergic synapse,GnRH signaling pathway | 5 | 25 | 0.0472131 |

| | | | |
|---|---|---|---|
| MAPK signaling pathway,GnRH signaling pathway | 9 | 57 | 0.0310636 |
| MAPK signaling pathway,GnRH signaling pathway,Calcium signaling pathway,Gap junction | 3 | 7 | 0.024676 |
| MAPK signaling pathway,GnRH signaling pathway,Focal adhesion,ErbB signaling pathway | 4 | 16 | 0.0416621 |
| MAPK signaling pathway,GnRH signaling pathway,Focal adhesion,ErbB signaling pathway,Endometrial cancer | 3 | 10 | 0.0491861 |
| MAPK signaling pathway,Pancreatic secretion | 5 | 21 | 0.028981 |
| MAPK signaling pathway,Focal adhesion | 7 | 40 | 0.0368106 |
| MAPK signaling pathway,Focal adhesion,ErbB signaling pathway | 5 | 26 | 0.0499864 |
| MAPK signaling pathway,Long-term potentiation | 6 | 31 | 0.0367723 |
| MAPK signaling pathway,ErbB signaling pathway | 6 | 32 | 0.0406529 |
| Vascular smooth muscle contraction,Glutamatergic synapse,Long-term depression,GnRH signaling pathway,Pancreatic secretion | 5 | 26 | 0.0499864 |
| Vascular smooth muscle contraction,Glutamatergic synapse,GnRH signaling pathway,Chemokine signaling pathway,Calcium signaling pathway,Endocrine and other factor-regulated calcium reabsorption,Wnt signaling pathway,Salivary secretion,Long-term potentiation,Gap junction,Melanogenesis,Gastric acid secretion,Amoebiasis | 3 | 9 | 0.0418808 |
| VEGF signaling pathway,Calcium signaling pathway,Natural killer cell mediated cytotoxicity,B cell receptor signaling pathway | 3 | 9 | 0.0418808 |
| Fc epsilon RI signaling pathway | 11 | 77 | 0.0311774 |
| Fc epsilon RI signaling pathway,Long-term depression | 5 | 26 | 0.0499864 |
| Fc epsilon RI signaling pathway,Natural killer cell mediated cytotoxicity,Osteoclast differentiation | 4 | 17 | 0.0468142 |
| Fc epsilon RI signaling pathway,B cell receptor signaling pathway | 6 | 34 | 0.04732 |
| Glutamatergic synapse | 15 | 124 | 0.0399482 |
| Glutamatergic synapse,Long-term depression | 7 | 39 | 0.0330655 |
| Glutamatergic synapse,Long-term depression,Toxoplasmosis | 5 | 21 | 0.028981 |

| | | | |
|---|---|---|---|
| Glutamatergic synapse,Long-term depression,Chemokine signaling pathway,Leukocyte transendothelial migration,Gap junction,Melanogenesis,Gastric acid secretion,Tight junction | 3 | 4 | 0.00562131 |
| Glutamatergic synapse,Long-term depression,Chemokine signaling pathway,Gap junction,Melanogenesis,Gastric acid secretion | 4 | 8 | 0.00561468 |
| Glutamatergic synapse,Long-term depression,Chemokine signaling pathway,Gap junction,Melanogenesis,Gastric acid secretion,Chagas disease (American trypanosomiasis) | 3 | 7 | 0.024676 |
| Glutamatergic synapse,Chemokine signaling pathway | 7 | 42 | 0.0421101 |
| Glutamatergic synapse,Chemokine signaling pathway,Gap junction,Melanogenesis,Gastric acid secretion | 5 | 21 | 0.028981 |
| Long-term depression | 10 | 69 | 0.0365157 |
| Long-term depression,Chemokine signaling pathway | 5 | 17 | 0.0145043 |
| Long-term depression,Tight junction | 4 | 13 | 0.0243223 |
| Long-term depression,Tight junction,Chagas disease (American trypanosomiasis) | 3 | 7 | 0.024676 |
| Long-term depression,Chagas disease (American trypanosomiasis) | 4 | 17 | 0.0468142 |
| GnRH signaling pathway,Calcium signaling pathway,Wnt signaling pathway,Long-term potentiation,Melanogenesis,Gastric acid secretion | 4 | 14 | 0.0298996 |
| GnRH signaling pathway,Calcium signaling pathway,ErbB signaling pathway,Glioma | 3 | 7 | 0.024676 |
| Pancreatic secretion,Chemokine signaling pathway,Long-term potentiation | 3 | 9 | 0.0418808 |
| Pancreatic secretion,Endocrine and other factor-regulated calcium reabsorption,Salivary secretion,Aldosterone-regulated sodium reabsorption,Gastric acid secretion,Carbohydrate digestion and absorption | 3 | 9 | 0.0418808 |
| Toxoplasmosis | 18 | 123 | 0.00563489 |
| Toxoplasmosis,Measles | 7 | 37 | 0.0270366 |
| Toxoplasmosis,Measles,Leishmaniasis | 6 | 20 | 0.00683128 |
| Toxoplasmosis,Measles,Leishmaniasis,Osteoclast differentiation | 4 | 11 | 0.0153891 |
| Toxoplasmosis,Measles,Osteoclast differentiation | 5 | 24 | 0.0413546 |

| | | | |
|---|---|---|---|
| Toxoplasmosis,Chagas disease (American trypanosomiasis) | 8 | 48 | 0.0327381 |
| Citrate cycle (TCA cycle) | 8 | 30 | 0.00356975 |
| Citrate cycle (TCA cycle),Glyoxylate and dicarboxylate metabolism | 3 | 5 | 0.0108321 |
| Dorso-ventral axis formation | 7 | 23 | 0.00337763 |
| Dorso-ventral axis formation,Pathways in cancer | 3 | 9 | 0.0418808 |
| Pathways in cancer | 56 | 324 | 3.05E-09 |
| Pathways in cancer,Non-small cell lung cancer | 11 | 50 | 0.00243617 |
| Pathways in cancer,Non-small cell lung cancer,Calcium signaling pathway,ErbB signaling pathway,Glioma | 3 | 6 | 0.0168365 |
| Pathways in cancer,Non-small cell lung cancer,Leukocyte transendothelial migration | 4 | 14 | 0.0298996 |
| Pathways in cancer,Non-small cell lung cancer,Melanoma,Bladder cancer,Cell cycle,Small cell lung cancer,Pancreatic cancer,Chronic myeloid leukemia,Glioma | 3 | 7 | 0.024676 |
| Pathways in cancer,Non-small cell lung cancer,Melanoma,Bladder cancer,Pancreatic cancer,Glioma | 4 | 17 | 0.0468142 |
| Pathways in cancer,Non-small cell lung cancer,Melanoma,Cell cycle,Small cell lung cancer,Pancreatic cancer,Chronic myeloid leukemia,Glioma | 4 | 8 | 0.00561468 |
| Pathways in cancer,Non-small cell lung cancer,Melanoma,Small cell lung cancer,Pancreatic cancer,Chronic myeloid leukemia,Glioma | 5 | 19 | 0.0208959 |
| Pathways in cancer,Non-small cell lung cancer,Melanoma,Pancreatic cancer,Glioma | 6 | 29 | 0.0286494 |
| Pathways in cancer,Non-small cell lung cancer,Small cell lung cancer | 6 | 23 | 0.0118268 |
| Pathways in cancer,Non-small cell lung cancer,Chronic myeloid leukemia,Glioma | 6 | 33 | 0.0426965 |
| Pathways in cancer,Non-small cell lung cancer,Glioma | 9 | 41 | 0.00569375 |
| Pathways in cancer,Measles,p53 signaling pathway | 4 | 8 | 0.00561468 |
| Pathways in cancer,Measles,p53 signaling pathway,Cell cycle,Small cell lung cancer | 3 | 7 | 0.024676 |
| Pathways in cancer,Natural killer cell mediated cytotoxicity | 6 | 33 | 0.0426965 |

| | | | |
|---|---|---|---|
| Pathways in cancer,Arrhythmogenic right ventricular cardiomyopathy (ARVC),Wnt signaling pathway,Prostate cancer,Melanogenesis,Adherens junction,Colorectal cancer,Endometrial cancer,Basal cell carcinoma,Acute myeloid leukemia,Thyroid cancer | 3 | 4 | 0.00562131 |
| Pathways in cancer,Endocytosis | 7 | 28 | 0.00835914 |
| Pathways in cancer,Endocytosis,Adherens junction | 3 | 9 | 0.0418808 |
| Pathways in cancer,Wnt signaling pathway | 12 | 65 | 0.00547613 |
| Pathways in cancer,Wnt signaling pathway,Melanogenesis | 9 | 43 | 0.00782329 |
| Pathways in cancer,Wnt signaling pathway,Melanogenesis,Basal cell carcinoma | 8 | 38 | 0.0116878 |
| Pathways in cancer,Wnt signaling pathway,Adherens junction,Colorectal cancer | 4 | 12 | 0.0198264 |
| Pathways in cancer,Wnt signaling pathway,Colorectal cancer | 5 | 24 | 0.0413546 |
| Pathways in cancer,Wnt signaling pathway,Colorectal cancer,Endometrial cancer,Acute myeloid leukemia,Thyroid cancer | 4 | 6 | 0.00218312 |
| Pathways in cancer,Wnt signaling pathway,Acute myeloid leukemia | 5 | 7 | 0.00046345 |
| Pathways in cancer,p53 signaling pathway | 6 | 19 | 0.00580534 |
| Pathways in cancer,p53 signaling pathway,Small cell lung cancer | 4 | 10 | 0.0113403 |
| Pathways in cancer,Prostate cancer | 10 | 75 | 0.0494568 |
| Pathways in cancer,Prostate cancer,Adherens junction,Endometrial cancer | 4 | 9 | 0.00841625 |
| Pathways in cancer,Prostate cancer,Cell cycle,Small cell lung cancer | 3 | 10 | 0.0491861 |
| Pathways in cancer,Melanogenesis | 10 | 54 | 0.0106508 |
| Pathways in cancer,Melanogenesis,Acute myeloid leukemia | 4 | 13 | 0.0243223 |
| Pathways in cancer,Ubiquitin mediated proteolysis | 5 | 19 | 0.0208959 |
| Pathways in cancer,Ubiquitin mediated proteolysis,Small cell lung cancer | 3 | 9 | 0.0418808 |
| Pathways in cancer,Adherens junction | 6 | 28 | 0.0249821 |
| Pathways in cancer,Bladder cancer | 7 | 39 | 0.0330655 |

| | | | |
|---|---|---|---|
| Pathways in cancer,Bladder cancer,Cell cycle,Small cell lung cancer,Chronic myeloid leukemia | 4 | 8 | 0.00561468 |
| Pathways in cancer,Cell cycle | 9 | 32 | 0.00174548 |
| Pathways in cancer,Cell cycle,Small cell lung cancer | 7 | 15 | 0.00046167 |
| Pathways in cancer,Cell cycle,Small cell lung cancer,Chronic myeloid leukemia | 5 | 10 | 0.00235327 |
| Pathways in cancer,Cell cycle,Pancreatic cancer | 5 | 15 | 0.00934034 |
| Pathways in cancer,Cell cycle,Chronic myeloid leukemia | 6 | 21 | 0.00841347 |
| Pathways in cancer,Small cell lung cancer | 15 | 82 | 0.00220665 |
| Pathways in cancer,Small cell lung cancer,Chronic myeloid leukemia | 6 | 28 | 0.0249821 |
| Pathways in cancer,Colorectal cancer | 10 | 62 | 0.021515 |
| Pathways in cancer,Colorectal cancer,Endometrial cancer,Acute myeloid leukemia | 5 | 25 | 0.0472131 |
| Pathways in cancer,ErbB signaling pathway | 8 | 52 | 0.0430457 |
| Pathways in cancer,Chronic myeloid leukemia | 10 | 67 | 0.0313831 |
| Pathways in cancer,Endometrial cancer,Acute myeloid leukemia | 6 | 31 | 0.0367723 |
| Pathways in cancer,Basal cell carcinoma | 11 | 55 | 0.00444567 |
| Pathways in cancer,Basal cell carcinoma,Hedgehog signaling pathway | 6 | 32 | 0.0406529 |
| Pathways in cancer,Acute myeloid leukemia | 9 | 52 | 0.0204198 |
| Pathways in cancer,Thyroid cancer | 6 | 28 | 0.0249821 |
| Non-small cell lung cancer | 12 | 53 | 0.00161663 |
| beta-Alanine metabolism | 5 | 26 | 0.0499864 |
| Spliceosome | 24 | 118 | 2.84E-05 |
| Chemokine signaling pathway | 37 | 186 | 1.02E-07 |
| Chemokine signaling pathway,Leukocyte transendothelial migration | 7 | 31 | 0.0128622 |
| Chemokine signaling pathway,Gap junction | 6 | 31 | 0.0367723 |

| | | | |
|---|---|---|---|
| Chemokine signaling pathway,Epithelial cell signaling in Helicobacter pylori infection | 5 | 16 | 0.0114 |
| Chemokine signaling pathway,Chagas disease (American trypanosomiasis) | 6 | 32 | 0.0406529 |
| Chemokine signaling pathway,Amoebiasis | 6 | 23 | 0.0118268 |
| Toll-like receptor signaling pathway,Measles,Tuberculosis,African trypanosomiasis,Chagas disease (American trypanosomiasis) | 3 | 6 | 0.0168365 |
| Toll-like receptor signaling pathway,Malaria,Chagas disease (American trypanosomiasis) | 3 | 8 | 0.0330655 |
| Toll-like receptor signaling pathway,Chagas disease (American trypanosomiasis),Amoebiasis | 4 | 17 | 0.0468142 |
| Cytosolic DNA-sensing pathway | 9 | 60 | 0.0400532 |
| Measles | 32 | 130 | 5.09E-09 |
| Measles,Natural killer cell mediated cytotoxicity | 8 | 36 | 0.00888154 |
| Measles,Natural killer cell mediated cytotoxicity,T cell receptor signaling pathway,Osteoclast differentiation | 3 | 10 | 0.0491861 |
| Measles,Natural killer cell mediated cytotoxicity,Chagas disease (American trypanosomiasis) | 5 | 14 | 0.00714564 |
| Measles,Natural killer cell mediated cytotoxicity,Osteoclast differentiation | 6 | 15 | 0.00231127 |
| Measles,Tuberculosis,African trypanosomiasis,Malaria,Chagas disease (American trypanosomiasis) | 3 | 6 | 0.0168365 |
| Measles,Tuberculosis,African trypanosomiasis,Chagas disease (American trypanosomiasis) | 4 | 7 | 0.00338038 |
| Measles,Tuberculosis,Chagas disease (American trypanosomiasis) | 6 | 20 | 0.00683128 |
| Measles,African trypanosomiasis,Chagas disease (American trypanosomiasis) | 5 | 9 | 0.00157116 |
| Measles,Hematopoietic cell lineage | 4 | 8 | 0.00561468 |
| Measles,Hematopoietic cell lineage,T cell receptor signaling pathway,Chagas disease (American trypanosomiasis) | 3 | 3 | 0.00219324 |
| Measles,T cell receptor signaling pathway | 8 | 29 | 0.00290804 |
| Measles,T cell receptor signaling pathway,Chagas disease (American trypanosomiasis) | 5 | 20 | 0.0247545 |

| | | | |
|---|---|---|---|
| Measles,Adherens junction | 3 | 4 | 0.00562131 |
| Measles,Tight junction | 4 | 7 | 0.00338038 |
| Measles,Chagas disease (American trypanosomiasis) | 11 | 36 | 0.00031692 |
| Measles,Osteoclast differentiation | 7 | 33 | 0.0174564 |
| Measles,Primary immunodeficiency | 4 | 4 | 0.00043898 |
| Phagosome,Protein processing in endoplasmic reticulum | 3 | 6 | 0.0168365 |
| Calcium signaling pathway | 19 | 175 | 0.0471051 |
| Calcium signaling pathway,Focal adhesion | 3 | 10 | 0.0491861 |
| Calcium signaling pathway,Wnt signaling pathway,Long-term potentiation | 5 | 22 | 0.0329673 |
| Calcium signaling pathway,ErbB signaling pathway,Glioma | 4 | 10 | 0.0113403 |
| Ribosome biogenesis in eukaryotes | 30 | 73 | 7.93E-15 |
| Alanine, aspartate and glutamate metabolism | 6 | 32 | 0.0406529 |
| Arginine and proline metabolism | 8 | 54 | 0.0498951 |
| Nitrogen metabolism | 5 | 23 | 0.0386519 |
| Ribosome | 73 | 86 | 3.16E-70 |
| Protein processing in endoplasmic reticulum | 28 | 159 | 4.85E-05 |
| Protein processing in endoplasmic reticulum,Ubiquitin mediated proteolysis | 7 | 19 | 0.00162975 |
| Protein processing in endoplasmic reticulum,Protein export | 3 | 7 | 0.024676 |
| Bile secretion,Mineral absorption,Carbohydrate digestion and absorption | 3 | 10 | 0.0491861 |
| Natural killer cell mediated cytotoxicity | 23 | 125 | 0.00016837 |
| Natural killer cell mediated cytotoxicity,Tuberculosis | 6 | 34 | 0.04732 |
| Natural killer cell mediated cytotoxicity,Tuberculosis,Osteoclast differentiation | 4 | 12 | 0.0198264 |
| Natural killer cell mediated cytotoxicity,African trypanosomiasis | 4 | 7 | 0.00338038 |
| Natural killer cell mediated cytotoxicity,Type I diabetes mellitus,Allograft rejection,Graft-versus-host disease | 3 | 8 | 0.0330655 |

| | | | |
|---|---|---|---|
| Natural killer cell mediated cytotoxicity,Leishmaniasis | 4 | 10 | 0.0113403 |
| Natural killer cell mediated cytotoxicity,Chagas disease (American trypanosomiasis) | 6 | 17 | 0.00350999 |
| Natural killer cell mediated cytotoxicity,Osteoclast differentiation | 9 | 32 | 0.00174548 |
| Natural killer cell mediated cytotoxicity,Graft-versus-host disease | 4 | 13 | 0.0243223 |
| Tuberculosis | 24 | 172 | 0.00266876 |
| Tuberculosis,African trypanosomiasis | 5 | 9 | 0.00157116 |
| Tuberculosis,African trypanosomiasis,Malaria | 4 | 8 | 0.00561468 |
| Tuberculosis,Malaria | 5 | 15 | 0.00934034 |
| Tuberculosis,Type I diabetes mellitus | 4 | 16 | 0.0416621 |
| Cell adhesion molecules (CAMs) | 17 | 125 | 0.0127262 |
| Cell adhesion molecules (CAMs),T cell receptor signaling pathway | 4 | 9 | 0.00841625 |
| Cell adhesion molecules (CAMs),T cell receptor signaling pathway,Primary immunodeficiency | 3 | 6 | 0.0168365 |
| African trypanosomiasis | 12 | 33 | 2.54E-05 |
| African trypanosomiasis,Malaria | 6 | 14 | 0.00177582 |
| African trypanosomiasis,Malaria,Rheumatoid arthritis | 3 | 5 | 0.0108321 |
| African trypanosomiasis,Leukocyte transendothelial migration | 3 | 5 | 0.0108321 |
| African trypanosomiasis,Leishmaniasis,Amoebiasis | 4 | 6 | 0.00218312 |
| African trypanosomiasis,Chagas disease (American trypanosomiasis) | 6 | 15 | 0.00231127 |
| African trypanosomiasis,Chagas disease (American trypanosomiasis),Amoebiasis | 4 | 11 | 0.0153891 |
| African trypanosomiasis,Amoebiasis | 5 | 15 | 0.00934034 |
| Malaria | 11 | 48 | 0.00227156 |
| Malaria,Rheumatoid arthritis | 4 | 14 | 0.0298996 |
| Malaria,Chagas disease (American trypanosomiasis) | 4 | 14 | 0.0298996 |
| Hematopoietic cell lineage | 17 | 83 | 0.00048802 |

| | | | |
|---|---|---|---|
| Hematopoietic cell lineage,T cell receptor signaling pathway | 4 | 9 | 0.00841625 |
| Hematopoietic cell lineage,T cell receptor signaling pathway,Primary immunodeficiency | 3 | 5 | 0.0108321 |
| Hematopoietic cell lineage,Primary immunodeficiency | 4 | 7 | 0.00338038 |
| RNA degradation | 17 | 66 | 3.12E-05 |
| mRNA surveillance pathway | 15 | 79 | 0.00179499 |
| Arrhythmogenic right ventricular cardiomyopathy (ARVC),Adherens junction | 4 | 13 | 0.0243223 |
| Focal adhesion,Long-term potentiation | 4 | 15 | 0.0366428 |
| Focal adhesion,Gastric acid secretion | 3 | 7 | 0.024676 |
| Endocytosis | 22 | 193 | 0.0256267 |
| Endocrine and other factor-regulated calcium reabsorption,Salivary secretion,Gastric acid secretion | 5 | 23 | 0.0386519 |
| Homologous recombination | 7 | 26 | 0.00571677 |
| Vibrio cholerae infection | 8 | 53 | 0.0475913 |
| Vibrio cholerae infection,Leukocyte transendothelial migration | 3 | 7 | 0.024676 |
| Pathogenic Escherichia coli infection | 12 | 54 | 0.00172231 |
| T cell receptor signaling pathway | 23 | 107 | 1.99E-05 |
| T cell receptor signaling pathway,Chagas disease (American trypanosomiasis) | 6 | 33 | 0.0426965 |
| T cell receptor signaling pathway,Primary immunodeficiency | 6 | 11 | 0.00051547 |
| B cell receptor signaling pathway | 13 | 75 | 0.00565394 |
| Leukocyte transendothelial migration | 16 | 113 | 0.0113696 |
| Leukocyte transendothelial migration,Gastric acid secretion,Tight junction | 4 | 7 | 0.00338038 |
| Wnt signaling pathway | 27 | 149 | 4.37E-05 |
| Wnt signaling pathway,Oocyte meiosis | 6 | 29 | 0.0286494 |
| Wnt signaling pathway,Melanogenesis | 12 | 55 | 0.00198416 |

| | | | |
|---|---|---|---|
| Wnt signaling pathway,Adherens junction | 6 | 18 | 0.00433798 |
| Wnt signaling pathway,Tight junction | 4 | 11 | 0.0153891 |
| Wnt signaling pathway,Cell cycle,TGF-beta signaling pathway | 3 | 9 | 0.0418808 |
| Wnt signaling pathway,TGF-beta signaling pathway | 4 | 16 | 0.0416621 |
| Salivary secretion | 12 | 85 | 0.0270062 |
| Glutathione metabolism | 9 | 44 | 0.00880867 |
| p53 signaling pathway | 19 | 67 | 2.28E-06 |
| p53 signaling pathway,Oocyte meiosis,Progesterone-mediated oocyte maturation,Cell cycle | 3 | 4 | 0.00562131 |
| p53 signaling pathway,Oocyte meiosis,Cell cycle | 4 | 6 | 0.00218312 |
| p53 signaling pathway,Cell cycle | 8 | 24 | 0.00134996 |
| DNA replication | 9 | 35 | 0.00243855 |
| DNA replication,Cell cycle | 3 | 7 | 0.024676 |
| Mismatch repair | 6 | 23 | 0.0118268 |
| Prostate cancer | 13 | 88 | 0.0169944 |
| Mineral absorption | 9 | 51 | 0.0183953 |
| Oocyte meiosis | 25 | 110 | 2.11E-06 |
| Oocyte meiosis,Long-term potentiation | 6 | 34 | 0.04732 |
| Oocyte meiosis,Progesterone-mediated oocyte maturation | 9 | 51 | 0.0183953 |
| Oocyte meiosis,Progesterone-mediated oocyte maturation,Cell cycle | 7 | 22 | 0.00266209 |
| Oocyte meiosis,Ubiquitin mediated proteolysis,Cell cycle | 4 | 16 | 0.0416621 |
| Oocyte meiosis,Cell cycle | 15 | 40 | 1.03E-06 |
| Long-term potentiation | 11 | 68 | 0.0162363 |
| Pentose phosphate pathway | 7 | 27 | 0.00704145 |
| Type I diabetes mellitus | 6 | 34 | 0.04732 |
| Type I diabetes mellitus,Allograft rejection | 5 | 24 | 0.0413546 |

| | | | |
|---|---|---|---|
| Progesterone-mediated oocyte maturation | 15 | 86 | 0.00302358 |
| Progesterone-mediated oocyte maturation,Cell cycle | 9 | 29 | 0.00105293 |
| Melanogenesis | 16 | 98 | 0.00368463 |
| Melanogenesis,Gastric acid secretion | 6 | 33 | 0.0426965 |
| Gastric acid secretion | 12 | 72 | 0.0105397 |
| Glycosaminoglycan biosynthesis - heparan sulfate | 6 | 26 | 0.0194784 |
| Ubiquitin mediated proteolysis | 22 | 135 | 0.00103315 |
| Ubiquitin mediated proteolysis,Cell cycle | 5 | 19 | 0.0208959 |
| Adherens junction | 15 | 71 | 0.00080803 |
| Adherens junction,Bacterial invasion of epithelial cells | 4 | 17 | 0.0468142 |
| Tight junction | 17 | 130 | 0.0169558 |
| Tight junction,Chagas disease (American trypanosomiasis) | 4 | 14 | 0.0298996 |
| Rheumatoid arthritis | 13 | 84 | 0.0121721 |
| Glycosphingolipid biosynthesis - lacto and neolacto series | 8 | 25 | 0.00162214 |
| Epithelial cell signaling in Helicobacter pylori infection | 11 | 67 | 0.0149617 |
| Cell cycle | 39 | 123 | 9.11E-15 |
| Cell cycle,TGF-beta signaling pathway | 5 | 18 | 0.0171648 |
| Leishmaniasis | 10 | 66 | 0.0292868 |
| Chagas disease (American trypanosomiasis) | 22 | 102 | 2.41E-05 |
| Chagas disease (American trypanosomiasis),Amoebiasis | 6 | 34 | 0.04732 |
| Vitamin digestion and absorption | 5 | 24 | 0.0413546 |
| TGF-beta signaling pathway | 11 | 82 | 0.0415532 |
| ErbB signaling pathway | 13 | 87 | 0.0155366 |
| Bacterial invasion of epithelial cells | 13 | 70 | 0.00337183 |
| Bacterial invasion of epithelial cells,Shigellosis | 9 | 26 | 0.00048849 |

| | | | |
|---|---|---|---|
| Endometrial cancer | 9 | 52 | 0.0204198 |
| Intestinal immune network for IgA production,Primary immunodeficiency | 3 | 6 | 0.0168365 |
| Prion diseases | 6 | 34 | 0.04732 |
| Amoebiasis | 13 | 102 | 0.0400509 |
| Non-homologous end-joining | 4 | 13 | 0.0243223 |
| Carbohydrate digestion and absorption | 7 | 41 | 0.0409441 |
| Protein export | 7 | 23 | 0.00337763 |
| Tryptophan metabolism | 9 | 41 | 0.00569375 |
| Shigellosis | 15 | 61 | 0.00018009 |
| Acute myeloid leukemia | 10 | 57 | 0.0138373 |
| Allograft rejection | 6 | 29 | 0.0286494 |
| Primary immunodeficiency | 12 | 33 | 2.54E-05 |
| Glioma | 10 | 63 | 0.0240063 |
| Aminoacyl-tRNA biosynthesis | 13 | 40 | 3.26E-05 |
| One carbon pool by folate | 5 | 18 | 0.0171648 |

**Table S20. Differential gene expression in PABPC1 silenced HCC1806 cells versus control.**

Results of paired SAMR analysis (3 replicates) are shown. Significant downregulated genes (q< 5%) in silenced cells are shown. See Methods for further experimental and analysis details.

| Gene ID | Gene Name | Score(d) | Numerator(r) | Denominator(s+s0) | Fold Change | q-value(%) |
|---|---|---|---|---|---|---|
| LOC341315 | 22528 | -28.879 | -2.996 | 0.104 | 0.125 | 0 |
| PABPC1 | 35531 | -21.686 | -2.488 | 0.115 | 0.178 | 0 |

| | | | | | | |
|---|---|---|---|---|---|---|
| PABPC3 | 35536 | -14.431 | -1.491 | 0.103 | 0.356 | 0 |
| LOC100134273 | 21689 | -13.559 | -0.759 | 0.056 | 0.591 | 0 |
| PABPC1 | 35530 | -11.129 | -2.358 | 0.212 | 0.195 | 0 |
| PGAM4 | 36266 | -10.568 | -0.882 | 0.083 | 0.543 | 0 |
| MDK | 31704 | -8.84 | -0.725 | 0.082 | 0.605 | 0 |
| ASH2L | 1824 | -8.821 | -1.333 | 0.151 | 0.397 | 0 |
| CKB | 6035 | -8.168 | -0.778 | 0.095 | 0.583 | 0 |
| EFNB3 | 8692 | -8.165 | -0.731 | 0.09 | 0.602 | 0 |
| ASAP3 | 1771 | -8.159 | -0.58 | 0.071 | 0.669 | 0 |
| SLC48A1 | 41131 | -8.11 | -0.536 | 0.066 | 0.69 | 0 |
| SLC9A3R1 | 41230 | -8.028 | -0.493 | 0.061 | 0.71 | 0 |
| LOC388564 | 22746 | -7.973 | -0.706 | 0.089 | 0.613 | 0 |
| CD24 | 5313 | -7.814 | -0.887 | 0.114 | 0.541 | 0 |
| UBL5 | 45122 | -7.337 | -0.699 | 0.095 | 0.616 | 0 |
| SUMF2 | 42667 | -7.326 | -0.965 | 0.132 | 0.512 | 0 |
| ATP6V0E2 | 2059 | -7.204 | -0.752 | 0.104 | 0.594 | 0 |
| AXL | 2167 | -7.187 | -0.793 | 0.11 | 0.577 | 0 |
| MT1X | 33221 | -7.162 | -1.188 | 0.166 | 0.439 | 0 |
| LOC653888 | 29054 | -6.94 | -0.543 | 0.078 | 0.686 | 0 |
| KRT5 | 18549 | -6.934 | -0.632 | 0.091 | 0.645 | 0 |
| CCDC72 | 5079 | -6.687 | -1.115 | 0.167 | 0.462 | 0 |
| CACNB3 | 4616 | -6.657 | -0.402 | 0.06 | 0.757 | 0 |
| FKBP1A | 10402 | -6.611 | -0.552 | 0.084 | 0.682 | 0 |
| LOC728188 | 29442 | -6.514 | -0.619 | 0.095 | 0.651 | 0 |
| LOC728026 | 29358 | -6.431 | -0.587 | 0.091 | 0.666 | 0 |

| | | | | | | |
|---|---|---|---|---|---|---|
| FLOT2 | 10948 | -6.393 | -0.622 | 0.097 | 0.65 | 0 |
| HCFC1R1 | 12726 | -6.309 | -1.424 | 0.226 | 0.373 | 0.612 |
| C17orf89 | 3375 | -6.303 | -0.556 | 0.088 | 0.68 | 0.612 |
| FLJ22184 | 10548 | -6.302 | -0.823 | 0.131 | 0.565 | 0.612 |
| PTGFRN | 37912 | -6.29 | -0.442 | 0.07 | 0.736 | 0.612 |
| LOC441763 | 23821 | -6.289 | -1.185 | 0.188 | 0.44 | 0.612 |
| FBLN2 | 10046 | -6.286 | -0.565 | 0.09 | 0.676 | 0.612 |
| MXRA7 | 33388 | -6.248 | -0.814 | 0.13 | 0.569 | 0.612 |
| MT1E | 33210 | -6.153 | -0.737 | 0.12 | 0.6 | 0.612 |
| PTMS | 37942 | -6.122 | -0.726 | 0.119 | 0.605 | 0.612 |
| MATN2 | 31567 | -6.078 | -0.856 | 0.141 | 0.552 | 0.612 |
| LOC643319 | 24764 | -6.04 | -0.464 | 0.077 | 0.725 | 0.612 |
| LOC205251 | 22151 | -6.013 | -0.502 | 0.083 | 0.706 | 0.612 |
| PCBD1 | 35770 | -6.01 | -0.757 | 0.126 | 0.592 | 0.612 |
| DBI | 7423 | -5.978 | -0.586 | 0.098 | 0.666 | 0.612 |
| LOC100133565 | 21405 | -5.93 | -1.302 | 0.22 | 0.406 | 0.612 |
| ATP5E | 2013 | -5.915 | -0.727 | 0.123 | 0.604 | 0.612 |
| MGC4677 | 32046 | -5.89 | -0.848 | 0.144 | 0.555 | 0.612 |
| TIMP2 | 43535 | -5.883 | -0.788 | 0.134 | 0.579 | 0.612 |
| LGALS1 | 18942 | -5.787 | -1.38 | 0.238 | 0.384 | 0.612 |
| ATP5I | 2026 | -5.773 | -0.823 | 0.142 | 0.565 | 0.612 |
| LOC730382 | 30483 | -5.715 | -0.986 | 0.173 | 0.505 | 0.612 |
| C10orf116 | 2812 | -5.707 | -0.763 | 0.134 | 0.589 | 0.612 |
| LOC100129599 | 19848 | -5.678 | -0.787 | 0.139 | 0.58 | 0.612 |
| SUCLA2 | 42622 | -5.661 | -0.434 | 0.077 | 0.74 | 0.612 |

| | | | | | | |
|---|---|---|---|---|---|---|
| SPATA20 | 42040 | -5.654 | -0.416 | 0.074 | 0.75 | 0.612 |
| DLK2 | 8024 | -5.646 | -1.229 | 0.218 | 0.427 | 0.612 |
| LOC732007 | 30697 | -5.523 | -0.561 | 0.102 | 0.678 | 0.612 |
| PGAM4 | 36267 | -5.492 | -0.654 | 0.119 | 0.635 | 0.612 |
| A4GALT | 19 | -5.434 | -0.614 | 0.113 | 0.654 | 0.978 |
| TMEM191B | 43811 | -5.417 | -0.548 | 0.101 | 0.684 | 0.978 |
| FAM195B | 9708 | -5.347 | -0.398 | 0.074 | 0.759 | 0.978 |
| CRIP1 | 6727 | -5.338 | -1.705 | 0.319 | 0.307 | 0.978 |
| NPW | 34385 | -5.332 | -0.633 | 0.119 | 0.645 | 0.978 |
| PDCD6IP | 36010 | -5.306 | -0.444 | 0.084 | 0.735 | 0.978 |
| LOC730102 | 30387 | -5.275 | -0.401 | 0.076 | 0.758 | 0.978 |
| IFI6 | 16911 | -5.252 | -0.569 | 0.108 | 0.674 | 0.978 |
| PKM2 | 36618 | -5.119 | -0.438 | 0.086 | 0.738 | 0.978 |
| LAMP2 | 18750 | -5.111 | -0.699 | 0.137 | 0.616 | 0.978 |
| MYL9 | 33464 | -5.104 | -0.33 | 0.065 | 0.795 | 0.978 |
| TSPO | 44688 | -5.101 | -0.677 | 0.133 | 0.626 | 0.978 |
| LOC643272 | 24738 | -5.093 | -0.657 | 0.129 | 0.634 | 0.978 |
| LOC400948 | 23259 | -5.039 | -0.67 | 0.133 | 0.629 | 0.978 |
| PBX2 | 35760 | -5.012 | -0.482 | 0.096 | 0.716 | 0.978 |
| LOC439953 | 23456 | -5.011 | -0.677 | 0.135 | 0.625 | 0.978 |
| NUTF2 | 34689 | -5.007 | -0.746 | 0.149 | 0.596 | 0.978 |
| AP2M1 | 1317 | -4.971 | -0.473 | 0.095 | 0.72 | 0.978 |
| S100A13 | 39710 | -4.96 | -0.868 | 0.175 | 0.548 | 0.978 |
| LOC729645 | 30173 | -4.943 | -1.032 | 0.209 | 0.489 | 0.978 |
| MID1IP1 | 32119 | -4.937 | -0.497 | 0.101 | 0.709 | 2.362 |

| | | | | | | |
|---|---|---|---|---|---|---|
| PPDPF | 37144 | -4.937 | -0.738 | 0.149 | 0.6 | 2.362 |
| CSTB | 6926 | -4.936 | -0.705 | 0.143 | 0.613 | 2.362 |
| LOC729082 | 29899 | -4.926 | -0.455 | 0.092 | 0.73 | 2.362 |
| C1QTNF6 | 3696 | -4.915 | -0.81 | 0.165 | 0.57 | 2.362 |
| GTF2H5 | 12520 | -4.882 | -0.396 | 0.081 | 0.76 | 2.362 |
| KRT13 | 18494 | -4.85 | -0.386 | 0.08 | 0.765 | 2.362 |
| CAV2 | 4861 | -4.849 | -0.495 | 0.102 | 0.71 | 2.362 |
| ZNF358 | 46734 | -4.845 | -0.721 | 0.149 | 0.607 | 2.362 |
| FAM69B | 9848 | -4.84 | -0.517 | 0.107 | 0.699 | 2.362 |
| C10orf75 | 2884 | -4.8 | -0.631 | 0.131 | 0.646 | 2.362 |
| GAMT | 11500 | -4.796 | -0.802 | 0.167 | 0.573 | 2.362 |
| LOC100132499 | 20972 | -4.789 | -0.702 | 0.147 | 0.615 | 2.362 |
| CRYBB2 | 6791 | -4.781 | -0.717 | 0.15 | 0.608 | 2.362 |
| VTI1B | 45722 | -4.779 | -0.439 | 0.092 | 0.738 | 2.362 |
| EIF5A | 8829 | -4.776 | -0.955 | 0.2 | 0.516 | 2.362 |
| PDPN | 36128 | -4.762 | -0.696 | 0.146 | 0.617 | 2.362 |
| LOC643997 | 25105 | -4.746 | -0.689 | 0.145 | 0.62 | 2.362 |
| LOC653219 | 28695 | -4.727 | -0.617 | 0.131 | 0.652 | 2.362 |
| MTE | 33234 | -4.717 | -0.883 | 0.187 | 0.542 | 2.362 |
| HBQ1 | 12715 | -4.708 | -0.818 | 0.174 | 0.567 | 2.362 |
| FKBP1A | 10401 | -4.687 | -0.442 | 0.094 | 0.736 | 2.362 |
| TRMT112 | 44523 | -4.673 | -0.536 | 0.115 | 0.69 | 2.362 |
| MAGEA9B | 31254 | -4.664 | -0.446 | 0.096 | 0.734 | 2.362 |
| GAMT | 11498 | -4.647 | -0.989 | 0.213 | 0.504 | 2.362 |
| FBLN2 | 10045 | -4.638 | -0.445 | 0.096 | 0.734 | 2.362 |

| | | | | | | |
|---|---|---|---|---|---|---|
| OLFML2A | 34816 | -4.633 | -1.169 | 0.252 | 0.445 | 2.362 |
| GCNT1 | 11609 | -4.63 | -0.664 | 0.143 | 0.631 | 2.362 |
| LOC643287 | 24749 | -4.59 | -0.927 | 0.202 | 0.526 | 2.362 |
| MT1A | 33208 | -4.583 | -0.845 | 0.184 | 0.557 | 2.362 |
| COMMD7 | 6495 | -4.579 | -0.6 | 0.131 | 0.66 | 2.362 |
| LSS | 31067 | -4.566 | -0.402 | 0.088 | 0.757 | 2.362 |
| LOC729859 | 30271 | -4.552 | -0.334 | 0.073 | 0.794 | 2.362 |
| SLC27A3 | 40950 | -4.549 | -0.288 | 0.063 | 0.819 | 2.362 |
| LOC728453 | 29562 | -4.539 | -0.571 | 0.126 | 0.673 | 2.362 |
| C2orf7 | 3977 | -4.537 | -0.813 | 0.179 | 0.569 | 2.362 |
| C19orf12 | 3420 | -4.532 | -0.398 | 0.088 | 0.759 | 2.362 |
| EFCAB4A | 8658 | -4.522 | -0.673 | 0.149 | 0.627 | 2.362 |
| LAMA3 | 18725 | -4.521 | -0.42 | 0.093 | 0.747 | 2.362 |
| TRAPPC1 | 44330 | -4.513 | -0.839 | 0.186 | 0.559 | 2.362 |
| LOC730029 | 30348 | -4.494 | -0.745 | 0.166 | 0.597 | 2.362 |
| LOC100133328 | 21324 | -4.494 | -0.882 | 0.196 | 0.543 | 2.362 |
| LOC100133390 | 21333 | -4.489 | -0.481 | 0.107 | 0.717 | 2.362 |
| SUCLG2 | 42624 | -4.488 | -0.457 | 0.102 | 0.729 | 2.362 |
| LOC643161 | 24686 | -4.482 | -0.364 | 0.081 | 0.777 | 2.362 |
| PLAC8 | 36675 | -4.468 | -1.07 | 0.239 | 0.476 | 2.362 |
| ESPN | 9254 | -4.466 | -0.504 | 0.113 | 0.705 | 2.362 |
| SEPX1 | 40236 | -4.448 | -0.306 | 0.069 | 0.809 | 2.362 |
| BNIP3 | 2592 | -4.442 | -0.746 | 0.168 | 0.596 | 2.362 |
| INF2 | 17272 | -4.432 | -0.37 | 0.083 | 0.774 | 2.362 |
| GPX2 | 12292 | -4.398 | -0.618 | 0.14 | 0.652 | 2.362 |

| | | | | | | |
|---|---|---|---|---|---|---|
| NME4 | 34203 | -4.39 | -0.765 | 0.174 | 0.588 | 2.362 |
| TOMM5 | 44164 | -4.374 | -0.341 | 0.078 | 0.789 | 2.362 |
| LOC388707 | 22758 | -4.373 | -0.631 | 0.144 | 0.646 | 2.362 |
| GNAI2 | 11937 | -4.369 | -0.66 | 0.151 | 0.633 | 2.362 |
| UCA1 | 45174 | -4.346 | -0.533 | 0.123 | 0.691 | 4.253 |
| OSTF1 | 35406 | -4.342 | -0.452 | 0.104 | 0.731 | 4.253 |
| MID1IP1 | 32118 | -4.337 | -0.382 | 0.088 | 0.767 | 4.253 |
| CDC26 | 5457 | -4.334 | -0.463 | 0.107 | 0.726 | 4.253 |
| PPP2R1B | 37277 | -4.329 | -0.531 | 0.123 | 0.692 | 4.253 |
| LPCAT1 | 30833 | -4.323 | -0.464 | 0.107 | 0.725 | 4.253 |
| TSPAN4 | 44677 | -4.316 | -0.465 | 0.108 | 0.725 | 4.253 |
| XAGE1A | 46055 | -4.312 | -0.502 | 0.117 | 0.706 | 4.253 |
| LOC100132037 | 20813 | -4.306 | -0.536 | 0.125 | 0.69 | 4.253 |
| SKA2 | 40679 | -4.304 | -0.544 | 0.126 | 0.686 | 4.253 |
| SRPK2 | 42285 | -4.289 | -0.991 | 0.231 | 0.503 | 4.253 |
| TMEM219 | 43854 | -4.275 | -0.509 | 0.119 | 0.703 | 4.253 |
| ITGAE | 17490 | -4.274 | -0.871 | 0.204 | 0.547 | 4.253 |
| TUBB6 | 44893 | -4.265 | -0.526 | 0.123 | 0.695 | 4.253 |
| DSC3 | 8352 | -4.256 | -0.365 | 0.086 | 0.776 | 4.253 |
| SERF2 | 40247 | -4.242 | -1.066 | 0.251 | 0.478 | 4.253 |
| NME2 | 34200 | -4.234 | -0.854 | 0.202 | 0.553 | 4.253 |
| CDC42 | 5473 | -4.229 | -0.468 | 0.111 | 0.723 | 4.253 |
| IQGAP1 | 17396 | -4.208 | -0.509 | 0.121 | 0.703 | 4.253 |
| DCAKD | 7454 | -4.202 | -0.322 | 0.077 | 0.8 | 4.253 |
| TARS2 | 42954 | -4.197 | -0.331 | 0.079 | 0.795 | 4.253 |

| | | | | | | |
|---|---|---|---|---|---|---|
| | 13947 | -4.195 | -0.429 | 0.102 | 0.743 | 4.253 |
| LOC440731 | 23604 | -4.178 | -1.019 | 0.244 | 0.493 | 4.253 |
| S1PR5 | 39740 | -4.151 | -0.471 | 0.114 | 0.721 | 4.253 |
| MXD1 | 33379 | -4.145 | -0.396 | 0.096 | 0.76 | 4.253 |
| POLE4 | 36967 | -4.142 | -0.643 | 0.155 | 0.641 | 4.253 |
| PACSIN1 | 35552 | -4.135 | -0.765 | 0.185 | 0.589 | 4.253 |
| CALML5 | 4673 | -4.131 | -0.721 | 0.174 | 0.607 | 4.253 |
| KRT10 | 18490 | -4.116 | -0.595 | 0.145 | 0.662 | 4.253 |
| TAGLN2 | 42919 | -4.116 | -0.771 | 0.187 | 0.586 | 4.253 |
| SSH3 | 42320 | -4.11 | -0.709 | 0.173 | 0.612 | 4.253 |
| NDUFS5 | 33881 | -4.107 | -0.358 | 0.087 | 0.78 | 4.253 |
| BANF1 | 2277 | -4.078 | -0.656 | 0.161 | 0.634 | 4.253 |
| HCFC1R1 | 12725 | -4.073 | -1.247 | 0.306 | 0.421 | 4.253 |
| NRSN2 | 34483 | -4.069 | -0.438 | 0.108 | 0.738 | 4.253 |
| | 13469 | -4.057 | -0.263 | 0.065 | 0.833 | 4.253 |
| C14orf112 | 3095 | -4.048 | -0.581 | 0.143 | 0.669 | 4.253 |
| LOC729255 | 29979 | -4.04 | -0.969 | 0.24 | 0.511 | 4.253 |
| MXRA5 | 33386 | -4.029 | -0.644 | 0.16 | 0.64 | 4.253 |
| ZNF580 | 46949 | -4.016 | -0.478 | 0.119 | 0.718 | 4.253 |
| TRIM6 | 44472 | -4.008 | -0.436 | 0.109 | 0.739 | 4.253 |
| PGAM1 | 36262 | -4.008 | -0.566 | 0.141 | 0.676 | 4.253 |
| STMN1 | 42542 | -4.004 | -0.992 | 0.248 | 0.503 | 4.253 |
| EPAS1 | 8989 | -3.986 | -0.61 | 0.153 | 0.655 | 4.253 |
| PCDHGB6 | 35911 | -3.982 | -0.434 | 0.109 | 0.74 | 4.253 |
| LOC651064 | 27858 | -3.972 | -0.721 | 0.182 | 0.607 | 4.253 |

| | | | | | | |
|---|---|---|---|---|---|---|
| RAB28 | 38210 | -3.964 | -0.302 | 0.076 | 0.811 | 4.253 |
| CENPN | 5703 | -3.956 | -0.369 | 0.093 | 0.775 | 4.253 |
| LRRC26 | 30932 | -3.941 | -0.623 | 0.158 | 0.649 | 4.253 |
| PIK3C2B | 36494 | -3.935 | -0.321 | 0.082 | 0.801 | 4.253 |
| LOC130773 | 21939 | -3.929 | -0.618 | 0.157 | 0.651 | 4.253 |
| EEF1A2 | 8638 | -3.923 | -0.681 | 0.174 | 0.624 | 4.253 |
| CNTNAP1 | 6367 | -3.915 | -0.357 | 0.091 | 0.781 | 4.253 |

**Table S21. KEGG pathway analysis of genes differentially expressed in PABPC1 silenced HCC1806 cells versus control.**

Results of a KEGG pathway enrichment analysis using Genecois3 (http://www.genecodis.cnb.csic.es) for genes co-expressed with PABPC1 in the clinical samples (downregulated genes shown in Table S21). Only significantly (adjusted p<0.05) enriched pathway are show.

| Items | Pathway Details | Support | List size | Reference Support | Reference size | Hypergeometric Test p-value | Hypergeometric Test p-value (FDR adjusted) | Genes |
|---|---|---|---|---|---|---|---|---|
| Kegg:00010 | Glycolysis / Gluconeogenesis | 3 | 132 | 62 | 34208 | 0.00179863 | 0.032735 | PKM2,PGAM1,PGAM4 |
| Kegg:00190 | Oxidative phosphorylation | 4 | 132 | 130 | 34208 | 0.00165305 | 0.0376068 | NDUFS5,ATP5I,ATP6V0E2,ATP5E |

| | | | | | | | | |
|---|---|---|---|---|---|---|---|---|
| Kegg:03015 | mRNA surveillance pathway | 3 | 132 | 79 | 34208 | 0.00358597 | 0.0466176 | PABPC1,PABPC3,PPP2R1B |
| Kegg:04530 | Tight junction | 4 | 132 | 130 | 34208 | 0.00165305 | 0.0376068 | CDC42,PPP2R1B,GNAI2,MYL9 |

**Table S22. miRNA expression in control cells compared with AGO2 silenced HCC1806 cells under hypoxic conditions (1% O2).**

Results of Limma analysis are shown, positive t values correspond to lower expression in AGO2 silenced cells (hence downregulated by loss of AGO2 expression), negative t values correspond to higher expression in AGO2 silenced (hence upregulated by loss of AGO2 expression). See Methods for further experimental and analysis details.

| GeneName | ProbeName | AveExpr | t | P.Value | adj.P.Val | B |
|---|---|---|---|---|---|---|
| hsa-miR-4520-2-3p | A_25_P00016397 | 40.68 | 19.37 | 2.95E-14 | 1.43E-10 | 17.17 |
| hsa-miR-24-3p | A_25_P00010677 | 4480.34 | 16.03 | 9.78E-13 | 1.92E-09 | 15.38 |
| hsa-miR-324-5p | A_25_P00010154 | 271.75 | 14.92 | 3.60E-12 | 4.40E-09 | 14.63 |
| hsa-let-7i-5p | A_25_P00012145 | 316.11 | 14.89 | 3.73E-12 | 4.40E-09 | 14.61 |
| hsa-miR-4291 | A_25_P00015568 | 226.12 | 13.70 | 1.68E-11 | 1.42E-08 | 13.69 |
| hsa-miR-24-3p | A_25_P00010676 | 5186.72 | 13.69 | 1.69E-11 | 1.42E-08 | 13.69 |
| hsa-miR-4291 | A_25_P00015567 | 162.93 | 13.43 | 2.40E-11 | 1.77E-08 | 13.47 |
| hsa-miR-324-5p | A_25_P00010153 | 384.54 | 13.24 | 3.08E-11 | 2.02E-08 | 13.31 |
| hsa-miR-107 | A_25_P00011069 | 1001.22 | 12.67 | 6.70E-11 | 3.95E-08 | 12.80 |
| hsa-miR-4520-2-3p | A_25_P00016398 | 41.13 | 12.05 | 1.61E-10 | 7.91E-08 | 12.21 |
| hsa-miR-107 | A_25_P00011068 | 407.83 | 11.96 | 1.84E-10 | 8.33E-08 | 12.11 |
| hsa-let-7i-5p | A_25_P00012146 | 460.68 | 11.81 | 2.29E-10 | 9.00E-08 | 11.96 |
| hsa-miR-30d-5p | A_25_P00010682 | 158.87 | 11.68 | 2.77E-10 | 1.02E-07 | 11.83 |
| hsa-miR-183-5p | A_25_P00012098 | 227.92 | 11.37 | 4.39E-10 | 1.52E-07 | 11.51 |
| hsa-miR-99b-5p | A_25_P00014849 | 191.60 | 11.17 | 5.96E-10 | 1.95E-07 | 11.29 |
| hsa-miR-301a-3p | A_25_P00010839 | 258.07 | 10.49 | 1.72E-09 | 5.35E-07 | 10.52 |
| hsa-let-7d-5p | A_25_P00011980 | 258.87 | 10.30 | 2.34E-09 | 6.90E-07 | 10.29 |
| hsa-miR-148b-3p | A_25_P00010133 | 159.54 | 9.69 | 6.44E-09 | 1.81E-06 | 9.53 |
| hsa-miR-429 | A_25_P00010273 | 526.06 | 9.56 | 8.07E-09 | 2.16E-06 | 9.35 |
| hsa-miR-30d-5p | A_25_P00010683 | 129.74 | 9.42 | 1.02E-08 | 2.52E-06 | 9.17 |
| hsa-miR-183-5p | A_25_P00012099 | 231.41 | 9.41 | 1.04E-08 | 2.52E-06 | 9.16 |
| hsa-miR-505-3p | A_25_P00012654 | 62.74 | 9.39 | 1.07E-08 | 2.52E-06 | 9.14 |

| | | | | | | |
|---|---|---|---|---|---|---|
| hsa-miR-429 | A_25_P00014860 | 433.16 | 9.18 | 1.55E-08 | 3.38E-06 | 8.85 |
| hsa-miR-183-5p | A_25_P00012097 | 106.71 | 8.74 | 3.38E-08 | 6.95E-06 | 8.23 |
| hsa-miR-99b-5p | A_25_P00010597 | 109.28 | 8.73 | 3.42E-08 | 6.95E-06 | 8.22 |
| hsa-miR-105-5p | A_25_P00012043 | 67.27 | 8.69 | 3.69E-08 | 7.25E-06 | 8.16 |
| hsa-miR-221-3p | A_25_P00010690 | 859.49 | 8.58 | 4.49E-08 | 8.53E-06 | 8.01 |
| hsa-let-7d-5p | A_25_P00011981 | 620.69 | 8.57 | 4.63E-08 | 8.53E-06 | 7.98 |
| hsa-miR-1307-3p | A_25_P00015256 | 55.21 | 8.39 | 6.37E-08 | 1.10E-05 | 7.73 |
| hsa-miR-15b-5p | A_25_P00011101 | 2823.53 | 8.27 | 8.02E-08 | 1.35E-05 | 7.54 |
| hsa-miR-342-3p | A_25_P00012358 | 174.11 | 8.25 | 8.35E-08 | 1.37E-05 | 7.51 |
| hsa-miR-221-3p | A_25_P00010689 | 243.73 | 8.22 | 8.79E-08 | 1.40E-05 | 7.47 |
| hsa-miR-301b-3p | A_25_P00013025 | 43.92 | 8.16 | 9.84E-08 | 1.53E-05 | 7.37 |
| hsa-miR-301a-3p | A_25_P00013973 | 91.66 | 7.89 | 1.66E-07 | 2.51E-05 | 6.95 |
| hsa-miR-103a-3p | A_25_P00011005 | 876.76 | 7.77 | 2.07E-07 | 3.05E-05 | 6.77 |
| hsa-miR-425-5p | A_25_P00014045 | 61.19 | 7.65 | 2.60E-07 | 3.73E-05 | 6.58 |
| hsa-let-7g-5p | A_25_P00012142 | 563.62 | 7.58 | 2.98E-07 | 4.18E-05 | 6.46 |
| hsa-let-7e-5p | A_25_P00011984 | 499.08 | 7.46 | 3.83E-07 | 5.25E-05 | 6.25 |
| hsa-miR-23a-3p | A_25_P00010843 | 1294.36 | 7.31 | 5.12E-07 | 6.69E-05 | 6.01 |
| hsa-miR-342-3p | A_25_P00012357 | 89.58 | 7.30 | 5.20E-07 | 6.69E-05 | 6.00 |
| hsa-miR-301b-3p | A_25_P00013027 | 112.87 | 7.30 | 5.22E-07 | 6.69E-05 | 6.00 |
| hsa-miR-330-3p | A_25_P00011046 | 50.93 | 7.24 | 5.85E-07 | 7.33E-05 | 5.90 |
| hsa-miR-103a-3p | A_25_P00011004 | 2138.82 | 7.14 | 7.23E-07 | 8.70E-05 | 5.72 |
| hsa-miR-320c | A_25_P00015036 | 65.68 | 6.87 | 1.24E-06 | 0.000146654 | 5.26 |
| hsa-miR-320c | A_25_P00015037 | 360.35 | 6.85 | 1.29E-06 | 0.000149208 | 5.23 |
| hsa-miR-152-3p | A_25_P00012196 | 92.79 | 6.71 | 1.75E-06 | 0.000197891 | 4.97 |
| hsa-let-7e-5p | A_25_P00011985 | 826.98 | 6.61 | 2.16E-06 | 0.000239642 | 4.79 |
| hsa-miR-138-5p | A_25_P00012170 | 108.65 | 6.51 | 2.62E-06 | 0.000285608 | 4.63 |
| hsa-miR-183-3p | A_25_P00013323 | 69.85 | 6.42 | 3.20E-06 | 0.000343039 | 4.45 |
| hsa-miR-505-5p | A_25_P00013608 | 51.38 | 6.30 | 4.09E-06 | 0.000428755 | 4.24 |
| hsa-miR-301b-3p | A_25_P00013026 | 58.35 | 6.30 | 4.15E-06 | 0.000428755 | 4.23 |

| | | | | | | |
|---|---|---|---|---|---|---|
| hsa-miR-454-3p | A_25_P00012871 | 51.77 | 6.27 | 4.37E-06 | 0.000443817 | 4.18 |
| hsa-miR-98-5p | A_25_P00010047 | 127.77 | 6.18 | 5.35E-06 | 0.000528078 | 4.01 |
| hsa-miR-15b-5p | A_25_P00011102 | 1711.71 | 6.18 | 5.38E-06 | 0.000528078 | 4.00 |
| hsa-miR-3200-3p | A_25_P00015620 | 48.50 | 5.96 | 8.54E-06 | 0.000811766 | 3.60 |
| hsa-miR-222-3p | A_25_P00012126 | 246.51 | 5.91 | 9.56E-06 | 0.000894055 | 3.50 |
| hsa-miR-330-3p | A_25_P00014009 | 40.19 | 5.73 | 1.41E-05 | 0.001298915 | 3.16 |
| hsa-miR-155-5p | A_25_P00012270 | 86.52 | 5.66 | 1.66E-05 | 0.001503495 | 3.02 |
| hsa-let-7g-5p | A_25_P00012141 | 248.02 | 5.47 | 2.50E-05 | 0.002236312 | 2.66 |
| hsa-miR-532-5p | A_25_P00014179 | 60.27 | 5.46 | 2.55E-05 | 0.002243157 | 2.64 |
| hsa-miR-130a-3p | A_25_P00010439 | 2833.22 | 5.38 | 3.05E-05 | 0.002605614 | 2.48 |
| hsa-miR-205-5p | A_25_P00010504 | 19172.69 | 5.37 | 3.18E-05 | 0.002635766 | 2.45 |
| hsa-miR-23b-3p | A_25_P00010881 | 344.01 | 5.35 | 3.29E-05 | 0.002658014 | 2.41 |
| hsa-miR-744-5p | A_25_P00012986 | 72.71 | 5.31 | 3.57E-05 | 0.002804729 | 2.34 |
| hsa-miR-181a-5p | A_25_P00014832 | 158.37 | 5.30 | 3.71E-05 | 0.002877583 | 2.31 |
| hsa-miR-106b-5p | A_25_P00010434 | 438.24 | 5.13 | 5.34E-05 | 0.004037997 | 1.99 |
| hsa-miR-652-3p | A_25_P00012834 | 65.00 | 5.08 | 6.04E-05 | 0.004294047 | 1.88 |
| hsa-miR-222-3p | A_25_P00012125 | 57.85 | 5.08 | 6.05E-05 | 0.004294047 | 1.88 |
| hsa-miR-362-5p | A_25_P00013983 | 42.79 | 5.04 | 6.63E-05 | 0.004649074 | 1.79 |
| hsa-miR-148b-3p | A_25_P00010134 | 49.01 | 5.03 | 6.77E-05 | 0.004689774 | 1.78 |
| hsa-miR-138-5p | A_25_P00012169 | 68.32 | 5.03 | 6.84E-05 | 0.004689774 | 1.77 |
| hsa-miR-183-3p | A_25_P00013322 | 46.39 | 5.00 | 7.24E-05 | 0.004901974 | 1.72 |
| hsa-miR-130b-3p | A_25_P00010437 | 497.29 | 4.98 | 7.56E-05 | 0.005059891 | 1.68 |
| hsa-miR-320e | A_25_P00015664 | 360.91 | 4.96 | 7.89E-05 | 0.00522229 | 1.64 |
| hsa-miR-98-5p | A_25_P00010048 | 47.49 | 4.85 | 0.000101764 | 0.006663295 | 1.41 |
| hsa-miR-320e | A_25_P00015663 | 83.00 | 4.84 | 0.000105394 | 0.00680176 | 1.38 |
| hsa-miR-128-3p | A_25_P00012161 | 64.53 | 4.83 | 0.000106187 | 0.00680176 | 1.37 |
| hsa-miR-3200-3p | A_25_P00015619 | 42.28 | 4.79 | 0.000117092 | 0.007419579 | 1.29 |

| | | | | | | |
|---|---|---|---|---|---|---|
| hsa-miR-182-5p | A_25_P00012095 | 87.58 | 4.78 | 0.00011948 | 0.007490406 | 1.27 |
| hsa-miR-744-5p | A_25_P00012987 | 88.86 | 4.74 | 0.000129951 | 0.007894869 | 1.19 |
| hsa-miR-505-5p | A_25_P00013607 | 46.73 | 4.66 | 0.000157816 | 0.00939404 | 1.02 |
| hsa-miR-769-5p | A_25_P00011954 | 66.14 | 4.66 | 0.000159474 | 0.009397831 | 1.01 |
| hsa-miR-877-5p | A_25_P00012999 | 47.50 | 4.64 | 0.000164174 | 0.009579007 | 0.99 |
| hsa-miR-106b-5p | A_25_P00010433 | 1084.13 | 4.62 | 0.000171225 | 0.009892439 | 0.95 |
| hsa-let-7c-5p | A_25_P00010072 | 655.84 | 4.60 | 0.000182574 | 0.010445716 | 0.89 |
| hsa-miR-17-5p | A_25_P00014819 | 1723.12 | 4.57 | 0.000195526 | 0.011079178 | 0.83 |
| hsa-miR-320d | A_25_P00015271 | 481.06 | 4.54 | 0.000206854 | 0.01160944 | 0.78 |
| hsa-miR-182-5p | A_25_P00012094 | 52.39 | 4.50 | 0.000226628 | 0.012599231 | 0.70 |
| hsa-miR-182-5p | A_25_P00012093 | 53.14 | 4.48 | 0.0002405 | 0.013245496 | 0.64 |
| hsa-miR-137 | A_25_P00010616 | 101.76 | 4.40 | 0.000290343 | 0.015842536 | 0.48 |
| hsa-miR-769-5p | A_25_P00011955 | 50.88 | 4.39 | 0.000294018 | 0.015895858 | 0.46 |
| hsa-miR-181a-5p | A_25_P00010285 | 80.46 | 4.30 | 0.000364914 | 0.019030405 | 0.27 |
| hsa-miR-203a-3p | A_25_P00010628 | 141.79 | 4.25 | 0.000407923 | 0.021086759 | 0.17 |
| hsa-miR-331-3p | A_25_P00014027 | 423.51 | 4.21 | 0.000450105 | 0.022866726 | 0.08 |
| hsa-miR-4690-5p | A_25_P00017267 | 41.31 | 4.20 | 0.000453484 | 0.022866726 | 0.08 |
| hsa-miR-194-5p | A_25_P00011007 | 45.97 | 4.20 | 0.000456082 | 0.022866726 | 0.07 |
| hsa-let-7b-5p | A_25_P00010070 | 2052.89 | 4.20 | 0.000457878 | 0.022866726 | 0.07 |
| hsa-let-7f-5p | A_25_P00010088 | 3360.81 | 4.19 | 0.000464575 | 0.023006229 | 0.05 |
| hsa-miR-455-5p | A_25_P00014182 | 60.99 | 4.18 | 0.000477576 | 0.023367835 | 0.03 |
| hsa-miR-574-5p | A_25_P00012724 | 125.58 | 4.16 | 0.000504343 | 0.024257141 | -0.02 |
| hsa-miR-93-5p | A_25_P00010610 | 1183.24 | 4.16 | 0.0005063 | 0.024257141 | -0.02 |
| hsa-miR-425-5p | A_25_P00010977 | 105.73 | 4.14 | 0.000524734 | 0.024937579 | -0.05 |
| hsa-miR-514b-5p | A_25_P00015585 | 36.79 | 4.13 | 0.000534878 | 0.025216288 | -0.07 |
| hsa-miR-23b-3p | A_25_P00010882 | 121.66 | 4.13 | 0.000542034 | 0.025225366 | -0.08 |
| hsa-miR-135b-5p | A_25_P00012383 | 101.90 | 4.12 | 0.000543632 | 0.025225366 | -0.09 |
| hsa-miR-486-5p | A_25_P00014064 | 57.88 | 4.09 | 0.000587483 | 0.026837498 | -0.16 |

| | | | | | | |
|---|---|---|---|---|---|---|
| hsa-miR-200a-3p | A_25_P00010208 | 153.96 | 4.07 | 0.000622968 | 0.028024053 | -0.21 |
| hsa-miR-503-5p | A_25_P00014885 | 54.04 | 4.03 | 0.000669719 | 0.029588647 | -0.27 |
| hsa-miR-4739 | A_25_P00016507 | 45.87 | 4.03 | 0.000672812 | 0.029588647 | -0.28 |
| hsa-miR-186-5p | A_25_P00012243 | 47.71 | 4.03 | 0.000683527 | 0.02983722 | -0.29 |
| hsa-miR-331-3p | A_25_P00014026 | 229.98 | 3.99 | 0.000737214 | 0.031710948 | -0.36 |
| hsa-miR-137 | A_25_P00010617 | 88.78 | 3.98 | 0.000756416 | 0.032301137 | -0.38 |
| hsa-miR-450a-5p | A_25_P00012437 | 40.37 | 3.96 | 0.000791994 | 0.033553429 | -0.42 |
| hsa-miR-23a-3p | A_25_P00014820 | 4287.60 | 3.96 | 0.000797129 | 0.033553429 | -0.43 |
| hsa-miR-1273f | A_25_P00016636 | 42.52 | 3.93 | 0.000849711 | 0.035513082 | -0.49 |
| hsa-miR-486-5p | A_25_P00014063 | 50.07 | 3.84 | 0.001056526 | 0.043236863 | -0.68 |
| hsa-miR-362-5p | A_25_P00013984 | 57.84 | 3.78 | 0.001215498 | 0.049399503 | -0.81 |
| hsa-miR-6809-3p | A_25_P00018565 | 36.47 | -3.86 | 0.001006968 | 0.041496925 | -0.64 |
| hsa-miR-21-3p | A_25_P00013173 | 307.23 | -3.93 | 0.00085673 | 0.035554307 | -0.49 |
| hsa-miR-19a-3p | A_25_P00010997 | 481.12 | -4.00 | 0.00073132 | 0.031688744 | -0.35 |
| hsa-miR-1273g-3p | A_25_P00017530 | 2245.75 | -4.04 | 0.000664311 | 0.029588647 | -0.27 |
| hsa-miR-181a-2-3p | A_25_P00013315 | 51.90 | -4.09 | 0.000594517 | 0.026949891 | -0.17 |
| hsa-miR-19b-1-5p | A_25_P00013163 | 55.52 | -4.10 | 0.000578776 | 0.0266463 | -0.14 |
| hsa-miR-4286 | A_25_P00015774 | 3845.49 | -4.18 | 0.000479808 | 0.023367835 | 0.03 |
| hsa-miR-1202 | A_25_P00015076 | 217.28 | -4.35 | 0.000324772 | 0.017088208 | 0.38 |
| hsa-miR-1260a | A_25_P00015172 | 269.11 | -4.35 | 0.000321934 | 0.017088208 | 0.38 |
| hsa-miR-193b-5p | A_25_P00013596 | 49.75 | -4.37 | 0.000306644 | 0.016427731 | 0.43 |
| hsa-miR-6881-3p | A_25_P00018175 | 35.94 | -4.73 | 0.000135657 | 0.008157395 | 1.16 |
| hsa-miR-30b-3p | A_25_P00013382 | 53.38 | -4.77 | 0.000122527 | 0.007521374 | 1.25 |
| hsa-miR-1260a | A_25_P00015173 | 3216.88 | -4.77 | 0.000121602 | 0.007521374 | 1.25 |
| hsa-miR-193b-5p | A_25_P00013597 | 99.60 | -5.10 | 5.84E-05 | 0.004249593 | 1.91 |
| hsa-miR-7-1-3p | A_25_P00013295 | 61.99 | -5.11 | 5.67E-05 | 0.004180216 | 1.93 |
| hsa-miR-22-5p | A_25_P00013178 | 46.43 | -5.12 | 5.58E-05 | 0.004162807 | 1.95 |

| | | | | | | |
|---|---|---|---|---|---|---|
| hsa-miR-141-5p | A_25_P00013414 | 49.69 | -5.28 | 3.86E-05 | 0.002957185 | 2.27 |
| hsa-miR-210-5p | A_25_P00018244 | 43.92 | -5.32 | 3.53E-05 | 0.002804729 | 2.35 |
| hsa-miR-629-3p | A_25_P00010832 | 45.84 | -5.35 | 3.26E-05 | 0.002658014 | 2.42 |
| hsa-miR-4286 | A_25_P00015773 | 310.55 | -5.38 | 3.10E-05 | 0.002606483 | 2.47 |
| hsa-miR-1271-5p | A_25_P00015043 | 43.35 | -5.46 | 2.59E-05 | 0.002246106 | 2.63 |
| hsa-miR-30e-3p | A_25_P00014610 | 48.44 | -6.12 | 6.01E-06 | 0.000580445 | 3.91 |
| hsa-miR-141-5p | A_25_P00013415 | 56.19 | -7.21 | 6.22E-07 | 7.64E-05 | 5.85 |
| hsa-miR-30e-3p | A_25_P00014611 | 78.78 | -8.50 | 5.24E-08 | 9.36E-06 | 7.88 |
| hsa-miR-205-3p | A_25_P00015381 | 127.02 | -9.33 | 1.19E-08 | 2.69E-06 | 9.06 |
| hsa-miR-486-3p | A_25_P00012468 | 44.33 | -11.90 | 1.99E-10 | 8.36E-08 | 12.06 |
| hsa-miR-205-3p | A_25_P00015382 | 226.49 | -12.21 | 1.28E-10 | 6.85E-08 | 12.36 |
| hsa-miR-486-3p | A_25_P00012469 | 54.71 | -18.86 | 4.86E-14 | 1.43E-10 | 16.93 |

**Table S23. Driver genes in 8q22 and 24 regions.**

| Gene Symbol | Entrez GeneId | Genome Location | Chr Band | Name |
|---|---|---|---|---|
| RUNX1T1 | 862 | 8:91960242-92017340 | 8q22 | runt-related transcription factor 1; translocated to, 1 (cyclin D-related) |
| UBR5 | 51366 | 8:102254302-102412234 | 8q22 | ubiquitin protein ligase E3 component n-recognin 5 |
| COX6C | 1345 | 8:99878148-99892021 | 8q22-q23 | cytochrome c oxidase subunit VIc |
| EIF3E | 3646 | 8:108201885-108248702 | 8q22-q23 | eukaryotic translation initiation factor 3, subunit E |
| RAD21 | 5885 | 8:116847500-116866729 | 8q24.11 | RAD21 homolog (S. pombe) |
| EXT1 | 2131 | 8:117799712-118111046 | 8q24.11-q24.13 | multiple exostoses type 1 gene |
| MYC | 4609 | 8:127738263-127740958 | 8q24.12-q24.13 | v-myc myelocytomatosis viral oncogene homolog (avian) |
| NDRG1 | 10397 | 8:133238878-133284311 | 8q24.3 | N-myc downstream regulated 1 |
| RECQL4 | 9401 | 8:144511431-144517784 | 8q24.3 | RecQ protein-like 4 |

**Table S24. Prognostic Genes in the 8q24 Region.**

Expression of prognostic genes in the Metabric patient set. Genes were ranked by prognostic significance of their expression levels (genes with p-value < 0.0001 are shown). Amongst top ranked genes are some of the known drivers and AGO2. COSMIC database cancer drivers (Table S10) highlighted in blue. Spearman Rank Correlation between mRNA levels of genes in this region and AGO2 shown. Only patients with matched genomic (CN), transcriptomic (both coding and non-coding mRNA) and clinical follow-up data were considered.

| **Gene Symbols** | **Correlation between Amplification and Expression (rho)** | **Correlation between Amplification and Expression (p-value)** | **Cox survival outcome by Amplification (p-value)** | **Cox survival outcome by Amplification (HR)** | **Cox survival outcome by Expression (p-value)** | **Cox survival outcome by Expression (HR)** | **Correlation of gene expression with AGO2 expression (rho)** |
|---|---|---|---|---|---|---|---|
| SLC52A2 | 0.56 | 7.45E-109 | 0.04082 | 1.3 | 1.66E-08 | 3.2 | 0.60 |
| TBC1D31 | 0.50 | 3.39E-80 | 0.00031 | 1.5 | 7.33E-08 | 3.1 | 0.41 |
| AGO2 | 0.42 | 5.65E-54 | 0.01396 | 1.4 | 8.19E-08 | 3.1 | 1.00 |
| RECQL4 | 0.42 | 2.95E-57 | 0.05419 | 1.3 | 1.11E-07 | 3.0 | 0.63 |
| YWHAZ | 0.62 | 1.83E-139 | 0.00824 | 1.4 | 7.06E-07 | 2.8 | 0.44 |
| ATAD2 | 0.56 | 3.20E-105 | 0.00005 | 1.6 | 7.41E-07 | 2.8 | 0.31 |
| SQLE | 0.55 | 4.86E-104 | 0.00017 | 1.6 | 1.17E-06 | 2.7 | 0.29 |

| | | | | | | | |
|---|---|---|---|---|---|---|---|
| FBXL6 | 0.51 | 8.73E-87 | 0.04082 | 1.3 | 1.76E-06 | 2.7 | 0.64 |
| CCNE2 | 0.49 | 1.89E-80 | 0.01858 | 1.3 | 3.97E-06 | 2.6 | 0.29 |
| DERL1 | 0.65 | 1.34E-155 | 0.00018 | 1.6 | 5.41E-06 | 2.5 | 0.26 |
| POP1 | 0.35 | 1.40E-38 | 0.00578 | 1.4 | 5.69E-06 | 2.6 | 0.39 |
| LAPTM4 B | 0.42 | 2.63E-55 | 0.01943 | 1.3 | 1.77E-05 | 2.4 | 0.16 |
| FAM91A 1 | 0.54 | 2.51E-96 | 0.00036 | 1.5 | 1.80E-05 | 2.4 | 0.24 |
| EXT1 | 0.31 | 3.21E-29 | 0.00021 | 1.6 | 2.18E-05 | 2.4 | 0.51 |
| PUF60 | 0.60 | 3.06E-125 | 0.08942 | 1.2 | 2.95E-05 | 2.3 | 0.66 |
| LY6E | 0.23 | 9.50E-17 | 0.03503 | 1.3 | 3.32E-05 | 2.3 | 0.24 |
| TAF2 | 0.61 | 1.49E-125 | 0.00021 | 1.6 | 6.35E-05 | 2.3 | 0.25 |
| PLEC | 0.33 | 1.85E-34 | 0.10430 | 1.2 | 8.15E-05 | 2.2 | 0.53 |
| C8orf76 | 0.68 | 5.47E-178 | 0.00007 | 1.6 | 0.00013 | 2.2 | 0.28 |
| RAD21 | 0.57 | 4.51E-112 | 0.00042 | 1.5 | 0.00013 | 2.2 | 0.57 |
| NDRG1 | 0.36 | 2.18E-39 | 0.01499 | 1.3 | 0.00014 | 2.2 | 0.43 |
| PTP4A3 | 0.24 | 1.83E-17 | 0.00514 | 1.4 | 0.00034 | 2.1 | 0.41 |
| DSCC1 | 0.46 | 1.32E-67 | 0.00060 | 1.5 | 0.00036 | 2.0 | 0.34 |
| CPSF1 | 0.46 | 2.37E-68 | 0.04551 | 1.3 | 0.00044 | 2.0 | 0.67 |
